# Comprehensive genome-wide identification of angiosperm upstream ORFs with peptide sequences conserved in various taxonomic ranges using a novel pipeline, ESUCA

**DOI:** 10.1101/524090

**Authors:** Hiro Takahashi, Noriya Hayashi, Yui Yamashita, Satoshi Naito, Anna Takahashi, Kazuyuki Fuse, Kenji Satou, Toshinori Endo, Shoko Kojima, Hitoshi Onouchi

## Abstract

**Background:** Upstream open reading frames (uORFs) in the 5′-untranslated regions (5′-UTRs) of certain eukaryotic mRNAs encode evolutionarily conserved functional peptides, such as cis-acting regulatory peptides that control translation of downstream main ORFs (mORFs). For genome-wide searches for uORFs with conserved peptide sequences (CPuORFs), comparative genomic studies have been conducted, in which uORF sequences were compared between selected species. To increase chances of identifying CPuORFs, we previously developed an approach in which uORF sequences were compared using BLAST between *Arabidopsis* and any other plant species with available transcript sequence databases. If this approach is applied to multiple plant species belonging to phylogenetically distant clades, it is expected to further comprehensively identify CPuORFs conserved in various plant lineages, including those conserved among relatively small taxonomic groups.

**Results:** To efficiently compare uORF sequences among many species and efficiently identify CPuORFs conserved in various taxonomic lineages, we developed a novel pipeline, ESUCA. We applied ESUCA to the genomes of five angiosperm species, which belong to phylogenetically distant clades, and selected CPuORFs conserved among at least three different orders. Through these analyses, we identified 88 novel CPuORF families. As expected, ESUCA analysis of each of the five angiosperm genomes identified many CPuORFs that were not identified from ESUCA analyses of the other four species. However, unexpectedly, these CPuORFs include those conserved in wide taxonomic ranges, indicating that the approach used here is useful not only for comprehensive identification of narrowly conserved CPuORFs but also for that of widely conserved CPuORFs. Examination of the effects of 11 selected CPuORFs on mORF translation revealed that CPuORFs conserved only in relatively narrow taxonomic ranges can have sequence-dependent regulatory effects, suggesting that most of the identified CPuORFs are conserved because of functional constraints of their encoded peptides.

**Conclusions:** This study demonstrates that ESUCA is capable of efficiently identifying CPuORFs likely to be conserved because of the functional importance of their encoded peptides. Furthermore, our data show that the approach in which uORF sequences from multiple species are compared with those of many other species, using ESUCA, is highly effective in comprehensively identifying CPuORFs conserved in various taxonomic ranges.

## Background

The 5′-untranslated regions (5′-UTRs) of many eukaryotic mRNAs contain upstream open reading frames (uORFs) [1–4]. Although most uORFs are not thought to encode functional proteins or peptides, certain uORFs encode regulatory peptides that have roles in post-transcriptional regulation of gene expression [5–8]. During translation of some of these regulatory uORFs, nascent peptides act inside the ribosomal exit tunnel to cause ribosome stalling [9]. Ribosome stalling on a uORF results in translational repression of the downstream main ORF (mORF) because stalled ribosomes block the access of subsequently loaded ribosomes to the mORF start codon [10]. Additionally, if ribosome stalling occurs at the stop codon of a uORF, nonsense-mediated mRNA decay may be induced [11, 12]. In some genes, uORF-encoded nascent peptides cause ribosome stalling in response to metabolites to down-regulate mORF translation under specific cellular conditions [10, 12–17]. In contrast to the uORFs encoding cis-acting regulatory nascent peptides, a uORF in the *Medicago truncatula MtHAP2-1* gene encodes a trans-acting regulatory peptide, which binds to the 5′-UTR of *MtHAP2-1* mRNA and causes mRNA degradation [18].

To comprehensively identify uORFs that encode functional peptides, genome-wide searches for uORFs with conserved peptide sequences (CPuORFs) have been conducted using comparative genomic approaches in various organisms [19–23]. In plants, approximately 40 CPuORF families have been identified by comparing the uORF-encoded amino acid sequences of orthologous genes in some of *Arabidopsis*, rice, cotton, orange, soybean, grape and tobacco, or those of paralogous genes in *Arabidopsis* [20, 22, 23]. Recently, 29 additional CPuORF families, which include CPuORFs with non-canonical initiation codons, have been identified by comparing 5′-UTR sequences between *Arabidopsis* and 31 other plant species [24].

In conventional comparative genomic approaches, uORF sequences are compared among selected species. Therefore, homology detection depends on the selection of species for comparison. In searches using this approach, if a uORF amino acid sequence is not conserved among the selected species, this uORF is not identified as a CPuORF, even if it is evolutionarily conserved between one of the selected species and other unselected species. To overcome this problem, we previously developed the BAIUCAS (for BLAST-based algorithm for identification of uORFs with conserved amino acid sequences) pipeline [25]. In BAIUCAS, homology searches of uORF amino acid sequences are performed using BLAST between a certain species and any other species for which expressed sequence tag (EST) databases are available, and uORFs conserved beyond a certain taxonomic range are selected. Using BAIUCAS, we searched for *Arabidopsis* CPuORFs conserved beyond the order Brassicales, which *Arabidopsis* belongs to, and identified 13 novel CPuORF families [25]. We examined the sequence-dependent effects of the CPuORFs identified by BAIUCAS on mORF translation using a transient expression assay, and identified six regulatory CPuORFs that repress mORF translation in an amino acid sequence-dependent manner [26, 27]. These sequence-dependent regulatory CPuORFs include ones conserved only among relatively small taxonomic groups, such as a part of eudicots. Therefore, it is expected that sequence-dependent regulatory CPuORFs conserved in various plant lineages, including narrowly conserved ones, will be more comprehensively identified if BAIUCAS is applied to many plant species.

Before applying BAIUCAS to many species, improvement of BAIUCAS was desired to more efficiently identify CPuORFs that were conserved because of the functional importance of their encoded peptides. One major problem with identifying CPuORFs is that there are cases where a uORF found in the 5′-UTR of a transcript is fused to the mORF in an isoform of the transcript, and in some of these cases, such uORF sequences are conserved because they actually encode parts of mORF-encoded protein sequences. In other words, there are cases where the protein-coding mORF is split into multiple ORFs in a splice variant and the ORF coding for the N-terminal region of the protein appears like a uORF. Such an ORF can be extracted as a CPuORF if the amino acid sequence in the N-terminal region of the protein is evolutionarily conserved. It is difficult to distinguish between this type of ‘spurious’ CPuORFs and ‘true’ CPuORFs because even ‘true’ CPuORF-containing genes produce splice variants in which a CPuORF is fused to the mORF, as seen in the At2g31280, At5g01710, and At5g03190 genes [20, 25, 28]. Another major point to be improved is the method of calculating nonsynonymous to synonymous nucleotide substitution (*K*_a_/*K*_s_) ratios for CPuORF sequences. These *K*_a_/*K*_s_ ratios are used to evaluate whether uORF sequences are conserved at the amino acid level or at the nucleotide level. However, *K*_a_/*K*_s_ ratios largely depend on the selection of uORF sequences used for their calculation. If uORF sequences used for the calculation of a *K*_a_/*K*_s_ ratio include many sequences from closely related species, the *K*a/*K*s ratio tends to be high. For appropriate calculation of *K*_a_/*K*_s_ ratios, uORF sequences need to be selected using proper criteria.

Here, we present an improved BAIUCAS version ESUCA (for evolutionary search for uORFs with conserved amino acid sequences) and genome-wide identification of CPuORFs from five angiosperm genomes using ESUCA. To distinguish between ‘spurious’ CPuORFs conserved because they code for parts of mORF-encoded proteins and ‘true’ CPuORFs conserved because of functional constraints of their encoded small peptides, ESUCA includes an algorithm to assess whether, for each uORF, transcripts bearing a uORF-mORF fusion are minor or major forms among orthologous transcripts. Another new function of ESUCA is systematic calculation of *K*_a_/*K*_s_ ratios for CPuORF sequences. ESUCA includes an algorithm to select a uORF sequence from each order for calculation of the *K*_a_/*K*_s_ ratio of each CPuORF. Additionally, ESUCA is capable of determining the taxonomic range within which each CPuORF is conserved. Although ESUCA can identify CPuORFs conserved only among a small taxonomic group because ESUCA compares uORF sequences between a certain species and any other species with available transcript databases, CPuORFs conserved among a small taxonomic group may be less likely to encode functional peptides than those conserved across a wide taxonomic range. The automatic determination of the taxonomic range of CPuORF conservation provides useful information for the selection of CPuORFs likely to encode functional peptides. The current study demonstrates that ESUCA efficiently identifies CPuORFs likely to be conserved because of functional constraints of their encoded peptides. Furthermore, the data presented here show that the approach in which uORF sequences from multiple species are compared with those of many other species, using ESUCA, is highly effective in comprehensively identifying CPuORFs conserved in various taxonomic lineages.

## Results

### The ESUCA pipeline

In this study, to efficiently identify CPuORFs likely to be conserved because of functional importance of their encoded peptides, we developed a novel pipeline, ESUCA, which consists of a six-step procedure (Fig. 1). The first step is extraction of uORF sequences from a transcript sequence dataset. The uORFs are extracted by searching the 5′-UTR sequence of each transcript for an ATG codon and its nearest downstream in-frame stop codon. Although uORFs overlapping their downstream mORFs are also usually considered uORFs, we focus on the type of uORFs that has both the start and stop codons within the 5′-UTR to avoid including uORFs whose sequences are conserved because of functional constraints of mORF-encoded proteins. When there are splice variants of a gene, uORFs in all splice variants are extracted. The second step assesses whether, for each uORF, uORF-mORF fusion type transcripts are minor or major forms among orthologous transcripts. If transcripts with a uORF-mORF fusion are found as a major form in a majority of species with their orthologs, the uORF sequence is likely to code for a part of the mORF-encoded protein. Therefore, such a uORF should be discarded as a ‘spurious’ uORF. In contrast, if transcripts with a uORF-mORF fusion are found in only a small proportion of species with their orthologs, the uORF-mORF fusion type transcripts are considered minor form transcripts and therefore can be ignored. For this assessment, the NCBI reference sequence (RefSeq) database is used, which provides curated non-redundant transcript sequences [29]. For each uORF, the ratio of RefSeq RNAs with a uORF-mORF fusion to all RefSeq RNAs with both sequences similar to the uORF and its downstream mORF is calculated (Fig. 2). We define this ratio as the uORF-mORF fusion ratio. If the uORF-mORF fusion ratio of a uORF is equal to or greater than 0.3, then the uORF is discarded. The third step is uORF amino acid sequence homology searches. In this step, tBLASTn searches are performed against a transcript sequence database, using the amino acid sequences of the uORFs as queries (uORF-tBLASTn analysis). The uORFs with tBLASTn hits from other species are selected. The fourth step is selection of uORFs conserved among homologous genes. To confirm whether the uORF-tBLASTn hits are derived from homologs of the original uORF-containing gene, the downstream sequences of putative uORFs in the uORF-tBLASTn hits are subjected to another tBLASTn analysis, which uses the mORF amino acid sequence of the original uORF-containing transcript as a query (mORF-tBLASTn analysis) (Fig. 3). If a uORF-tBLASTn hit has a partial or intact ORF that contains a sequence similar to the mORF amino acid sequence downstream of the putative uORF, it is considered to be derived from a homolog of the original uORF-containing gene. If uORF-tBLASTn and mORF-tBLASTn hits are found in at least two orders other than that of the original uORF, then the uORF is selected as a candidate CPuORF. This is because at least three uORF sequences from different orders are necessary to confirm at the later manual validation step that the same region is conserved among homologous uORF sequences. The fifth step is *K*_a_/*K*_s_ analysis. In this step, *K*_a_/*K*_s_ ratios for the selected candidate CPuORFs is calculated to assess whether the candidate CPuORF sequences are conserved at the nucleotide or amino acid level. A *K*_a_/*K*_s_ ratio close to 1 indicates neutral evolution, whereas a *K*_a_/*K*_s_ ratio close to 0 suggests that purifying selection acted on the amino acid sequences. For each candidate CPuORF, a representative uORF-tBLASTn and mORF-tBLASTn hit is selected from each order, and the putative uORF sequences in the representative uORF-tBLASTn and mORF-tBLASTn hits are used for the calculation of the *K*_a_/*K*_s_ ratio. If the *K*_a_/*K*_s_ ratio of a candidate CPuORF is less than 0.5 and significantly different from that of the negative control with *q* less than 0.05, then the candidate CPuORF is selected for further analysis. The final step is to determine the taxonomic range of uORF sequence conservation. In this step, the representative uORF-tBLASTn and mORF-tBLASTn hits selected in the fifth step are classified into taxonomic categories (Fig. 4). On the basis of the presence of the uORF-tBLASTn and mORF-tBLASTn hits in each taxonomic category, the taxonomic range of sequence conservation is determined for each CPuORF.

**Figure 1.**
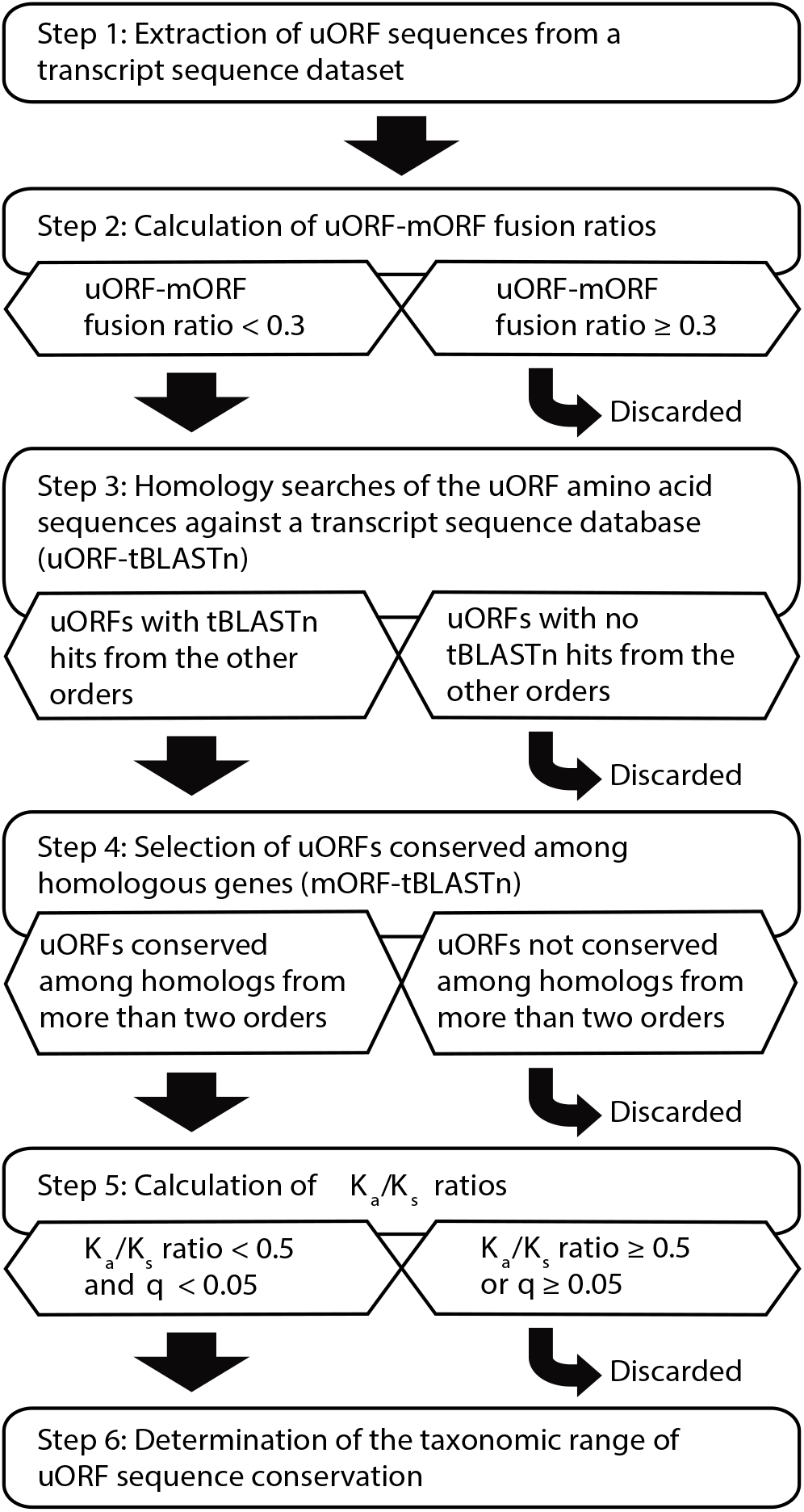
Outline of the ESUCA pipeline.

**Figure 2.**
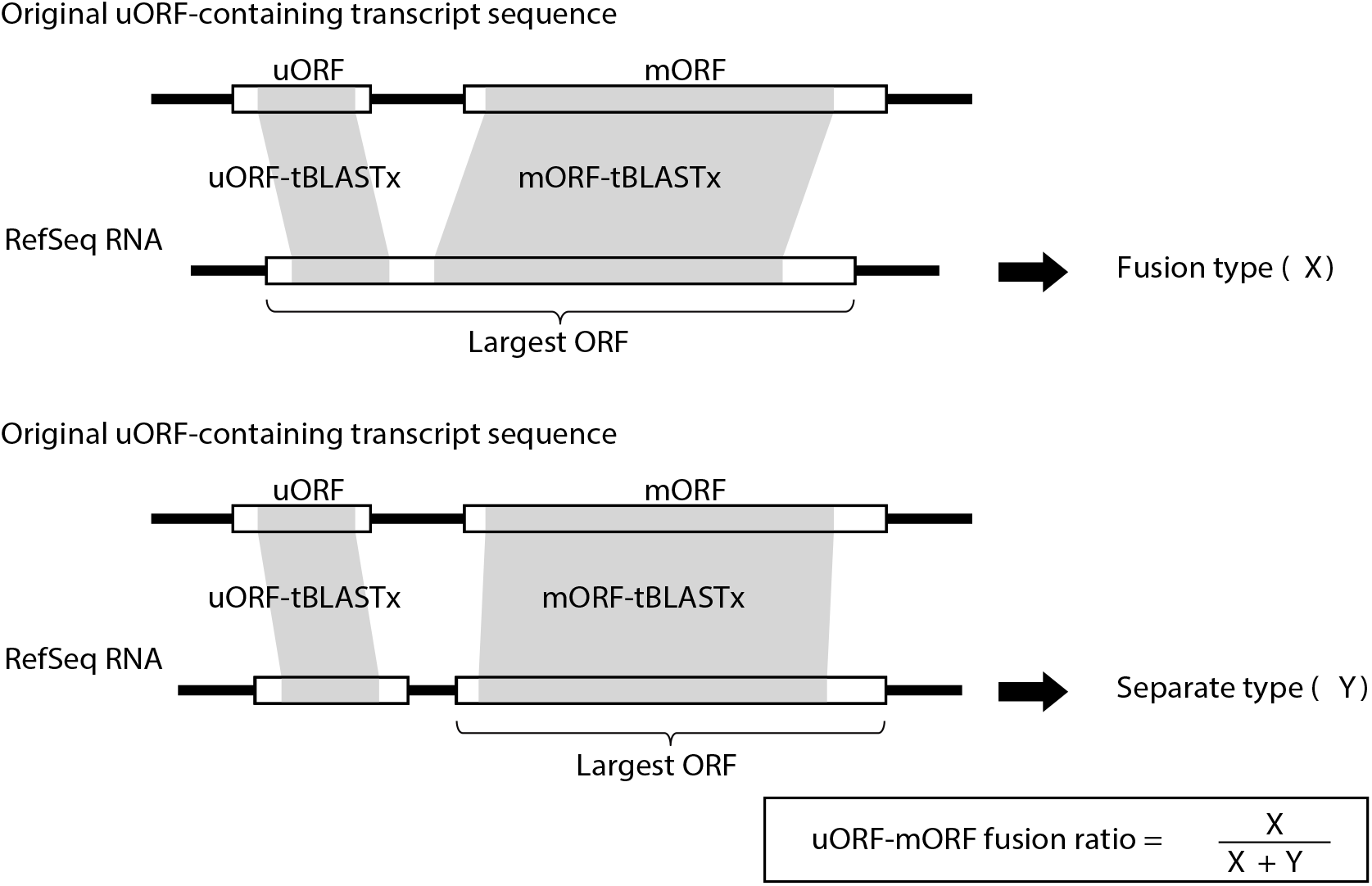
Schematic representation of the algorithm to calculate uORF-mORF fusion ratios. For each original uORF-containing transcript sequence, RefSeq RNAs are selected that match an original uORF sequence, irrespective of the reading frame, and the original mORF sequence in the same reading frame as the largest ORF of the RefSeq RNA, using tBLASTx. The shaded regions in the open boxes represent the tBLASTx-matching regions. If the uORF-tBLASTx-matching region is within the largest ORF, the RefSeq RNA is considered a uORF-mORF fusion type. The number of this type of RefSeq RNA is defined as ‘*X*’. If the uORF-tBLASTx-matching region is not within the largest ORF, the RefSeq RNA is considered a uORF-mORF separate type. The number of this type of RefSeq RNA is defined as ‘*Y*’. For each of the original uORF-containing transcripts, the uORF-mORF fusion ratio is calculated as *X* / (*X* + *Y*).

**Figure 3.**
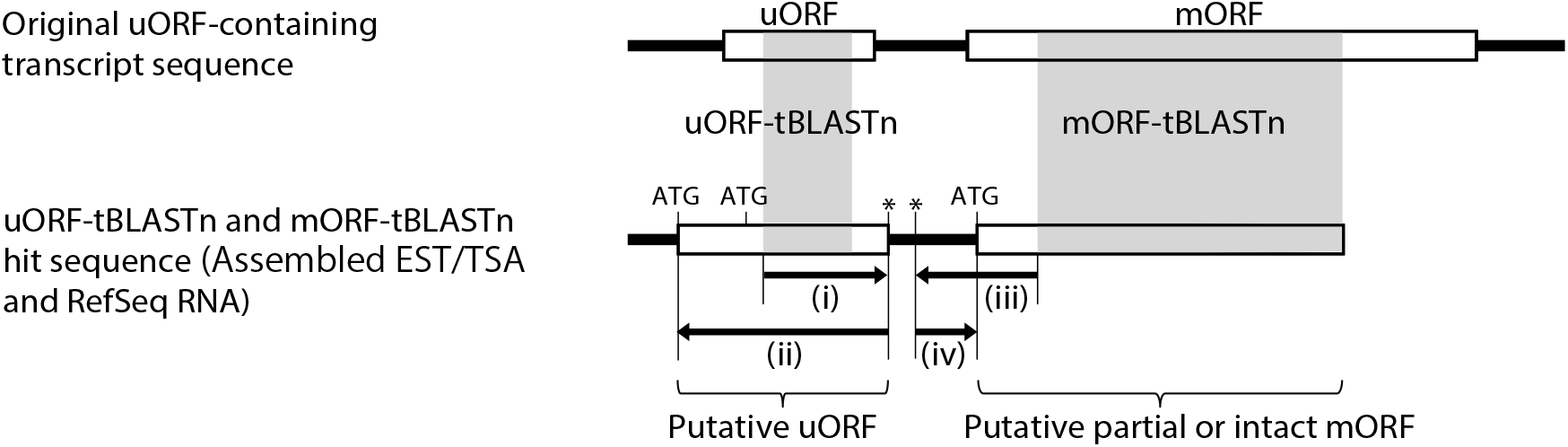
Schematic representation of BLAST-based search for uORFs conserved between homologous genes. In the third step of ESUCA, tBLASTn searches are conducted against a transcript sequence database that consists of assembled EST/TSA contigs, unclustered singleton EST/TSA sequences and RefSeq RNAs, using original uORF sequences as queries (uORF-tBLASTn). The shaded regions in the open boxes show the tBLASTn-matching regions. Asterisks represent stop codons. (i) The downstream in-frame stop codon closest to the 5′-end of the matching region of each uORF-tBLASTn hit is selected. (ii) The 5′-most in-frame ATG codon located upstream of the stop codon is selected. The ORF beginning with the selected ATG codon and ending with the selected stop codon is extracted as a putative uORF. In the fourth step of ESUCA, the downstream sequences of putative uORFs in the transcript sequences are subjected to mORF-tBLASTn analysis. Transcript sequences matching the original mORF with an *E*-value less than 10^−1^ are extracted. (iii) For each of the uORF-tBLASTn and mORF-tBLASTn hits, the upstream in-frame stop codon closest to the 5′-end of the matching region is selected. (iv) The 5′-most in-frame ATG codon located downstream of the selected stop codon is identified as the initiation codon of the putative partial or intact mORF. If the putative mORF overlaps with the putative uORF, the uORF-tBLASTn and mORF-tBLASTn hit is discarded as a uORF-mORF fusion type.

### Identification of angiosperm CPuORFs using ESUCA

We applied ESUCA to five angiosperm species, *Arabidopsis*, rice, tomato, poplar and grape, which belong to phylogenetically distant clades of angiosperm, and for which entire genomic DNA and transcript sequence datasets were available. Rice is a monocot, whereas the others are eudicots. *Arabidopsis* and poplar belong to two different groups of rosids (marvids and fabids), whereas tomato belongs to asterids. Grape belongs to neither rosids nor asterids. In the first step of ESUCA, we extracted uORF sequences from the 5′-UTR sequence of each transcript of these species, using the transcript sequence datasets described in the Materials and Methods. In these datasets, different transcript IDs are assigned to each splice variant from the same gene. To extract sequences of uORFs and their downstream mORFs from all splice variants, we extracted uORF and mORF sequences from each of the transcripts with different transcript IDs. In the second step, we calculated the uORF-mORF fusion ratio of each uORF-containing transcript, using the extracted uORF and mORF sequences, and removed uORFs with uORF-mORF fusion ratios equal to or greater than 0.3 (Supplementary Table S1). We also discarded uORFs whose numbers of RefSeq RNAs containing both sequences similar to the uORF and its downstream mORF were less than 10. This was done because appropriate evaluations of uORF-mORF fusion ratios were difficult with a few related RefSeq RNAs and such uORFs are unlikely to be evolutionarily conserved. In the third step, using the amino acid sequences of the remaining uORFs as queries, we performed uORF-tBLASTn searches against a plant transcript sequence database that contained contigs of assembled EST and transcriptome shotgun assembly (TSA), singleton EST/TSA sequences, and RefSeq RNAs (See Materials and Methods for details). In the fourth step, the uORF-tBLASTn hits were subjected to mORF-tBLASTn analysis, and uORF-tBLASTn and mORF-tBLASTn hits were extracted. Plant EST and TSA databases can include contaminant sequences from other organisms, such as parasites, plant-feeding insects and infectious microorganisms. We checked the possibility that the extracted uORF-tBLASTn and mORF-tBLASTn hits included contaminant EST/TSA sequences, using BLASTn searches. The BLASTn searches were performed using each uORF-tBLASTn and mORF-tBLASTn hit EST/TSA sequence as a query against EST/TSA and RefSeq RNA sequences from all organisms, with an *E*-value cutoff of 10^−100^ and an identity threshold of 95%. Contaminant EST/TSA sequences were identified by this analysis, as described in Materials and Methods, and were removed from the uORF-tBLASTn and mORF-tBLASTn hits. We selected uORFs whose remaining uORF-tBLASTn and mORF-tBLASTn hits were found in homologs from at least two orders other than that of the original uORF. Thereafter, we generated amino acid sequence alignments of each selected uORF and its homolog, using a putative homologous uORF sequence from each order in which uORF-tBLASTn and mORF-tBLASTn hits were found. When multiple original uORFs derived from splice variants of the same gene partially or completely shared amino acid sequences, the one with the longest conserved region was manually selected on the basis of the uORF amino acid sequence alignments. In the fifth step, the remaining uORFs were subjected to *K*_a_/*K*_s_ analysis. The uORFs with *K*_a_/*K*_s_ ratios less than 0.5 showing significant differences from those of negative controls (*q* < 0.05) were selected as candidate CPuORFs (Supplementary Table S1). Through ESUCA analyses of *Arabidopsis*, rice, tomato, poplar, and grape genomes, 104, 59, 42, 148, and 79 candidate CPuORFs were extracted, respectively. Of these, 87 *Arabidopsis*, 53 rice, 30 tomato, 77 poplar, and 44 grape uORFs belong to the previously identified CPuORF families, homology groups (HGs) 1 to 53 [20, 22–25] (Supplementary Table S1). The amino acid sequences of the remaining candidate CPuORFs are not similar to those of the known CPuORFs. Therefore, 17, 6, 12, 71, and 35 novel candidate CPuORFs were extracted from *Arabidopsis*, rice, tomato, poplar, and grape genomes, respectively.

### Validation of candidate CPuORFs

If the amino acid sequence of a uORF is evolutionarily conserved because of functional constraints of the uORF-encoded peptide, it is expected that the amino acid sequence in the functionally important region of the peptide is conserved among the uORF and its orthologous uORFs. Therefore, we manually checked whether the amino acid sequences in the same region are conserved among uORF sequences in the alignment of each novel candidate CPuORF. We found that the alignments of 13 novel candidate CPuORFs contain sequences that do not share the consensus amino acid sequence in the conserved region, and removed these sequences from the alignments. We also removed sequences derived from genes not related to the corresponding original uORF-containing gene from the alignments of five novel candidate CPuORFs. When these changes resulted in the number of orders with the uORF-tBLASTn and mORF-tBLASTn hits becoming less than two, the candidate CPuORFs were discarded. Eight novel candidate CPuORFs were discarded for this reason. Supplementary Figure S1 shows the uORF amino acid sequence alignments without the removed sequences. The *K*_a_/*K*_s_ ratios were recalculated after the manual removal of the sequences (Supplementary Table S1), and six additional novel candidate CPuORFs were discarded because their *K*_a_/*K*_s_ ratios were greater than 0.5.

Using genomic position information from Ensembl Plants (http://plants.ensembl.org/index.html) [30] and Phytozome v12.1 (https://phytozome.jgi.doe.gov/pz/portal.html) [31], we manually checked whether the positions of the remaining novel candidate CPuORFs overlap with those of the mORFs of other genes or the mORFs of splice variants of the same genes. We found that the genomic position of the candidate CPuORF of the *Arabidopsis ROA1* (AT1G60200) gene overlaps with that of an intron in the mORF region of a splice variant. Protein sequences with an N-terminal region similar to the amino acid sequence encoded by the 5′-extended region of the mORF in this splice variant are found in most orders from which the uORF-tBALASTn and mORF-tBLASTn hits of this candidate CPuORF were extracted, suggesting that the splice variant with the 5′-extended mORF is not a minor form among orthologous transcripts. Therefore, this candidate CPuORF was discarded.

In the second step of ESUCA, we excluded uORF sequences likely to encode parts of the mORF-encoded proteins, by removing uORFs with high uORF-mORF fusion ratios. To confirm that the novel candidate CPuORFs do not code for parts of the mORF-encoded proteins, each of the putative uORF sequences used for the alignment and *K*_a_/*K*_s_ analysis was queried against the UniProt protein database (https://www.uniprot.org/), using BLASTx. When putative uORF sequences matched protein sequences with low *E*-values, we manually checked whether amino acid sequences similar to those encoded by the putative uORFs were contained within mORF-encoded protein sequences. In this analysis, mORF-encoded proteins with N-terminal sequences similar to the amino acid sequences encoded by the candidate CPuORFs of the rice *OsUAM2* gene and its poplar ortholog, POPTR_0019s07850, were identified in many orders. This suggests that the sequences encoded by these candidate CPuORFs are likely to function as parts of the mORF-encoded proteins. Therefore, we discarded these candidate CPuORFs. For some other novel candidate CPuORFs, mORF-encoded proteins with sequences similar to those encoded by the candidate CPuORF and/or its homologous putative uORFs were also found. However, we did not exclude these candidate CPuORFs, because such uORF-mORF fusion type proteins were found in only a few species for each candidate.

After manual validation, 12, 4, 11, 70, and 34 uORFs were identified as novel CPuORFs in *Arabidopsis*, rice, tomato, poplar and grape, respectively. Among these novel CPuORFs, those with both similar uORF and mORF amino acid sequences were classified into the same HGs. Also, using OrthoFinder ver. 1.1.4 [32], an algorithm for ortholog group inference, we classified the genes with novel CPuORFs and those with previously identified CPuORFs into ortholog groups. The same HG number with a different sub-number was assigned to CPuORFs of genes in the same ortholog group with dissimilar uORF sequences (e.g. HG56.1 and HG56.2). Of the newly identified CPuORF genes, six were classified into the same ortholog groups as previously identified CPuORF genes, but the amino acid sequences of these six CPuORFs are dissimilar to those of the known CPuORFs. Including this type of CPuORFs, we identified 131 novel CPuORFs that belong to 88 novel HGs (HG2.2, HG9.2, HG16.2, HG43.2, HG50.2, HG52.2 and HG54-HG130) (Supplementary Table S1).

### Determination of the taxonomic range of CPuORF sequence conservation

As the final step of ESUCA, we determined the taxonomic range of the sequence conservation of each CPuORF identified, including previously identified CPuORFs. For this purpose, the uORF-tBLASTn and mORF-tBLASTn hits selected for the *K*_a_/*K*_s_ analysis and retained after manual validation were classified into 13 plant taxonomic categories (See Materials and Methods for details.), on the basis of taxonomic lineage information of EST, TSA, and RefSeq RNA sequences (Fig. 4). Fig. 5 and Supplementary Table S2 show the taxonomic range of sequence conservation for each HG and each CPuORF, respectively. In general, CPuORFs belonging to previously identified HGs tend to be conserved in a wider range of taxonomic categories than those belonging to the newly identified HGs. For 19 of the novel HGs, CPuORF sequences are conserved both in eudicots and monocots or in wider taxonomic ranges. In contrast, for 69 of the novel HGs, CPuORF sequences are conserved only among eudicots. For 12 of these, CPuORF sequences are conserved in narrower taxonomic ranges, only among rosids or asterids. These results indicate that the taxonomic range of CPuORF sequence conservation varies, and that ESUCA can identify CPuORFs conserved in a relatively narrow taxonomic range.

**Figure 4.**
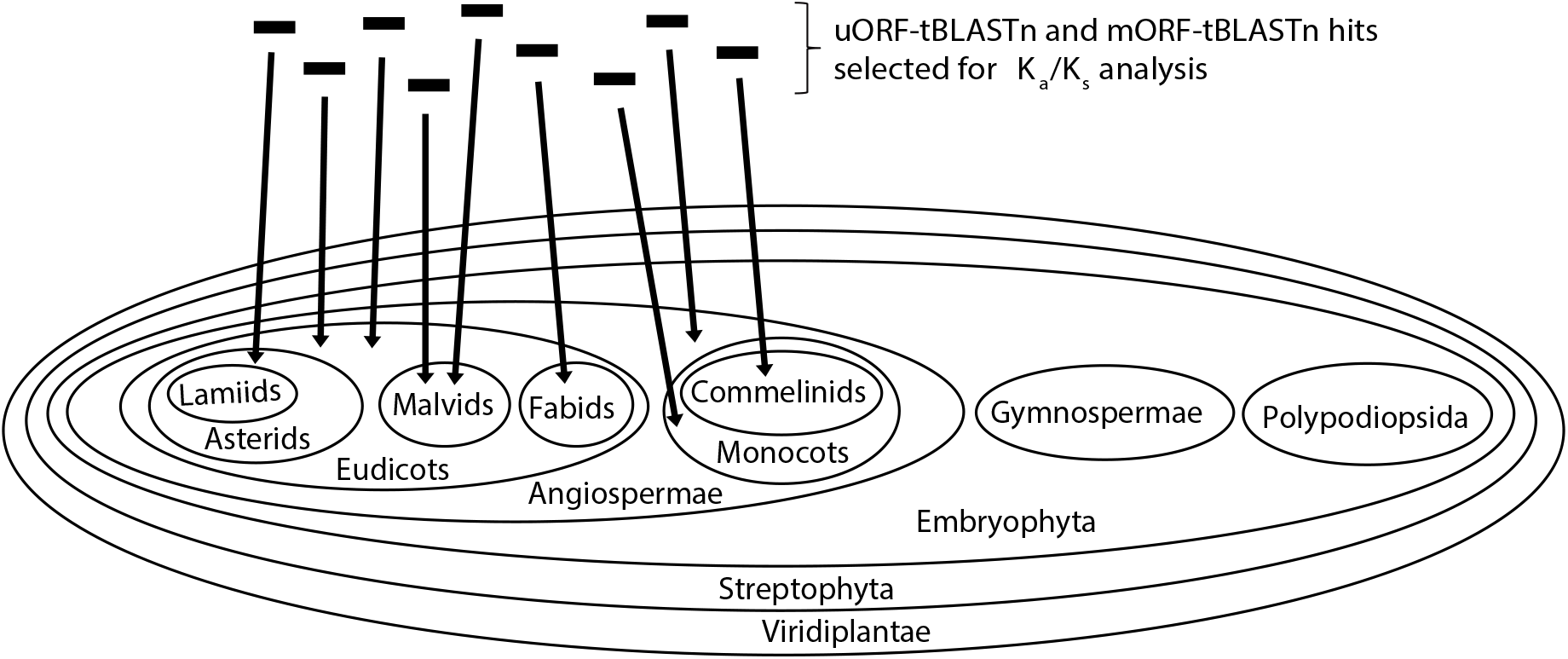
Schematic representation of the algorithm to determine the taxonomic range of uORF sequence conservation. In the fifth step of ESUCA, for *K*_a_/*K*_s_ analysis of each candidate CPuORF, a uORF-tBLASTn and mORF-tBLASTn hit sequence is selected from each order that has uORF-tBLASTn and mORF-tBLASTn hits (See Materials and Methods for the criteria for the selection). In the sixth step of ESUCA, the selected transcript sequences are classified into the 13 plant taxonomic categories to determine the taxonomic range of uORF sequence conservation, using taxonomic lineage information of EST, TSA, and RefSeq RNA sequences.

**Figure 5.**
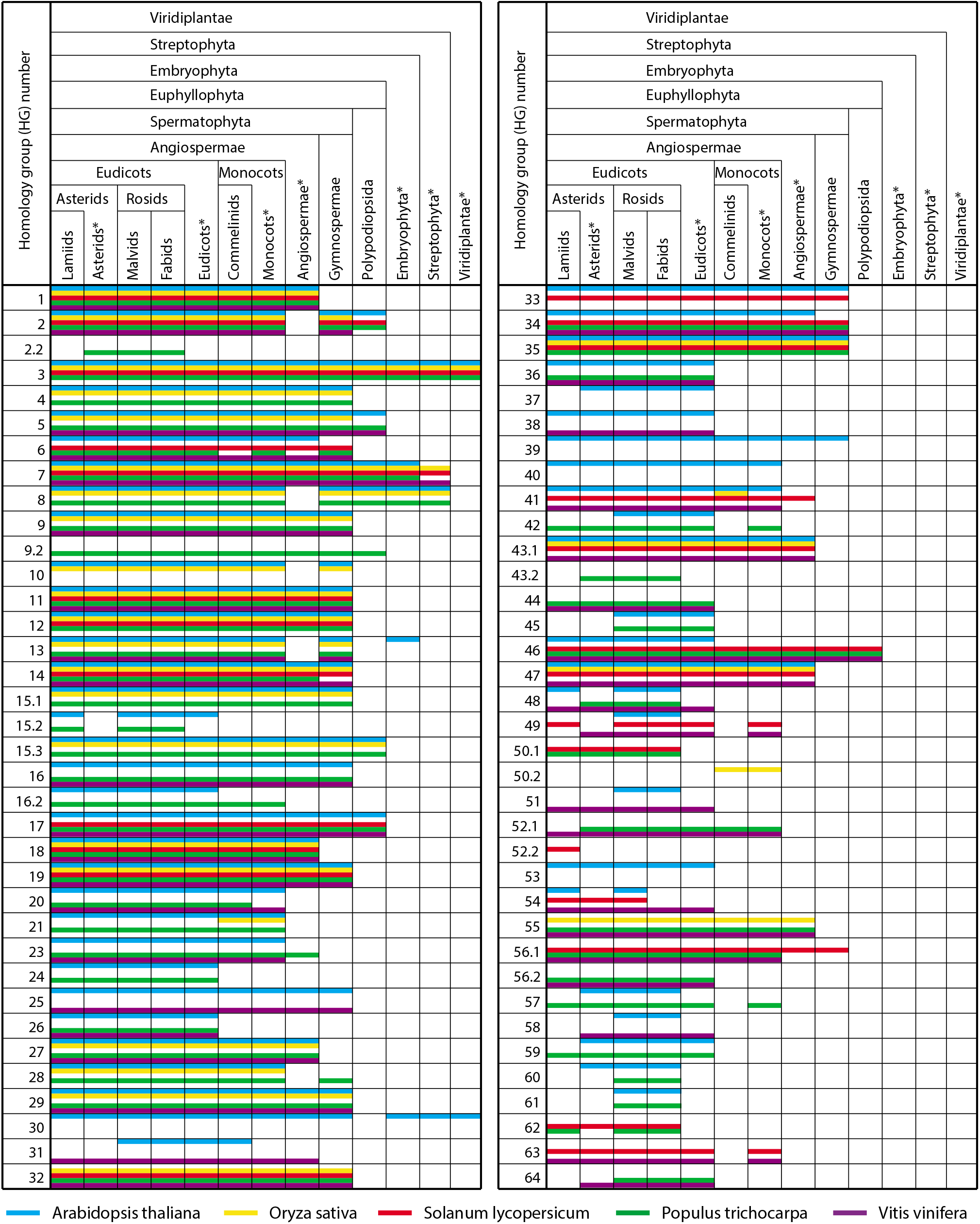
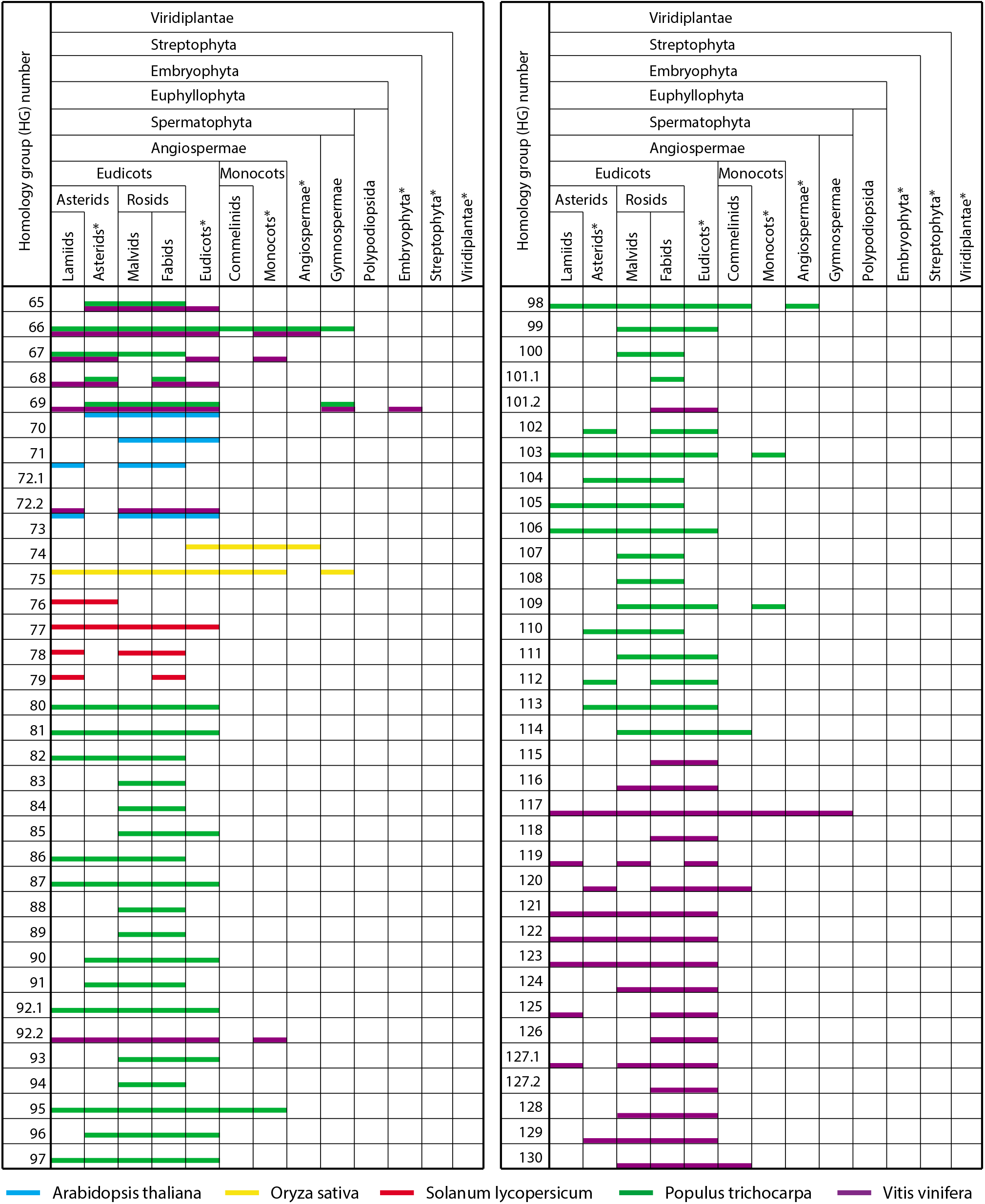
Taxonomic range of the sequence conservation of the CPuORF families. The blue, yellow, red, green, and purple lines show the conservation range of CPuORF HGs determined by applying ESUCA to *Arabidopsis* (*Arabidopsis thaliana*), rice (*Oryza sativa*), tomato (*Solanum lycopersicum*), poplar (*Populus trichocarpa*), and grape (*Vitis vinifera*) genomes, respectively. The presence of a line within a cell in each taxonomic category indicates the presence of uORF-tBLASTn and mORF-tBLASTn hits for any of the CPuORFs that belong to each HG. In taxonomic categories with a category name with an asterisk, uORF-tBLASTn and mORF-tBLASTn hits found in lower taxonomic categories were excluded. In the case where no uORF-tBLASTn and mORF-tBLASTn hit was found in the taxonomic category that contain a species from which the original uORF was derived, the line showing the species was still drawn in the cell of the taxonomic category because this category contained the species with the original uORF. *Arabidopsis*, rice, tomato, poplar and grape belong to malvids, commelinids, lamiids, fabids and eudicots*, respectively. HG1-HG53 are previously identified HGs, except for HG2.2, HG9.2, HG16.2, HG43.2, HG50.2 and HG52.2, whereas HG2.2, HG9.2, HG16.2, HG43.2, HG50.2, HG52.2 and HG54-HG130 are newly identified HGs.

### Sequence-dependent effects of CPuORFs on mORF translation

To address the relationship between the taxonomic range of CPuORF sequence conservation and the sequence-dependent effects of CPuORFs on mORF translation, we selected 11 poplar CPuORFs and examined their sequence-dependent effects on expression of the downstream reporter gene using a transient expression assay. Of the selected CPuORFs, those belonging to HG46, HG55, HG57, HG66 and HG103 are conserved in diverse angiosperms or in wider taxonomic ranges (Fig. 5, Supplementary Table S2). The CPuORFs belonging to HG65, HG80, HG81 and HG87 are conserved in a wide range of eudicots, whereas the CPuORFs belonging to HG88 and HG107 are conserved only among rosids (Fig. 5, Supplementary Table S2). In the 5′-UTR of the poplar gene with the HG107 CPuORF, there is another uORF immediately upstream of the CPuORF (Supplementary Table S2K). To focus on the sequence-dependent effect of the CPuORF on mORF translation, the upstream uORF was eliminated by mutating its initiation codon because the presence of the immediate upstream uORF may reduce the translation efficiency of the CPuORF and therefore potentially make the effect of the CPuORF ambiguous. The 5′-UTR sequences containing the selected CPuORFs were fused to the firefly luciferase (Fluc) coding sequence and were placed under the control of the 35S promoter to generate the wild-type (WT) reporter constructs (Fig. 6a, Supplementary Figure S2). To assess the importance of the amino acid sequences for the effects of these CPuORFs on mORF translation, frameshift mutations were introduced into the CPuORFs so that the amino acid sequences of their conserved regions could be altered. A + 1 or - 1 frameshift was introduced upstream or within the conserved region of each CPuORF, and another frameshift was introduced before the stop codon to shift the reading frame back to the original frame (Supplementary Figure S2). These reporter constructs were each transfected into protoplasts from *Arabidopsis thaliana* MM2d suspension-cultured cells. After 24 h of incubation, cells were harvested and disrupted to analyze luciferase activity. In five of the 11 CPuORFs, the introduced frameshift mutations significantly upregulated the expression of the reporter gene, indicating that these CPuORFs repress mORF translation in a sequence-dependent manner (Fig. 6b). These five novel sequence-dependent regulatory CPuORF include the HG107 CPuORF, which is one of the CPuORFs conserved only among rosids. Therefore, this result suggests that CPuORFs conserved only among rosids can have sequence-dependent regulatory effects.

**Figure 6.**
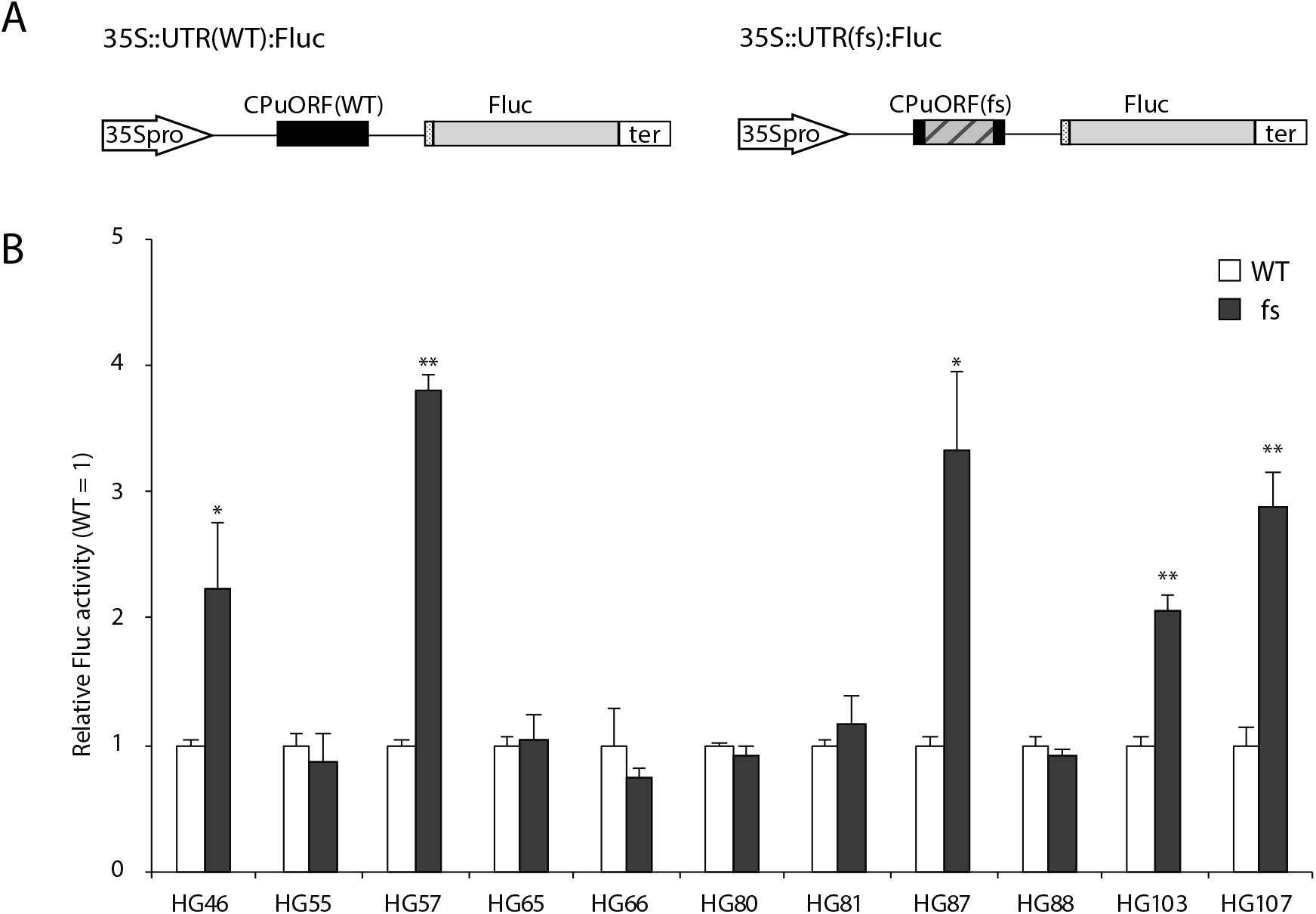
Sequence-dependent effects of novel CPuORFs on main ORF translation. (A) Schematic representation of the WT (*35S::UTR(WT):Fluc*) and frameshift mutant (*35S::UTR(fs):Fluc*) reporter constructs. The 5′-UTR sequence containing each of the 11 poplar CPuORFs was inserted between the 35S promoter (*35Spro*) and the *Fluc* coding sequence. The hatched box in the frameshift (fs) mutant CPuORF indicates the frame-shifted region. The dotted boxes represent the first five nucleotides of the mORF associated with each of the 11 poplar CPuORFs. See Supplementary Figure S2 for the exact position and length of each CPuORF and the exact frame-shifted region. The polyadenylation signal of the *A. thaliana HSP18.2* gene is designated as ‘ter’. (B) Transient expression assay. Each reporter plasmid containing a WT or fs mutant CPuORF, belonging to an indicated HG, was co-transfected into MM2d protoplasts with the *35S::Rluc* internal control plasmid by PEG treatment. After incubation for 24 h, cells were collected and disrupted for dual luciferase assay. Fluc activity was normalized to Rluc activity, and the activity relative to that of the corresponding WT reporter construct was calculated. Means ± SD of three biological replicates are shown. Single and double asterisks indicate significant differences between the WT and fs constructs at *p* < 0.05 and *p* < 0.01, respectively, as determined by Student’s *t*-test. Representative results of two experiments using independently prepared protoplasts are shown.

## Discussion

### Comprehensive identification of angiosperm CPuORFs by ESUCA analyses of multiple species’ genomes

In this study, we developed ESUCA, a pipeline for efficient genome-wide identification of CPuORFs. By applying ESUCA to five angiosperm genomes, we identified 131 CPuORFs that belong to 88 novel HGs. Of these HGs, 70 were identified through ESUCA analysis of only one of the *Arabidopsis*, rice, tomato, poplar or grape genomes (Fig. 5). This means that CPuORFs belonging to these HGs cannot be identified by comparing uORF sequences of orthologous genes between these five species. Therefore, this result demonstrates that the approach used in this study, in which uORF sequences from multiple species are compared with those of many other species, is highly effective in comprehensively identifying CPuORFs. We expected this approach to be particularly useful for comprehensive identification of CPuORFs conserved in relatively narrow taxonomic ranges. Therefore, we used the five angiosperm species belonging to relatively distant lineages in order to identify CPuORFs conserved only among the taxonomic categories to which each of the five species belong. However, unexpectedly, in 41 of 70 HGs identified through ESUCA analysis of only one of the five plant genomes, CPuORF sequences are conserved in both rosids and asterids, two major groups of eudicots, indicating that these CPuORFs are conserved across diverse eudicots (Fig. 5). One possible explanation of this observation is that sequence conservation of uORFs is often lost in small taxonomic groups, such as orders or families, during evolution. For example, while the CPuORF of the tomato LOC101264451 gene, which belongs to HG43.1, exerts a sequence-dependent repressive effect on mORF translation, the CPuORF of its *Arabidopsis* ortholog, *ANAC096*, lacks the C-terminal half of the amino acid sequence in the highly conserved region and does not have a sequence-dependent regulatory effect [25–27]. In contrast to the *Arabidopsis ANAC096* CPuORF, all the critical amino acid residues in the highly conserved region are retained in the HG43.1 CPuORF of *Tarenaya hassleriana*, which belongs to the same order as *Arabidopsis*, Brassicales, but a different family. CPuORFs involved in preferable but not essential post-transcriptional regulations may be lost in some taxonomic clades during evolution. Such CPuORFs would be difficult to be identified by comparing uORF sequences between a few selected species if a selected species is included in clades where the CPuORF sequences were lost. Therefore, the ESUCA method is advantageous for comprehensive identification of CPuORFs compared with conventional comparative genomic approaches. Our results reveal that the approach used here (*i.e*. ESUCA analyses of multiple species’ genomes) is further advantageous and is highly useful not only for comprehensive identification of narrowly conserved CPuORFs but also for that of widely conserved CPuORFs.

The transient expression assays in the current study identified five novel poplar regulatory CPuORFs that exert sequence-dependent repressive effects on mORF translation. One of the identified regulatory CPuORFs is conserved only among rosids (*i.e*. only among fabids and malvids). This result suggests that at least, CPuORFs conserved only among fabids and malvids can have sequence-dependent regulatory effects, although we cannot rule out the possibility that CPuORFs conserved in narrower taxonomic ranges can have sequence-dependent regulatory effects. Of the 92 CPuORF HGs identified through ESUCA analysis of the poplar genome, HG101.1 is conserved only among fabids, whereas the others are conserved in at least one other taxonomic category in addition to fabids (Fig. 5). Likewise, all the HGs identified through ESUCA analysis of the other four plant genomes are conserved in multiple taxonomic categories. Altogether, these results suggest that most the CPuORFs identified in this study, in which CPuORFs conserved in more than two orders were extracted, are likely to be conserved because of functional constraints of their encoded peptides. Of the 11 poplar CPuORFs analyzed by the transient expression assays, five are conserved beyond eudicots, and three of them exhibited sequence-dependent repressive effects (Figs. 5 and 6b). In addition, we previously examined the effects of 16 CPuORFs, which belong to HG27-HG29, HG33-HG43.1, HG44, and HG45, on mORF translation and identified six sequence-dependent regulatory CPuORFs. In this analysis, five of 11 CPuORFs conserved beyond eudicots showed sequence-dependent repressive effects, whereas one of five CPuORFs conserved only among eudicots showed a sequence-dependent repressive effect [26, 27]. These results may suggest that CPuORFs conserved beyond eudicots are more likely to encode functional peptides than CPuORFs conserved only among eudicots. ESUCA is capable of selecting CPuORFs conserved in certain taxonomic ranges on the basis of two criteria, the numbers of orders in which CPuORFs are conserved and/or taxonomic categories in which CPuORFs are conserved. Our results demonstrate that this function of ESUCA is highly useful for the efficient selection of CPuORFs likely to encode functional peptides.

Of the CPuORFs analyzed for their sequence-dependent regulatory effects in the present study, the CPuORFs belonging to HG55 and HG66 showed no significant sequence-dependent effect on mORF translation, despite their widespread sequence conservation beyond eudicots (Figs. 5 and 6b). These CPuORFs might encode peptides that have functions other than the control of mORF translation, or they might exert sequence-dependent regulatory effects only under certain conditions. In fact, many known sequence-dependent regulatory uORFs repress mORF translation in response to metabolites, such as polyamine, arginine, and sucrose [10, 14, 15, 17]. Likewise, the other CPuORFs that exhibited no significant sequence-dependent regulatory effect might encode peptides that have other functions or exert regulatory effects only under specific conditions.

### Filtering using the uORF-mORF fusion ratio

To distinguish between ‘spurious’ CPuORFs conserved because they code for parts of mORF-encoded proteins and ‘true’ CPuORFs conserved because of functional constraints of their encoded small peptides, we employed the criterion of the uORF-mORF fusion ratio and discarded uORFs with uORF-mORF fusion ratios equal to or greater than 0.3. We checked how effectively the ‘spurious’ CPuORFs were removed with this criterion, using a protein sequence database. Although uORF-mORF fusion type protein sequences were found in OsUAM2 homologs from widespread angiosperm species, no other candidate CPuORFs were extracted in which uORF-mORF fusion type protein sequences were found in many species that have sequences similar to the candidate CPuORFs. This indicates that the uORF-mORF fusion ratio filtering worked effectively to exclude ‘spurious’ CPuORFs that code for parts of the mORF-encoded proteins.

ESUCA extracted all known plant cis-acting regulatory CPuORFs that control mORF translation in a sequence-dependent manner (*i.e*. HG1, HG3, HG6, HG12, HG13, HG14, HG15.3, HG18, HG19, HG27, HG34, HG36, HG41, HG42, and HG43.1 CPuORFs [15, 17, 26, 27, 33–38]) (Supplementary Table S1). Of these, the HG3 CPuORFs, associated with an mORF coding for an *S*-adenosylmethionine decarboxylase (AdoMetDC; EC 4.1.1.50) [20, 39], showed relatively high uORF-mORF fusion ratios, although still below 0.3 (Supplementary Table S1). In this study, 11 CPuORFs belonging to this HG were extracted using ESUCA and the uORF-mORF fusion ratios of these CPuORFs were in the range of 0.17 to 0.27, with a median value of 0.26 (Supplementary Table S1). Consistent with these relatively high uORF-mORF fusion ratios, uORF-mORF fusion type protein sequences that contain amino acid sequences resembling HG3 CPuORFs are found in widespread angiosperm species. The uORF-mORF fusion ratios of the candidate CPuORFs of rice *OsUAM2* and its poplar ortholog were 0.28 (Supplementary Table S1). The rice *OsUAM2* gene codes for a protein similar to UDP-arabinopyranose mutases (EC 5.4.99.30) [40]. Two protein isoforms, a uORF-mORF fusion type and one lacking the uORF-encoded region, produced from splice variants of an *OsUAM2* orthologous gene, are found in diverse angiosperm species. The functions of both of isoforms are not yet known. Considering the example of HG3 CPuORFs, we cannot rule out the possibility that a candidate CPuORF functions as a regulatory uORF even if uORF-mORF fusion type protein sequences are found in widespread species. However, to avoid including potential ‘spurious’ CPuORFs whose amino acid sequences are likely to be evolutionarily conserved because of their function as N-terminal regions of the mORF-encoded proteins, we excluded the candidate CPuORFs of the rice *OsUAM2* gene and its poplar ortholog. In conclusion, the criterion of the uORF-mORF fusion ratio used in this study appears appropriate because all known cis-acting sequence-dependent regulatory CPuORFs were extracted and most ‘spurious’ CPuORFs were removed.

## Conclusion

The present study demonstrates that the approach in which uORF sequences from multiple species are compared with those of many other species, using ESUCA, is highly effective in comprehensive identification of CPuORFs. Using this approach, we identified many novel angiosperm CPuORFs, which include CPuORFs conserved among limited clades and widely conserved CPuORFs. Our results also showed that ESUCA is capable of efficiently selecting CPuORFs likely to be conserved because of functional importance of their encoded peptides. The approach used here can be applied to any eukaryotic organism with available genome and transcript sequence databases and therefore is expected to contribute to the comprehensive identification of CPuORFs encoding functional peptides in various organisms. Furthermore, besides CPuORFs, the algorithms developed for ESUCA and the approach used here can be applied to the identification of other sequences conserved in various taxonomic ranges.

## Materials and Methods

### Extraction of uORF sequences

We used genome sequence files in FASTA format and genomic coordinate files in GFF3 format obtained from Ensemble Plants Release 33 (https://plants.ensembl.org/index.html) [30] to extract *Arabidopsis* (*Arabidopsis thaliana*), tomato (*Solanum lycopersicum*), poplar (*Populus trichocarpa*), and grape (*Vitis vinifera*) uORF sequences. We used a genome sequence file in FASTA format and a genomic coordinate files in GFF3 format obtained from Phytozome v11 (https://phytozome.jgi.doe.gov/pz/portal.html) [31] for rice (*Oryza sativa*). We extracted exon sequences from genome sequences, on the basis of genomic coordinate information, and constructed transcript sequence datasets by combining exon sequences. On the basis of the transcription start site and the translation initiation codon of each transcript in the genomic coordinate files, we extracted 5′-UTR sequences from the transcript sequence datasets. Then, we searched the 5′-UTR sequences for an ATG codon and its nearest downstream in-frame stop codon. Sequences starting with an ATG codon and ending with the nearest in-frame stop codon were extracted as uORF sequences. When multiple uORFs from a gene shared the same stop codon, only the longest uORF sequence was used for further analyses.

### Assembly of EST and TSA sequences

EST, TSA, and RefSeq RNA sequence datasets were obtained from the International Nucleotide Sequence Database Collaboration databases (NCBI release of 2016-12-03 and DDBJ release 106.0 for EST and TSA sequences, NCBI release 79 for RefSeq RNA sequences). On the basis of taxonomic lineage information provided by NCBI Taxonomy (https://www.ncbi.nlm.nih.gov/taxonomy), EST and TSA sequences derived from Viridiplantae were extracted from the databases. EST and TSA sequences from the same species were assembled using Velvet ver. 1.2.10 [41] with k-mer length of 99. The k-mer length was optimized using *A. thaliana* EST and TSA sequences to minimize the total numbers of assembled contigs and unclustered singleton sequences. Unclustered singleton EST/TSA sequences, derived from species for which RefSeq RNA sequences were available, were mapped to the RefSeq RNA sequences using Bowtie2 ver. 2.2.9 [42] and the default parameters. We discarded singleton EST/TSA sequences that matched any RefSeq RNA sequences from the same species. We created a plant transcript sequence database for BLAST searches in our local computers by using the remaining singleton EST/TSA sequences, assembled contigs, Viridiplantae RefSeq RNA sequences, and makeblastdb, a program contained in the NCBI-BLAST package.

### Calculation of the uORF-mORF fusion ratio

The uORF-mORF fusion ratio for each of the extracted uORFs was assessed as follows. We performed tBLASTx using each uORF sequence as a query against plant (Viridiplantae) RefSeq RNA sequences with an *E*-value cutoff of 2000 (uORF-tBLASTx). We used standalone NCBI-BLAST + ver. 2.6.0 [43] for all BLAST analyses. We next performed tBLASTx with an *E*-value threshold of 10^−1^ using the mORF sequence associated with each uORF as a query against the uORF-tBLASTx hit sequences (mORF-tBLASTx). Using these two-step tBLASTx searches, we selected RefSeq RNAs that contain both sequences similar to the original uORF and its downstream mORF. Then, we examined whether the largest ORF of each of the selected RefSeq RNAs included the region that matched the original mORF in the same reading frame (Fig. 2). We also examined whether the largest ORF included the region that matched the original uORF, irrespective of the reading frame. The RefSeq RNA was considered to have a uORF-mORF fusion if the largest ORF contained both regions that matched the original uORF and mORF (Fig. 2). The RefSeq RNA was considered to have a uORF separated from the downstream mORF if the largest ORF contained the region that matched the original mORF but not the region that matched the original uORF (Fig. 2). RefSeq RNA numbers of the former and latter types were defined as *X* and *Y*, respectively. We calculated a uORF-mORF fusion ratio as *X* / (*X* + *Y*) for each of the original uORF-containing transcript sequences.

### BLAST-based search for uORFs conserved between homologous genes

To search for uORFs with amino acid sequences conserved between homologous genes, we first performed tBLASTn searches against the assembled plant transcript sequence database, using the amino acid sequences of the uORFs as queries. In these uORF-tBLASTn searches, we extracted transcript sequences that matched a uORF with an *E*-value less than 2000 and derived from species other than that of the original uORF. The downstream in-frame stop codon closest to the 5′-end of the matching region of each uORF-tBLASTn hit was selected (Fig. 3). Then, we looked for an in-frame ATG codon upstream of the selected stop codon, without any other in-frame stop codon between them. uORF-tBLASTn hits without such an ATG codon were discarded. If one or more in-frame ATG codons were identified, the 5′-most ATG codon was selected. The ORF beginning with the selected ATG codon and ending with the selected stop codon was extracted as a putative uORF (Fig. 3). The downstream sequences of putative uORFs were subjected to another tBLASTn analysis to examine whether the transcripts were derived from homologs of the original uORF-containing gene. In this analysis, the amino acid sequence of the mORF associated with the original uORF was used as a query sequence, and transcript sequences matching the mORF with an *E*-value less than 10^−1^ were extracted. For each of the uORF-tBLASTn and mORF-tBLASTn hits, the upstream in-frame stop codon closest to the 5′-end of the region matching the original mORF was selected, and the 5′-most in-frame ATG codon located downstream of the selected stop codon was identified as the putative mORF initiation codon (Fig. 3). uORF-tBLASTn and mORF-tBLASTn hits were discarded as uORF-mORF fusion type sequences if the putative mORF overlapped with the putative uORF. The original uORF was selected as a candidate CPuORF if the remaining uORF-tBLASTn and mORF-tBLASTn hits belonged to at least two orders other than that from which the original uORF was derived.

### Identification of contaminant ESTs and TSAs

In the uORF-tBLASTn and mORF-tBLASTn analyses described above, we excluded tBLASTn hit ESTs and TSAs derived from contaminated organisms. To examine whether each uORF-tBLASTn and mORF-tBLASTn hit sequence is derived from contaminated organisms, we performed BLASTn searches against EST, TSA, and RefSeq RNA sequences from all organisms except for those of metagenomes, using each uORF-tBLASTn and mORF-tBLASTn hit EST/TSA sequence as a query. If a uORF-tBLASTn hit EST/TSA sequence matched an EST, TSA, or RefSeq RNA sequence of a different order from the species of the uORF-tBLASTn and mORF-tBLASTn hit, with an *E*-value less than 10^−100^ and an identity equal to or greater than 95%, it was considered a candidate contaminant sequence. In this case, either the uORF-tBLASTn and mORF-tBLASTn hit or the BLASTn hit may be a contaminant sequence. To distinguish these possibilities, we compared the ratio of the BLASTn hit number to the total EST/TSA and RefSeq RNA sequence number between the species of each uORF-tBLASTn and mORF-tBLASTn hit and the species of its BLASTn hits. Appropriate comparisons are difficult unless species used for this comparison have enough number of EST/TSA and RefSeq RNA sequences. Therefore, if the total EST/TSA and RefSeq RNA sequence number of a species is less than 5000, BLASTn hits derived from the species were not used for this analysis. If the ratio of the BLASTn hit number to the total EST/TSA and RefSeq RNA sequence number of a uORF-tBLASTn and mORF-tBLASTn hit species is less than that of any other BLASTn hit species, the uORF-tBLASTn and mORF-tBLASTn hit sequence was identified as a contaminant sequence.

### *K*_a_/*K*_s_ analysis

For *K*_a_/*K*_s_ analysis of each candidate CPuORF, a putative uORF sequence was selected from each order in which uORF-tBLASTn and mORF-tBLASTn hits were found, using the following criteria. First, we selected transcript sequences that matched the mORF associated with the candidate CPuORF with an *E*-value less than 10^−20^ in mORF-tBLASTn analysis. Of these sequences, we then selected the one with the smallest geometric means of mORF-tBLASTn and uORF-tBLASTn *E*-values. When mORF-tBLASTn *E*-values of all uORF-tBLASTn and mORF-tBLASTn hits in an order were equal to or greater than 10^−20^, we selected the transcript sequence with the smallest geometric means of mORF-tBLASTn and uORF-tBLASTn *E*-values in the order. Putative uORF sequences in the selected transcript sequences were used for generation of uORF amino acid sequence alignments and for *K*_a_/*K*_s_ analysis. Multiple alignments were generated by using standalone Clustal Omega (ClustalO) ver. 1.2.2 [44]. For each candidate CPuORF, a median *K*_a_/*K*_s_ ratio for all pairwise combinations of the original uORF and its homologous putative uORFs was calculated using the LWL85 algorithm [45] in the seqinR package [46].

For statistical tests of *K*_a_/*K*_s_ ratios, we calculated the distribution of mutation rates between the original uORF and its homologous putative uORFs and those between the original uORF and its artificially generated mutants, using the observed mutation rate distribution. Then, observed empirical *K*_a_/*K*_s_ ratio distributions were compared with null distributions (negative controls) using the Mann-Whitney *U* test to validate statistical significance. The one-sided *U* test was used to investigate whether the observed distributions were significantly lower than the null distributions. Adjustment for multiple comparisons was achieved by controlling the false discovery rate using the Benjamini and Hochberg procedure [47].

### Determination of the taxonomic range of uORF sequence conservation

To automatically determine the taxonomic range of the sequence conservation of each CPuORF, we first defined 13 plant taxonomic categories. The 13 defined taxonomic categories are lamiids, asterids other than lamiids, mavids, fabids, eudicots other than rosids and asterids, commelinids, monocots other than commelinids, Angiospermae other than eudicots and monocots, Gymnospermae, Polypodiopsida, Embryophyta other than Euphyllophyta, Streptophyta other than Embryophyta, and Viridiplantae other than Streptophyta. On the basis of taxonomic lineage information of EST, TSA, and RefSeq RNA sequences, which were provided by NCBI Taxonomy, the uORF-tBLASTn and mORF-tBLASTn hit sequences selected for *K*_a_/*K*_s_ analysis were classified into the 13 taxonomic categories (Fig. 4). It should be noted that, in NCBI taxonomy, eudicots, Angiospermae, and Ggymnospermae are referred to as eudicotyledons, Magnoliophyta, and Acrogymnospermae, respectively. For each CPuORF, the numbers of transcript sequences classified into each category were counted and shown in Supplementary Table S2. These numbers represent the numbers of orders in which the amino acid sequence of each CPuORF is conserved.

### Statistical and informatic analyses

All programs, except for existing stand-alone programs, such as BLAST [43], ClustalO [44] and Jalview [48], were written in R (www.r-project.org). We also used R libraries, GenomicRanges [49], exactRankTests, Biostrings and seqinr [46].

### Plasmid construction

Plasmid pNH006 harbors the cauliflower mosaic virus 35S RNA (35S) promoter, the Fluc coding sequence, and the polyadenylation signal of the *A. thaliana HSP18.2* gene in pUC19. To construct this plasmid, pMT61 [26] was digested with *Sal*I and *Sac*I, and the *Sal*I-*Sac*I fragment containing the Fluc coding sequence was ligated into the *Sal*I and *Sac*I sites between the 35S promoter and the *HSP18.2* polyadenylation signal of plasmid pKM56 [38]. To generate reporter plasmids pNH92-pNH101 (Supplementary Table S3) for transient expression assays, the 5′-UTR sequences of ten poplar genes were amplified by PCR from poplar (*Populus nigra*) full-length cDNA clones pds25559, pds10965, pds14390, pds12940, pds13862, pds15817, pds28294, pds26157, pds14623 and pds23234 (Supplementary Table S4), obtained from RIKEN [50]. Primer sets used are shown in Supplementary Tables S3 and S5. To construct pNH92, pNH94-pNH98 and pNH100-pNH101, amplified fragments containing the 5′-UTR sequences were digested with *Xba*I and *Sal*I and ligated between the *Xba*I and *Sal*I sites of pNH006. To create pNH93 and pNH99, amplified fragments containing the 5′-UTR sequences were inserted between the *Xba*I and *Sal*I sites of pNH006 using the SLiCE method [51]. To make pHN102, the 5′-UTR sequence of the *Populus trichocarpa* POPTR_0013s08000 gene with a mutation at the initiation codon of the uORF located immediately upstream of the HG107 CPuORF was synthesized by Fasmac (Atsugi, Japan) on the basis of NCBI RefSeq accession no. XM_002319213.3. The synthesized 5′-UTR sequence was amplified by PCR using primers 35S_*Xba*I_SLiCE-F and FLUC_*Sal*I_SLiCE-R (Supplementary Table S5) and were inserted between the *Xba*I and *Sal*I sites of pNH006 using the SLiCE method [51]. Frameshift mutations were introduced into each CPuORF using overlap extension PCR [52], with primers listed in Supplementary Tables S4 and S5, to yield pNH103-pNH112. Sequence analysis confirmed the integrity of the PCR-amplified regions of all constructs.

### Transient expression assay

Transient expression assays were performed as described in Hayashi *et al*. 2017 [38]. Protoplasts from *A. thaliana* MM2d suspension cells [53] were used, as were the reporter plasmids described above and the pKM5 [38] internal control plasmid. pKM5 contains the 35S promoter, the *Renilla* luciferase (Rluc) coding sequence, and the *NOS* polyadenylation signal in pUC19. For each experiment, 5 μg each of a reporter plasmid and pKM5 were transfected into protoplasts.

## Declarations

### Authors’ contributions

HT, HO, SK, KS, and SN conceived and designed the experiments. NH and YY performed the experiments. HT, HO, TE, and AT analyzed the data. HT, NH, YY, AT, and KF contributed to reagents/materials/analysis tools. HO and HT wrote the paper. All authors read and approved the final manuscript.

### Ethics approval and consent to participate

Not applicable.

### Consent for publication

Not applicable.

### Availability of data and materials

Not applicable.

## Acknowledgements

We thank Ms. Kazuko Harada for general assistance.

## List of Abbreviations

35S: cauliflower mosaic virus 35S RNA;
5′-UTR: 5′-untranslated region;
CPuORF: conserved peptide upstream open reading frame;
EST: expressed sequence tag;
Fluc: firefly luciferase;
HG: homology group;
mORF: main open reading frame;
RefSeq: NCBI reference sequence;
Rluc: Renilla luciferase;
TSA: transcriptome shotgun assembly;
uORF: upstream open reading frame

## References

1. Churbanov A, Rogozin IB, Babenko VN, Ali H, Koonin EV. Evolutionary conservation suggests a regulatory function of AUG triplets in 5’-UTRs of eukaryotic genes. Nucleic Acids Res. 2005; 33:5512–5520.

2. Galagan JE, Calvo SE, Cuomo C, Ma LJ, Wortman JR, Batzoglou S, et al. Sequencing of *Aspergillus nidulans* and comparative analysis with *A. fumigatus* and *A. oryzae*. Nature 2005; 438:1105–1115.

3. Kawaguchi R, Bailey-Serres J. mRNA sequence features that contribute to translational regulation in *Arabidopsis*. Nucleic Acids Res. 2005; 33:955–965.

4. Rogozin IB, Kochetov AV, Kondrashov FA, Koonin EV, Milanesi L. Presence of ATG triplets in 5’ untranslated regions of eukaryotic cDNAs correlates with a ‘weak’ context of the start codon. Bioinformatics 2001; 17:890–900.

5. Morris DR, Geballe AP. Upstream open reading frames as regulators of mRNA translation. Mol. Cell Biol. 2000; 20:8635–8642.

6. Cruz-Vera LR, Sachs MS, Squires CL, Yanofsky C. Nascent polypeptide sequences that influence ribosome function. Curr. Opin. Microbiol. 2011; 14:160–166.

7. Ito K, Chiba S. Arrest peptides: cis-acting modulators of translation. Annu. Rev. Biochem. 2013; 82:171–202.

8. Somers J, Poyry T, Willis AE. A perspective on mammalian upstream open reading frame function. Int. J. Biochem. Cell Biol. 2013; 45:1690–1700.

9. Bhushan S, Meyer H, Starosta AL, Becker T, Mielke T, Berninghausen O, et al. Structural basis for translational stalling by human cytomegalovirus and fungal arginine attenuator peptide. Mol. Cell 2010; 40:138–146.

10. Wang Z, Sachs MS. Ribosome stalling is responsible for arginine-specific translational attenuation in *Neurospora crassa*. Mol. Cell Biol. 1997; 17:4904–4913.

11. Gaba A, Jacobson A, Sachs MS. Ribosome occupancy of the yeast *CPA1* upstream open reading frame termination codon modulates nonsense-mediated mRNA decay. Mol. Cell 2005; 20:449–460.

12. Uchiyama-Kadokura N, Murakami K, Takemoto M, Koyanagi N, Murota K, Naito S, et al. Polyamine-responsive ribosomal arrest at the stop codon of an upstream open reading frame of the AdoMetDC1 gene triggers nonsense-mediated mRNA decay in *Arabidopsis thaliana*. Plant Cell Physiol. 2014; 55:1556–1567.

13. Wang Z, Gaba A, Sachs MS. A highly conserved mechanism of regulated ribosome stalling mediated by fungal arginine attenuator peptides that appears independent of the charging status of arginyl-tRNAs. J. Biol. Chem. 1999; 274:37565–37574.

14. Law GL, Raney A, Heusner C, Morris DR. Polyamine regulation of ribosome pausing at the upstream open reading frame of S-adenosylmethionine decarboxylase. J. Biol. Chem. 2001; 276:38036–38043.

15. Hanfrey C, Elliott KA, Franceschetti M, Mayer MJ, Illingworth C, Michael AJ. A dual upstream open reading frame-based autoregulatory circuit controlling polyamine-responsive translation. J. Biol. Chem. 2005; 280:39229–39237.

16. Yamashita Y, Takamatsu S, Glasbrenner M, Becker T, Naito S, Beckmann R. Sucrose sensing through nascent peptide-meditated ribosome stalling at the stop codon of Arabidopsis *bZIP11* uORF2. FEBS Lett. 2017; 591:1266–1277.

17. Rahmani F, Hummel M, Schuurmans J, Wiese-Klinkenberg A, Smeekens S, Hanson J. Sucrose control of translation mediated by an upstream open reading frame-encoded peptide. Plant Physiol. 2009; 150:1356–1367.

18. Combier JP, de Billy F, Gamas P, Niebel A, Rivas S. Trans-regulation of the expression of the transcription factor MtHAP2-1 by a uORF controls root nodule development. Genes Dev. 2008; 22:1549–1559.

19. Crowe ML, Wang XQ, Rothnagel JA. Evidence for conservation and selection of upstream open reading frames suggests probable encoding of bioactive peptides. BMC genomics 2006; 7:16.

20. Hayden CA, Jorgensen RA. Identification of novel conserved peptide uORF homology groups in *Arabidopsis* and rice reveals ancient eukaryotic origin of select groups and preferential association with transcription factor-encoding genes. BMC Biol. 2007; 5:32.

21. Hayden CA, Bosco G. Comparative genomic analysis of novel conserved peptide upstream open reading frames in *Drosophila melanogaster* and other dipteran species. BMC genomics 2008; 9:61.

22. Tran MK, Schultz CJ, Baumann U. Conserved upstream open reading frames in higher plants. BMC genomics 2008; 9:361.

23. Vaughn JN, Ellingson SR, Mignone F, Arnim A. Known and novel post-transcriptional regulatory sequences are conserved across plant families. Rna 2012; 18:368–384.

24. van der Horst S, Snel B, Hanson J, Smeekens S. Novel pipeline identifies new upstream ORFs and non-AUG initiating main ORFs with conserved amino acid sequences in the 5’ leader of mRNAs in *Arabidopsis thaliana*. Rna 2018; 25:292–304.

25. Takahashi H, Takahashi A, Naito S, Onouchi H. BAIUCAS: a novel BLAST-based algorithm for the identification of upstream open reading frames with conserved amino acid sequences and its application to the *Arabidopsis thaliana* genome. Bioinformatics 2012; 28:2231–2241.

26. Ebina I, Takemoto-Tsutsumi M, Watanabe S, Koyama H, Endo Y, Kimata K, et al. Identification of novel *Arabidopsis thaliana* upstream open reading frames that control expression of the main coding sequences in a peptide sequence-dependent manner. Nucleic Acids Res. 2015; 43:1562–1576.

27. Noh AL, Watanabe S, Takahashi H, Naito S, Onouchi H. An upstream open reading frame represses expression of a tomato homologue of Arabidopsis *ANAC096*, a NAC domain transcription factor gene, in a peptide sequence-dependent manner. Plant Biotechnol. 2015; 32:157–163.

28. Jorgensen RA, Dorantes-Acosta AE. Conserved Peptide Upstream Open Reading Frames are Associated with Regulatory Genes in Angiosperms. Front. Plant Sci. 2012; 3:191.

29. Pruitt KD, Tatusova T, Maglott DR. NCBI reference sequences (RefSeq): a curated non-redundant sequence database of genomes, transcripts and proteins. Nucleic Acids Res. 2007; 35:D61–D65.

30. Zerbino DR, Achuthan P, Akanni W, Amode MR, Barrell D, Bhai J, et al. Ensembl 2018. Nucleic Acids Res. 2018; 46:D754–D761.

31. Goodstein DM, Shu S, Howson R, Neupane R, Hayes RD, Fazo J, et al. Phytozome: a comparative platform for green plant genomics. Nucleic Acids Res. 2012; 40:D1178–D1186.

32. Emms DM, Kelly S. OrthoFinder: solving fundamental biases in whole genome comparisons dramatically improves orthogroup inference accuracy. Genome Biol. 2015; 16:157.

33. Alatorre-Cobos F, Cruz-Ramirez A, Hayden CA, Perez-Torres CA, Chauvin AL, Ibarra-Laclette E, et al. Translational regulation of Arabidopsis XIPOTL1 is modulated by phosphocholine levels via the phylogenetically conserved upstream open reading frame 30. J. Exp. Bot. 2012; 63:5203–5221.

34. Guerrero-Gonzalez ML, Ortega-Amaro MA, Juarez-Montiel M, Jimenez-Bremont JF. Arabidopsis Polyamine oxidase-2 uORF is required for downstream translational regulation. Plant Physiol. Biochem. 2016; 108:381–390.

35. Imai A, Hanzawa Y, Komura M, Yamamoto KT, Komeda Y, Takahashi T. The dwarf phenotype of the Arabidopsis acl5 mutant is suppressed by a mutation in an upstream ORF of a bHLH gene. Development 2006; 133:3575–3585.

36. Ribone PA, Capella M, Arce AL, Chan RL. A uORF Represses the Transcription Factor AtHB1 in Aerial Tissues to Avoid a Deleterious Phenotype. Plant Physiol. 2017; 175:1238–1253.

37. Tabuchi T, Okada T, Azuma T, Nanmori T, Yasuda T. Posttranscriptional regulation by the upstream open reading frame of the phosphoethanolamine N-methyltransferase gene. Biosci. Biotechnol. Biochem. 2006; 70:2330–2334.

38. Hayashi N, Sasaki S, Takahashi H, Yamashita Y, Naito S, Onouchi H. Identification of *Arabidopsis thaliana* upstream open reading frames encoding peptide sequences that cause ribosomal arrest. Nucleic Acids Res. 2017; 45:8844–8858.

39. Franceschetti M, Hanfrey C, Scaramagli S, Torrigiani P, Bagni N, Burtin D, et al. Characterization of monocot and dicot plant S-adenosyl-l-methionine decarboxylase gene families including identification in the mRNA of a highly conserved pair of upstream overlapping open reading frames. Biochem. J. 2001; 353:403–409.

40. Konishi T, Takeda T, Miyazaki Y, Ohnishi-Kameyama M, Hayashi T, O’Neill MA, et al. A plant mutase that interconverts UDP-arabinofuranose and UDP-arabinopyranose. Glycobiology 2007; 17:345–354.

41. Zerbino DR, Birney E. Velvet: algorithms for de novo short read assembly using de Bruijn graphs. Genome Res. 2008; 18:821–829.

42. Langmead B, Salzberg SL. Fast gapped-read alignment with Bowtie 2. Nat. Methods 2012; 9:357–359.

43. Altschul SF, Madden TL, Schaffer AA, Zhang J, Zhang Z, Miller W, et al. Gapped BLAST and PSI-BLAST: a new generation of protein database search programs. Nucleic Acids Res. 1997; 25:3389–3402.

44. Sievers F, Wilm A, Dineen D, Gibson TJ, Karplus K, Li W, et al. Fast, scalable generation of high-quality protein multiple sequence alignments using Clustal Omega. Mol. Syst. Biol. 2011; 7:539.

45. Li WH. Unbiased estimation of the rates of synonymous and nonsynonymous substitution. J. Mol. Evol. 1993; 36:96–99.

46. Charif D, Lobry JR SeqinR 1.0-2: a contributed package to the R project for statistical computing devoted to biological sequences retrieval and analysis. In Structural Approaches to Sequence Evolution: Molecules, Networks, Populations. Edited by Bastolla U, Porto M, Roman HE, Vendruscolo M. New York: Springer Verlag; 2007: 207–232

47. Benjamini Y, Hochberg Y. Controlling the false discovery rate: a practical and powerful approach to multiple testing. J. R. Statist. Soc. ser.B 1995; 57:298–300.

48. Clamp M, Cuff J, Searle SM, Barton GJ. The Jalview Java alignment editor. Bioinformatics 2004; 20:426–427.

49. Lawrence M, Huber W, Pages H, Aboyoun P, Carlson M, Gentleman R, et al. Software for computing and annotating genomic ranges. PLoS Comput. Biol. 2013; 9:e1003118.

50. Nanjo T, Futamura N, Nishiguchi M, Igasaki T, Shinozaki K, Shinohara K. Characterization of full-length enriched expressed sequence tags of stress-treated poplar leaves. Plant Cell Physiol. 2004; 45:1738–1748.

51. Motohashi K. Seamless Ligation Cloning Extract (SLiCE) Method Using Cell Lysates from Laboratory Escherichia coli Strains and its Application to SLiP Site-Directed Mutagenesis. Methods Mol. Biol. 2017; 1498:349–357.

52. Ho SN, Hunt HD, Horton RM, Pullen JK, Pease LR. Site-directed mutagenesis by overlap extension using the polymerase chain reaction. Gene 1989; 77:51–59.

53. Menges M, Murray JA. Synchronous *Arabidopsis* suspension cultures for analysis of cell-cycle gene activity. Plant J. 2002; 30:203–212.

