## Supplementary data for "Comprehensive genome-wide identification of angiosperm upstream ORFs with peptide sequences conserved in various taxonomic ranges using a novel pipeline, ESUCA"

### Supplementary Figure S1

## HG002.2

POPTR_0001s22450

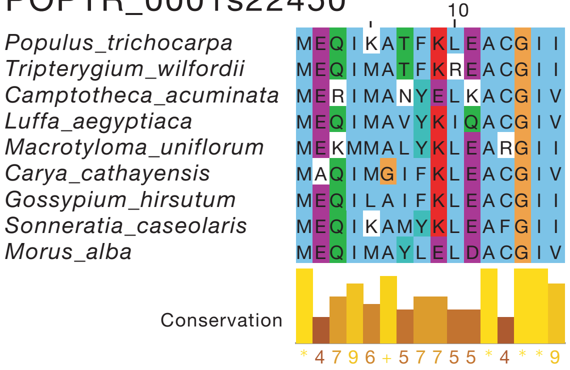


## HG009.2

POPTR_0003s01750

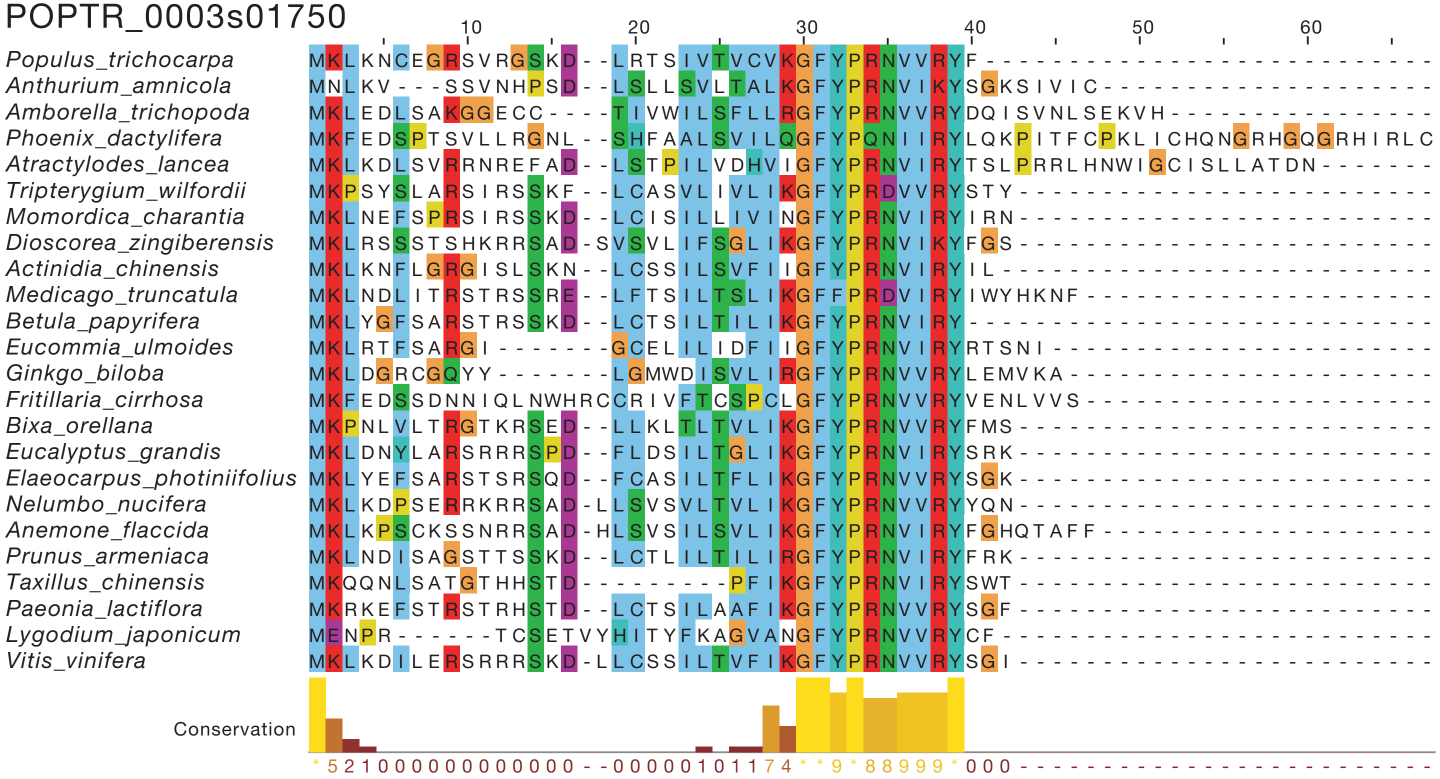

**HG016.2**

AT3G51630

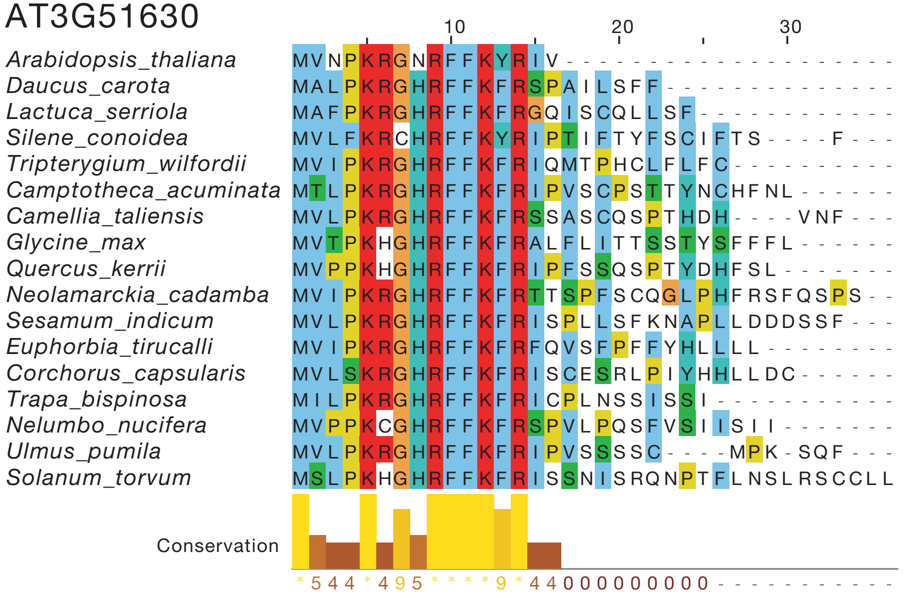


**HG016.2**

POPTR_0013s15110

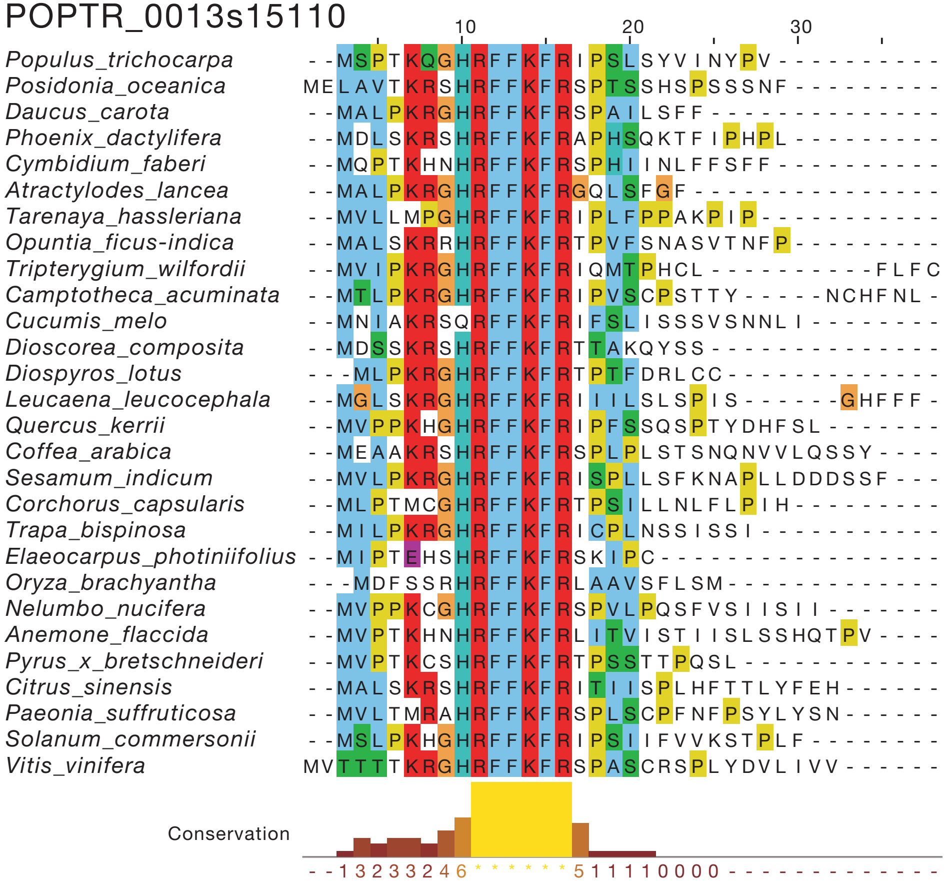

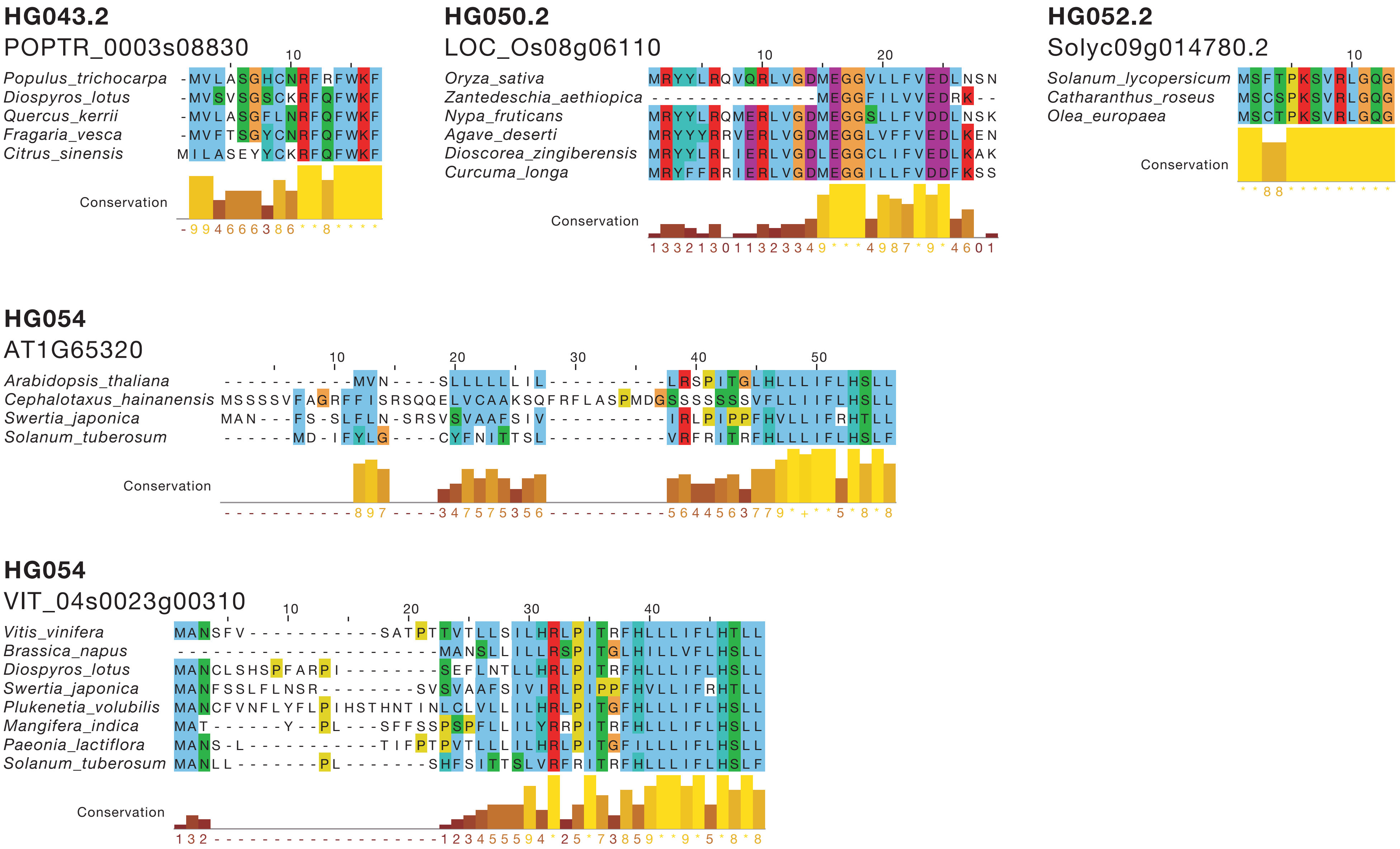
HG043.2
POPTR_0003s08830
Populus_trichocarpa
Diospyros_lotus
Quercus_kerrii
Fragaria_vesca
Citrus_sinensis
Conservation
HG050.2
LOC_Os08g06110
Oryza_sativa
Zantedeschia_aethiopica
Nypa_fruticans
Agave_deserti
Dioscorea_zingiberensis
Curcuma_longa
Conservation
HG052.2
Solyc09g014780.2
Solanum_lycopersicum
Catharanthus_roseus
Olea_europaea
Conservation
HG054
AT1G65320
Arabidopsis_thaliana
Cephalotaxus_hainanensis
Swertia_japonica
Solanum_tuberosum
Conservation
HG054
VIT_04s0023g00310
Vitis_vinifera
Brassica_napus
Diospyros_lotus
Swertia_japonica
Plukenetia_volubilis
Mangifera_indica
Paeonia_lactiflora
Solanum_tuberosum
Conservation

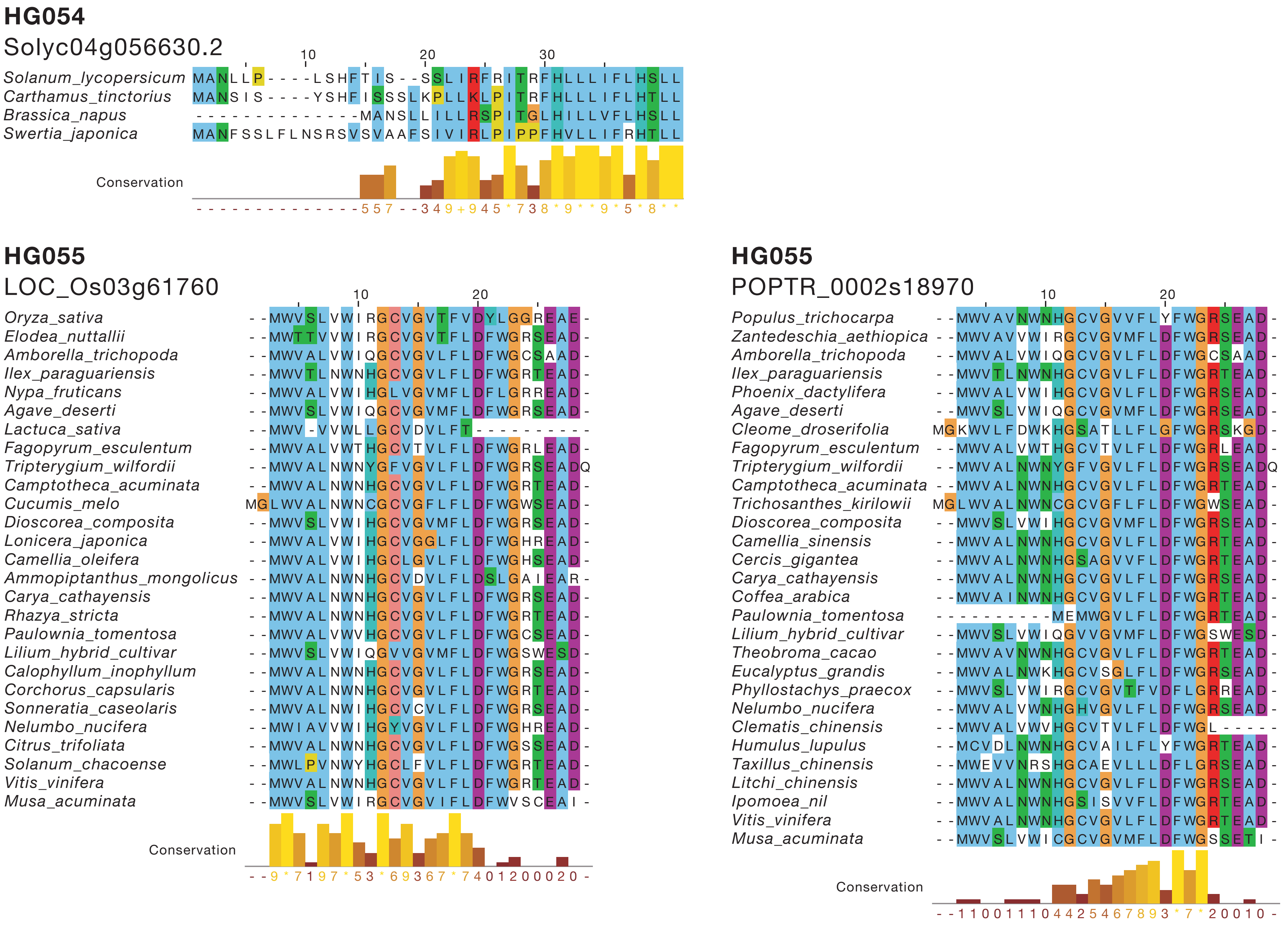
HG054
Solyc04g056630.2
Solanum_lycopersicum
Carthamus_tinctorius
Brassica_napus
Swertia_japonica
Conservation
HG055
LOC_Os03g61760
Oryza_sativa
Elodea_nuttallii
Amborella_trichopoda
Ilex_paraguariensis
Nypa_fruticans
Agave_deserti
Lactuca_sativa
Fagopyrum_esculentum
Tripterygium_wilfordii
Camptotheca_acuminata
Cucumis_melo
Dioscorea_composita
Lonicera_japonica
Camellia_oleifera
Ammopiptanthus_mongolicus
Carya_cathayensis
Rhazya_stricta
Paulownia_tomentosa
Lilium_hybrid_cultivar
Calophyllum_inophyllum
Corchorus_capsularis
Sonneratia_caseolaris
Nelumbo_nucifera
Citrus_trifoliata
Solanum_chacoense
Vitis_vinifera
Musa_acuminata
Conservation
HG055
POPTR_0002s18970
Populus_trichocarpa
Zantedeschia_aethiopica
Amborella_trichopoda
Ilex_paraguariensis
Phoenix_dactylifera
Agave_deserti
Cleome_droserifolia
Fagopyrum_esculentum
Tripterygium_wilfordii
Camptotheca_acuminata
Trichosanthes_kirilowii
Dioscorea_composita
Camellia_sinensis
Cercis_gigantea
Carya_cathayensis
Coffea_arabica
Paulownia_tomentosa
Lilium_hybrid_cultivar
Theobroma_cacao
Eucalyptus_grandis
Phyllostachys_praecox
Nelumbo_nucifera
Clematis_chinensis
Humulus_lupulus
Taxillus_chinensis
Litchi_chinensis
Ipomoea_nil
Vitis_vinifera
Musa_acuminata
Conservation

**HG055**

POPTR_0014s10960

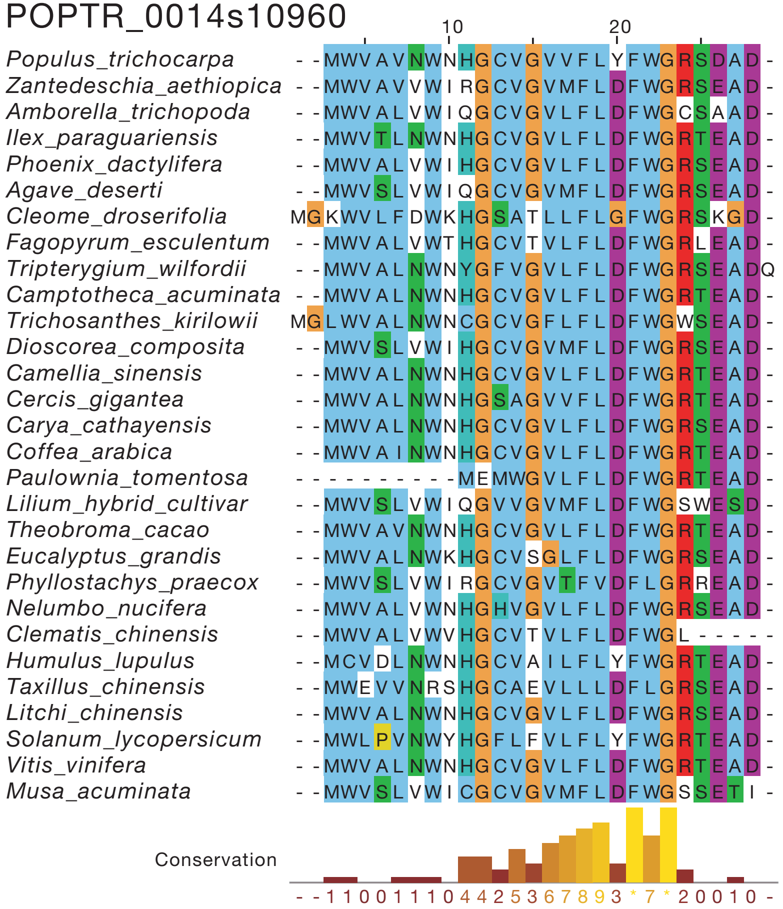


**HG055**

VIT_05s0020g00700

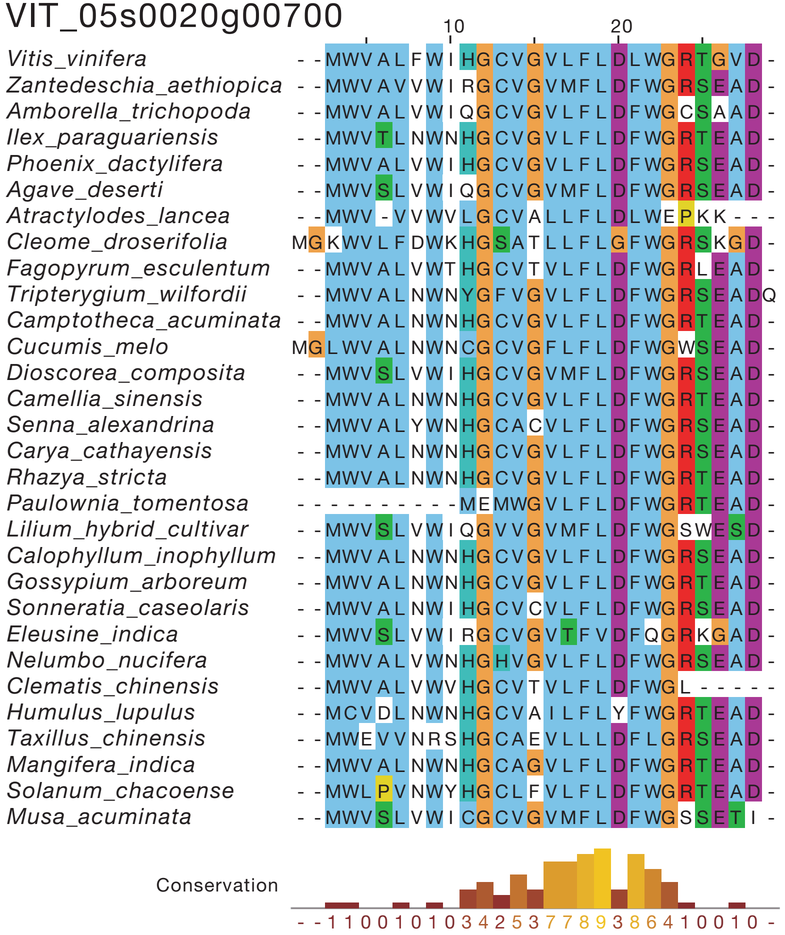

# HG055

VIT_07s0005g02260

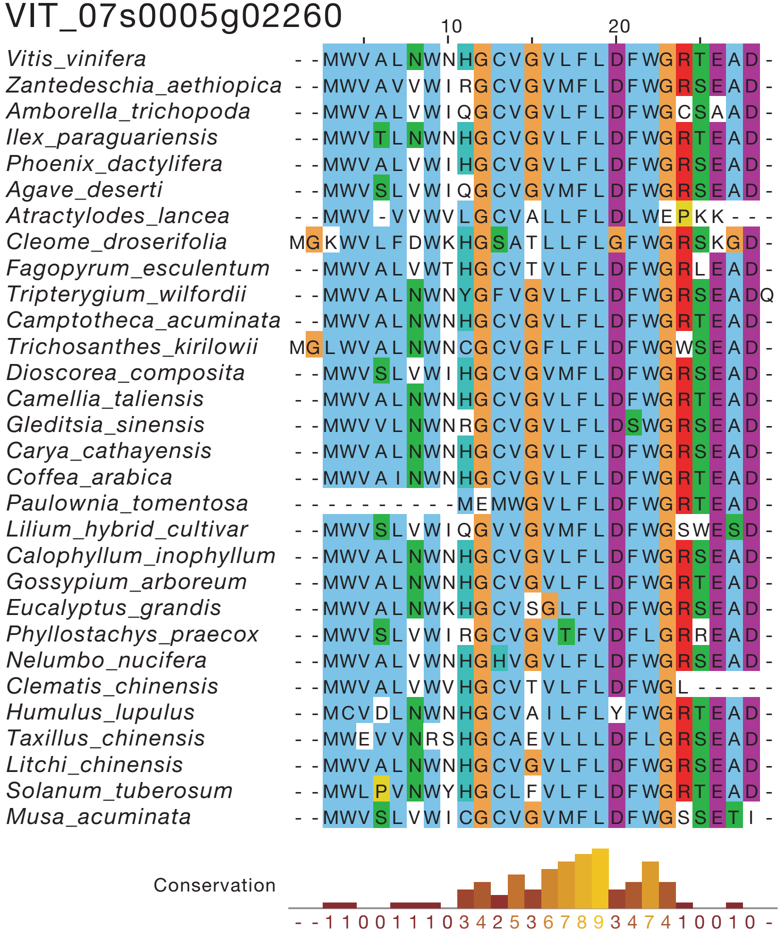

**HG056.1**

POPTR_0004s05490

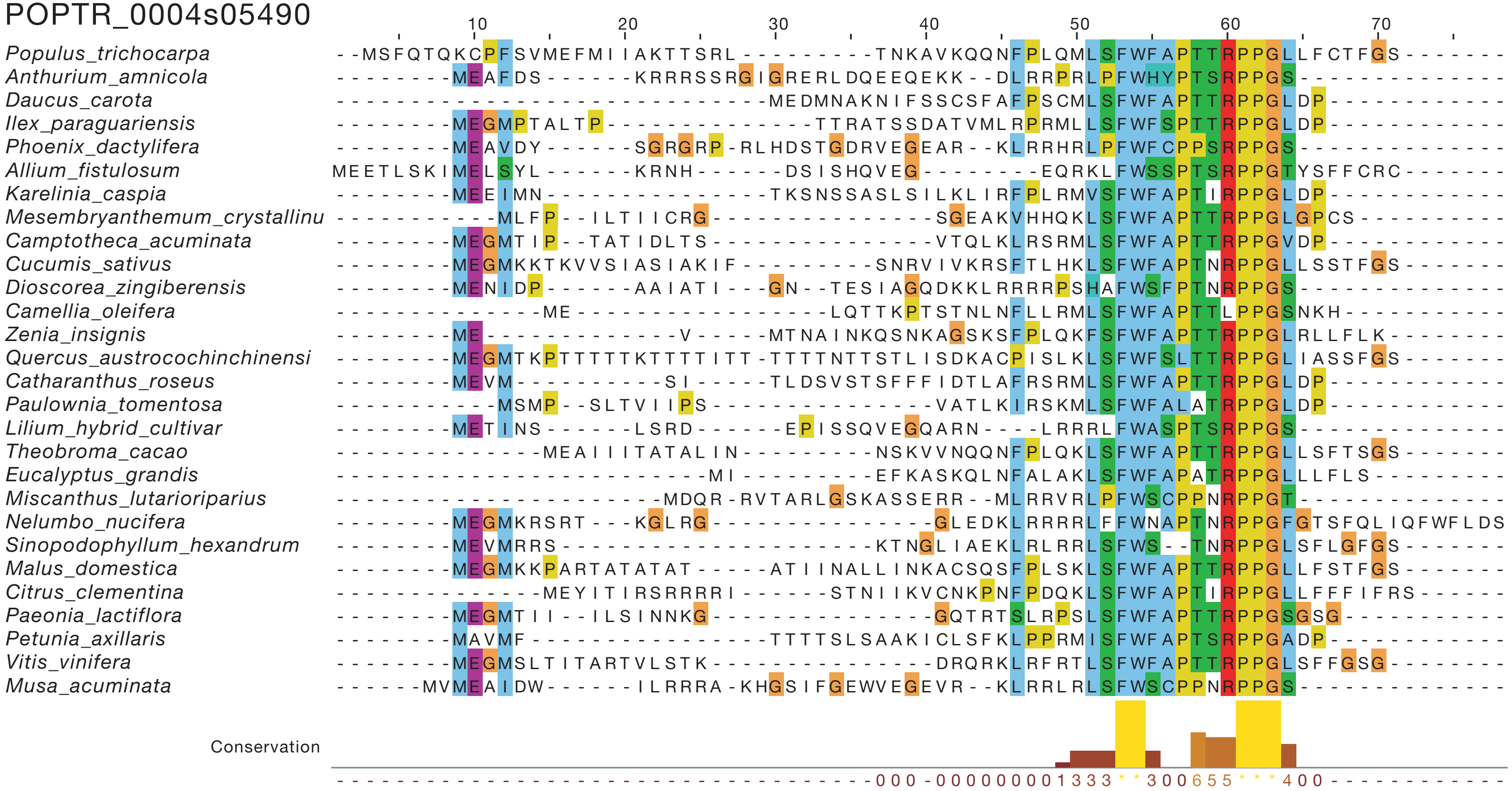

**HG056.1**

Solyc02g076920.2

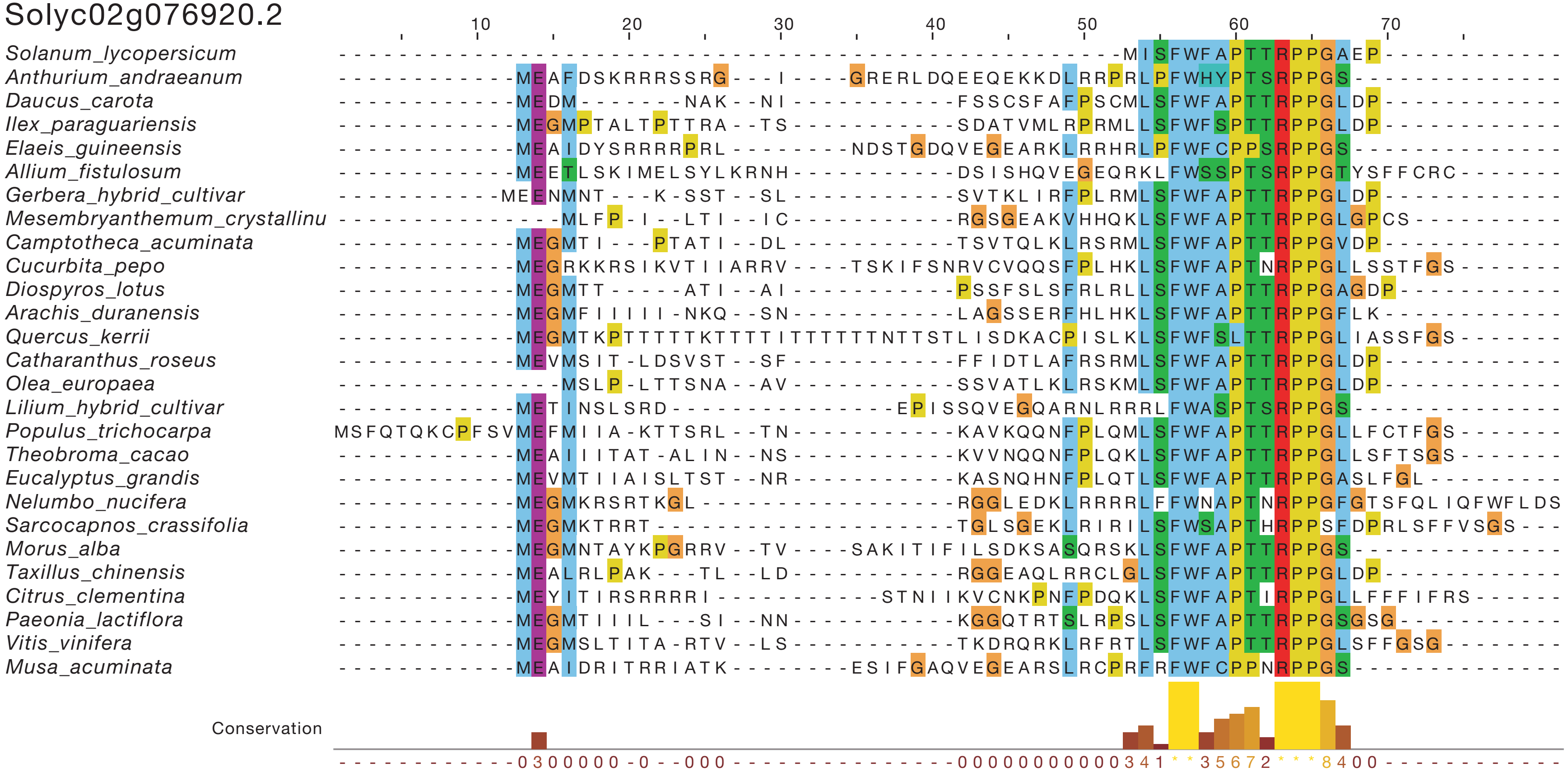

## HG056.1

Solyc07g053290.2

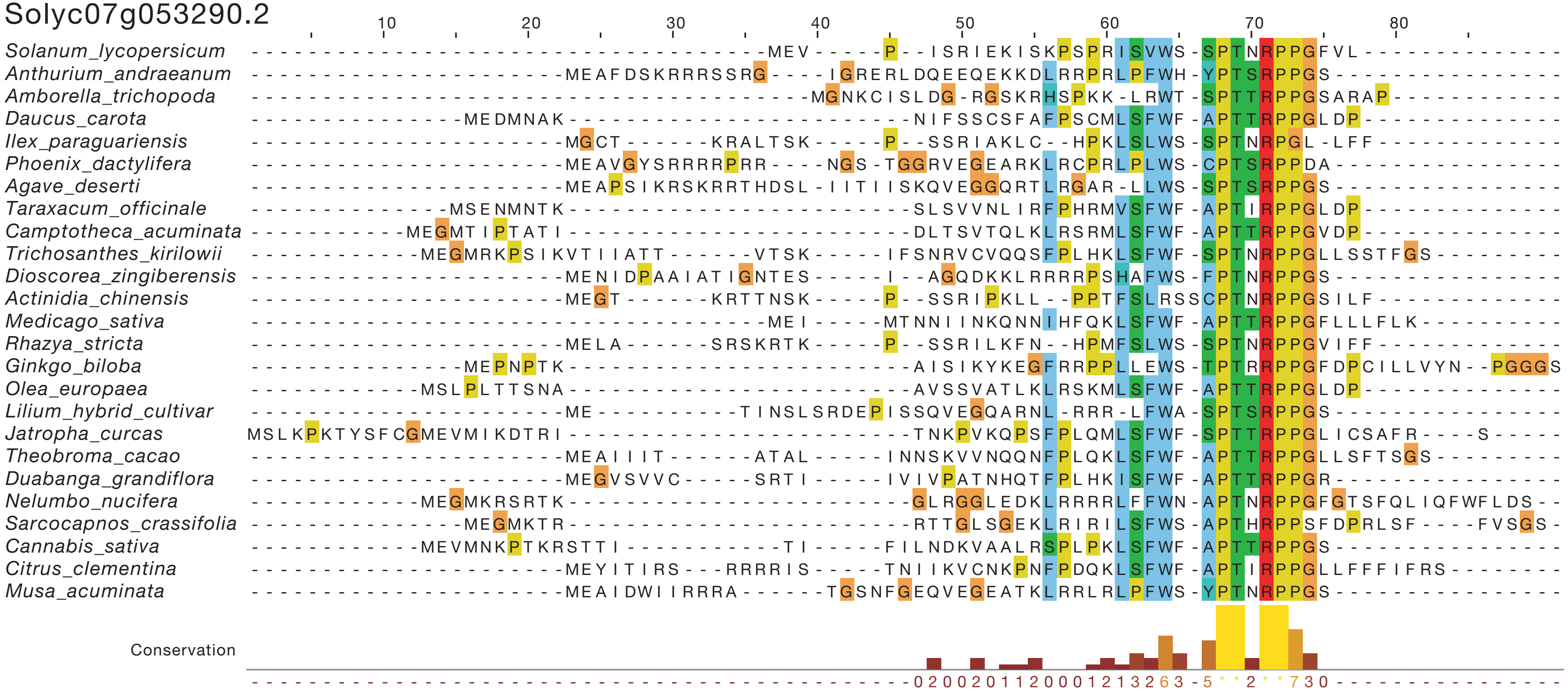

**HG056.1**

VIT_10s0003g01170

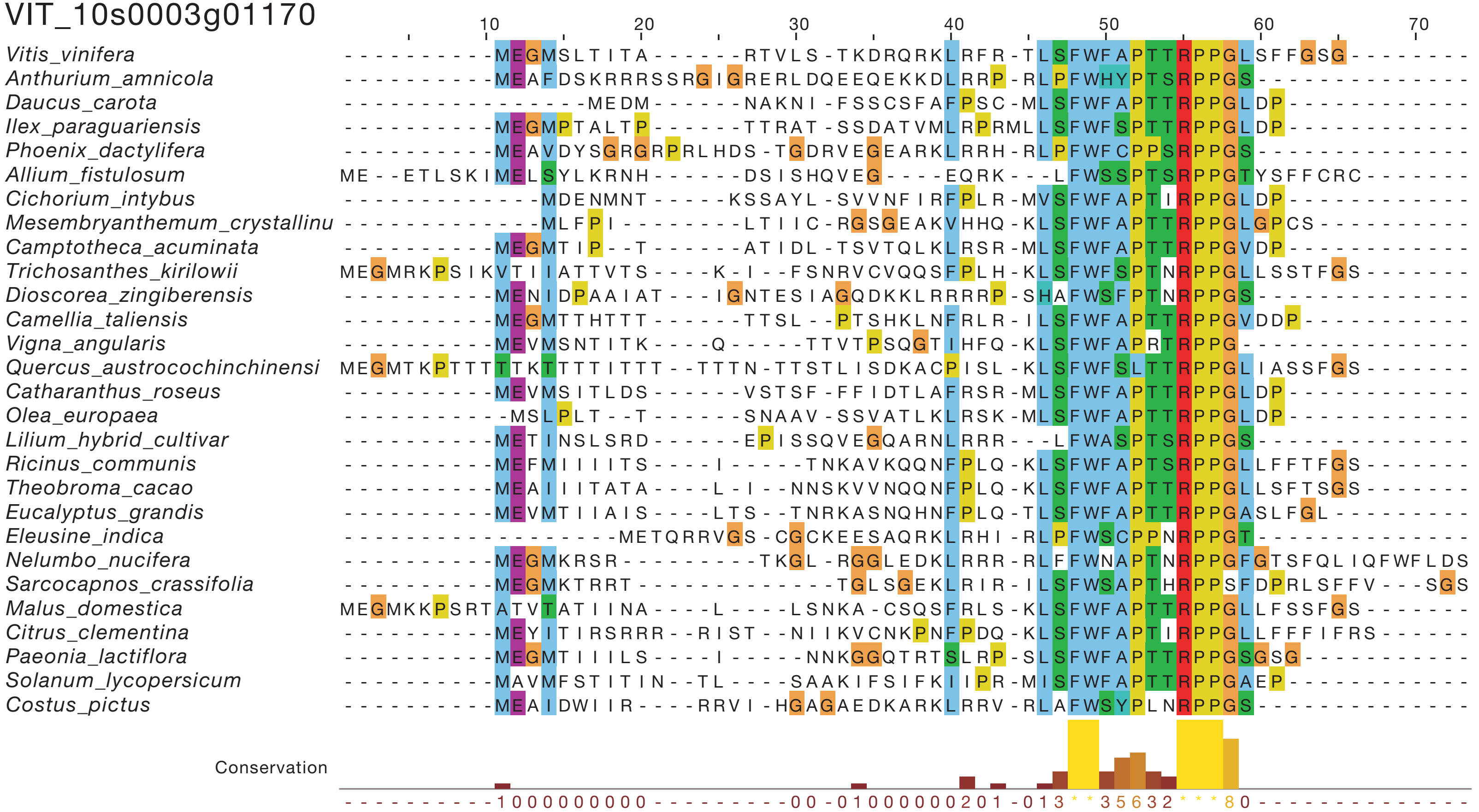

## HG056.2
POPTR_0004s05490

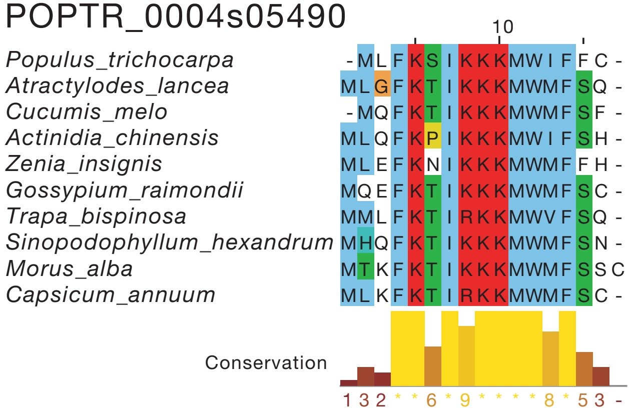


## HG056.2
VIT_10s0003g01170

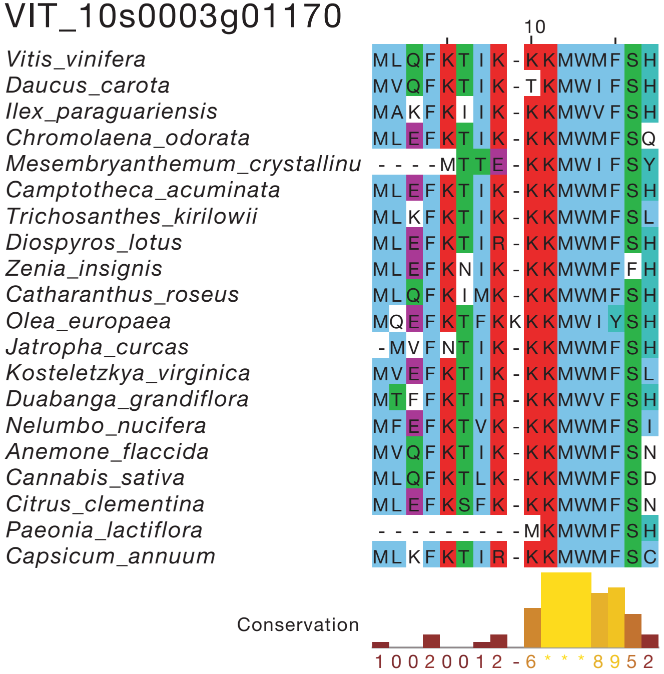


## HG057
AT1G54095

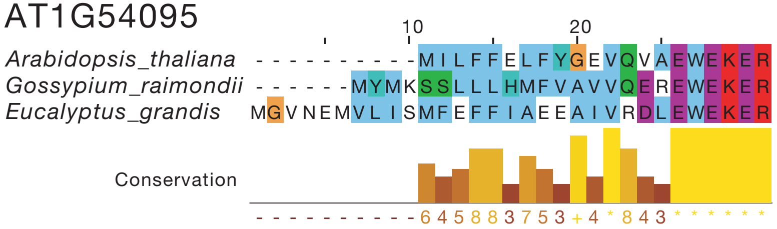


## HG057
AT1G72510

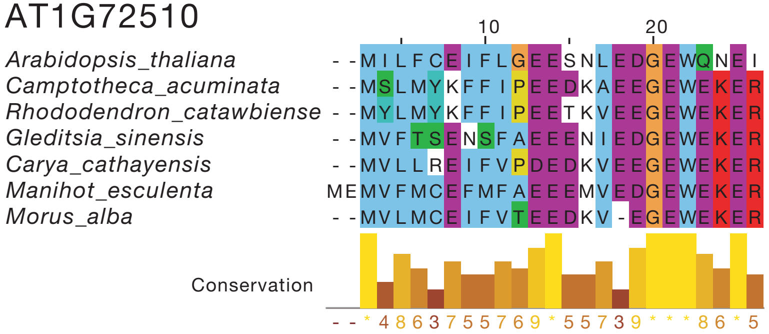

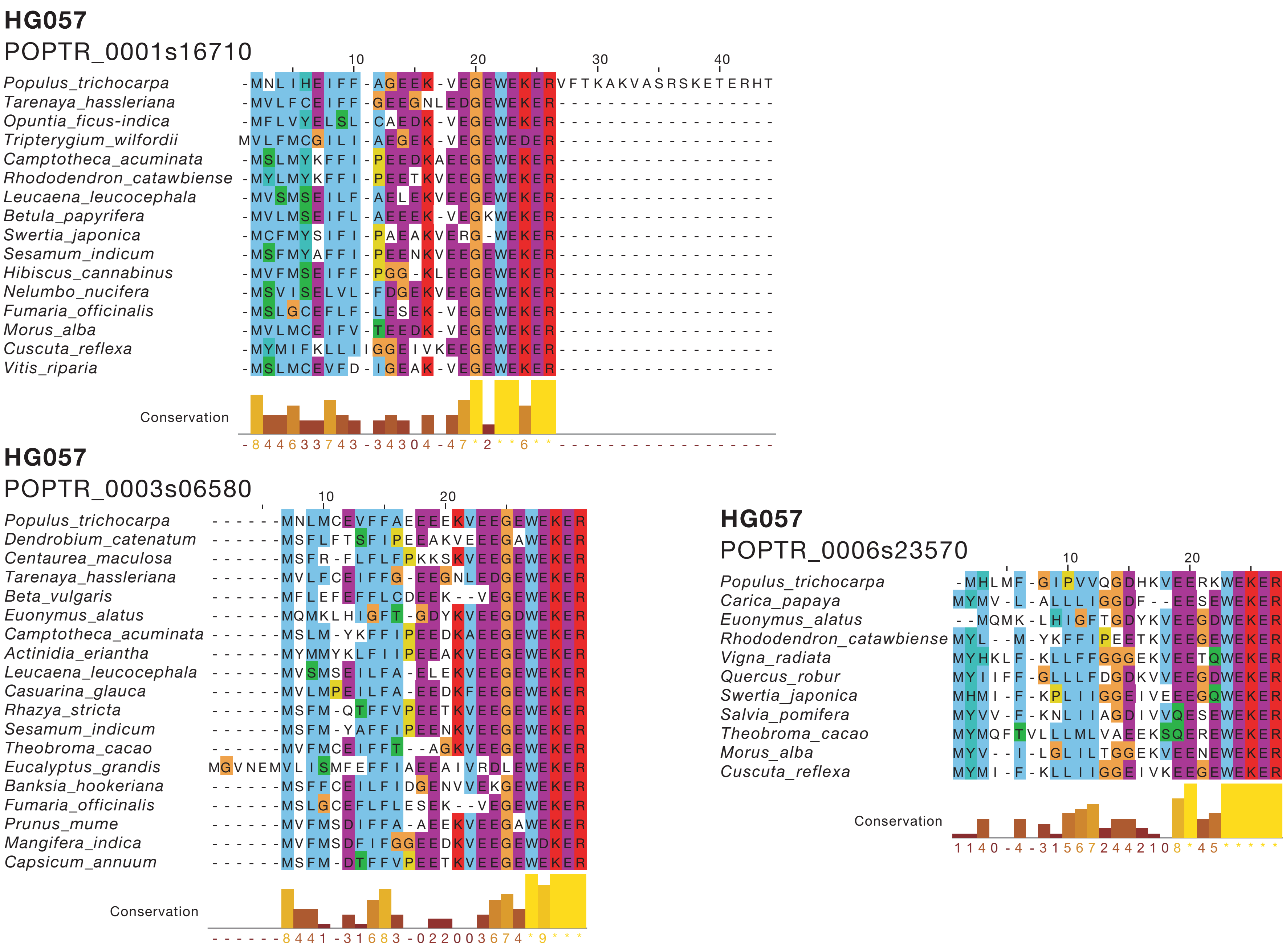
HG057
POPTR_0001s16710
Populus_trichocarpa
Tarenaya_hassleriana
Opuntia_ficus-indica
Tripterygium_wilfordii
Camptotheca_acuminata
Rhododendron_catawbiense
Leucaena_leucocephala
Betula_papyrifera
Swertia_japonica
Sesamum_indicum
Hibiscus_cannabinus
Nelumbo_nucifera
Fumaria_officinalis
Morus_alba
Cuscuta_reflexa
Vitis_riparia
Conservation
HG057
POPTR_0003s06580
Populus_trichocarpa
Dendrobium_catenatum
Centaurea_maculosa
Tarenaya_hassleriana
Beta_vulgaris
Euonymus_alatus
Camptotheca_acuminata
Actinidia_eriantha
Leucaena_leucocephala
Casuarina_glauca
Rhazya_stricta
Sesamum_indicum
Theobroma_cacao
Eucalyptus_grandis
Banksia_hookeriana
Fumaria_officinalis
Prunus_mume
Mangifera_indica
Capsicum_annuum
Conservation
HG057
POPTR_0006s23570
Populus_trichocarpa
Carica_papaya
Euonymus_alatus
Rhododendron_catawbiense
Vigna_radiata
Quercus_robur
Swertia_japonica
Salvia_pomifera
Theobroma_cacao
Morus_alba
Cuscuta_reflexa
Conservation

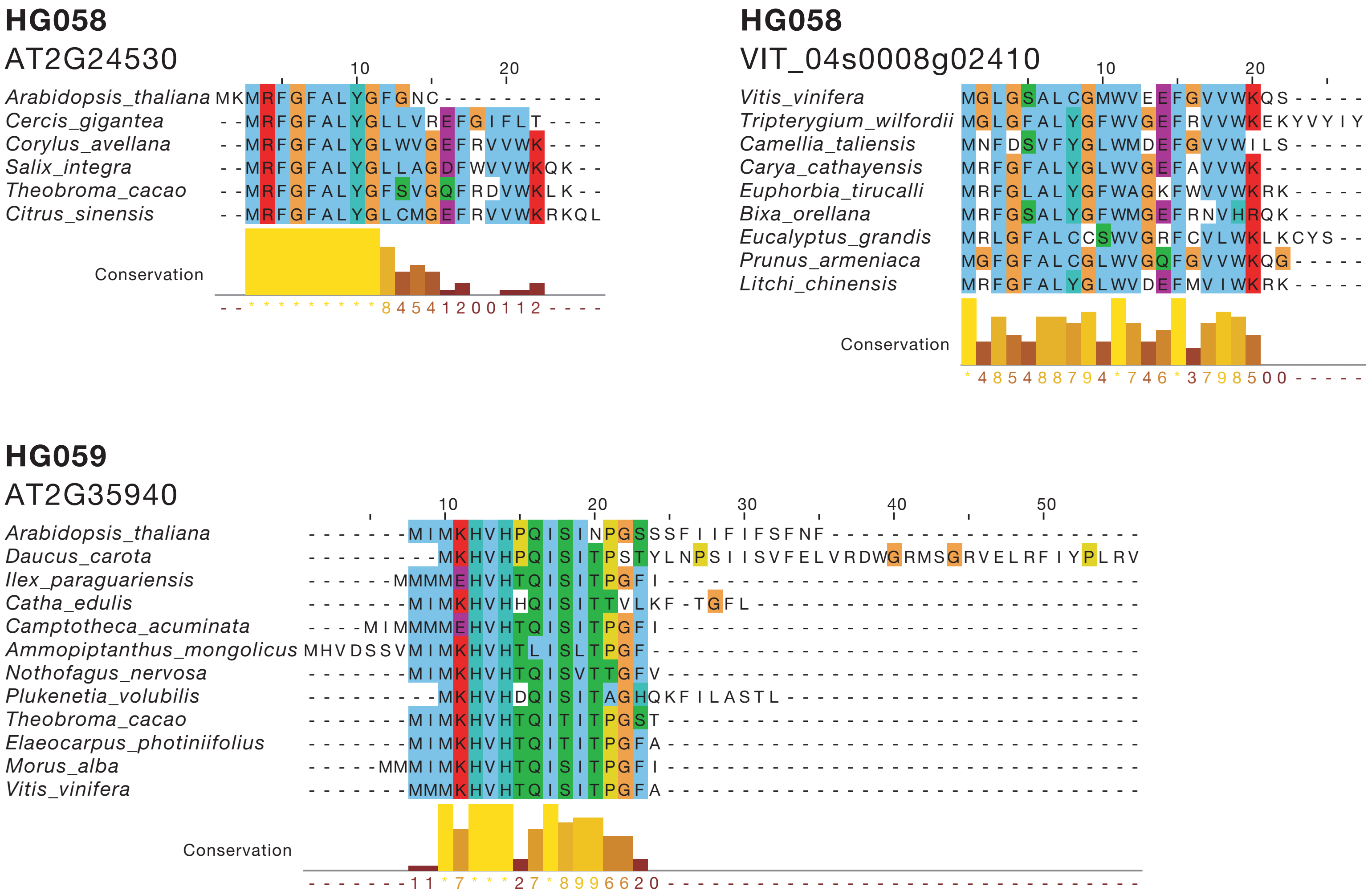
HG058
AT2G24530
Arabidopsis_thaliana
Cercis_gigantea
Corylus_avellana
Salix_integra
Theobroma_cacao
Citrus_sinensis
Conservation
HG058
VIT_04s0008g02410
Vitis_vinifera
Tripterygium_wilfordii
Camellia_taliensis
Carya_cathayensis
Euphorbia_tirucalli
Bixa_orellana
Eucalyptus_grandis
Prunus_armeniaca
Litchi_chinensis
Conservation
HG059
AT2G35940
Arabidopsis_thaliana
Daucus_carota
Ilex_paraguariensis
Catha_edulis
Camptotheca_acuminata
Ammopiptanthus_mongolicus
Nothofagus_nervosa
Plukenetia_volubilis
Theobroma_cacao
Elaeocarpus_photiniifolius
Morus_alba
Vitis_vinifera
Conservation

**HG059**

POPTR_0006s21950

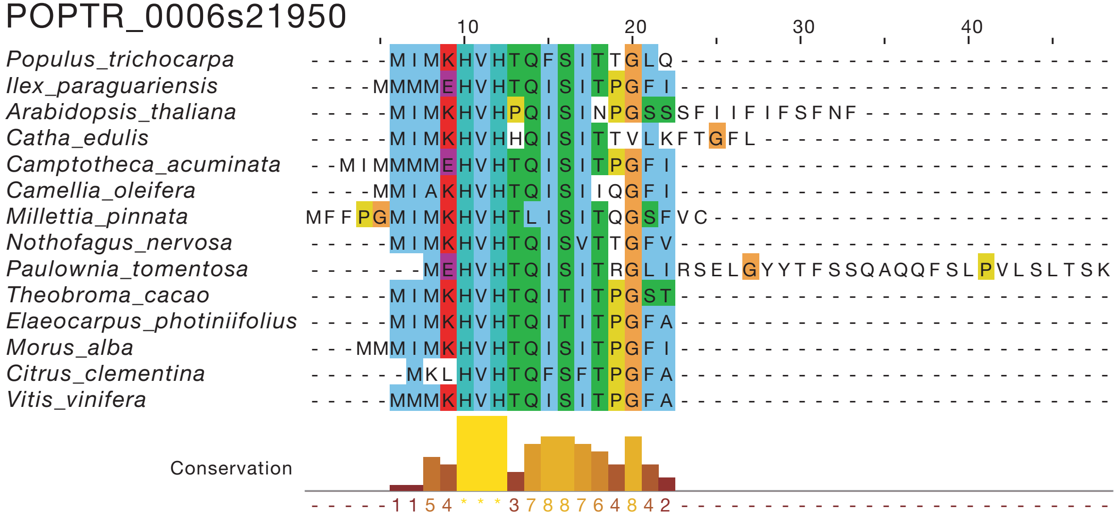


**HG059**

POPTR_0016s07040

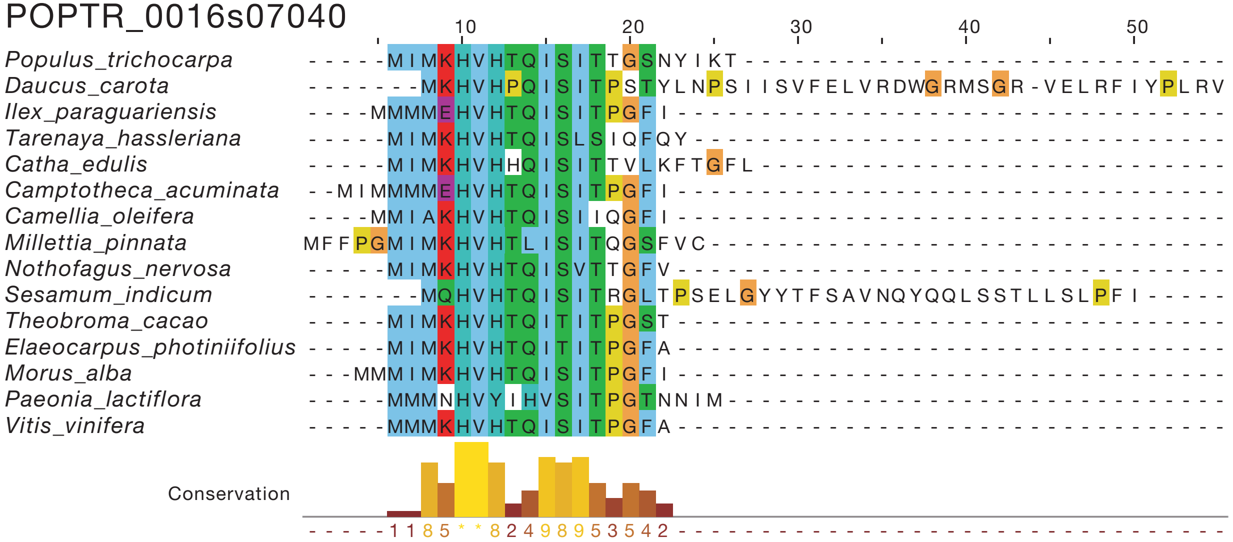

## HG060

AT3G12570

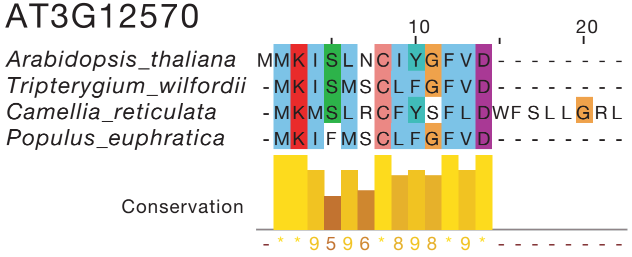


## HG060

POPTR_0008s05420

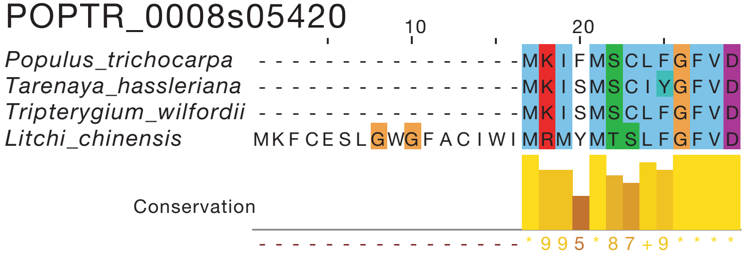


## HG060

POPTR_0010s21340

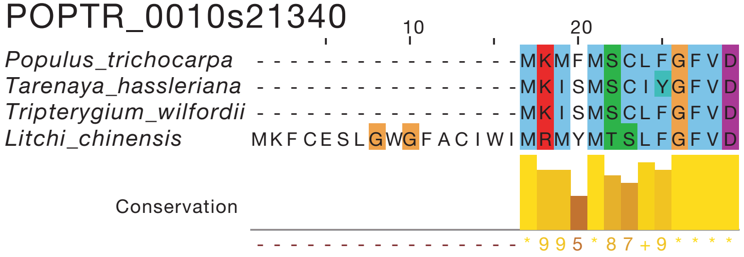

## HG061

AT5G55600

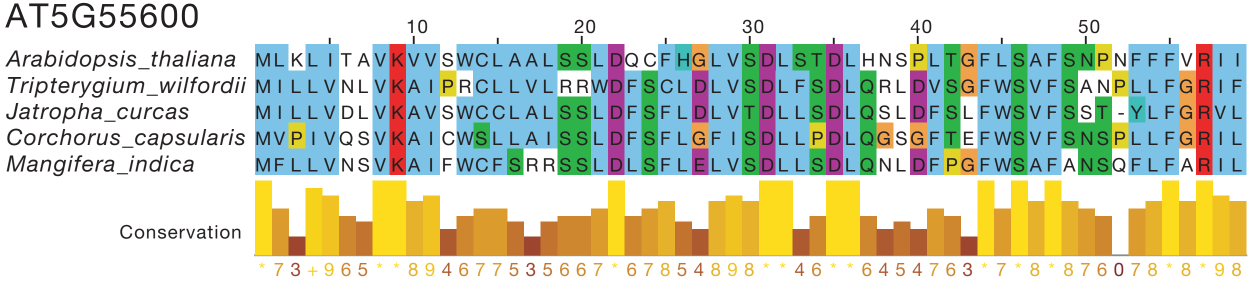


## HG061

POPTR_0001s37450

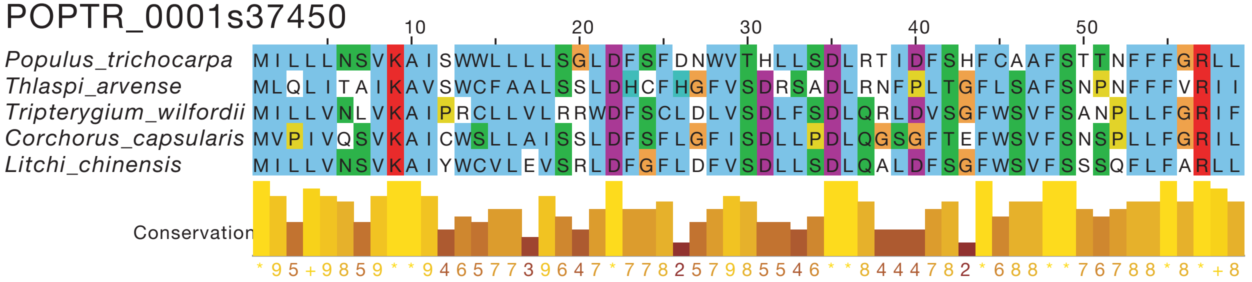


## HG062

POPTR_0008s09440

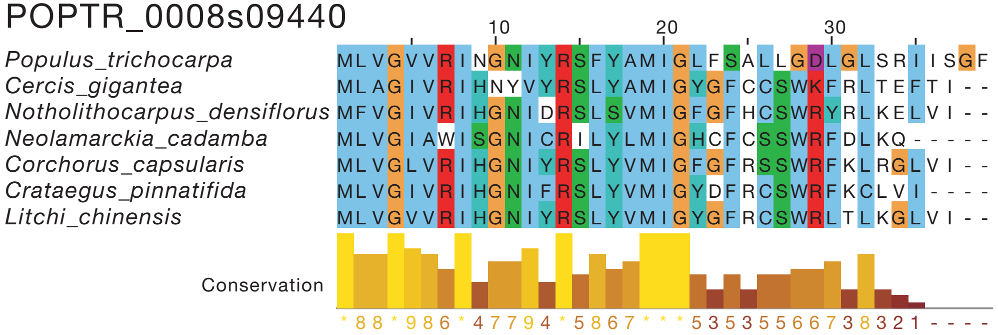

## HG062
Solyc04g005430.2

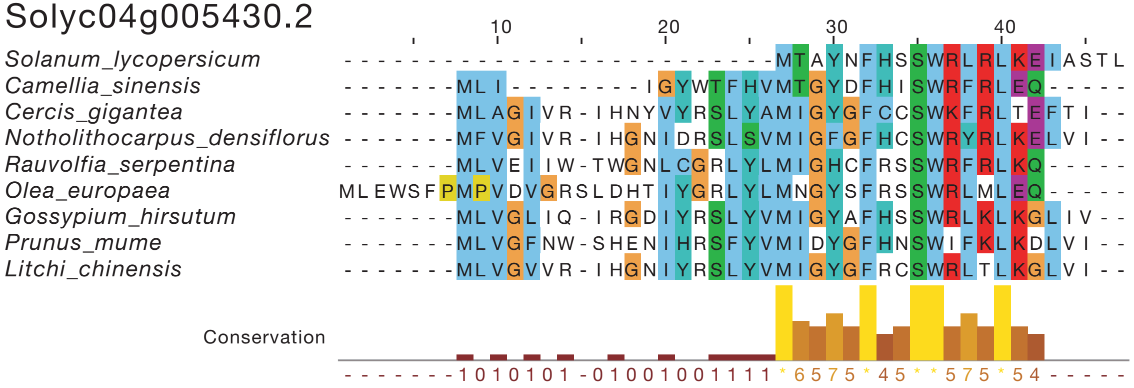


## HG062
Solyc05g012450.2

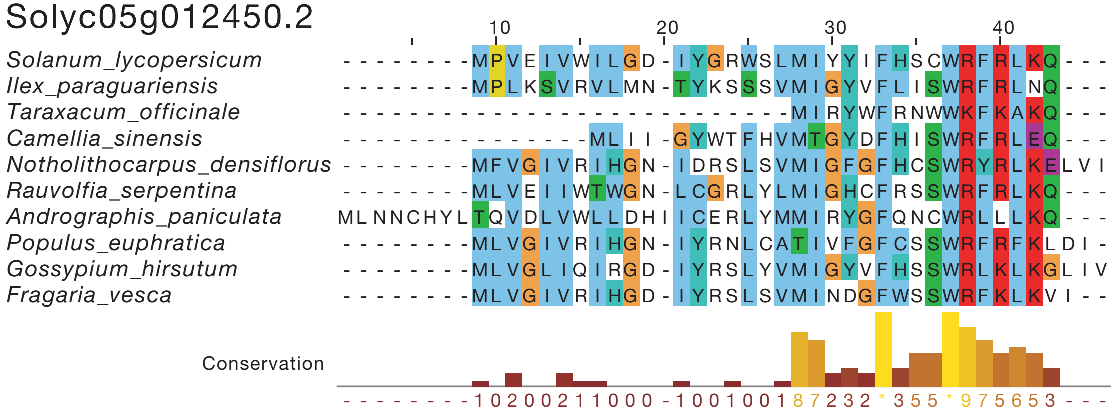


## HG063
Solyc04g071860.2

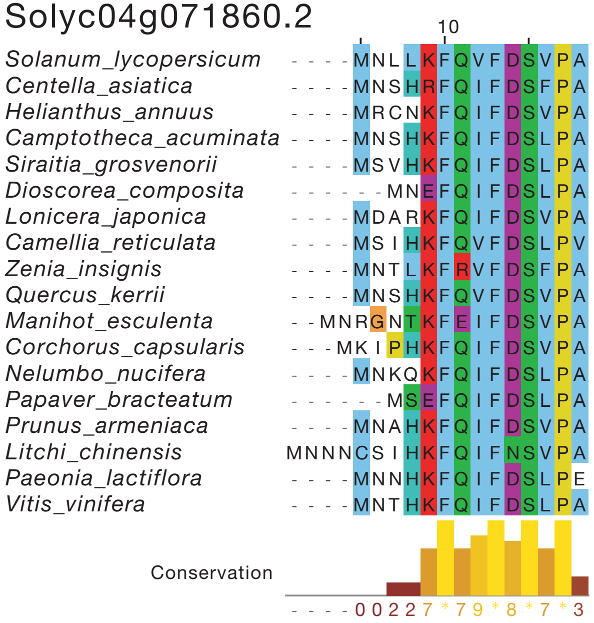

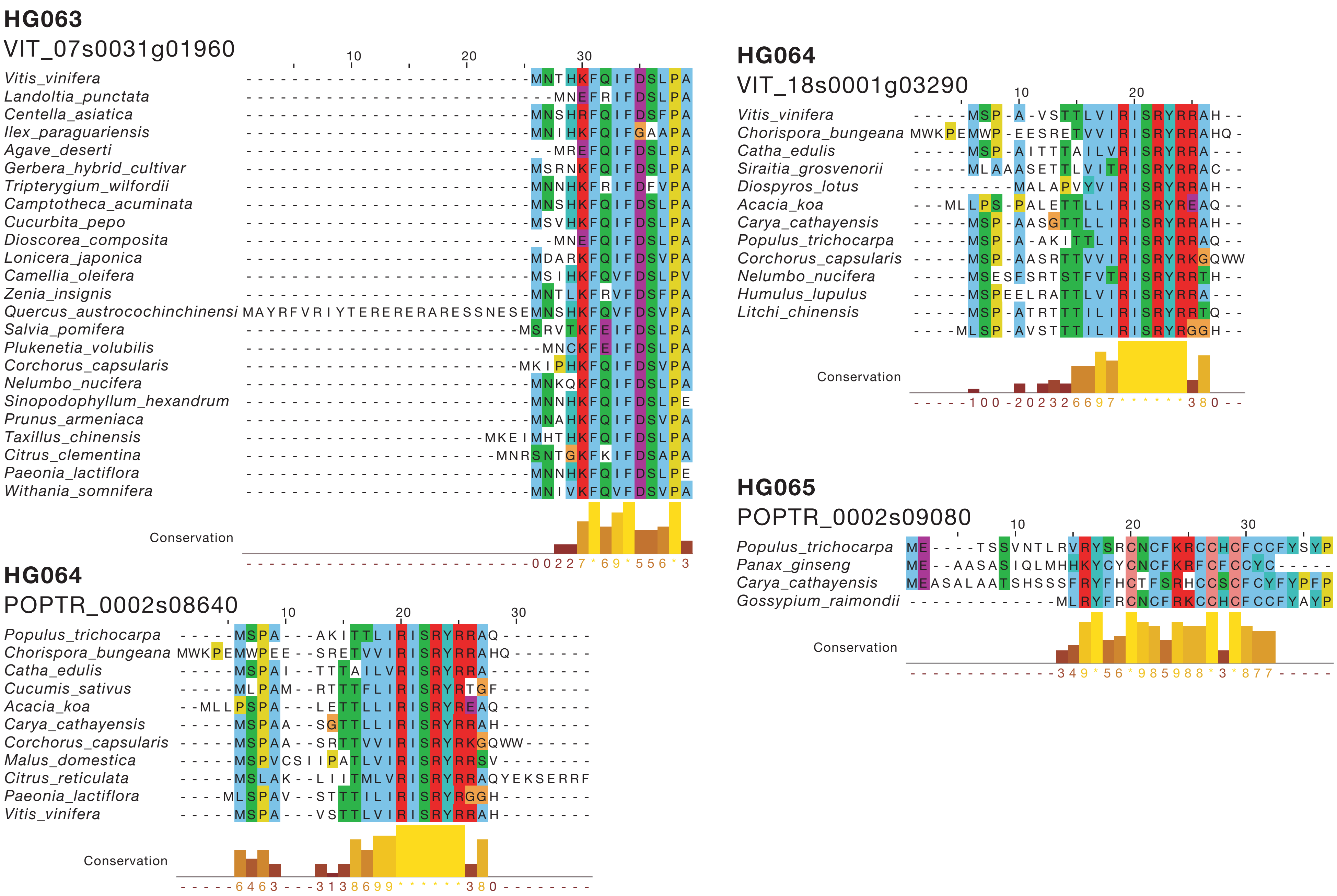
HG063
VIT_07s0031g01960
Vitis_vinifera
Landoltia_punctata
Centella_asiatica
Ilex_paraguariensis
Agave_deserti
Gerbera_hybrid_cultivar
Tripterygium_wilfordii
Camptotheca_acuminata
Cucurbita_pepo
Dioscorea_composita
Lonicera_japonica
Camellia_oleifera
Zenia_insignis
Quercus_austrocochinchinensi
Salvia_pomifera
Plukenetia_volubilis
Corchorus_capsularis
Nelumbo_nucifera
Sinopodophyllum_hexandrum
Prunus_armeniaca
Taxillus_chinensis
Citrus_clementina
Paeonia_lactiflora
Withania_somnifera
Conservation
HG064
POPTR_0002s08640
Populus_trichocarpa
Chorispora_bungeana
Catha_edulis
Cucumis_sativus
Acacia_koa
Carya_cathayensis
Corchorus_capsularis
Malus_domestica
Citrus_reticulata
Paeonia_lactiflora
Vitis_vinifera
Conservation
HG064
VIT_18s0001g03290
Vitis_vinifera
Chorispora_bungeana
Catha_edulis
Siraitia_grosvenorii
Diospyros_lotus
Acacia_koa
Carya_cathayensis
Populus_trichocarpa
Corchorus_capsularis
Nelumbo_nucifera
Humulus_lupulus
Litchi_chinensis
Conservation
HG065
POPTR_0002s09080
Populus_trichocarpa
Panax_ginseng
Carya_cathayensis
Gossypium_raimondii
Conservation

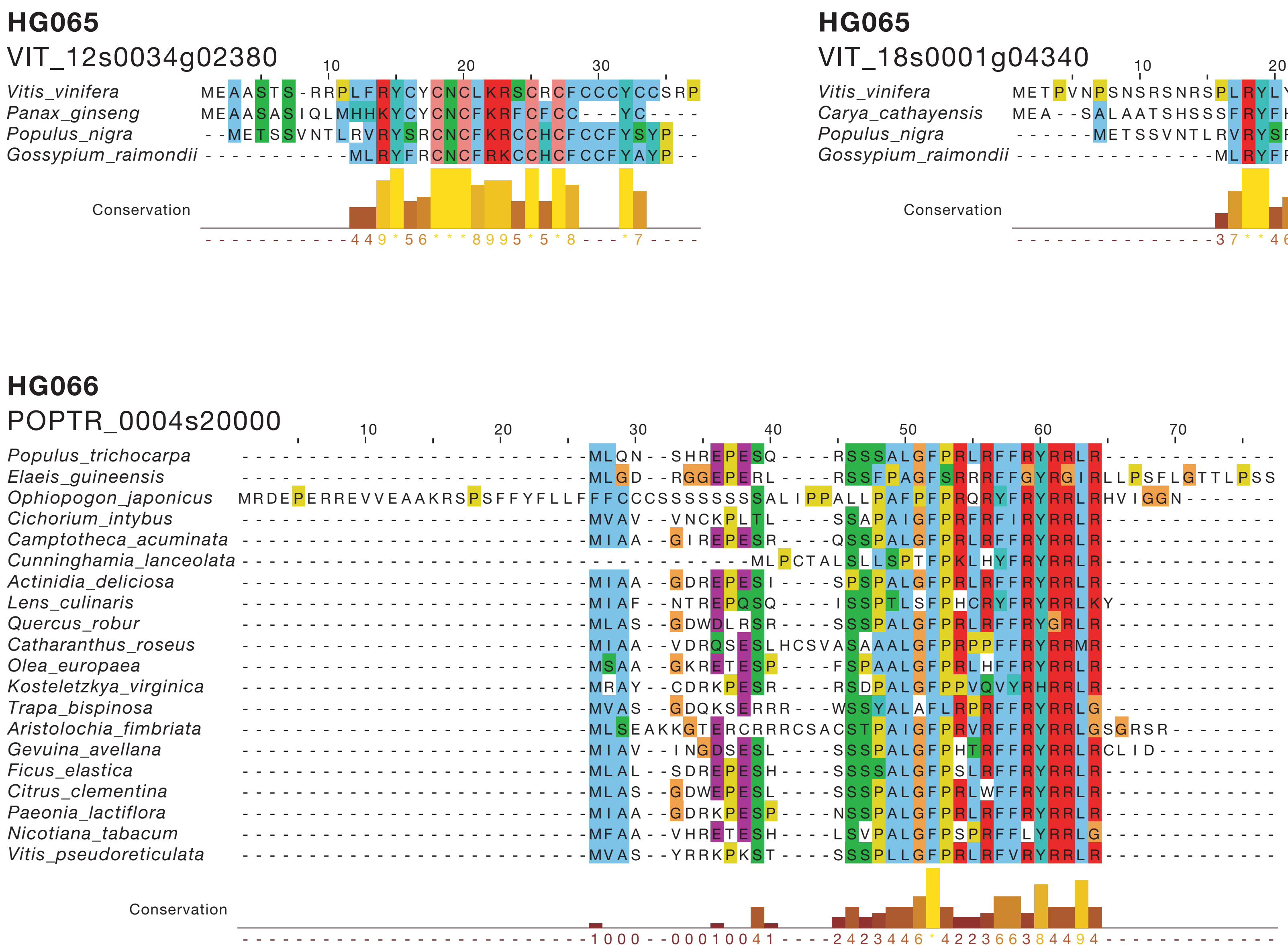

HG065
VIT_12s0034g02380
10
20
30
Vitis_vinifera
Panax_ginseng
Populus_nigra
Gossypium_raimondii
Conservation
HG065
VIT_18s0001g04340
10
20
30
Vitis_vinifera
Carya_cathayensis
Populus_nigra
Gossypium_raimondii
Conservation
HG066
POPTR_0004s20000
10
20
30
40
50
60
70
Populus_trichocarpa
Elaeis_guineensis
Ophiopogon_japonicus
Cichorium_intybus
Camptotheca_acuminata
Cunninghamia_lanceolata
Actinidia_deliciosa
Lens_culinaris
Quercus_robur
Catharanthus_roseus
Olea_europaea
Kosteletzkya_virginica
Trapa_bispinosa
Aristolochia_fimbriata
Gevuina_avellana
Ficus_elastica
Citrus_clementina
Paeonia_lactiflora
Nicotiana_tabacum
Vitis_pseudoreticulata
Conservation

**HG066**

POPTR_0009s15140

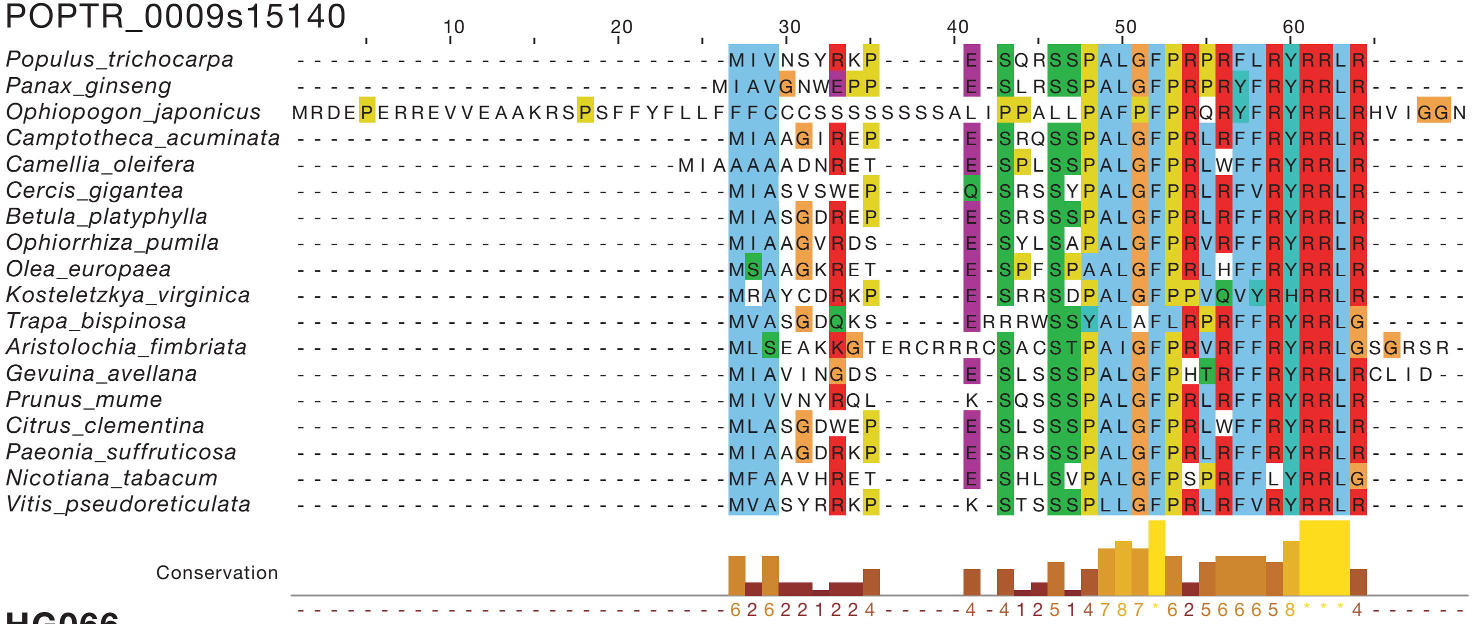


**HG066**

VIT_06s0061g01250

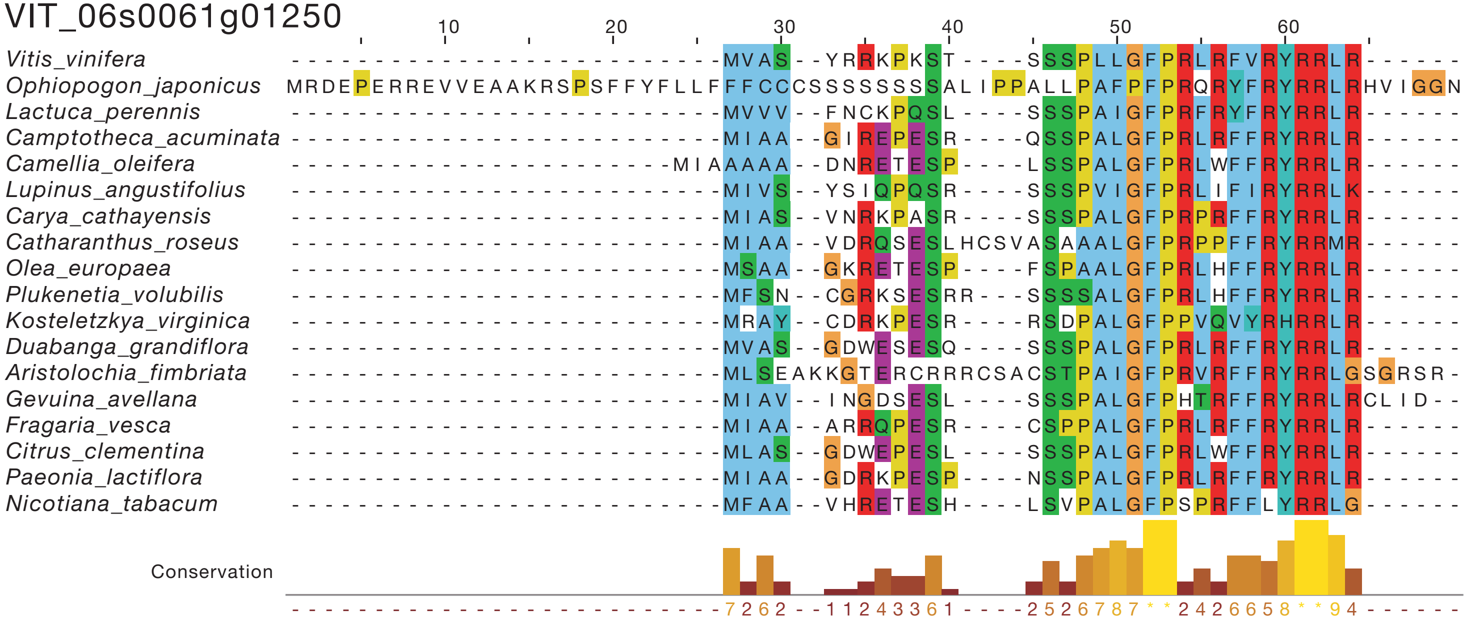

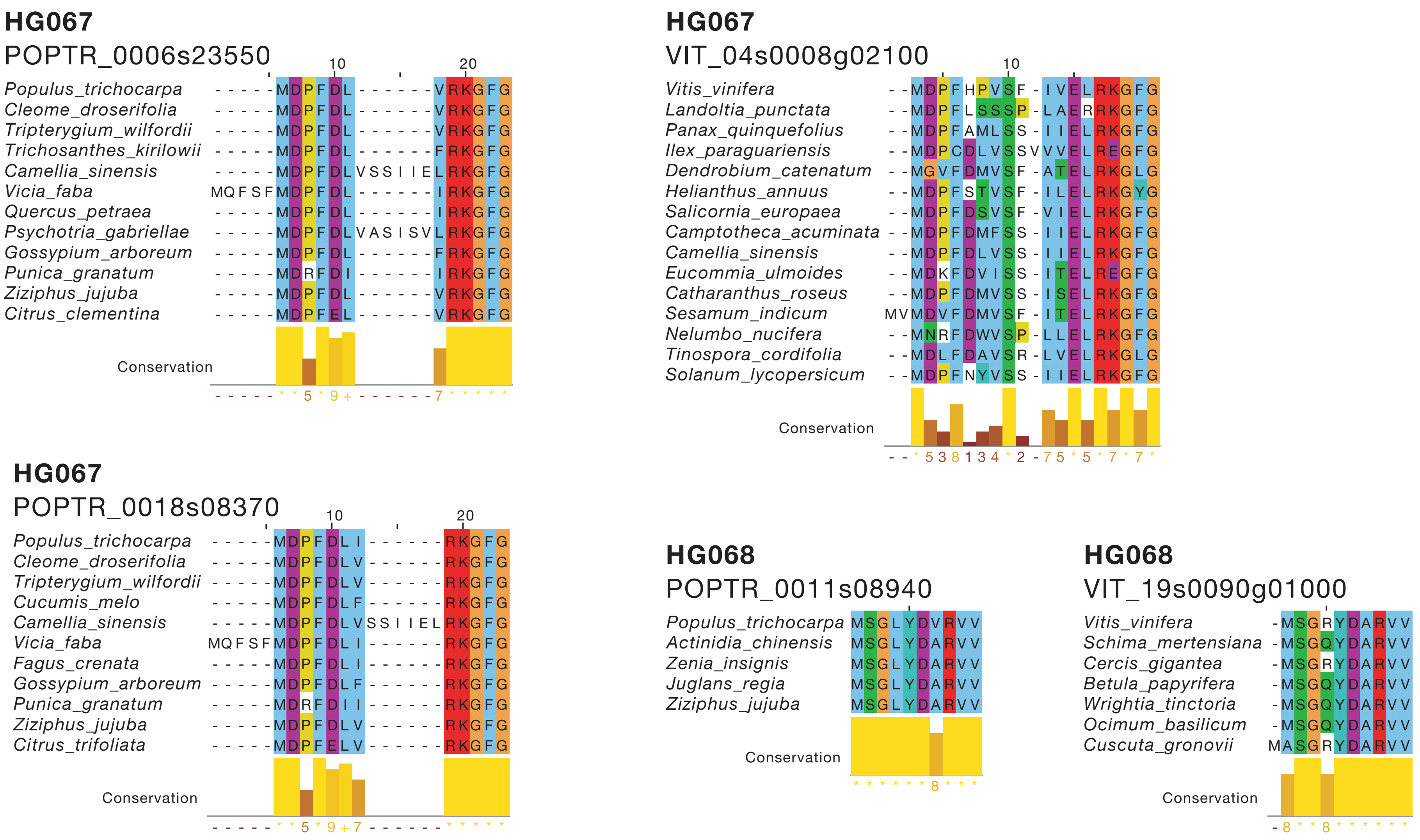
HG067
POPTR_0006s23550
Populus_trichocarpa
Cleome_droserifolia
Tripterygium_wilfordii
Trichosanthes_kirilowii
Camellia_sinensis
Vicia_faba
Quercus_petraea
Psychotria_gabriellae
Gossypium_arboreum
Punica_granatum
Ziziphus_jujuba
Citrus_clementina
Conservation
HG067
VIT_04s0008g02100
Vitis_vinifera
Landoltia_punctata
Panax_quinquefolius
Ilex_paraguariensis
Dendrobium_catenatum
Helianthus_annuus
Salicornia_europaea
Camptotheca_acuminata
Camellia_sinensis
Eucommia_ulmoides
Catharanthus_roseus
Sesamum_indicum
Nelumbo_nucifera
Tinospora_cordifolia
Solanum_lycopersicum
Conservation
HG067
POPTR_0018s08370
Populus_trichocarpa
Cleome_droserifolia
Tripterygium_wilfordii
Cucumis_melo
Camellia_sinensis
Vicia_faba
Fagus_crenata
Gossypium_arboreum
Punica_granatum
Ziziphus_jujuba
Citrus_trifoliata
Conservation
HG068
POPTR_0011s08940
Populus_trichocarpa
Actinidia_chinensis
Zenia_insignis
Juglans_regia
Ziziphus_jujuba
Conservation
HG068
VIT_19s0090g01000
Vitis_vinifera
Schima_mertensiana
Cercis_gigantea
Betula_papyrifera
Wrightia_tinctoria
Ocimum_basilicum
Cuscuta_gronovii
Conservation

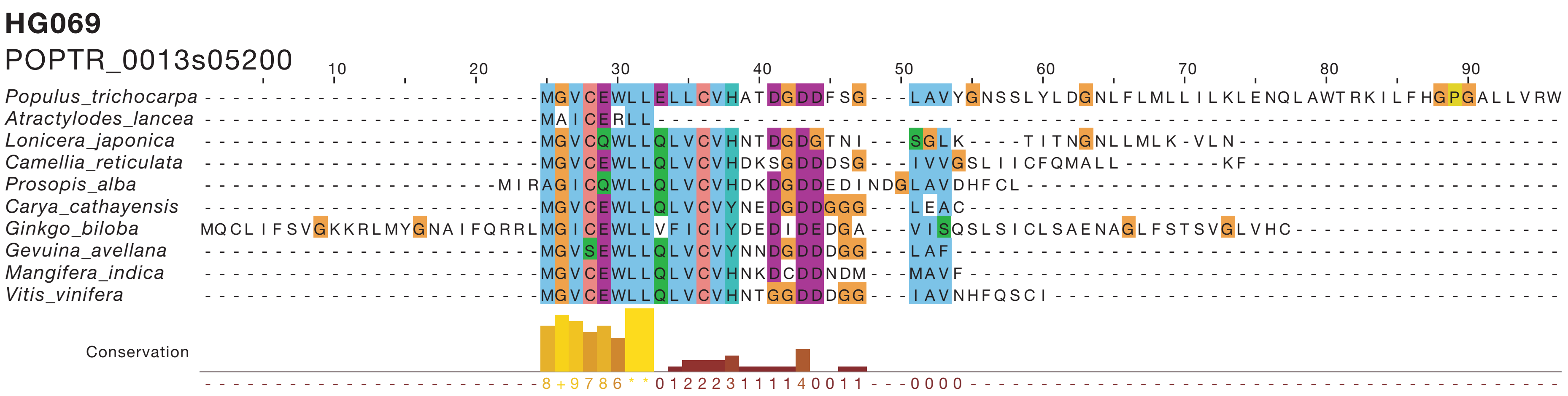
HG069
POPTR_0013s05200
Populus_trichocarpa
Atractylodes_lancea
Lonicera_japonica
Camellia_reticulata
Prosopis_alba
Carya_cathayensis
Ginkgo_biloba
Gevuina_avellana
Mangifera_indica
Vitis_vinifera
Conservation

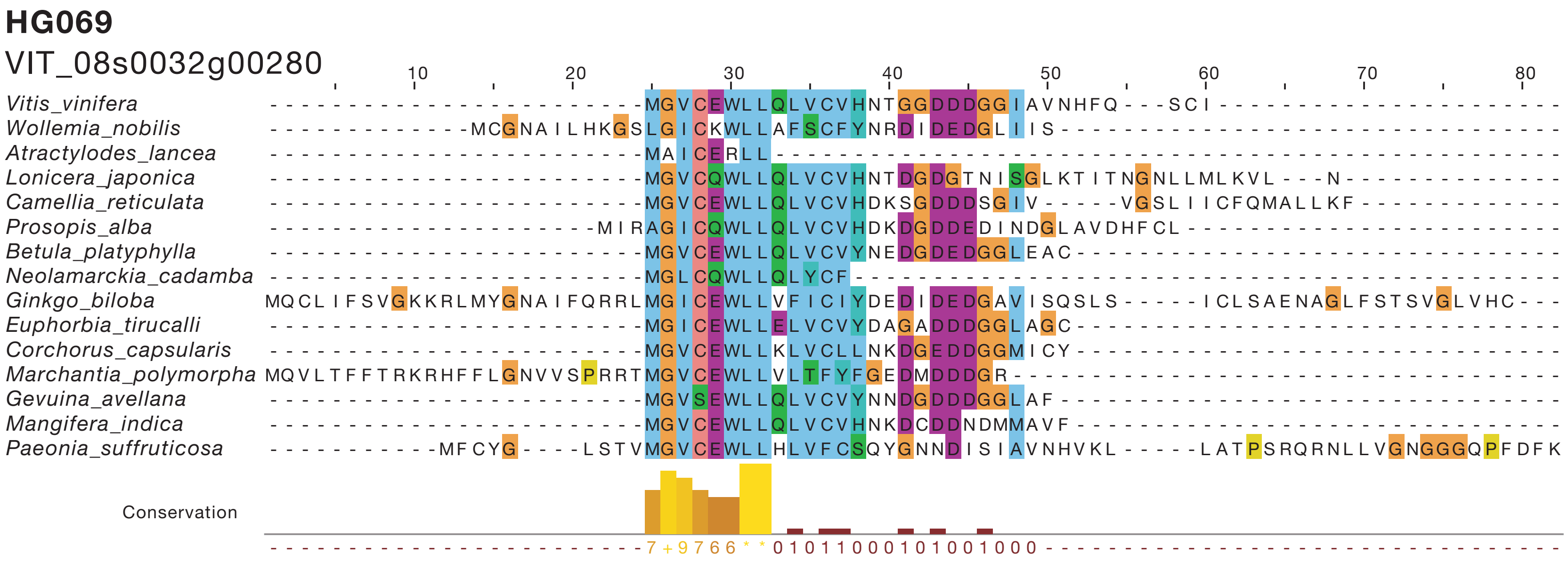
HG069
VIT_08s0032g00280
Vitis_vinifera
Wollemia_nobilis
Atractylodes_lancea
Lonicera_japonica
Camellia_reticulata
Prosopis_alba
Betula_platyphylla
Neolamarckia_cadamba
Ginkgo_biloba
Euphorbia_tirucalli
Corchorus_capsularis
Marchantia_polymorpha
Gevuina_avellana
Mangifera_indica
Paeonia_suffruticosa
Conservation

## HG070

AT3G52490

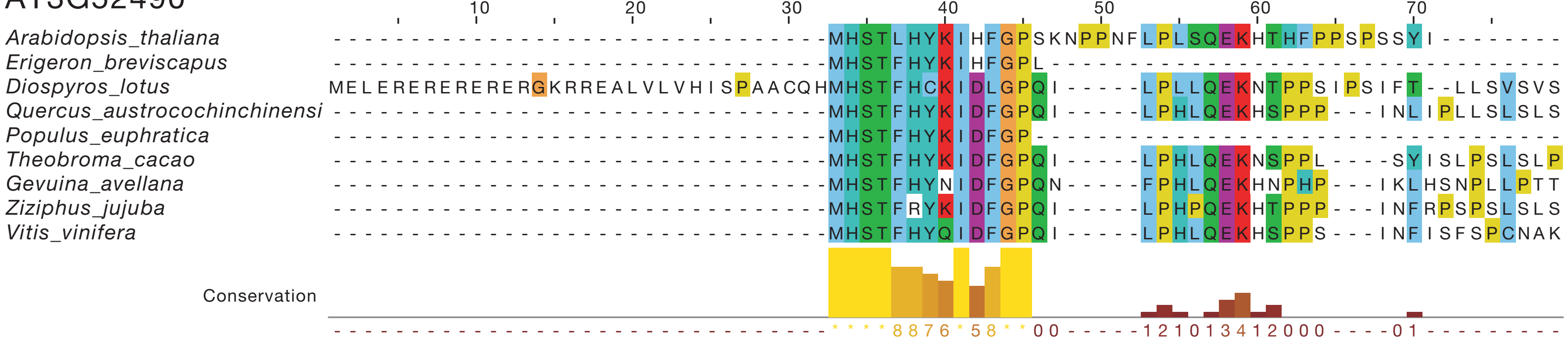


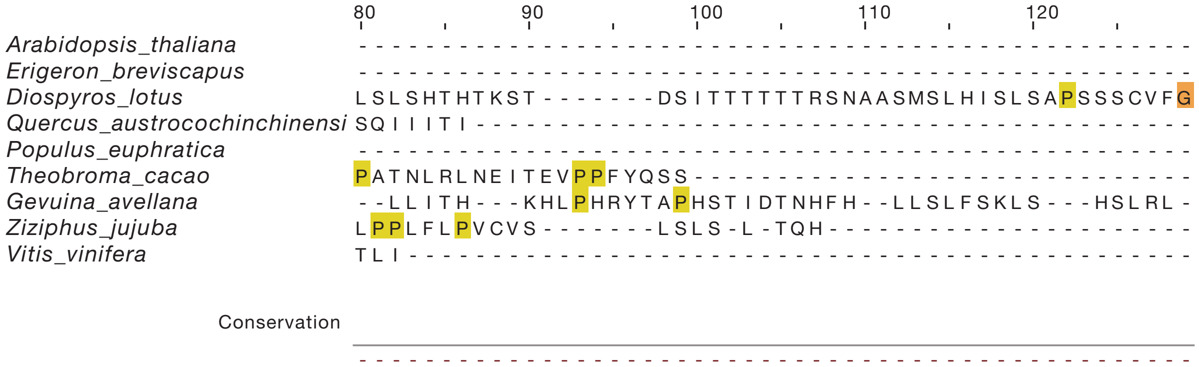


## HG071

AT3G61970

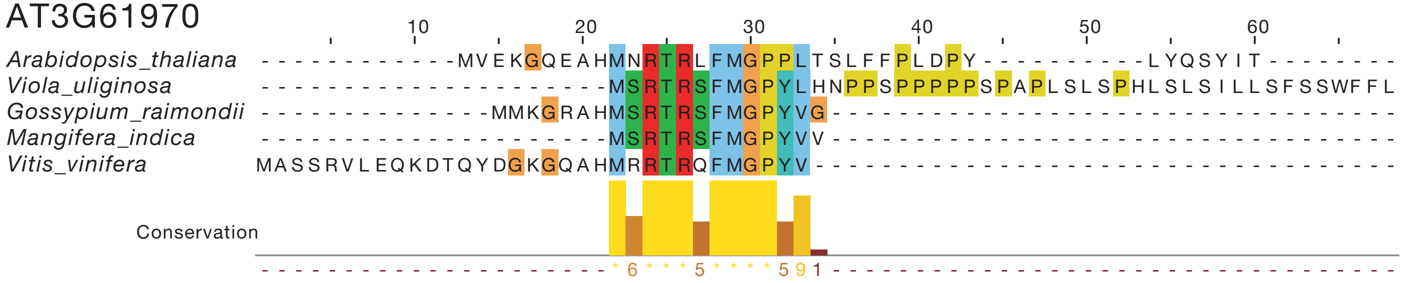

## HG072.1
AT4G34610

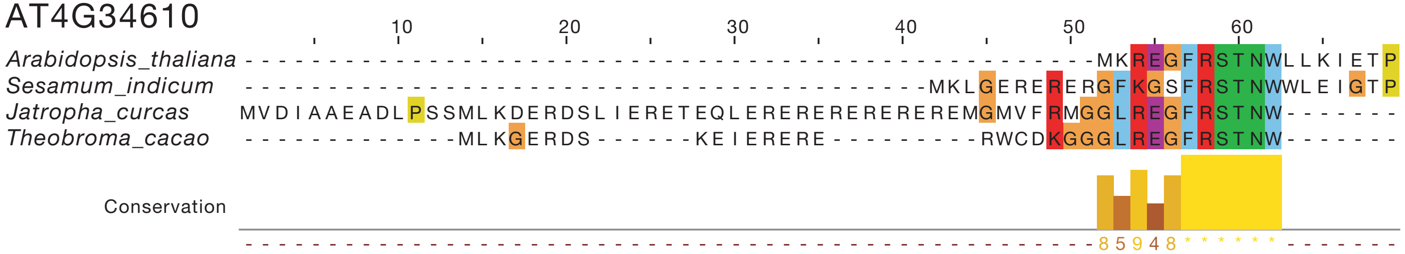


## HG072.2
VIT_03s0038g00050

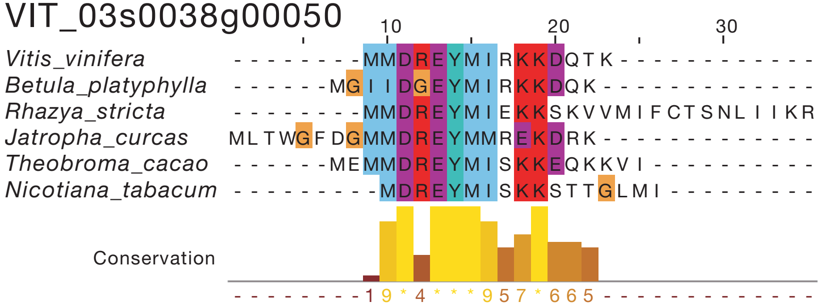


## HG073
AT5G53660

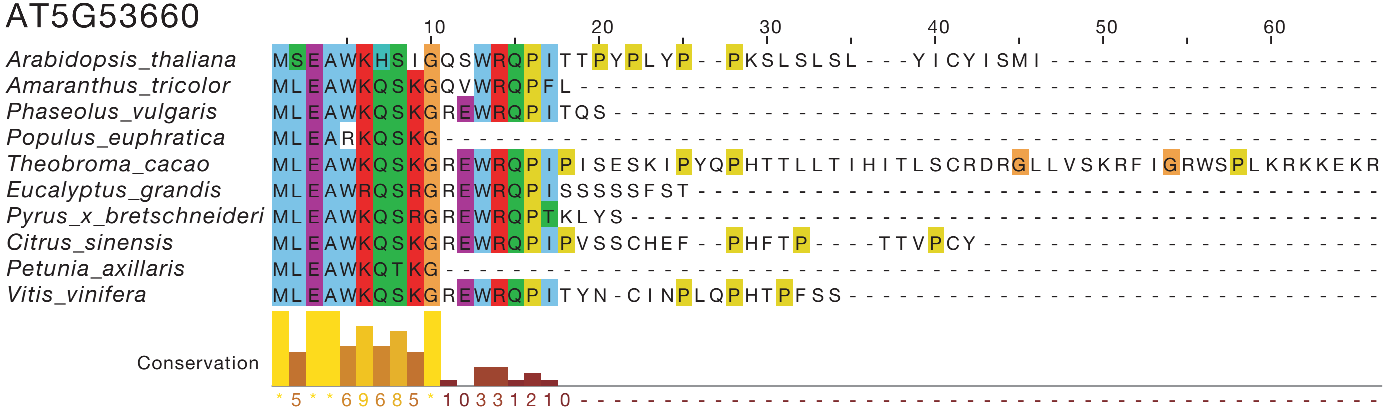

## HG074

LOC_Os04g36058

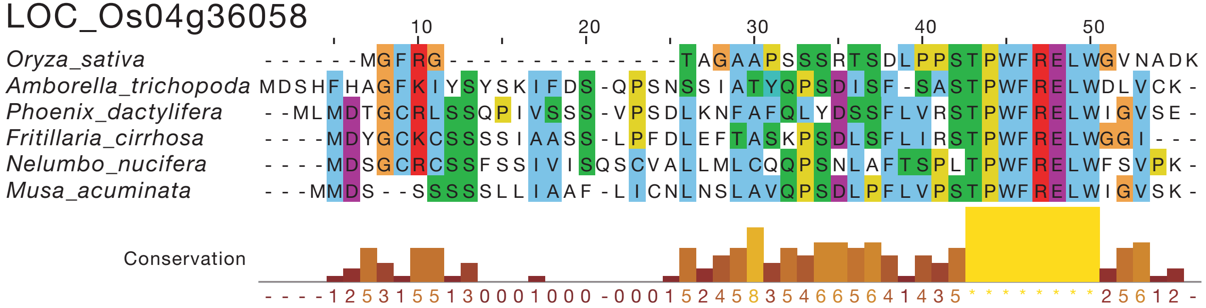


## HG075

LOC_Os11g16280

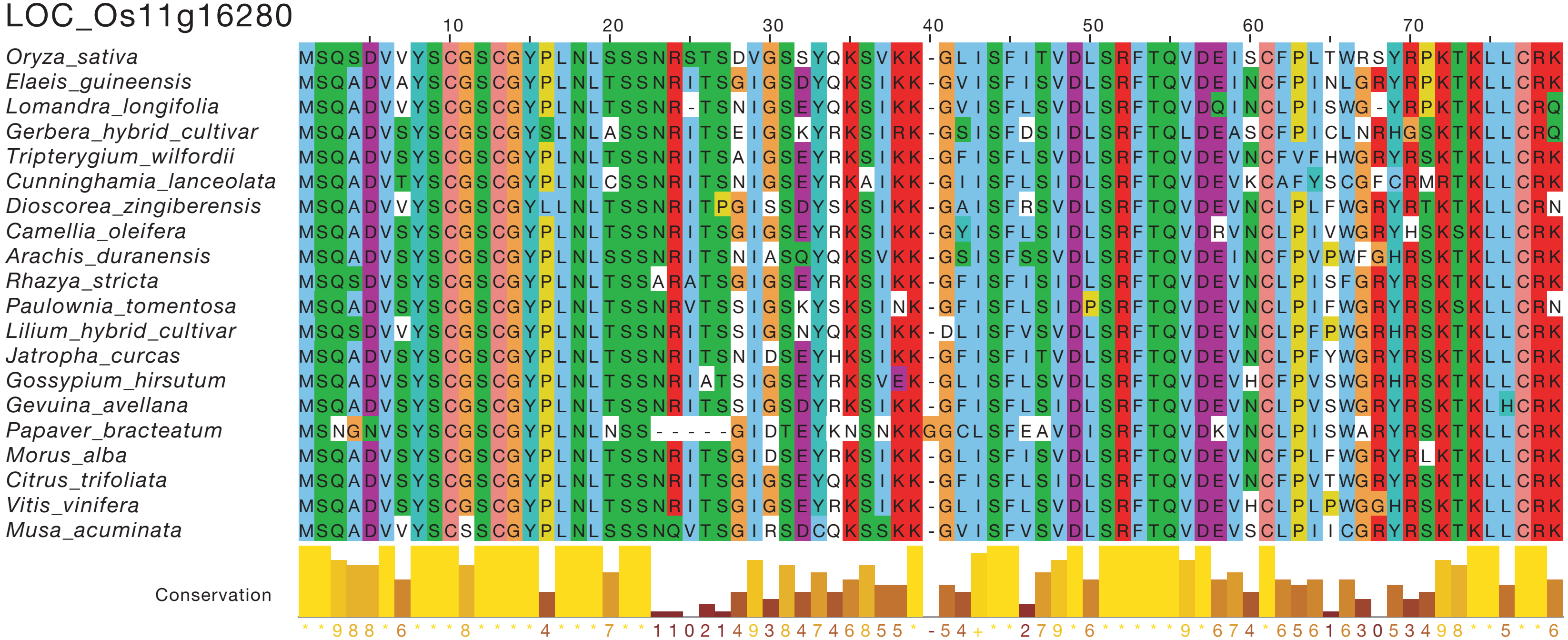

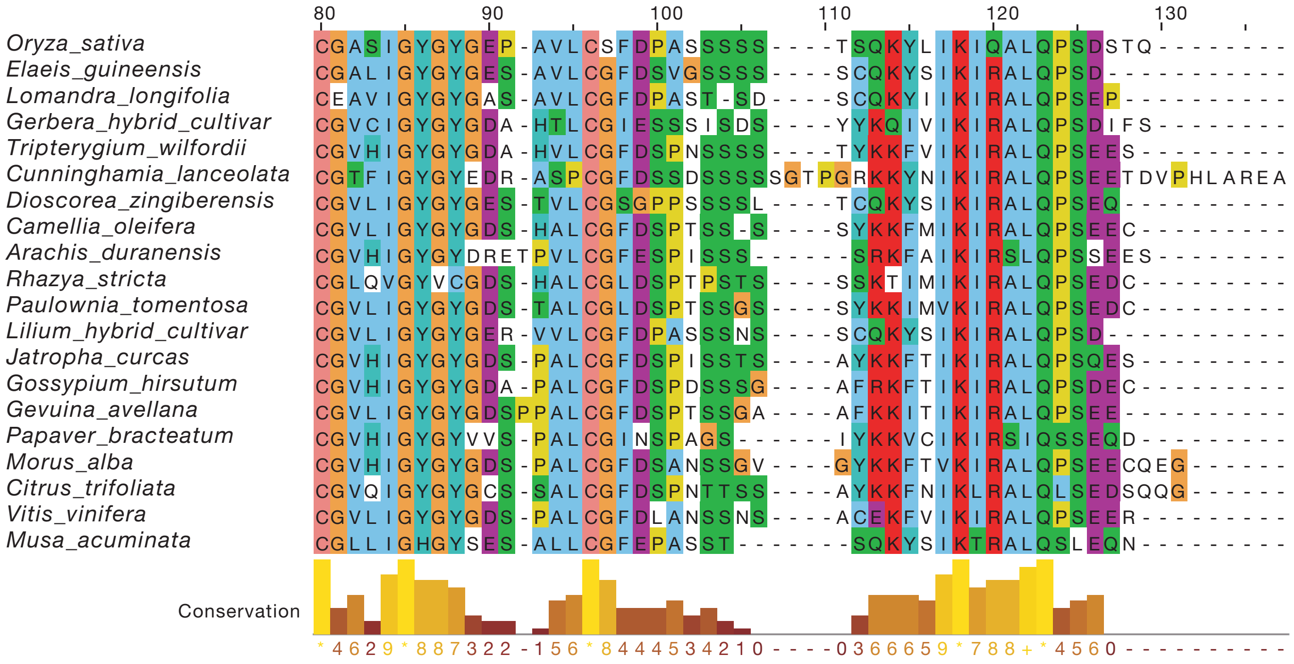


**HG076**

Solyc02g091660.2

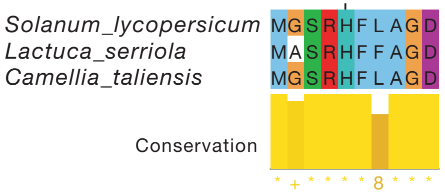


**HG077**

Solyc03g118620.2

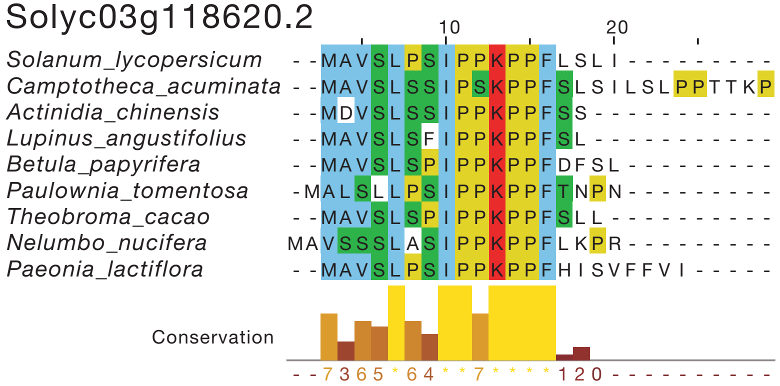

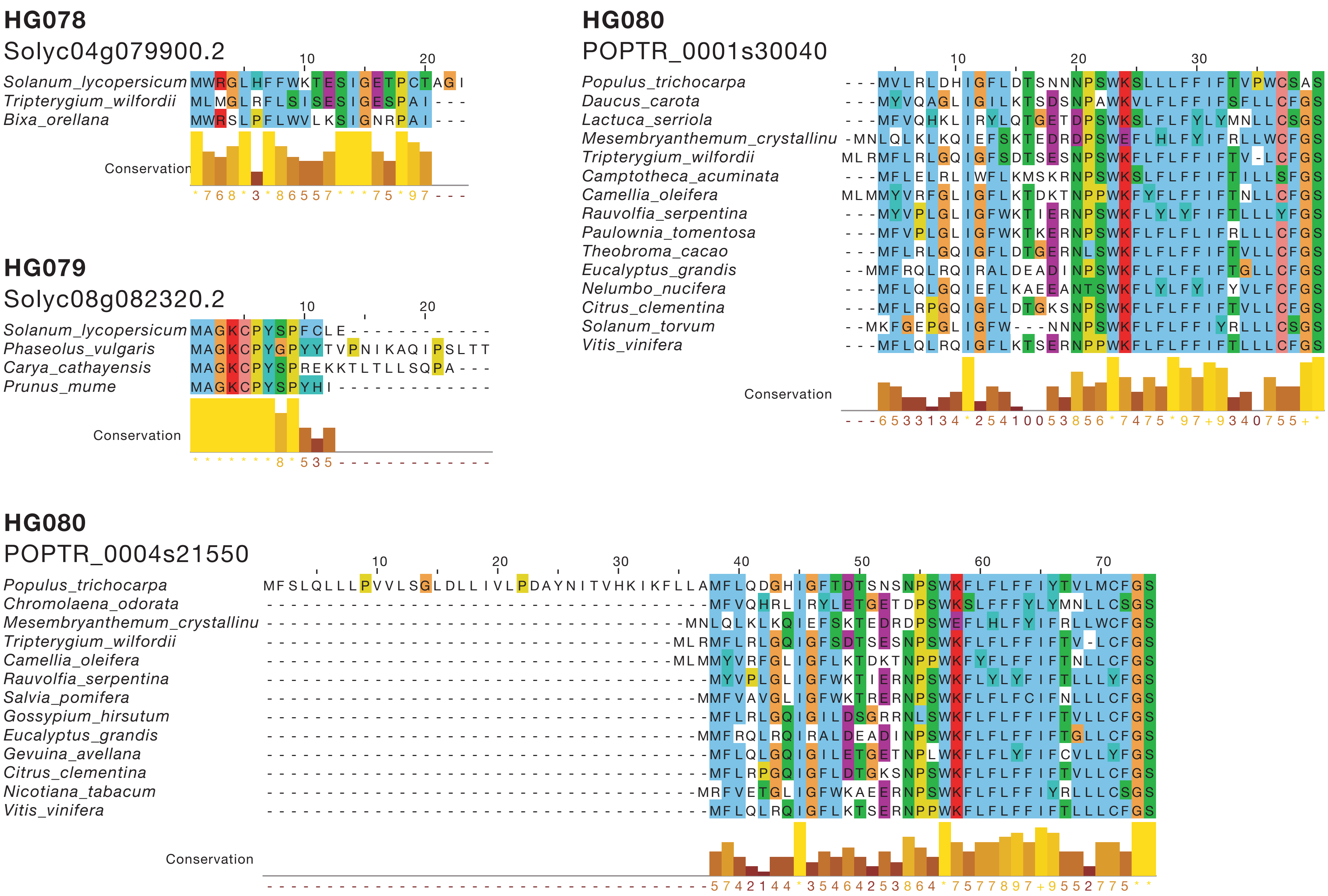
HG078
Solyc04g079900.2
Solanum_lycopersicum
Tripterygium_wilfordii
Bixa_orellana
Conservation
HG079
Solyc08g082320.2
Solanum_lycopersicum
Phaseolus_vulgaris
Carya_cathayensis
Prunus_mume
Conservation
HG080
POPTR_0001s30040
Populus_trichocarpa
Daucus_carota
Lactuca_serriola
Mesembryanthemum_crystallinu
Tripterygium_wilfordii
Camptotheca_acuminata
Camellia_oleifera
Rauvolfia_serpentina
Paulownia_tomentosa
Theobroma_cacao
Eucalyptus_grandis
Nelumbo_nucifera
Citrus_clementina
Solanum_torvum
Vitis_vinifera
Conservation
HG080
POPTR_0004s21550
Populus_trichocarpa
Chromolaena_odorata
Mesembryanthemum_crystallinu
Tripterygium_wilfordii
Camellia_oleifera
Rauvolfia_serpentina
Salvia_pomifera
Gossypium_hirsutum
Eucalyptus_grandis
Gevuina_avellana
Citrus_clementina
Nicotiana_tabacum
Vitis_vinifera
Conservation

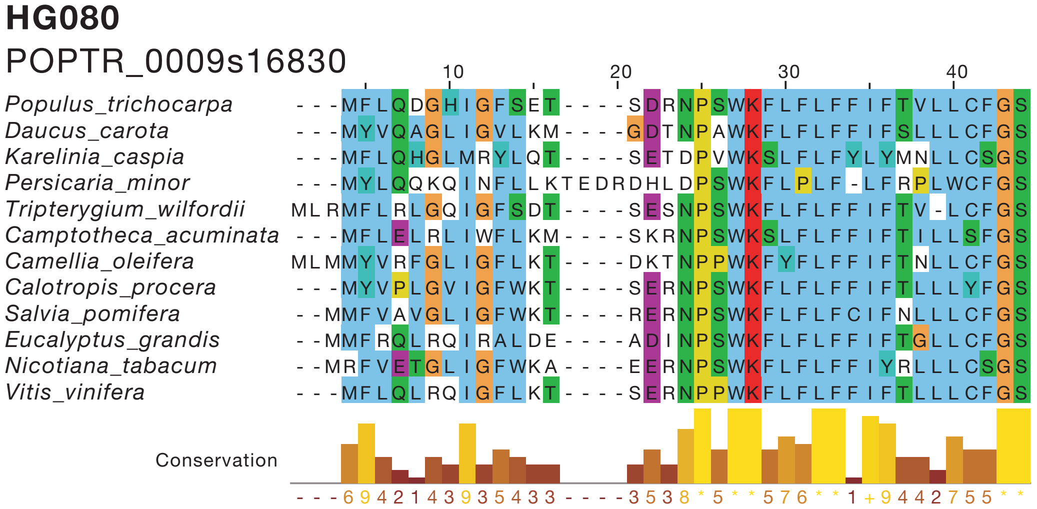
HG080
POPTR_0009s16830
Populus_trichocarpa
Daucus_carota
Karelinia_caspia
Persicaria_minor
Tripterygium_wilfordii
Camptotheca_acuminata
Camellia_oleifera
Calotropis_procera
Salvia_pomifera
Eucalyptus_grandis
Nicotiana_tabacum
Vitis_vinifera
Conservation

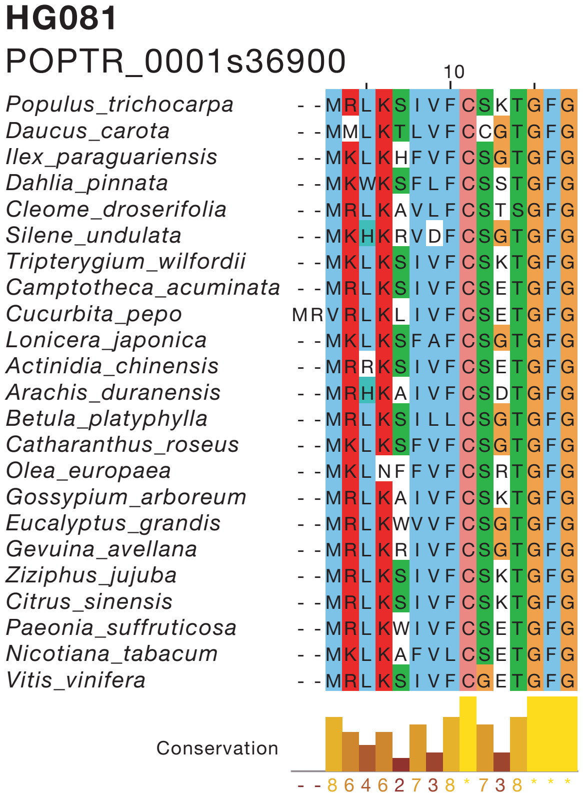
HG081
POPTR_0001s36900
Populus_trichocarpa
Daucus_carota
Ilex_paraguariensis
Dahlia_pinnata
Cleome_droserifolia
Silene_undulata
Tripterygium_wilfordii
Camptotheca_acuminata
Cucurbita_pepo
Lonicera_japonica
Actinidia_chinensis
Arachis_duranensis
Betula_platyphylla
Catharanthus_roseus
Olea_europaea
Gossypium_arboreum
Eucalyptus_grandis
Gevuina_avellana
Ziziphus_jujuba
Citrus_sinensis
Paeonia_suffruticosa
Nicotiana_tabacum
Vitis_vinifera
Conservation

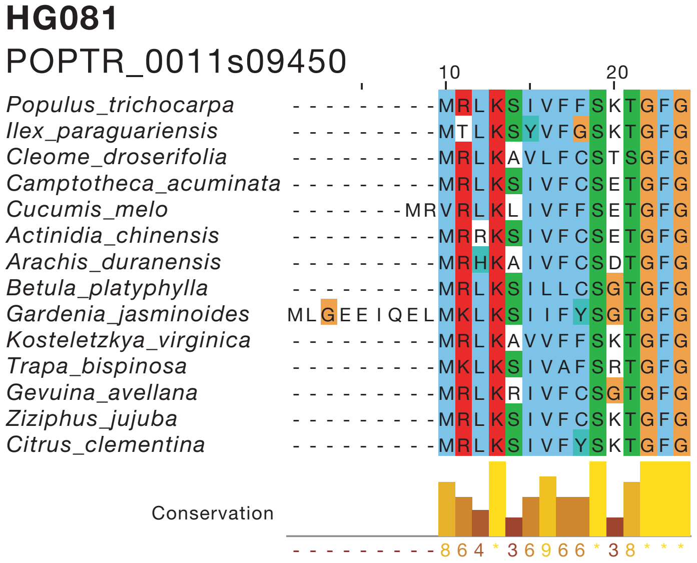
HG081
POPTR_0011s09450
Populus_trichocarpa
Ilex_paraguariensis
Cleome_droserifolia
Camptotheca_acuminata
Cucumis_melo
Actinidia_chinensis
Arachis_duranensis
Betula_platyphylla
Gardenia_jasminoides
Kosteletzkya_virginica
Trapa_bispinosa
Gevuina_avellana
Ziziphus_jujuba
Citrus_clementina
Conservation

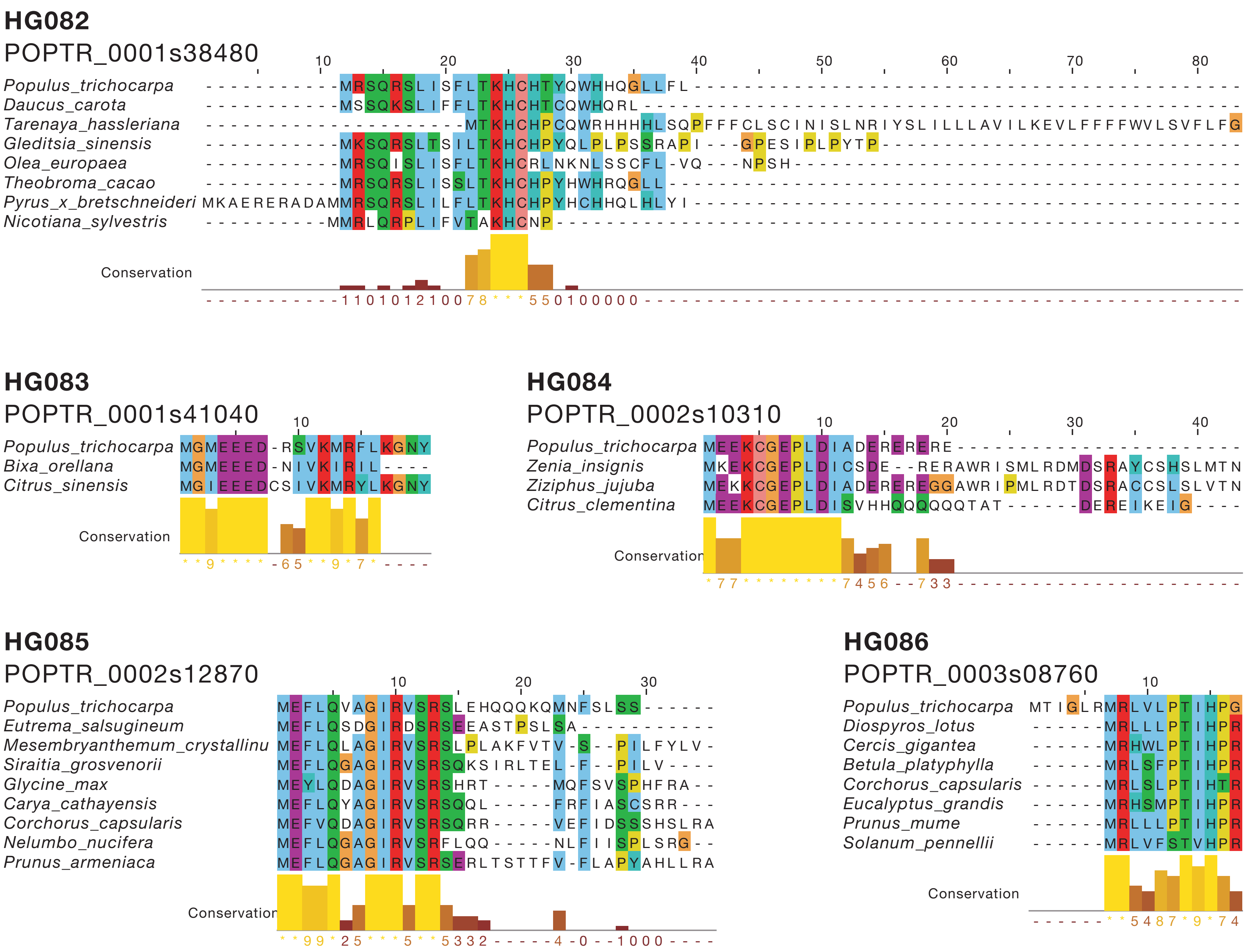
HG082
POPTR_0001s38480
Populus_trichocarpa
Daucus_carota
Tarenaya_hassleriana
Gleditsia_sinensis
Olea_europaea
Theobroma_cacao
Pyrus_x_bretschneideri
Nicotiana_sylvestris
Conservation
HG083
POPTR_0001s41040
Populus_trichocarpa
Bixa_orellana
Citrus_sinensis
Conservation
HG084
POPTR_0002s10310
Populus_trichocarpa
Zenia_insignis
Ziziphus_jujuba
Citrus_clementina
Conservation
HG085
POPTR_0002s12870
Populus_trichocarpa
Eutrema_salsugineum
Mesembryanthemum_crystallinu
Siraitia_grosvenorii
Glycine_max
Carya_cathayensis
Corchorus_capsularis
Nelumbo_nucifera
Prunus_armeniaca
Conservation
HG086
POPTR_0003s08760
Populus_trichocarpa
Diospyros_lotus
Cercis_gigantea
Betula_platyphylla
Corchorus_capsularis
Eucalyptus_grandis
Prunus_mume
Solanum_pennellii
Conservation

## HG087

POPTR_0004s03900

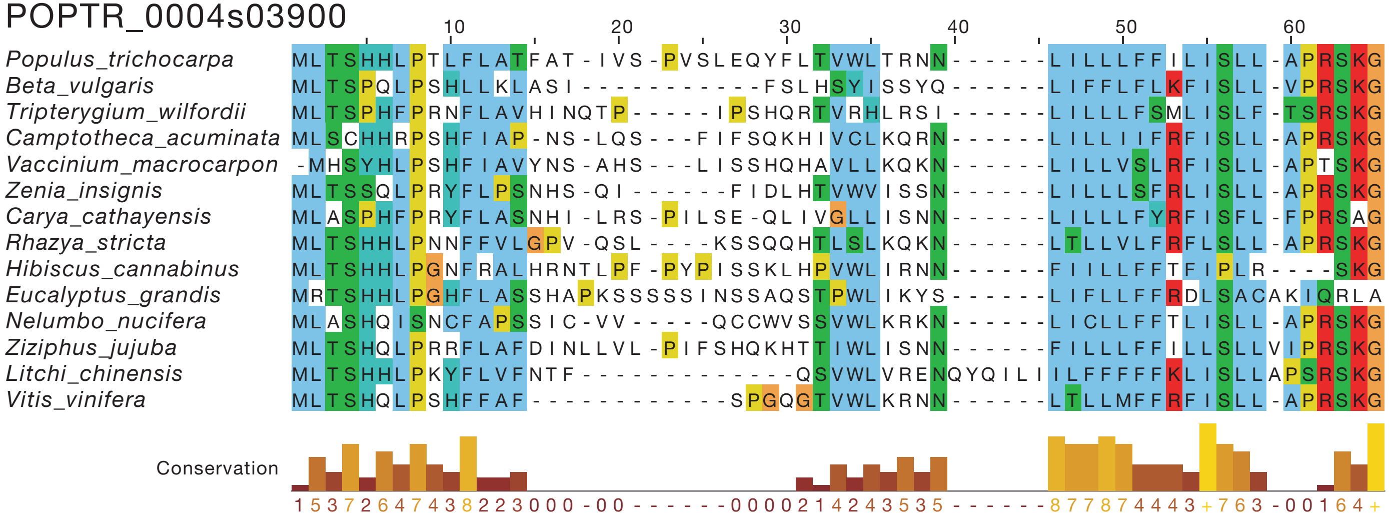


## HG087

POPTR_0011s04730

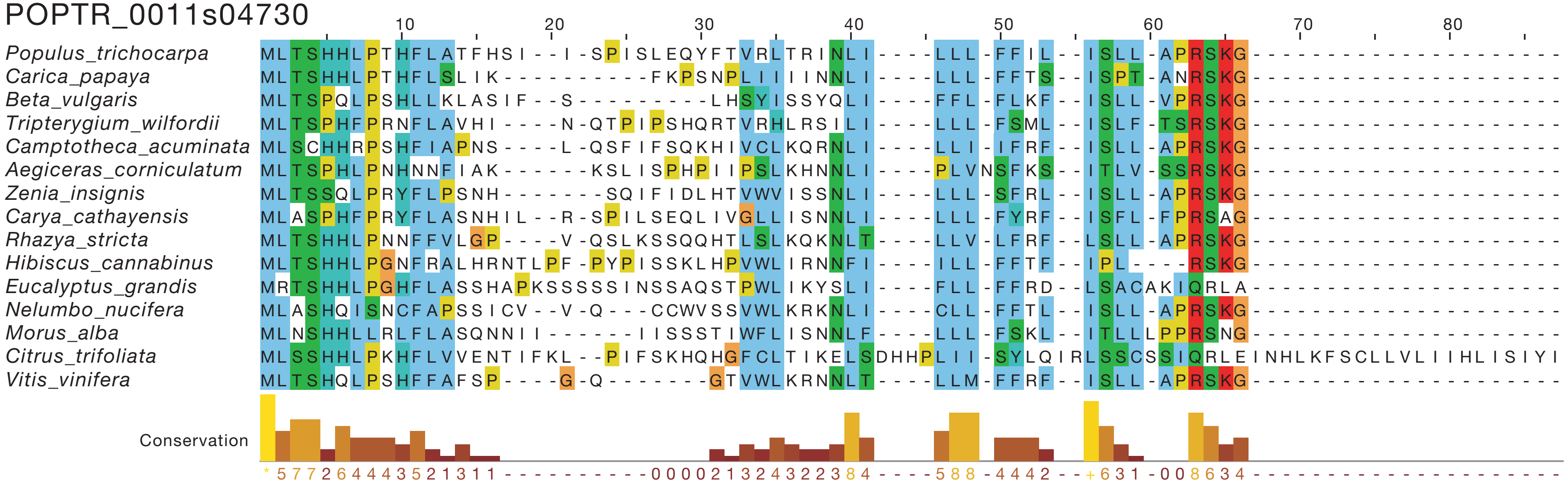

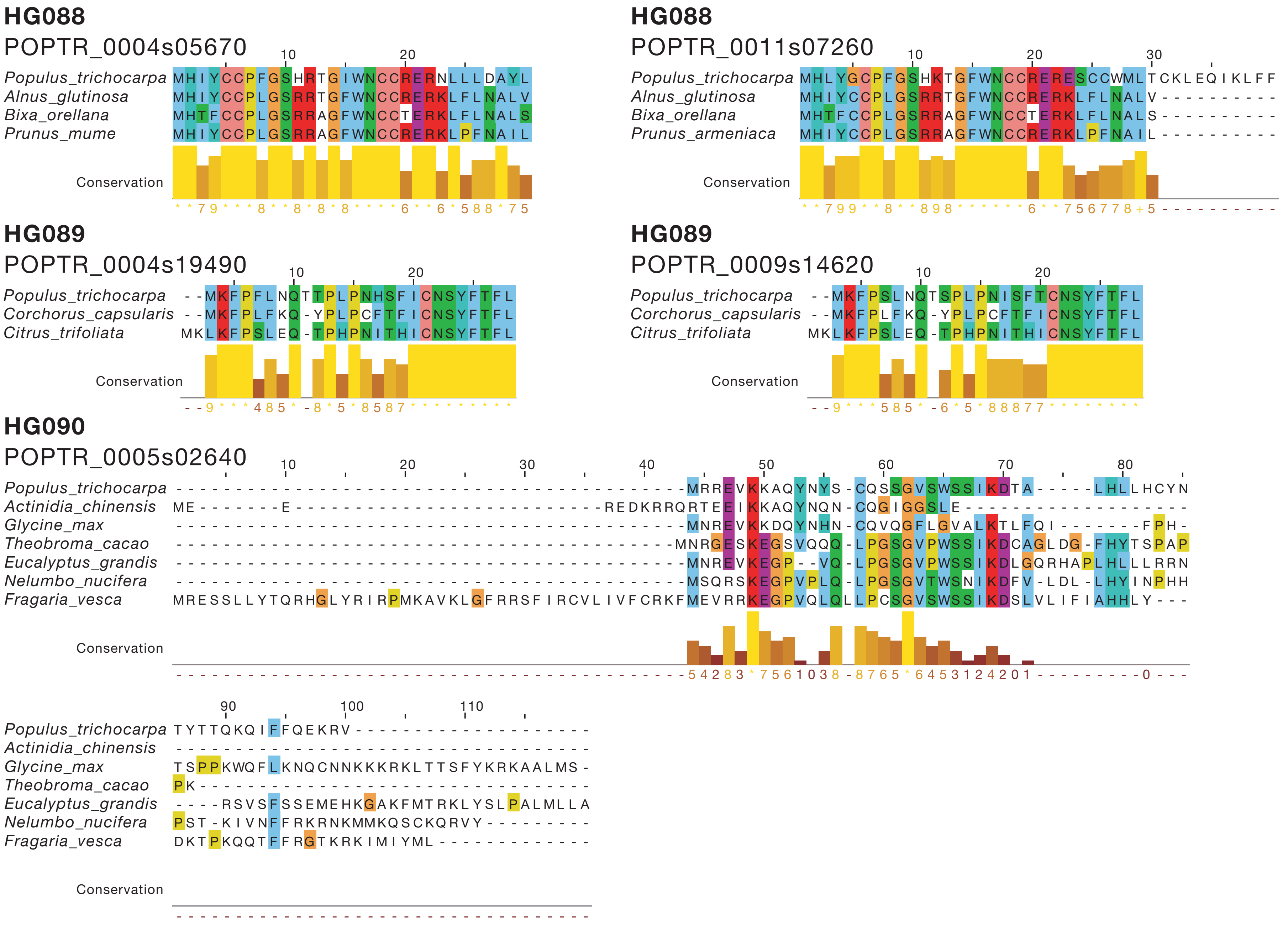

HG088
POPTR_0004s05670
Populus_trichocarpa
Alnus_glutinosa
Bixa_orellana
Prunus_mume
Conservation
HG088
POPTR_0011s07260
Populus_trichocarpa
Alnus_glutinosa
Bixa_orellana
Prunus_armeniaca
Conservation
HG089
POPTR_0004s19490
Populus_trichocarpa
Corchorus_capsularis
Citrus_trifoliata
Conservation
HG089
POPTR_0009s14620
Populus_trichocarpa
Corchorus_capsularis
Citrus_trifoliata
Conservation
HG090
POPTR_0005s02640
Populus_trichocarpa
Actinidia_chinensis
Glycine_max
Theobroma_cacao
Eucalyptus_grandis
Nelumbo_nucifera
Fragaria_vesca
Conservation

**HG091**

POPTR_0005s07290

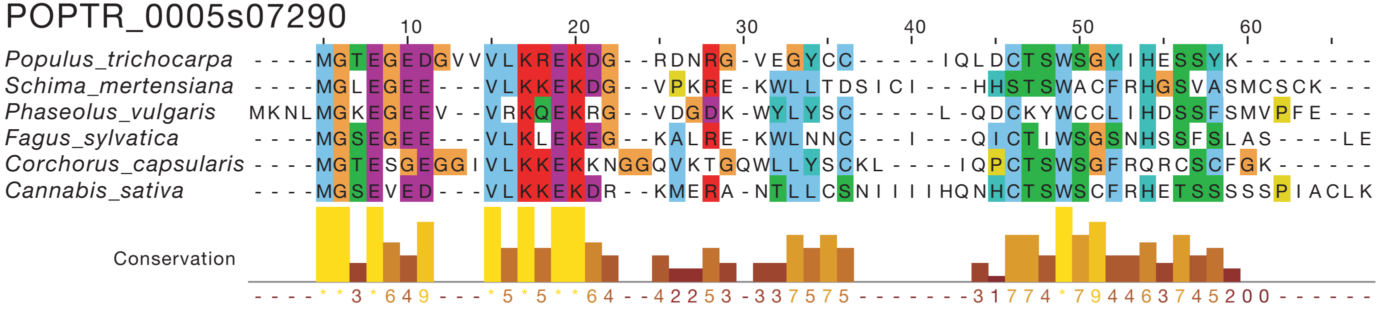


**HG091**

POPTR_0007s05010

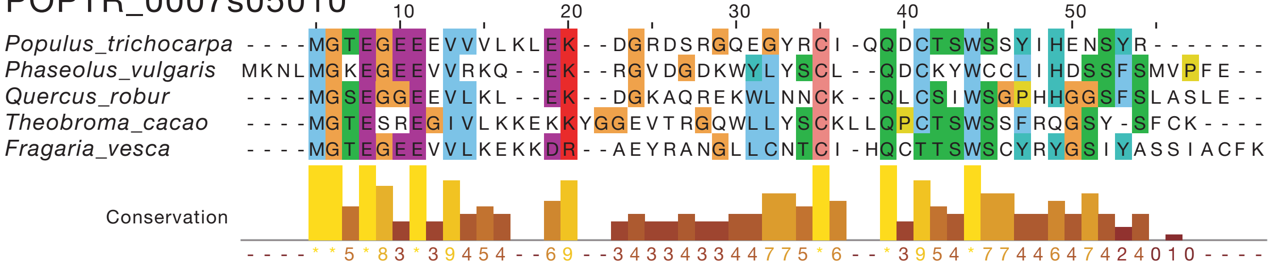

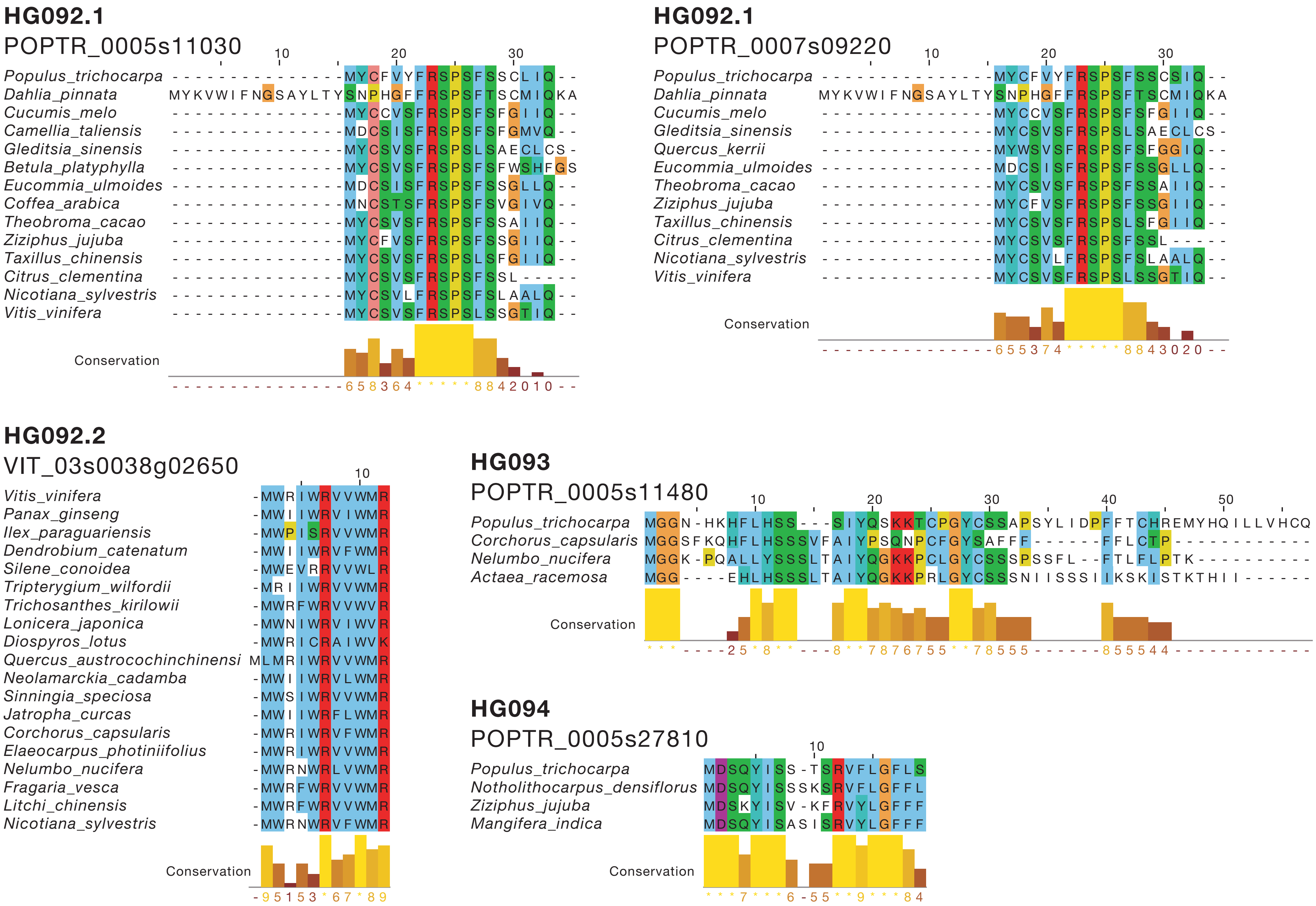
HG092.1
POPTR_0005s11030
Populus_trichocarpa
Dahlia_pinnata
Cucumis_melo
Camellia_taliensis
Gleditsia_sinensis
Betula_platyphylla
Eucommia_ulmoides
Coffea_arabica
Theobroma_cacao
Ziziphus_jujuba
Taxillus_chinensis
Citrus_clementina
Nicotiana_sylvestris
Vitis_vinifera
Conservation
HG092.1
POPTR_0007s09220
Populus_trichocarpa
Dahlia_pinnata
Cucumis_melo
Gleditsia_sinensis
Quercus_kerrii
Eucommia_ulmoides
Theobroma_cacao
Ziziphus_jujuba
Taxillus_chinensis
Citrus_clementina
Nicotiana_sylvestris
Vitis_vinifera
Conservation
HG092.2
VIT_03s0038g02650
Vitis_vinifera
Panax_ginseng
Ilex_paraguariensis
Dendrobium_catenatum
Silene_conoidea
Tripterygium_wilfordii
Trichosanthes_kirilowii
Lonicera_japonica
Diospyros_lotus
Quercus_austrocochinchinensi
Neolamarckia_cadamba
Sinningia_speciosa
Jatropha_curcas
Corchorus_capsularis
Elaeocarpus_photiniifolius
Nelumbo_nucifera
Fragaria_vesca
Litchi_chinensis
Nicotiana_sylvestris
Conservation
HG093
POPTR_0005s11480
Populus_trichocarpa
Corchorus_capsularis
Nelumbo_nucifera
Actaea_racemosa
Conservation
HG094
POPTR_0005s27810
Populus_trichocarpa
Notholithocarpus_densiflorus
Ziziphus_jujuba
Mangifera_indica
Conservation

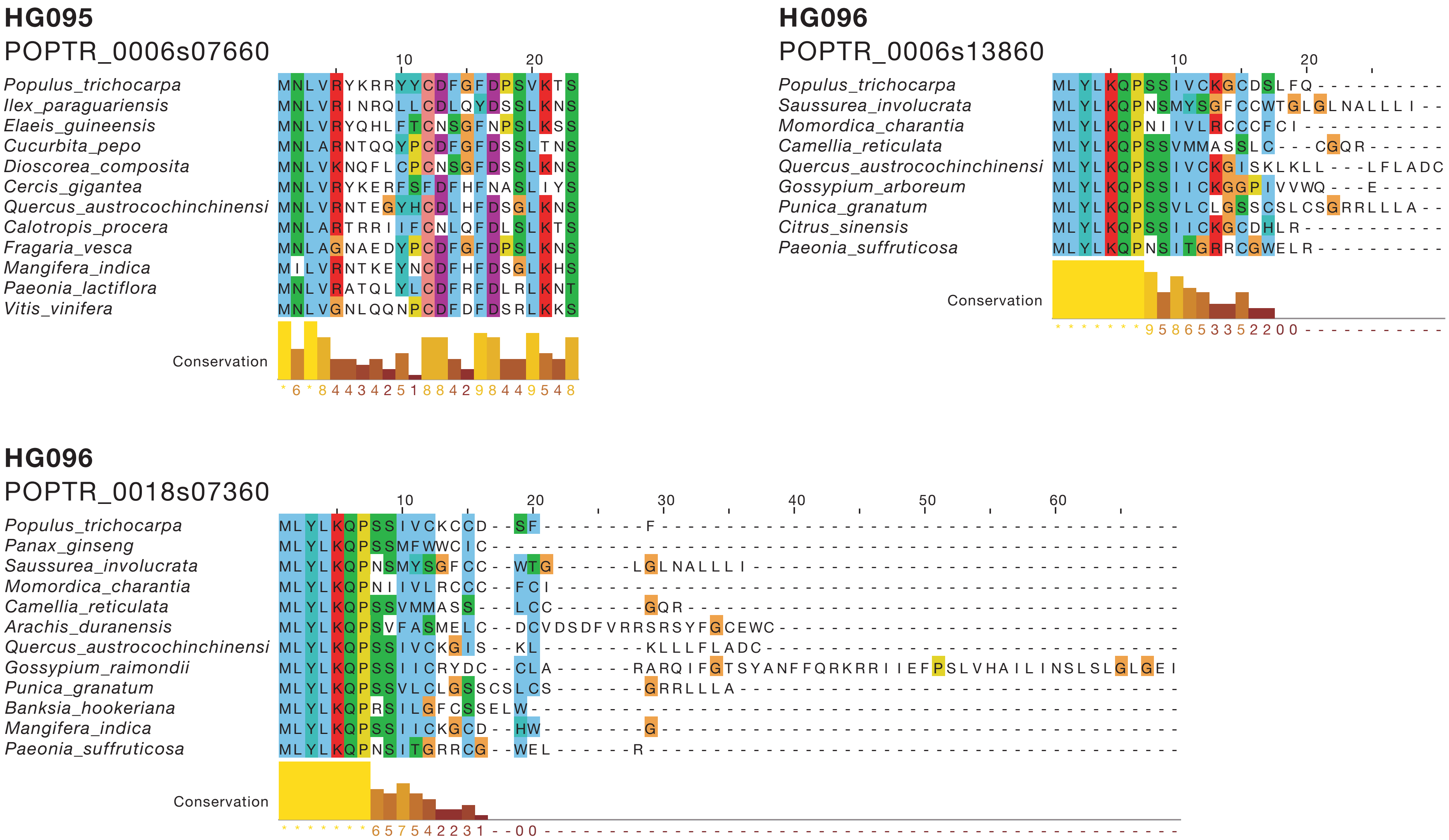
HG095
POPTR_0006s07660
Populus_trichocarpa
Ilex_paraguariensis
Elaeis_guineensis
Cucurbita_pepo
Dioscorea_composita
Cercis_gigantea
Quercus_austrocochinchinensi
Calotropis_procera
Fragaria_vesca
Mangifera_indica
Paeonia_lactiflora
Vitis_vinifera
Conservation
HG096
POPTR_0006s13860
Populus_trichocarpa
Saussurea_involucrata
Momordica_charantia
Camellia_reticulata
Quercus_austrocochinchinensi
Gossypium_arboreum
Punica_granatum
Citrus_sinensis
Paeonia_suffruticosa
Conservation
HG096
POPTR_0018s07360
Populus_trichocarpa
Panax_ginseng
Saussurea_involucrata
Momordica_charantia
Camellia_reticulata
Arachis_duranensis
Quercus_austrocochinchinensi
Gossypium_raimondii
Punica_granatum
Banksia_hookeriana
Mangifera_indica
Paeonia_suffruticosa
Conservation

## HG097

POPTR_0006s14740

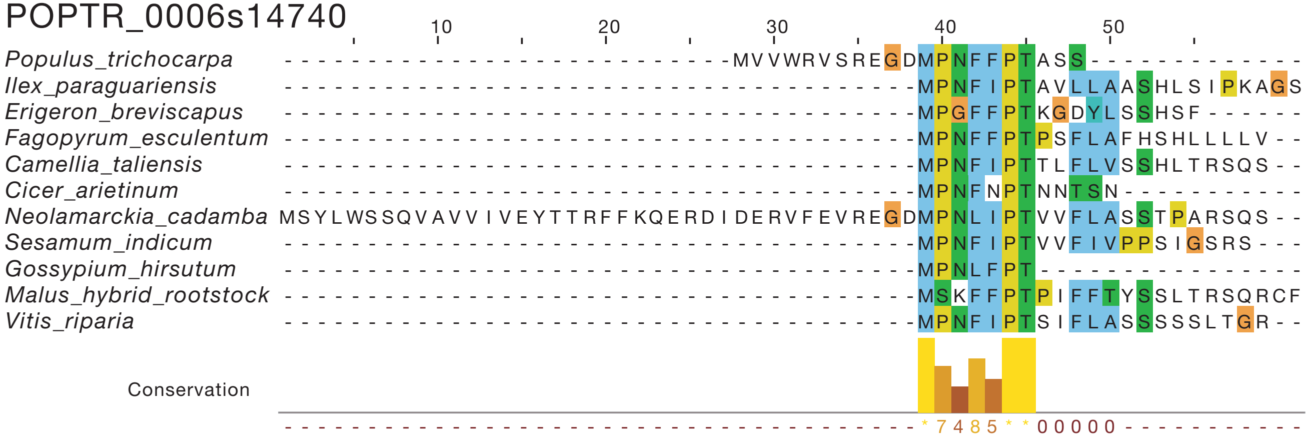


## HG098

POPTR_0006s22890

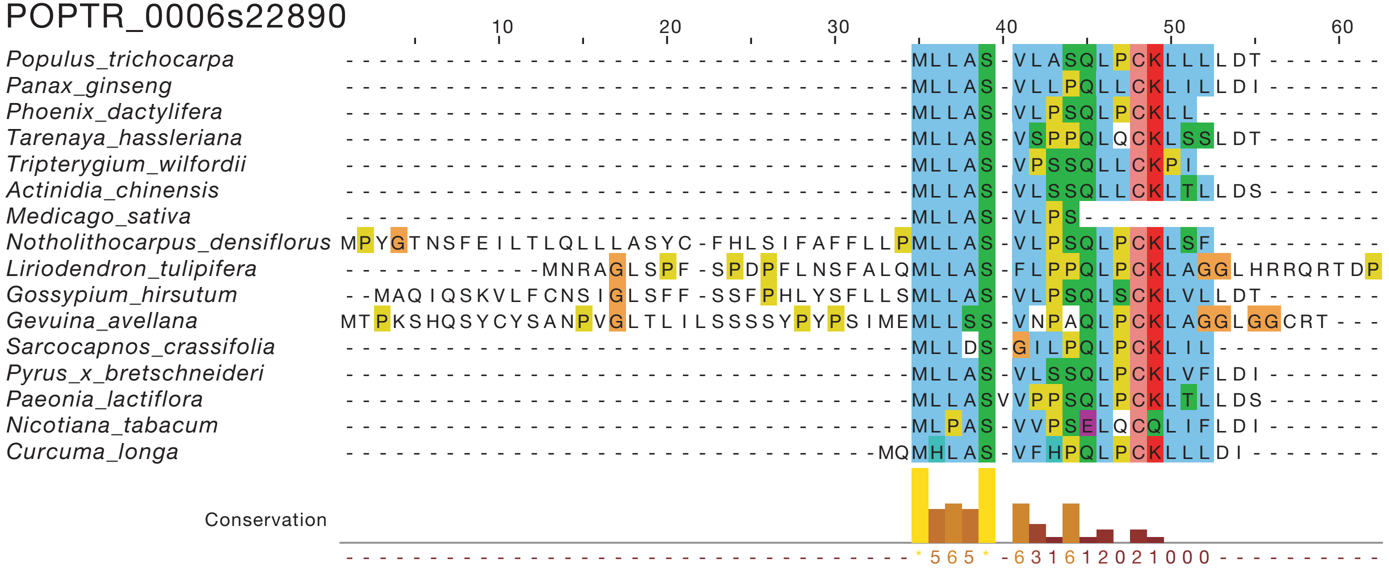

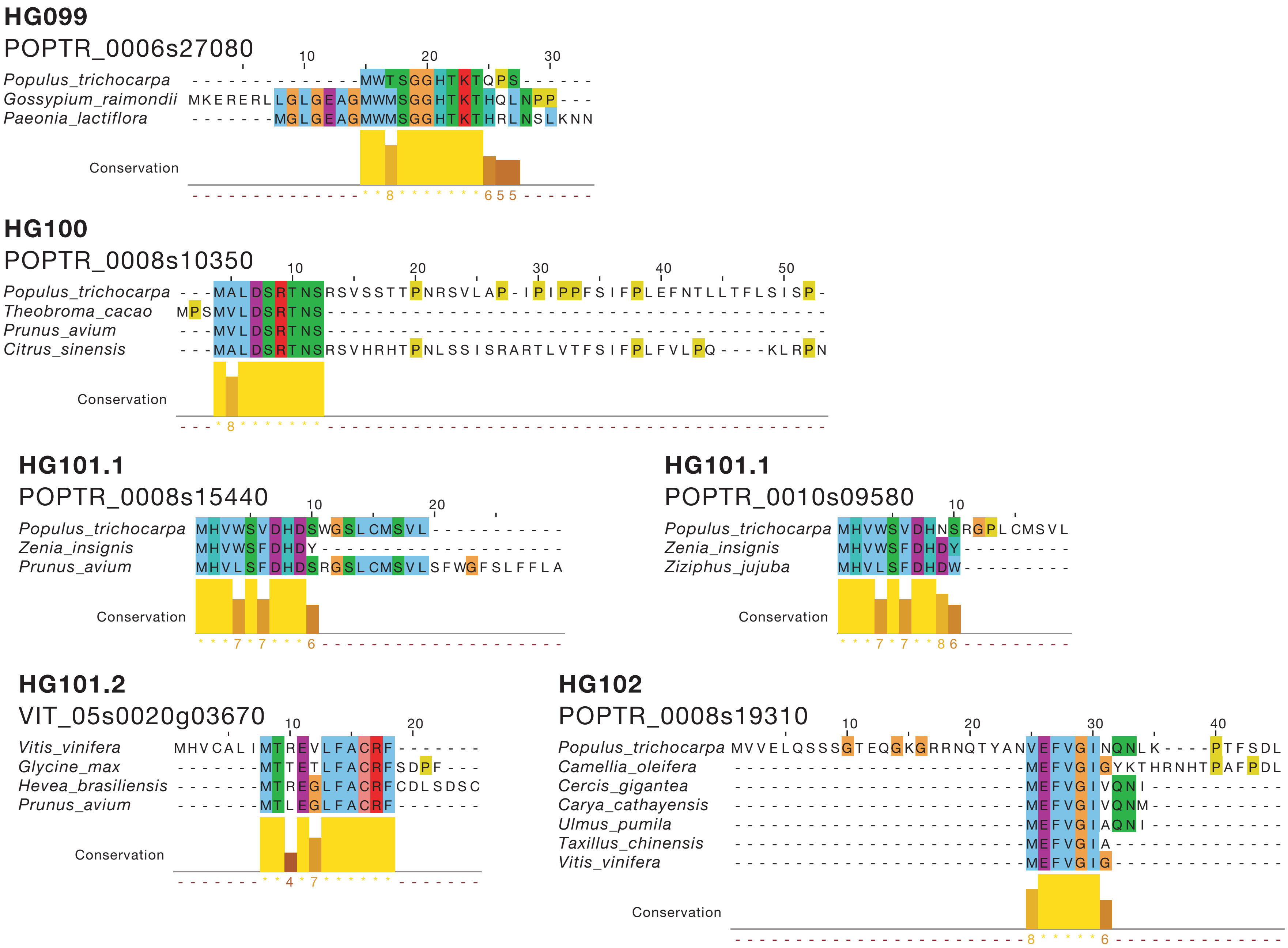
HG099
POPTR_0006s27080
Populus_trichocarpa - - - - - - - - - - - - - - MWTSGGHTKTQPS - - - - - -
Gossypium_raimondii MKERERLLGLGEAGMWMSGGHTKTHQLNPP - - -
Paeonia_lactiflora - - - - - - - MGLGEAGMWMSGGHTKTHRLNSLKNN
Conservation
* * 8 * * * * * * * 6 5 5
HG100
POPTR_0008s10350
Populus_trichocarpa - - - MALDSRTNSRSVSSTTPNRSVLAP - IPIPPFSIFPLEFNTLLTFLSISP -
Theobroma_cacao MPSMVLDSRTNS
Prunus_avium - - - MVLDSRTNS
Citrus_sinensis - - - MALDSRTNSRSVHRHTPNLSSISRARTLVTFSIFPLFVLPQ - - - - KLRPN
Conservation
* 8 * * * * * * *
HG101.1
POPTR_0008s15440
Populus_trichocarpa MHVWSVDHDSWGSLCMSVL
Zenia_insignis MHVWSFDHDY
Prunus_avium MHVLSFDHDSRGSLCMSVLSFWGFSLFFLA
Conservation
* * * 7 * 7 * * * 6
HG101.1
POPTR_0010s09580
Populus_trichocarpa MHVWSVDHNSRGPLCMSVL
Zenia_insignis MHVWSFDHDY
Ziziphus_jujuba MHVLSFDHDW
Conservation
* * * 7 * 7 * * 8 6
HG101.2
VIT_05s0020g03670
Vitis_vinifera MHVCALIMTREVLFACRF
Glycine_max - - - - - - - MTTETLFACRFSDPF - - -
Hevea_brasiliensis - - - - - - - MTREGLFACRFCDLSDSC
Prunus_avium - - - - - - - MTLEGLFACRF
Conservation
* * 4 * 7 * * * * * *
HG102
POPTR_0008s19310
Populus_trichocarpa MVVELQSSSGTEQGKGRRNQTYANVEFVGINQNLK - - - - PTFSDL
Camellia_oleifera MEFVGIGYKTHRNHTPAFPDL
Cercis_gigantea MEFVGIVQNI
Carya_cathayensis MEFVGIVQNM
Ulmus_pumila MEFVGIAQNI
Taxillus_chinensis MEFVGIA
Vitis_vinifera MEFVGIG
Conservation
8 * * * * * 6

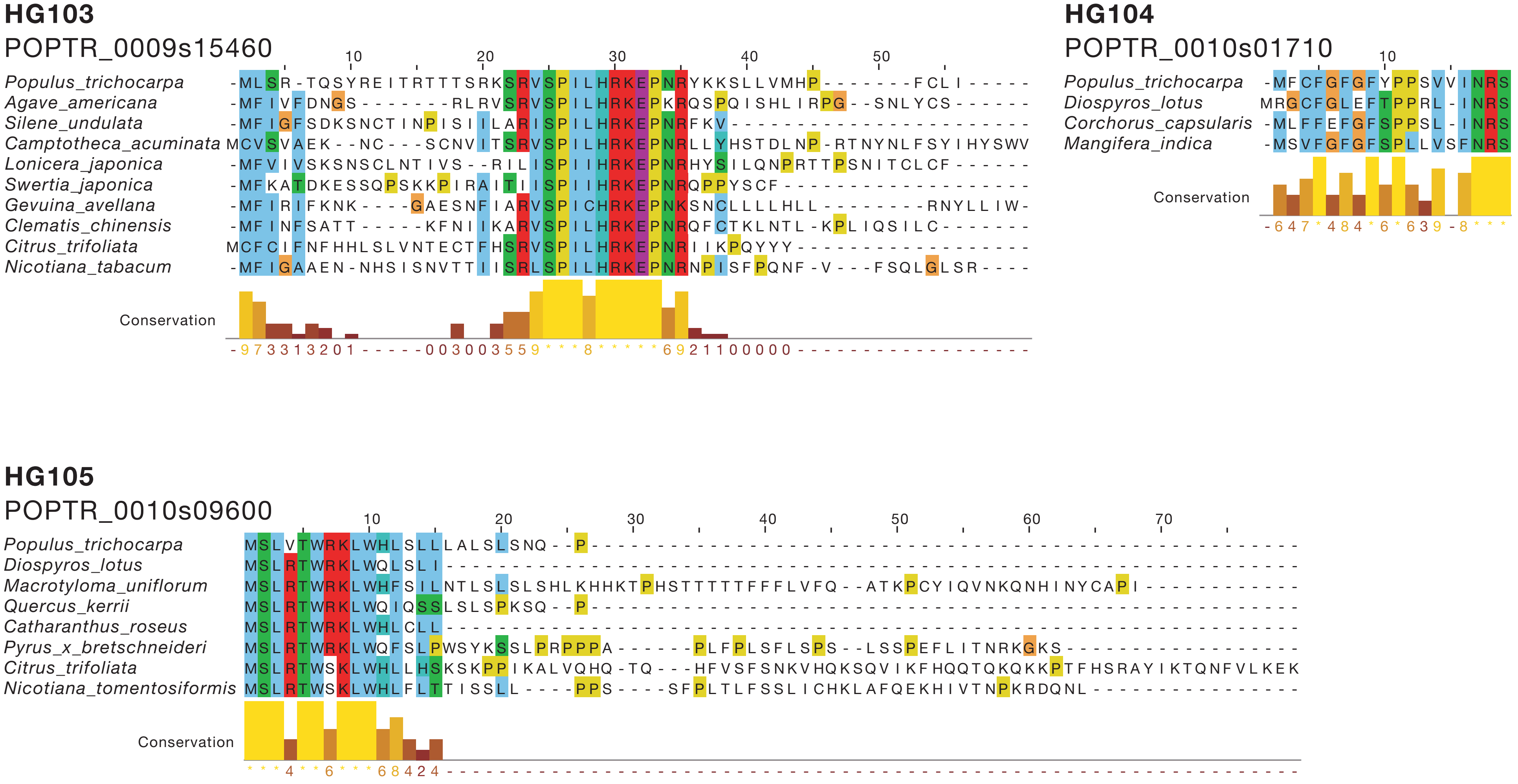
HG103
POPTR_0009s15460
Populus_trichocarpa
Agave_americana
Silene_undulata
Camptotheca_acuminata
Lonicera_japonica
Swertia_japonica
Gevuina_avellana
Clematis_chinensis
Citrus_trifoliata
Nicotiana_tabacum
Conservation
HG104
POPTR_0010s01710
Populus_trichocarpa
Diospyros_lotus
Corchorus_capsularis
Mangifera_indica
Conservation
HG105
POPTR_0010s09600
Populus_trichocarpa
Diospyros_lotus
Macrotyloma_uniflorum
Quercus_kerrii
Catharanthus_roseus
Pyrus_x_bretschneideri
Citrus_trifoliata
Nicotiana_tomentosiformis
Conservation

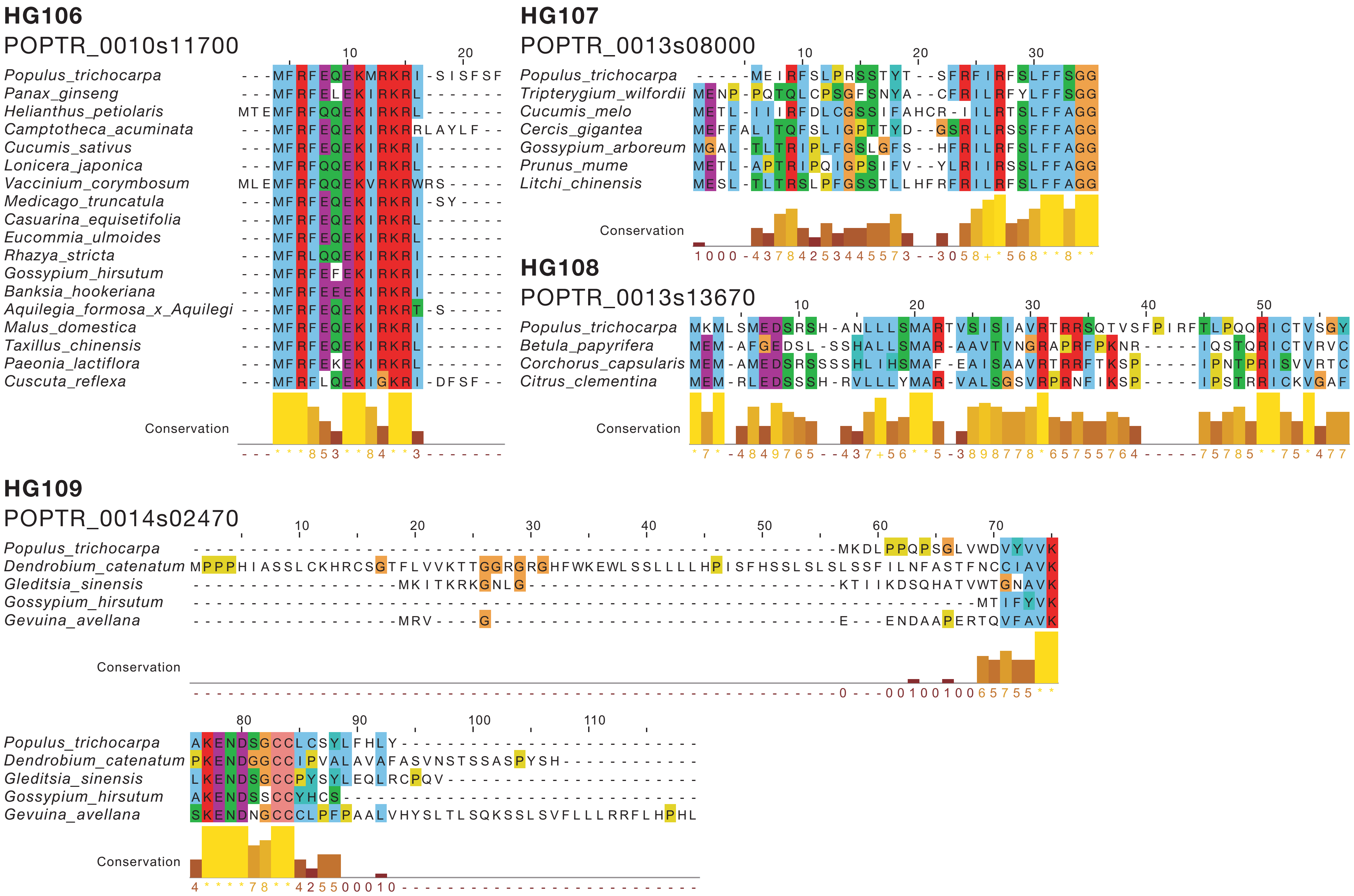
HG106
POPTR_0010s11700
Populus_trichocarpa
Panax_ginseng
Helianthus_petiolaris
Camptotheca_acuminata
Cucumis_sativus
Lonicera_japonica
Vaccinium_corymbosum
Medicago_truncatula
Casuarina_equisetifolia
Eucommia_ulmoides
Rhazya_stricta
Gossypium_hirsutum
Banksia_hookeriana
Aquilegia_formosa_x_Aquilegi
Malus_domestica
Taxillus_chinensis
Paeonia_lactiflora
Cuscuta_reflexa
Conservation
HG107
POPTR_0013s08000
Populus_trichocarpa
Tripterygium_wilfordii
Cucumis_melo
Cercis_gigantea
Gossypium_arboreum
Prunus_mume
Litchi_chinensis
Conservation
HG108
POPTR_0013s13670
Populus_trichocarpa
Betula_papyrifera
Corchorus_capsularis
Citrus_clementina
Conservation
HG109
POPTR_0014s02470
Populus_trichocarpa
Dendrobium_catenatum
Gleditsia_sinensis
Gossypium_hirsutum
Gevuina_avellana
Conservation
Populus_trichocarpa
Dendrobium_catenatum
Gleditsia_sinensis
Gossypium_hirsutum
Gevuina_avellana
Conservation

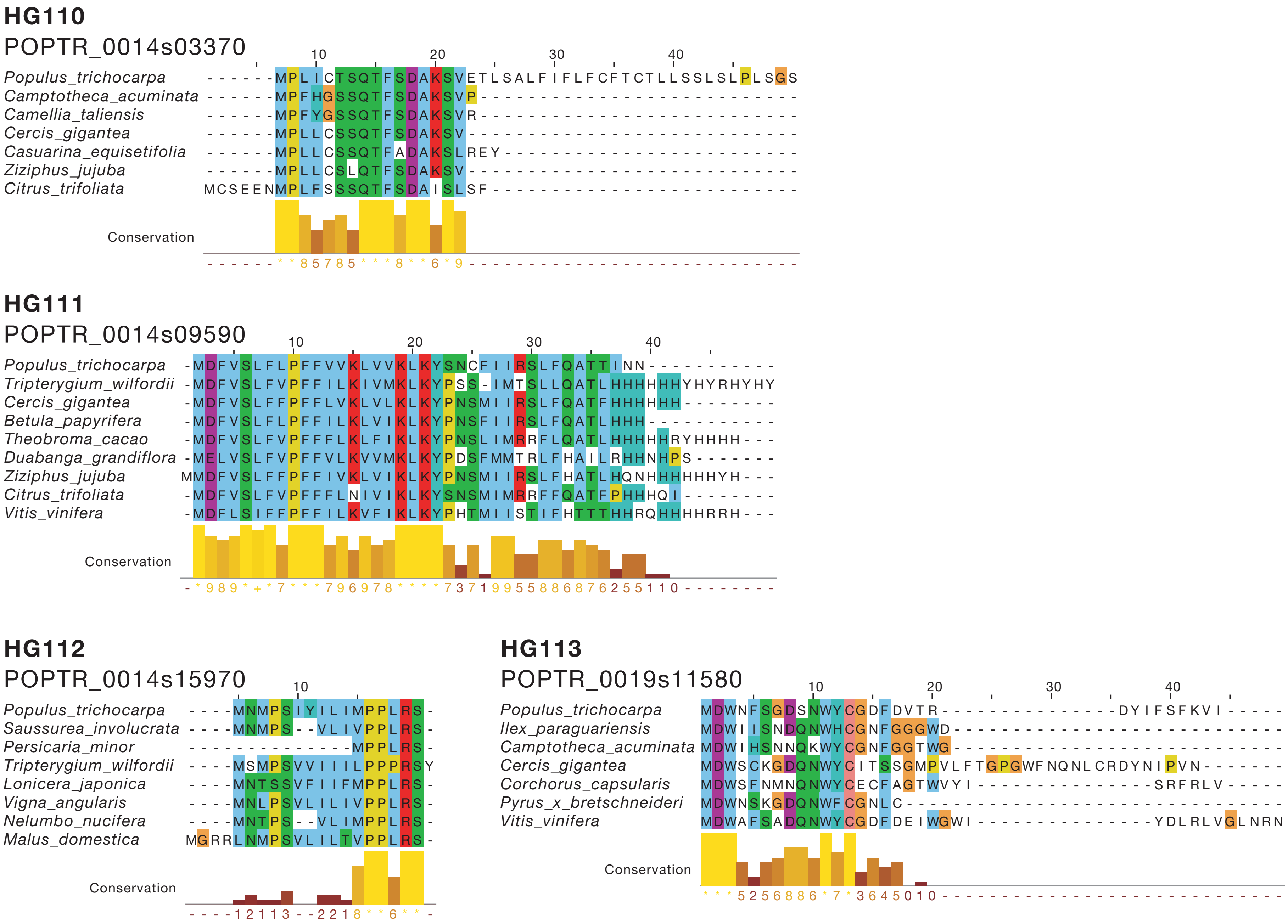
HG110
POPTR_0014s03370
Populus_trichocarpa
Camptotheca_acuminata
Camellia_taliensis
Cercis_gigantea
Casuarina_equisetifolia
Ziziphus_jujuba
Citrus_trifoliata
Conservation
HG111
POPTR_0014s09590
Populus_trichocarpa
Tripterygium_wilfordii
Cercis_gigantea
Betula_papyrifera
Theobroma_cacao
Duabanga_grandiflora
Ziziphus_jujuba
Citrus_trifoliata
Vitis_vinifera
Conservation
HG112
POPTR_0014s15970
Populus_trichocarpa
Saussurea_involucrata
Persicaria_minor
Tripterygium_wilfordii
Lonicera_japonica
Vigna_angularis
Nelumbo_nucifera
Malus_domestica
Conservation
HG113
POPTR_0019s11580
Populus_trichocarpa
Ilex_paraguariensis
Camptotheca_acuminata
Cercis_gigantea
Corchorus_capsularis
Pyrus_x_bretschneideri
Vitis_vinifera
Conservation

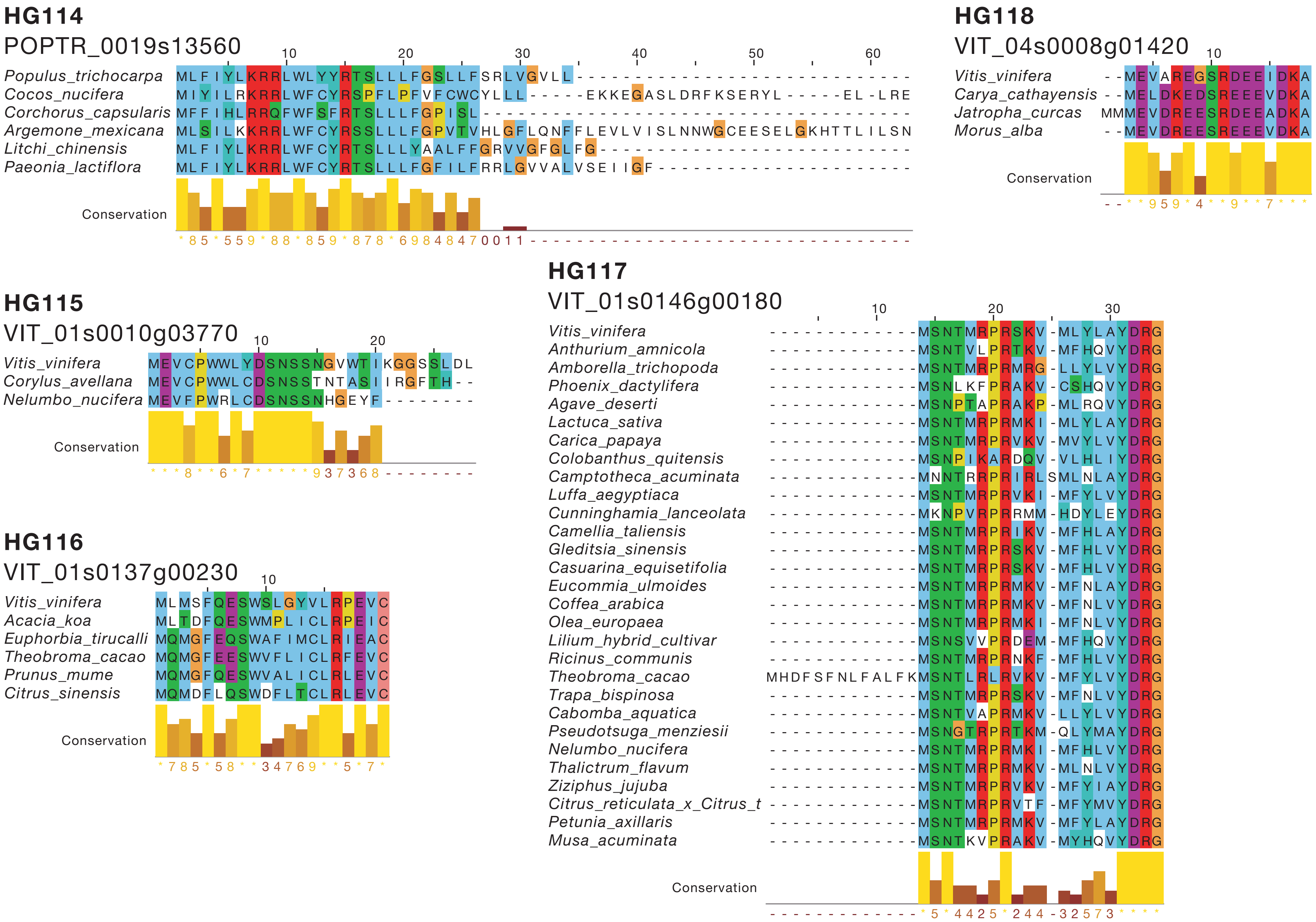
HG114
POPTR_0019s13560
Populus_trichocarpa
Cocos_nucifera
Corchorus_capsularis
Argemone_mexicana
Litchi_chinensis
Paeonia_lactiflora
Conservation
HG118
VIT_04s0008g01420
Vitis_vinifera
Carya_cathayensis
Jatropha_curcas
Morus_alba
Conservation
HG115
VIT_01s0010g03770
Vitis_vinifera
Corylus_avellana
Nelumbo_nucifera
Conservation
HG117
VIT_01s0146g00180
Vitis_vinifera
Anthurium_amnicola
Amborella_trichopoda
Phoenix_dactylifera
Agave_deserti
Lactuca_sativa
Carica_papaya
Colobanthus_quitensis
Camptotheca_acuminata
Luffa_aegyptiaca
Cunninghamia_lanceolata
Camellia_taliensis
Gleditsia_sinensis
Casuarina_equisetifolia
Eucommia_ulmoides
Coffea_arabica
Olea_europaea
Lilium_hybrid_cultivar
Ricinus_communis
Theobroma_cacao
Trapa_bispinosa
Cabomba_aquatica
Pseudotsuga_menziesii
Nelumbo_nucifera
Thalictrum_flavum
Ziziphus_jujuba
Citrus_reticulata_x_Citrus_t
Petunia_axillaris
Musa_acuminata
Conservation
HG116
VIT_01s0137g00230
Vitis_vinifera
Acacia_koa
Euphorbia_tirucalli
Theobroma_cacao
Prunus_mume
Citrus_sinensis
Conservation

**HG119**

VIT_04s0008g04480

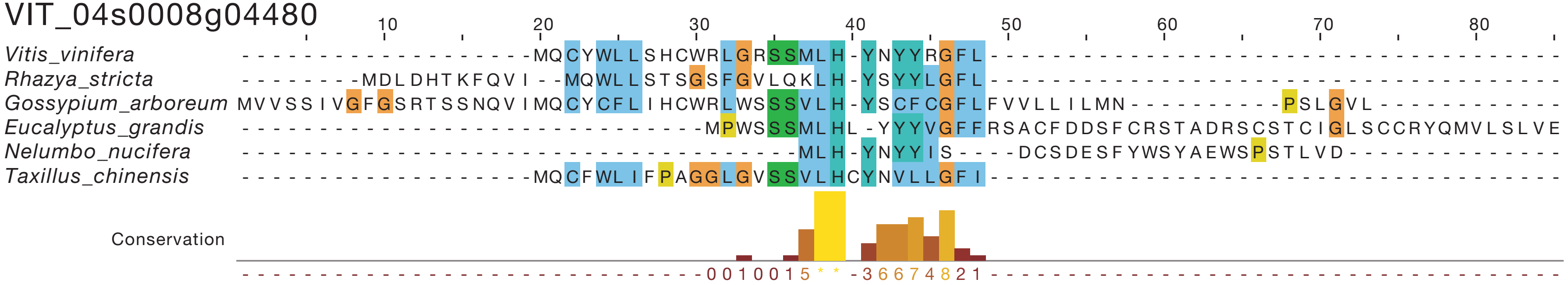


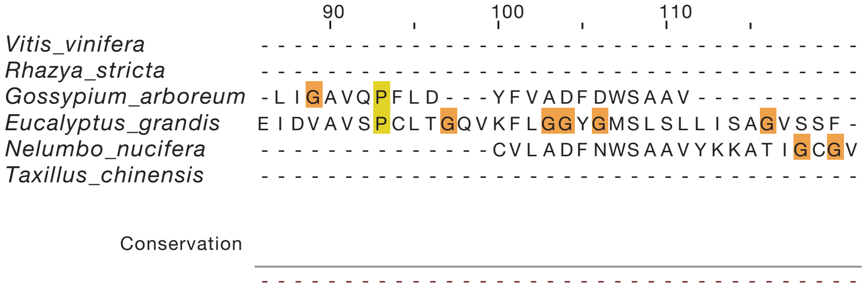


**HG120**

VIT_05s0020g00460

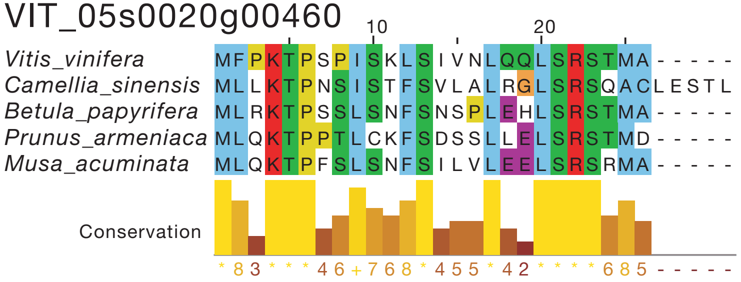

## HG121

VIT_05s0029g00470

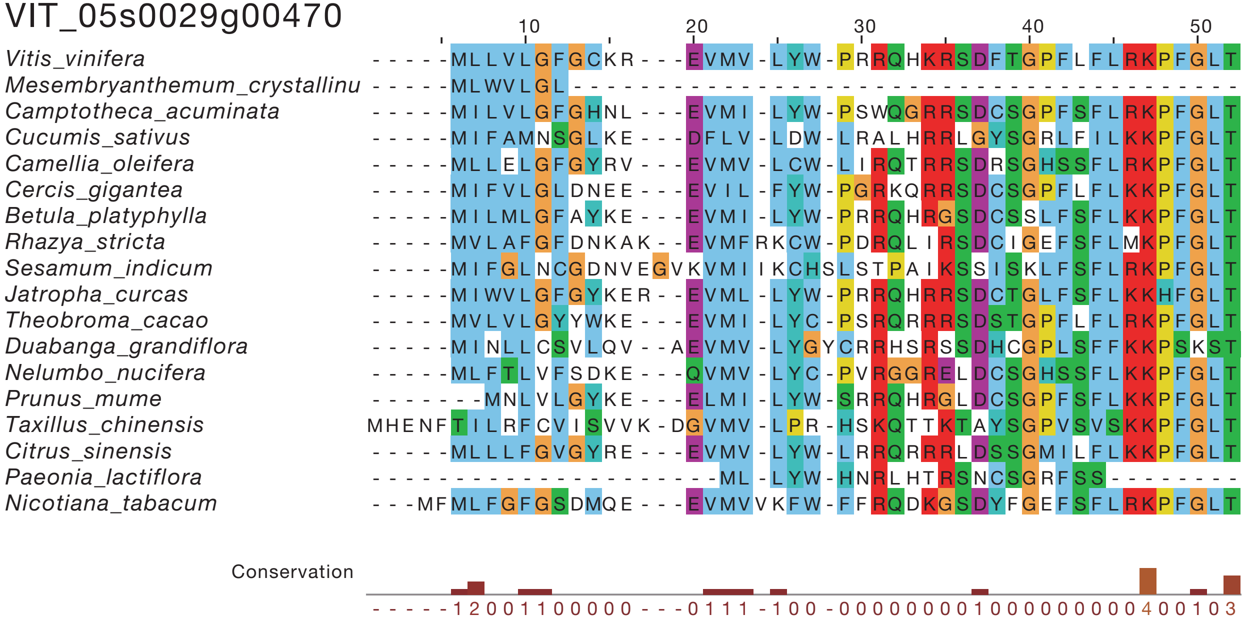


## HG122

VIT_06s0004g02980

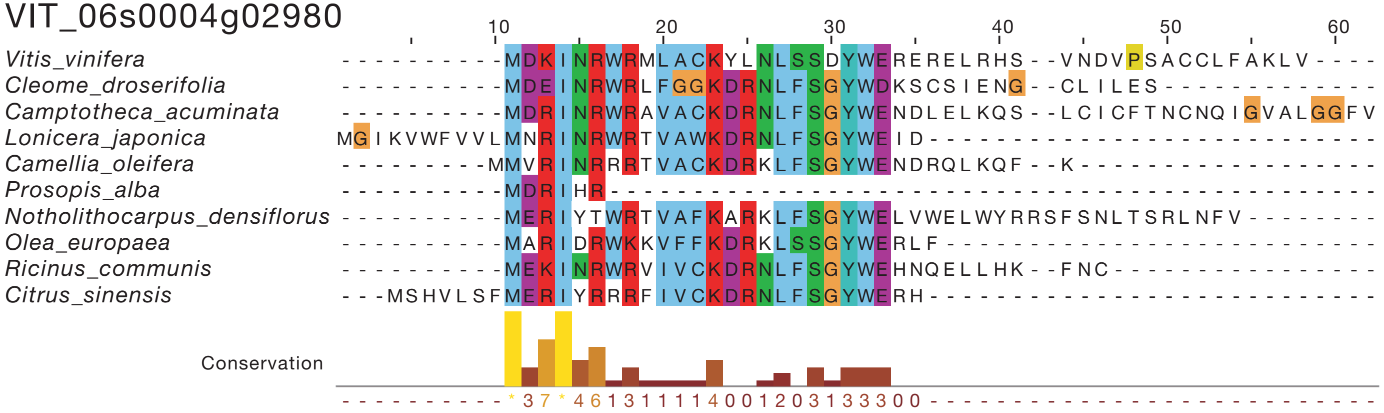

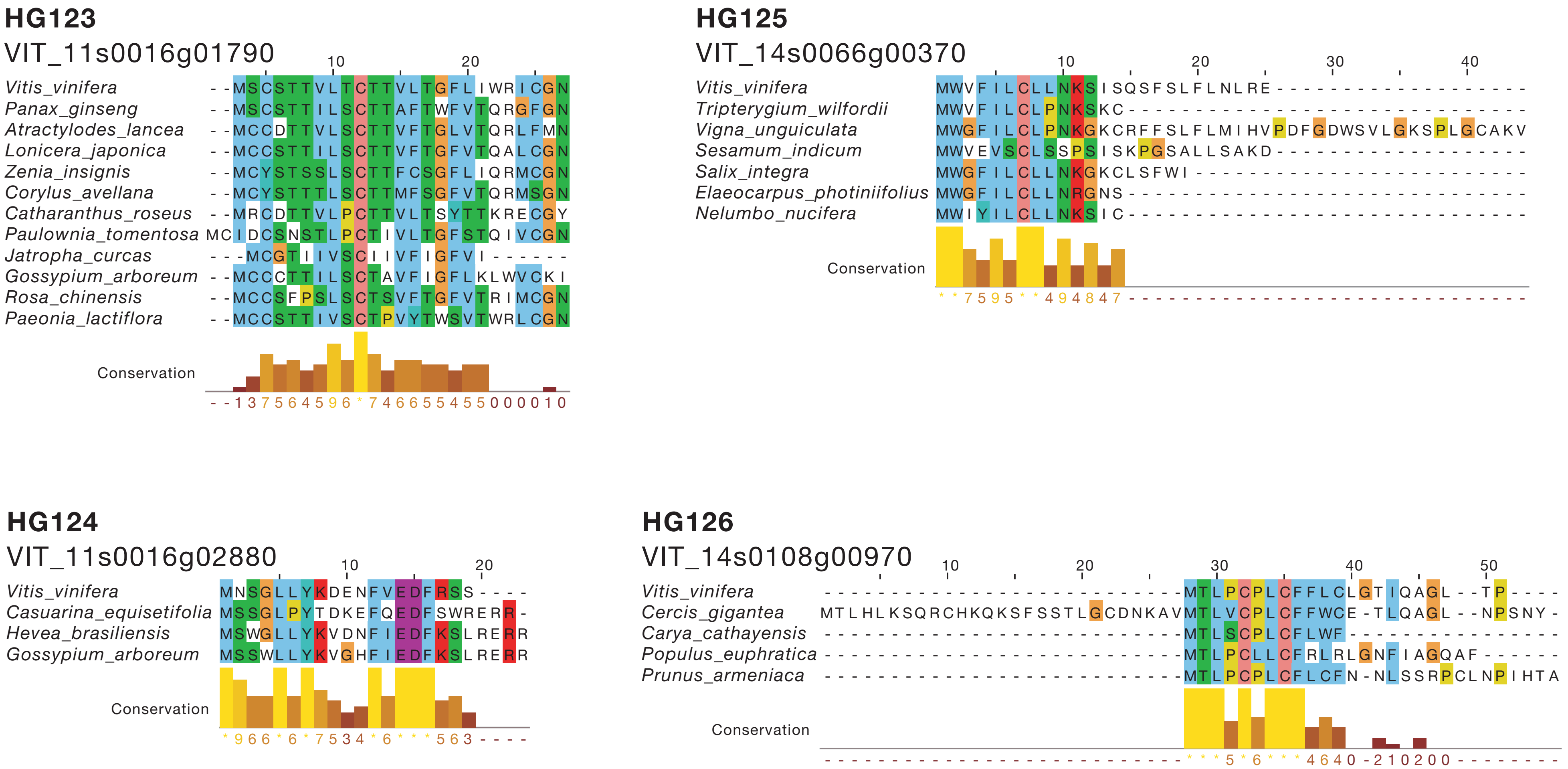
HG123
VIT_11s0016g01790
Vitis_vinifera
Panax_ginseng
Atractylodes_lancea
Lonicera_japonica
Zenia_insignis
Corylus_avellana
Catharanthus_roseus
Paulownia_tomentosa
Jatropha_curcas
Gossypium_arboreum
Rosa_chinensis
Paeonia_lactiflora
Conservation
HG125
VIT_14s0066g00370
Vitis_vinifera
Tripterygium_wilfordii
Vigna_unguiculata
Sesamum_indicum
Salix_integra
Elaeocarpus_photiniifolius
Nelumbo_nucifera
Conservation
HG124
VIT_11s0016g02880
Vitis_vinifera
Casuarina_equisetifolia
Hevea_brasiliensis
Gossypium_arboreum
Conservation
HG126
VIT_14s0108g00970
Vitis_vinifera
Cercis_gigantea
Carya_cathayensis
Populus_euphratica
Prunus_armeniaca
Conservation

**HG127.1**

VIT_18s0001g08100

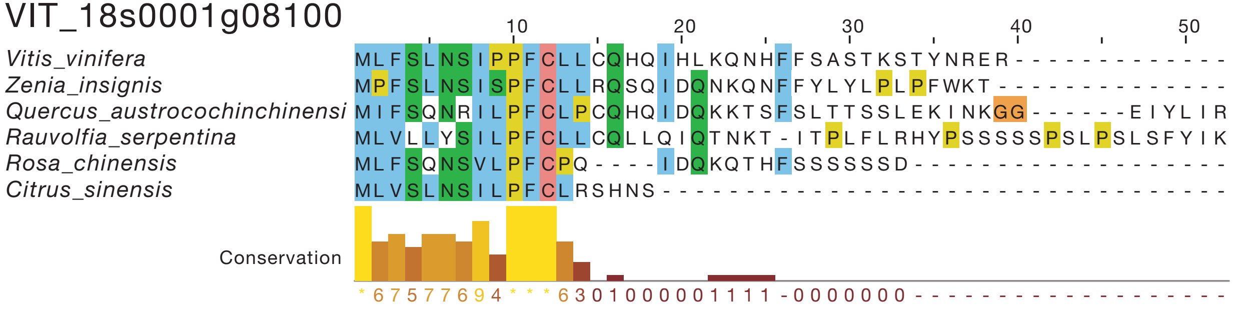


**HG127.2**

VIT_04s0008g04480

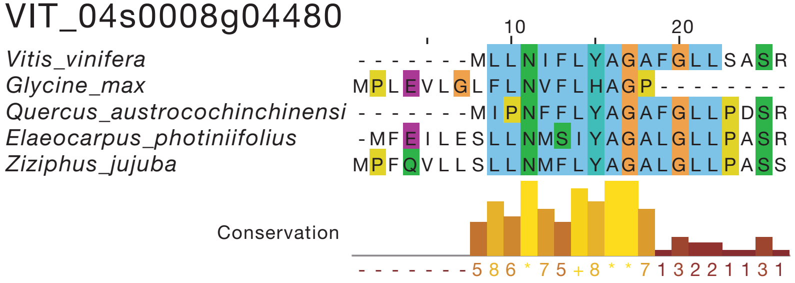


**HG128**

VIT_18s0001g10680

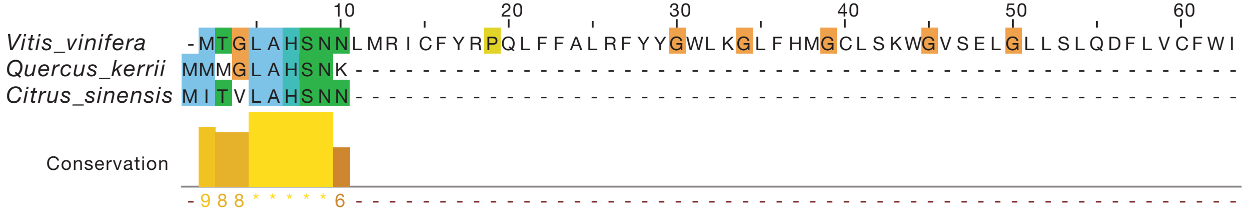

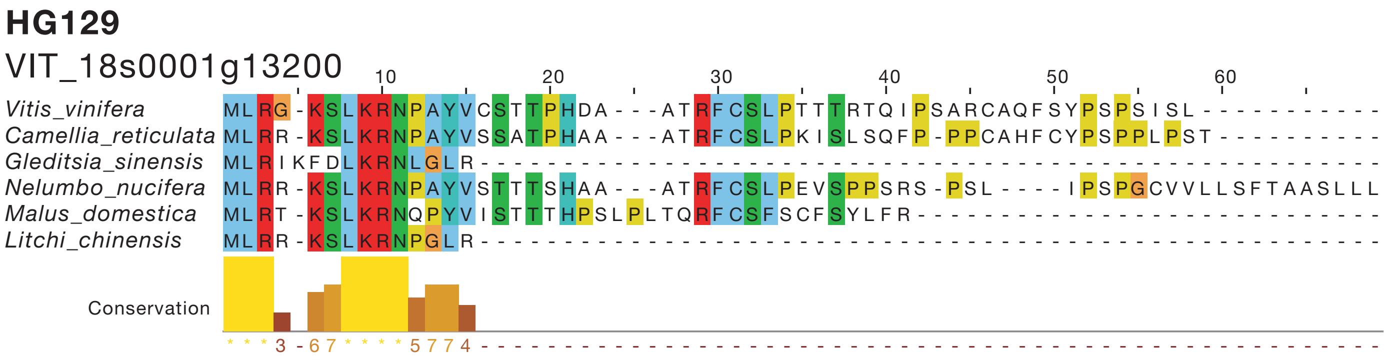


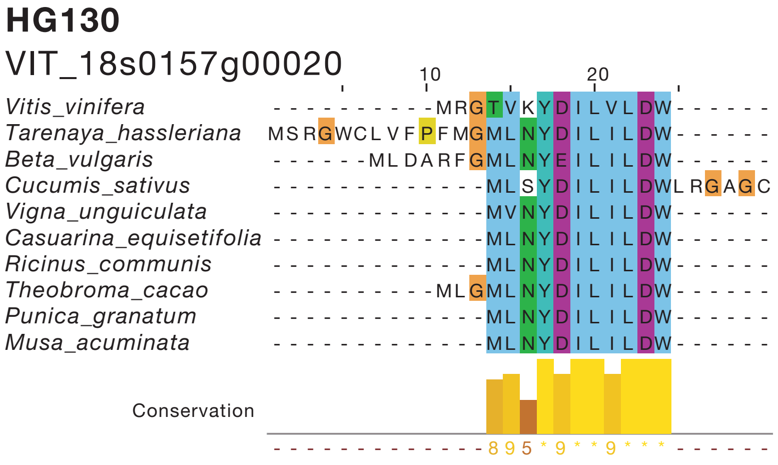


**Supplementary Figure S1.** Alignments of the newly identified CPuORF sequences. The amino acid sequences of the novel CPuORFs and their homologous putative uORFs were aligned using ClustalO ver. 1.2.2 and displayed using Jalview ver. 2.10.2. For each alignment, a putative uORF sequence was selected from each order in which uORF-tBLASTn and mORF-tBLASTn hits were found. In the alignments of the LOC_Os11g16280 (HG075), POPTR_0009s15460 (HG103), and POPTR_0019s11580 (HG113) CPuORFs, some putative uORF sequences were manually removed. After manual removal, $K_a/K_s$ analysis was conducted using the putative uORF sequences shown in the alignments.

### Supplementary Figure S2

## A HG46

1 AGAAGCCAAAAAAAAAAGAAAAGATACAGGACAGTAGAACGTTGGAGAAACCTCCCTGCC

61 TCCTTCCTTAATCTTTCCTTCATAGGTGAACAGAACAGAACAGGGTTGCCGATACTTTTA

121 TGCGGTTTTGCTAAATTCACTAGCCTGTTCTCCTGTGACCCAAGAAAGAATCATTTACTG

181 GTAGGTTTTCTCCAACAATTAATAAACAGAAGTGTAAAAACTGTGTTAATGTTGATTGAT

241 CTCAATTGGGTTTTGTTTATTTGGTCATTGGGTTAGCATTATAAAAGGGTTATGAGAATT

301 CTGTACTGTTTGATTGAAAAGTTTTGATTTTGAGACAAGGGTTTGGATTTTTTGGTCTGG

361 GATTGATTGGATTGTTGTTCTTGCTGGTTTGGCTGAGGTTATTGGATCTGGTTTTTTATT

421 TTGGATTGGTAGTGTTTGATAGAAGTGGTTATCTTGGAATGATTGTTGGTGGGTTTGTGA

481 TAGATCCATGGGAGAATTGTCGTGATTGAAGGGATTTGATTACTTGGTCTGGTGTGACTA

541 GTTTCTTTGTGTTTGTTG**ATGAATATTGTTATTTTTGAAGAGGAGGATCGGCTTGTGAAC**

**M N I V I F E E E D R L V N**

**M N I V I F E E E D R L V N**

601 **TTTCCCAATTCGCTTCTGGGTGTTCTCAGGCCCTTGGATTTACTTGTTGAACGAAAGGTC**

**F P N S L L G V L R P L D L L V E R K V**

**F P I R F W V F S G P W I Y L L N E R S**

G

661 **TTTTCTTTTTGTATCCCTAGAGATAGATGGGATAAGGAGTTTTTCAAGCTTCAGAGTGGA**

**F S F C I P R D R W D K E F F K L Q S G**

**F L F V S L E I D G I R S F S S F R V G**

721 **TAA**TTACTATTATCACCATCATCCATCTAATTTTCAAAGGTTGTGATTGTATAATTTTGG

*****

*****

781 TTTGGAAATTGGAATTTTGGGAAATGAG

**B** HG55

```
  1 CTCTTTCTATGTACTATACCTCTCACCTTACCATTTCCACATCTCTTCTACATCATCATC

 61 GTCATCATTACACCACCTTTTCGCTCACTTGACATCACTTTCCGGCAACAAGTCCGATCT

121 CTTTTCCTTCTCTTTAGACAGGATTATTAAATTACTTCCTGGCAAGTGAAGTCTTTTTTT

181 GCTTACTTTTCTTTCTATGATTGATTCAATTGAGGAAATTCTAGAGCTGGAAGTCGGCTA

241 AGGGTTTTTGGTTTTTTGTTAAGTGGGTTTTGGAGGAATAGGTAGTTTGATTAAAGGGTA

301 ATCTGGTATGTGGGTTGCGGTTAACTGGAATCATGGGTGTGTGGGTGTGGTTTTTTTGTA
           M  W  V  A  V  N  W  N  H  G  C  V  G  V  V  F  L  Y
           M  W  V  A  V  N  W  N  H  G  C  V  G   W  F  F  C  T
                        C
361 CTTTTGGGGAAGAAGCGACGCTGATTAGGTGTGGTTTTCATTTAAATCTTTCGTCTTTTT
     F  W  G  R  S  D  A  D  *
      F  G  E  E  A  T A  D  *

421 TGGTTTAGTTTATCTGGGGTTTTGCTGTTTTTCTTTTTAGTTCTCAGTGTTGGTTCAGTT

481 TGTGAAATTTTGGATCTTTTGGGTACTCGTTATGGA
```

**C** HG57

```
  1 AACAGATACAACACAATCTCCATACTGAAAACATTGTTTTTCAGATGAAATACTACCGGG

 61 TTTTAGACAGAATATCGCGAAAGTTCTGAGATGCATCTAATGTTTGGAGTTCCTGTTGTT
                                  M  H  L  M  F  G  V  P  V  V
                                  M  H  L  M  F  G  V  P   L  F
                                                       C
121 CAAGGAGATCATAAAGTTGAAGAGAGAAAGTGGGAGAAAGAGAGATAGATATTAGCTCAG
    Q  G  D  H  K  V  E  E  R  K  W  E  K  E  R  *
     K  E  I  I  K  L  K  R  E  S  G  R  K  SR  *

181 GCCAAGAAAACTTCAGATTTAGTATTAATTCTCTTTAGTGCGTAAAGGGTTTTTTCTTTT

241 TTCTATTTTTTTCTATTTTTTGCATTCAGACTCTTTTATTGCCCATAAAGCTGAAAGCAT

301 TGAAGGATTTTTAAGAGTGATTTAAGGGAAACGAATGGC
```

**D** HG65

```
  1 GACAACCCTCTCCAAACTCCAAAACAACAAGTCTCTCAAGAGTTTCCAAATTAGGGTTTG

 61 TTCTTATTTTAATGGAAACCTCCTCCGTTAACACATTAAGAGGTTCGTTACTCCCGTTGCA
                M  E  T  S  S  V  N  T  L  R  V  R  Y  S  R  C  N
                M  E  T  S  S  V  N  T  L  R   F  V  T  P  V  A
                                               T
121 ATTGCTTTAAACGTTGTTGTCACTGTTTCTGTTGTTTCTACTCTTACCCCTGAAAACTTA
      C  F  K  R  C  C  H  C  F  C  C  F  Y  S  Y  P  *
    I  A  L  N  V  V  V  T  V  S  V  V  S  T  S Y  P  *

181 TAACCAAAAATCCCTGAAAAAAAAGCCCCAATTTTTATTTTTATTAGAGATGGA
```

**E** HG66

```
  1 AGCCCAGAATGTCCATCTCCTATAATTCGCTTCTCAATTACCCCTTGATTTCCTTTTACT

 61 CACGGTCTTACATTACAGGTTATGATAGTAAACAGCTACCGGAAACCTGAGAGCCAACGC
                         M  I  V  N  S  Y  R  K  P  E  S  Q  R
                         M  I  V  N  S  Y  R  K  P  E  S  Q  R
                                                         C
121 TCCTCTCCCTGCTCTCGGTTTCCCTCGCCCGCGTTTCCTTCGCTACCGCCGTTTGAGGTAA
    S  S  P  A  L  G  F  P  R  P  R  F  L  R  Y  R  R  L  R  *
    S  S   L  L  S  V  S  L  A  R  V  S  F  A  T  A  VL  R  *

181 CCGGTGAAATAGAGTAACTGGAGGGAAAGAGATTTTTTATTGTATTTTTTTATTTTTTCG

241 TGTAACACCGTTACTGTCGCTTACAATTATGCC
```

**F** HG80

```
  1 GAGATTATAATCAAGGTGGTCAATTGATCATATTTTGTACTTGCTAATAGTATAATTAAC

 61 TTGTTGTGTTGGTTCTGAATGATGGTGTAGTAGGTATTTCTTCCTCTGAAATGCATTTGT

121 GAGTTGCAGCAGTTCACAAAAACAATTTTTTGTTGACAATGTTTCTGCAAGATGGGCATA
                                          M  F  L  Q  D  G  H  I
                                          M  F  L  Q  D  G  H  I
           C
181 TTGGGTTTTCGGATACAAGCGATCGCAATCCTTCCTGGAAGTTCCTCTTTTTGTTTTTCA
      G  F  S  D  T  S  D  R  N  P  S  W  K  F  L  F  L  F  F  I
      G  F F  G  Y  K  R  S  Q  S  F  L  E  V  P  L  F  V  F  H

241 TCTTTACAGTTCTTTTGTGTTTCGGATCGTGATAAAAGTTTGCAAATTCAGATTTACTAG
      F  T  V  L  L  C  F  G  S  *
     L  Y  S  S  F  V  F  R   S  *

301 GAGTTGTTGGATACAAATTTTCAATGCA
```

**G** HG81

```
  1 AAATGTCCTCTTCATTGATTGAGACCCCATTAAAAAAGAAAAGAGAAAAGAGAAAAACAA

 61 AGAAAGAGGTAAAAGCAGAAGCATCAGCAGCAAGCAACACAACCACGTACTATAATATCC

121 ATCGTTACTGGGAAAAATAAACTCTCCCCATAAATCTTCCGTTTCTTCTCCTCTGTTAAC

181 ATGTTGTGAATATAGGTAGCATACAAGTGCTTGAAGCAGTGTGGTGTGGTAGCTATGAAA

241 ATGTTGCGCACTCGGTTTTTAGTCCATGGTTAATGCTATTTGGGGTGTGAAAAGATCTGG

301 GTTTTCTCAGAGAACTCAAAATTTAGTGACATTCTACTAAGATCTAGCTTAAATGAGCCG

361 CACAAGTGAGTGCTTTTGTCGGCTGGTTAAAGCTTATTTGAAAAGCGGATGAGGCTGAAA
                                                    M   R   L   K
                                                    M   R   L   K
                                                  T
421 TCGATCGTGTTTTGTAGCAAAACCGGTTTCGGTTGAGGATTTAGAAACTGGGTCCGTTTC
    S   I   V   F   C   S   K   T   G   F   G   *
     R   S   C   F   V   A   K   P   V   S  G   *

481 TTTGCTTTTCTTGGCTTCCAGATGAACTTGTCTTTGATGGTTTAATAGTTTAGTCATTCT

541 TTTTTGGGTAAGAGGGAAGGTAAAAAGGTGAAGAACAGGAAGAAGAAGGCATGATAGTGT

601 CATTGTGTGAAGGGAAGAAATTAGTGAAGATTGTTTGGGTTGTGGGTCTAAACTTGTTGC

661 GATTGTATTATTAAATCTCTCCCCCTCTCTCAGTCTTAATGAG
```

**H** HG87

```
  1 AAGGAAAGGGTGCTGAGTATATCAATCAAGAATTTCTGAGACCACAAAGAAGCTAGGTAG

 61 CTTGAGTAAGCTTGATCTTCTCAAGTTCTTGTTATCGTCATTAATTTCCCTGTAAGAGAT

121 AGTTTAATATAGACTGGTATTTAGGGAGGAGACTATACTAATAAGATAGATAGTGACTCG

181 AATAGACATGCTCACTTCTCATCACCTTCCTACCCATTTCCTAGCAACATTTCACTCAAT
           M  L  T  S  H  H  L  P  T  H  F  L  A  T  F  H  S  I
           M  L  T  S  H  H  L  P  T  H  F  L  A  T  F  H  S  I
                                C
241 TATATCACCGATTTCACTTGAGCAATATTTCACTGTTCGCCTCACAAGGATCAACCTCAT
     I  S  P  I  S  L  E  Q  Y  F  T  V  R  L  T  R  I  N  L  I
     I  S  P  I  S  L  E  H I  F  H  C  S  P  H  K  D  Q  P  H

301 CCTCTTGTTATTCTTCATATTGATTTCTCTTCTTGCTCCAAGGTCCAAGGGCTAGCCAGT
     L  L  L  F  F  I  L  I  S  L  L  A  P  R  S  K  G  *
    P  L  V  I  L  H  I  D  F  S  S  C  S  K  V  Q   G  *

361 TAGAATTTGTTTGCTTTGTTCCTGTCTGAAAATAGAGAAAATTTGCTTCCTCTCACTCTC

421 TTGGTTGTTATTTAGTTGCTAATCTTGTAGTGTTTTAAGAGATGGG
```

**I** HG88

```
  1 TGATTAAACAATTTCGAAGACTTTCCTTTCTCTTCTCTGGTGCTATAAGCTATTTGAGAC

 61 TGGGCTTGATTAATGGCAATGCATATATATTGCTGTCCTTTTGGTTCGCATAGGACTGGA
                 M  A  M  H  I  Y  C  C  P  F  G  S  H  R  T  G
                 M  A  M  H   Y  I  A  V  L  L  V  R  I  G  L  E
                             C
121 ATTTGGAATTGTTGCAGAGAGAGAAACTTGCTTTTGGATGCTTACTTGTAAACTAGAGCA
    I  W  N  C  C  R  E  R  N  L  L  L  D  A  Y  L  *
     F  G  I  V  A  E  R  DN  L  L  L  D  A  Y  L  *

181 GCCCAAGTTGGTTTTTTTTTTTTAATTTTATTTGGATGCTCAATTTTTAATACAAGTTCG

241 GTGATTGATAAGTGGTTCTTCGAGTTAGTTTCAGTGATTTCTGTTATTAGATTTGGGTTG

301 GTGGGTTAGCTGCTACATTCTTTGCATCATGCC
```

**J** HG103

```
  1 ACTGGTATCTCTCTCTCCCTTTTCTAATTATGCTTTCAAGAACACAGTCATATAGAGAAA
                                 M  L  S  R  T  Q  S  Y  R  E  I
                                 M  L  S  R  T  Q  S  Y  R  E  I

 61 TAACAAGAACAACAACATCAAGAAAATCCAGGGTTTCTCCGATCCTCCATAGAAAAGAAC
      T  R  T  T  T  S  R  K  S  R  V  S  P  I  L  H  R  K  E  P
      T  R  T  T  T   Q  E  N  P  G  F  L  R  S  S  I  E  K  N
             C
             ↓
121 CTAACAGACATAAAAAATCTCTTCTTGTTATGCACCCCTTTTGCCTTATTTAGTGGGCCT
      N  R  H  K  K  S  L  L  V  M  H  P  F  C  L  I  *
    L  T  D   HK  K  S  L  L  V  M  H  P  F  C  L  I  *

181 ATACTTTCTCTTACACAAATTCCTAACTTGGGTTTTTCTTGTTTTGTCTAATTCAGTGGT

241 TCTTGTTTGTTTTTGTTGTTGTCGTTGTTGTTAAATTGTTGTTGAAGTTTGTTTTAGGGT

301 TTTAAGATGGG
```

**K** HG107

```
  1 GAGTTTCCGCGTCAAAAATCAGTGAGCCAGCAGAACACACCACTCGTCCTTCAATTTCAT

 61 TCTCTGCCGCCGATGCCGCTGCTGCTCGTTTGTTTTCGTTTTTAAAAAAACCAAACAAAA

121 ACCAATCCATCCATCTATCTGCCTTTATATCATATTAGATAAGAAAACAAGGGAATTTCG

                                                            A
181 AAAACAGATCGGTGAAGGGAAGGAAAAAAAACCAAACCCTAAAAGGGAAGAAAATGAAAA

241 GTCAGCAATTCTGAATGGAAATCCGATTCTCTTTACCCAGATCATCCACTTACACCTCTT
                  M  E  I  R  F  S  L  P  R  S  S  T  Y  T  S  F
                  M  E  I  R  F  S  L  P  R  S  S  T  Y  T   L
                                                C
301 TCCGGTTCATCCGCTTCTCTCTCTTCTTCTCTGGAGGCTGATTGATTTTCTTTTTCTTTT
      R  F  I  R  F  S  L  F  F  S  G  G  *
     S  G  S  S  A  S  L  S  S  S  L  E  A  *

361 TTGTTTCCTGTAAAATAAAAACTGCATTTACATATATATATATATATATAGAGAGAGAGA

421 GAGAGAGCGCGCGCGCTCGTGGATGGA
```

Supplementary Figure S2. 5´-UTR nucleotide and deduced amino acid sequences of the poplar CPuORFs analyzed in the transient expression study. (A-J) The 5´-UTR nucleotide sequences are based on the sequences of the *Populus nigra* full-length cDNA clones, pds25559 (A), pds10965 (B), pds14390 (C), pds12940 (D), pds13862 (E), pds15817 (F), pds28294 (G), pds26157 (H), pds14623 (I) and pds23234 (J) (GenBank accession no. DB890234, DB875826, DB879210, DB877779, DB878687, DB880616, DB892931, DB890822, DB879437 and DB887941, respectively). We determined the pds25559 and pds28294 5´-UTR nucleotide sequences by sequencing because sequence information of these regions was not available in public databases. Arrows indicate the positions of primers used for the 5´-UTR cloning. (K) The 5´-UTR nucleotide sequence of the *Poplus trichocarpa* POPTR_0013s08000 gene is based on NCBI Refseq XM_002319213.3. The nucleotide 'A' shown in a black background indicates the nucleotide change from T to A for the removal of the start codon of the uORF located immediately upstream of the HG107 CPuORF. In (A)-(K), the CPuORF nucleotide and deduced amino acid sequences are shown in bold. The nucleotide sequences of uORFs other than the CPuORFs are underlined. The initiation codon of the mORF in each sequence is boxed. The nucleotides that were deleted and inserted in the frameshift mutants are shaded, and the deduced amino sequences of the frameshift mutant CPuORFs are indicated.

Supplementary Table S1. CPuORFs extracted from *A. thaliana* , *O. sativa* , *S. lycopersicum* , *P. trichocarpa* and *V. vinifera* .

| HG number | Species | Gene ID | Gene symbo | Gene description† | uORF-mORF fusion ratio | $K_a/K_s$ analysis before manual validation | | | $K_a/K_s$ analysis after manual validation | |
|---|---|---|---|---|---|---|---|---|---|---|
| | | | | | | Median pairwise $K_a/K_s$ ratio | $U$-test $p$ value | $q$ value | Median pairwise $K_a/K_s$ ratio | $U$-test $p$ value |
| HG1 | *A. thaliana* | AT1G75390 | BZIP44 | bZIP transcription factor 44 | 0.01 | 0.05 | 0.00E+00 | 0.00E+00 | 0.05 | 0.00E+00 |
| | | AT2G18160 | BZIP2 | bZIP transcription factor 2 | 0.00 | 0.07 | 0.00E+00 | 0.00E+00 | 0.07 | 0.00E+00 |
| | | AT3G62420 | BZIP53 | bZIP transcription factor 53 | 0.00 | 0.03 | 0.00E+00 | 0.00E+00 | 0.03 | 0.00E+00 |
| | | AT4G34590 | BZIP11 | bZIP transcription factor 11 | 0.00 | 0.11 | 0.00E+00 | 0.00E+00 | 0.11 | 0.00E+00 |
| | | AT5G49450 | BZIP1 | Basic leucine zipper 1 | 0.00 | 0.03 | 6.29E-250 | 2.71E-249 | 0.03 | 6.29E-250 |
| | *O. sativa* | LOC_Os02g03960 | | Os02g0132500 protein | 0.00 | 0.05 | 0.00E+00 | 0.00E+00 | 0.05 | 0.00E+00 |
| | | LOC_Os03g19370 | | | 0.29 | 0.04 | 8.52E-269 | 2.13E-268 | 0.04 | 8.52E-269 |
| | | LOC_Os05g03860 | | bZIP transcription factor 53 | 0.00 | 0.04 | 0.00E+00 | 0.00E+00 | 0.04 | 0.00E+00 |
| | | LOC_Os08g26880 | | | 0.02 | 0.06 | 0.00E+00 | 0.00E+00 | 0.06 | 0.00E+00 |
| | | LOC_Os09g13570 | | Os09g0306400 protein | 0.00 | 0.08 | 0.00E+00 | 0.00E+00 | 0.08 | 0.00E+00 |
| | | LOC_Os12g37410 | | | 0.00 | 0.04 | 0.00E+00 | 0.00E+00 | 0.04 | 0.00E+00 |
| | *S. lycopersicum* | Solyc01g079480.2 | | BZIP DNA-binding protein | 0.00 | 0.03 | 0.00E+00 | 0.00E+00 | 0.03 | 0.00E+00 |
| | | Solyc01g100460.2 | | Anaerobic basic leucine zipper protein | 0.00 | 0.04 | 2.27E-196 | 8.04E-196 | 0.04 | 2.27E-196 |
| | | Solyc01g109880.2 | | BZIP transcription factor | 0.00 | 0.06 | 0.00E+00 | 0.00E+00 | 0.06 | 0.00E+00 |
| | *P. trichocarpa* | POPTR_0002s19720 | | BZIP transcription factor family protein | 0.00 | 0.03 | 0.00E+00 | 0.00E+00 | 0.03 | 0.00E+00 |
| | | POPTR_0004s16560 | | | 0.00 | 0.10 | 0.00E+00 | 0.00E+00 | 0.10 | 0.00E+00 |
| | | POPTR_0005s25290 | | BZIP transcription factor family protein | 0.01 | 0.05 | 0.00E+00 | 0.00E+00 | 0.05 | 0.00E+00 |
| | | POPTR_0007s13380 | | | 0.01 | 0.06 | 0.00E+00 | 0.00E+00 | 0.06 | 0.00E+00 |
| | | POPTR_0008s10620 | | BZIP family transcription factor family protein | 0.00 | 0.02 | 1.43E-300 | 1.34E-299 | 0.02 | 1.43E-300 |
| | | POPTR_0009s12250 | | BZIP transcription factor family protein | 0.00 | 0.10 | 0.00E+00 | 0.00E+00 | 0.10 | 0.00E+00 |
| | | POPTR_0010s15280 | | BZIP family transcription factor family protein | 0.00 | 0.04 | 7.13E-281 | 6.46E-280 | 0.04 | 7.13E-281 |
| | | POPTR_0014s11590 | | | 0.00 | 0.03 | 0.00E+00 | 0.00E+00 | 0.03 | 0.00E+00 |
| | *V. vinifera* | VIT_03s0038g04450 | | Putative uncharacterized protein | 0.00 | 0.09 | 0.00E+00 | 0.00E+00 | 0.09 | 0.00E+00 |
| | | VIT_04s0023g02430 | | Putative uncharacterized protein | 0.00 | 0.04 | 0.00E+00 | 0.00E+00 | 0.04 | 0.00E+00 |
| | | VIT_05s0077g01140 | | Putative uncharacterized protein | 0.00 | 0.03 | 0.00E+00 | 0.00E+00 | 0.03 | 0.00E+00 |
| | | VIT_07s0005g01450 | | Basic region/leucine zipper motif C22 protein | 0.00 | 0.03 | 0.00E+00 | 0.00E+00 | 0.03 | 0.00E+00 |
| | | VIT_14s0060g01210 | ATBZIP53 | Putative uncharacterized protein | 0.00 | 0.04 | 0.00E+00 | 0.00E+00 | 0.04 | 0.00E+00 |
| HG2 | *A. thaliana* | AT1G06150 | EMB1444 | basic helix-loop-helix (bHLH) DNA-binding superfamily protein | 0.04 | 0.04 | 0.00E+00 | 0.00E+00 | 0.04 | 0.00E+00 |
| | | AT2G27230 | LHW | Transcription factor LHW | 0.03 | 0.04 | 0.00E+00 | 0.00E+00 | 0.04 | 0.00E+00 |
| | | AT2G31280 | CPUORF7 | conserved peptide upstream open reading frame | 0.05 | 0.04 | 0.00E+00 | 0.00E+00 | 0.04 | 0.00E+00 |
| | *O. sativa* | LOC_Os11g06010 | | Os11g0158500 protein | 0.05 | 0.04 | 2.40E-305 | 8.56E-305 | 0.04 | 2.40E-305 |
| | | LOC_Os12g06330 | | Expressed protein | 0.05 | 0.04 | 0.00E+00 | 0.00E+00 | 0.04 | 0.00E+00 |
| | *S. lycopersicum* | Solyc06g074110.2 | | | 0.05 | 0.04 | 0.00E+00 | 0.00E+00 | 0.04 | 0.00E+00 |
| | | Solyc09g011170.2 | | | 0.02 | 0.05 | 0.00E+00 | 0.00E+00 | 0.05 | 0.00E+00 |
| | | Solyc09g066280.2 | | Transcription factor bHLH155 | 0.04 | 0.04 | 0.00E+00 | 0.00E+00 | 0.04 | 0.00E+00 |
| | *P. trichocarpa* | POPTR_0001s22450 | | Transcription factor-related family protein | 0.05 | 0.04 | 0.00E+00 | 0.00E+00 | 0.04 | 0.00E+00 |
| | | POPTR_0005s24390 | | Basic helix-loop-helix family protein | 0.04 | 0.04 | 0.00E+00 | 0.00E+00 | 0.04 | 0.00E+00 |
| | | POPTR_0006s09100 | | | 0.02 | 0.04 | 0.00E+00 | 0.00E+00 | 0.04 | 0.00E+00 |
| | *V. vinifera* | VIT_08s0007g00940 | | Putative uncharacterized protein | 0.01 | 0.05 | 0.00E+00 | 0.00E+00 | 0.05 | 0.00E+00 |
| HG2.2 | *P. trichocarpa* | POPTR_0001s22450 | | Transcription factor-related family protein | 0.00 | 0.07 | 7.81E-21 | 2.22E-20 | 0.07 | 7.81E-21 |
| HG3 | *A. thaliana* | AT3G02470 | SAMDC1 | S-adenosylmethionine decarboxylase proenzyme | 0.22 | 0.13 | 0.00E+00 | 0.00E+00 | 0.13 | 0.00E+00 |
| | | AT3G25570 | SAMDC3 | S-adenosylmethionine decarboxylase proenzyme | 0.24 | 0.12 | 0.00E+00 | 0.00E+00 | 0.12 | 0.00E+00 |
| | | AT5G15950 | SAMDC2 | S-adenosylmethionine decarboxylase proenzyme | 0.26 | 0.12 | 0.00E+00 | 0.00E+00 | 0.12 | 0.00E+00 |
| | *O. sativa* | LOC_Os02g39790 | | S-adenosylmethionine decarboxylase proenzyme | 0.24 | 0.11 | 0.00E+00 | 0.00E+00 | 0.11 | 0.00E+00 |
| | | LOC_Os04g42090 | SAMDC | S-adenosylmethionine decarboxylase proenzyme | 0.24 | 0.12 | 0.00E+00 | 0.00E+00 | 0.12 | 0.00E+00 |
| | | LOC_Os09g25620 | | S-adenosylmethionine decarboxylase proenzyme | 0.27 | 0.13 | 0.00E+00 | 0.00E+00 | 0.13 | 0.00E+00 |
| | *S. lycopersicum* | Solyc01g010050.2 | | S-adenosylmethionine decarboxylase proenzyme | 0.27 | 0.12 | 0.00E+00 | 0.00E+00 | 0.12 | 0.00E+00 |
| | *P. trichocarpa* | POPTR_0004s10660 | | S-adenosylmethionine decarboxylase proenzyme | 0.27 | 0.12 | 0.00E+00 | 0.00E+00 | 0.12 | 0.00E+00 |
| | | POPTR_0010s14390 | | S-adenosylmethionine decarboxylase proenzyme | 0.27 | 0.12 | 0.00E+00 | 0.00E+00 | 0.12 | 0.00E+00 |
| | | POPTR_0017s14280 | | S-adenosylmethionine decarboxylase proenzyme | 0.26 | 0.14 | 0.00E+00 | 0.00E+00 | 0.14 | 0.00E+00 |
| | | POPTR_0018s11040 | | S-adenosylmethionine decarboxylase proenzyme | 0.17 | 0.13 | 0.00E+00 | 0.00E+00 | 0.13 | 0.00E+00 |
| HG4 | *A. thaliana* | AT4G25670 | | | 0.02 | 0.19 | 0.00E+00 | 0.00E+00 | 0.19 | 0.00E+00 |
| | | AT4G25690 | | | 0.02 | 0.19 | 6.05E-292 | 3.39E-291 | 0.19 | 6.05E-292 |
| | | AT5G52550 | | AT5g52550/F6N7_3 | 0.02 | 0.18 | 0.00E+00 | 0.00E+00 | 0.18 | 0.00E+00 |
| | *O. sativa* | LOC_Os02g01360 | | | 0.02 | 0.18 | 2.26E-246 | 4.72E-246 | 0.18 | 2.26E-246 |
| | *P. trichocarpa* | POPTR_0004s07800 | | | 0.05 | 0.18 | 0.00E+00 | 0.00E+00 | 0.18 | 0.00E+00 |
| | | POPTR_0017s01640 | | | 0.05 | 0.19 | 0.00E+00 | 0.00E+00 | 0.19 | 0.00E+00 |
| HG5 | *A. thaliana* | AT5G07840 | PIA1 | PIA1 | 0.04 | 0.06 | 1.61E-276 | 8.74E-276 | 0.06 | 1.61E-276 |
| | | AT5G61230 | PIA2 | PIA2 | 0.05 | 0.06 | 1.26E-259 | 6.03E-259 | 0.06 | 1.26E-259 |
| | *O. sativa* | LOC_Os02g01240 | | Os02g0102600 protein | 0.06 | 0.06 | 6.81E-181 | 1.19E-180 | 0.06 | 6.81E-181 |
| | *P. trichocarpa* | POPTR_0001s01190 | | Ankyrin repeat family protein | 0.04 | 0.06 | 1.39E-216 | 8.34E-216 | 0.06 | 1.39E-216 |
| | | POPTR_0003s10350 | | Ankyrin repeat family protein | 0.04 | 0.06 | 2.91E-265 | 2.17E-264 | 0.06 | 2.91E-265 |
| | *V. vinifera* | VIT_02s0087g00100 | | Putative uncharacterized protein | 0.04 | 0.05 | 0.00E+00 | 0.00E+00 | 0.05 | 0.00E+00 |
| HG6 | *A. thaliana* | AT2G43020 | PAO2 | Probable polyamine oxidase 2 | 0.00 | 0.06 | 7.75E-32 | 1.63E-31 | 0.06 | 7.75E-32 |
| | | AT3G59050 | PAO3 | Polyamine oxidase 3 | 0.01 | 0.12 | 2.70E-06 | 4.44E-06 | 0.12 | 2.70E-06 |
| | *S. lycopersicum* | Solyc02g081390.2 | | | 0.00 | 0.13 | 6.53E-236 | 2.83E-235 | 0.13 | 6.53E-236 |
| | *P. trichocarpa* | POPTR_0004s07430 | | Amine oxidase family protein | 0.00 | 0.17 | 1.57E-121 | 7.00E-121 | 0.17 | 1.57E-121 |
| | *V. vinifera* | VIT_04s0043g00220 | | Putative uncharacterized protein | 0.00 | 0.11 | 6.53E-235 | 5.46E-234 | 0.11 | 6.53E-235 |
| HG7 | *A. thaliana* | AT1G36730 | | Probable eukaryotic translation initiation factor 5-1 | 0.01 | 0.40 | 2.61E-49 | 5.78E-49 | 0.40 | 2.61E-49 |
| | *O. sativa* | LOC_Os06g48350 | | Os06g0698674 protein | 0.00 | 0.42 | 5.38E-39 | 6.84E-39 | 0.42 | 5.38E-39 |
| | | LOC_Os09g15770 | | Os09g0326900 protein | 0.00 | 0.36 | 4.32E-88 | 6.00E-88 | 0.36 | 4.32E-88 |
| | *S. lycopersicum* | Solyc03g034440.2 | | | 0.00 | 0.41 | 9.46E-149 | 2.73E-148 | 0.41 | 9.46E-149 |
| | *P. trichocarpa* | POPTR_0004s11110 | | Eukaryotic translation initiation factor 5-2 family | 0.01 | 0.43 | 3.05E-177 | 1.57E-176 | 0.43 | 3.05E-177 |
| | | POPTR_0005s14880 | | | 0.01 | 0.47 | 5.20E-129 | 2.36E-128 | 0.47 | 5.20E-129 |
| | *V. vinifera* | VIT_14s0006g01990 | | Putative uncharacterized protein | 0.01 | 0.42 | 7.99E-240 | 7.16E-239 | 0.42 | 7.99E-240 |
| | | VIT_18s0001g04160 | | Putative uncharacterized protein | 0.00 | 0.41 | 2.85E-204 | 2.23E-203 | 0.41 | 2.85E-204 |
| HG8 | *A. thaliana* | AT3G12010 | | C18orf8 | 0.04 | 0.09 | 0.00E+00 | 0.00E+00 | 0.09 | 0.00E+00 |
| | *O. sativa* | LOC_Os10g26140 | | | 0.05 | 0.06 | 0.00E+00 | 0.00E+00 | 0.06 | 0.00E+00 |
| | *P. trichocarpa* | POPTR_0006s20960 | | | 0.05 | 0.09 | 0.00E+00 | 0.00E+00 | 0.09 | 0.00E+00 |
| HG9 | *A. thaliana* | AT1G64140 | | F22C12.10 | 0.00 | 0.03 | 0.00E+00 | 0.00E+00 | 0.03 | 0.00E+00 |
| | | AT5G09670 | | At5g09670 | 0.11 | 0.04 | 1.08E-260 | 5.35E-260 | 0.04 | 1.08E-260 |
| | | AT5G64550 | | Emb\|CAB89363.1 | 0.00 | 0.04 | 9.93E-230 | 3.97E-229 | 0.04 | 9.93E-230 |
| | *O. sativa* | LOC_Os01g43370 | | | 0.00 | 0.02 | 5.71E-277 | 1.53E-276 | 0.02 | 5.71E-277 |
| | | LOC_Os02g15880 | | | 0.00 | 0.04 | 3.34E-284 | 9.63E-284 | 0.04 | 3.34E-284 |
| | | LOC_Os02g36590 | | | 0.00 | 0.04 | 5.13E-249 | 1.10E-248 | 0.04 | 5.13E-249 |
| | | LOC_Os04g38520 | | | 0.00 | 0.04 | 5.96E-260 | 1.40E-259 | 0.04 | 5.96E-260 |
| | | LOC_Os04g54830 | | | 0.01 | 0.05 | 1.10E-242 | 2.22E-242 | 0.05 | 1.10E-242 |
| | | LOC_Os06g33180 | | | 0.00 | 0.04 | 7.67E-297 | 2.61E-296 | 0.04 | 7.67E-297 |
| | *P. trichocarpa* | POPTR_0001s10090 | | | 0.00 | 0.03 | 9.16E-306 | 8.83E-305 | 0.03 | 9.16E-306 |
| | | POPTR_0001s21410 | | | 0.07 | 0.04 | 1.25E-280 | 1.10E-279 | 0.04 | 1.25E-280 |
| | | POPTR_0003s01750 | | | 0.07 | 0.04 | 2.56E-269 | 2.02E-268 | 0.04 | 2.56E-269 |
| | | POPTR_0003s13400 | | | 0.00 | 0.04 | 0.00E+00 | 0.00E+00 | 0.04 | 0.00E+00 |
| | *V. vinifera* | VIT_02s0025g00710 | | | 0.00 | 0.04 | 0.00E+00 | 0.00E+00 | 0.04 | 0.00E+00 |
| | | VIT_11s0065g00090 | | | 0.00 | 0.04 | 0.00E+00 | 0.00E+00 | 0.04 | 0.00E+00 |
| HG9.2 | *P. trichocarpa* | POPTR_0003s01750 | | | 0.00 | 0.27 | 3.82E-85 | 1.56E-84 | 0.27 | 3.82E-85 |
| HG10 | *A. thaliana* | AT4G19110 | | Putative serine/threonine protein kinase | 0.01 | 0.20 | 1.73E-175 | 6.06E-175 | 0.20 | 1.73E-175 |
| | | AT5G45430 | | Protein kinase superfamily protein | 0.00 | 0.19 | 3.46E-173 | 1.19E-172 | 0.19 | 3.46E-173 |
| | *O. sativa* | LOC_Os02g47220 | CDKF-4 | Cyclin-dependent kinase F-4 | 0.13 | 0.22 | 1.88E-22 | 2.31E-22 | 0.22 | 1.88E-22 |
| | | LOC_Os06g02550 | | | 0.00 | 0.23 | 3.57E-65 | 4.78E-65 | 0.23 | 3.57E-65 |
| HG11 | *A. thaliana* | AT4G12430 | TPPF | Probable trehalose-phosphate phosphatase F | 0.01 | 0.06 | 0.00E+00 | 0.00E+00 | 0.06 | 0.00E+00 |
| | | AT4G22590 | TPPG | Probable trehalose-phosphate phosphatase G | 0.02 | 0.05 | 0.00E+00 | 0.00E+00 | 0.05 | 0.00E+00 |
| | *O. sativa* | LOC_Os02g44230 | TPP1 | Probable trehalose-phosphate phosphatase 1 | 0.01 | 0.14 | 1.25E-257 | 2.85E-257 | 0.14 | 1.25E-257 |
| | | LOC_Os10g40550 | TPP2 | Probable trehalose-phosphate phosphatase 2 | 0.01 | 0.12 | 3.94E-187 | 7.04E-187 | 0.12 | 3.94E-187 |
| | *S. lycopersicum* | Solyc03g083960.2 | | Trehalose 6-phosphate phosphatase | 0.00 | 0.05 | 0.00E+00 | 0.00E+00 | 0.05 | 0.00E+00 |
| | *P. trichocarpa* | POPTR_0003s11190 | | Trehalose 6-phosphate phosphatase | 0.02 | 0.10 | 0.00E+00 | 0.00E+00 | 0.10 | 0.00E+00 |
| | | POPTR_0012s14700 | | Trehalose 6-phosphate phosphatase | 0.00 | 0.05 | 0.00E+00 | 0.00E+00 | 0.05 | 0.00E+00 |
| | | POPTR_0015s14810 | | Trehalose 6-phosphate phosphatase | 0.00 | 0.05 | 0.00E+00 | 0.00E+00 | 0.05 | 0.00E+00 |
| | *V. vinifera* | VIT_02s0154g00110 | | Trehalose 6-phosphate phosphatase | 0.02 | 0.04 | 0.00E+00 | 0.00E+00 | 0.04 | 0.00E+00 |
| HG12 | *A. thaliana* | AT1G23150 | | Putative uncharacterized protein At1g23150 | 0.02 | 0.23 | 2.38E-181 | 8.68E-181 | 0.23 | 2.38E-181 |
| | | AT1G70780 | | At1g70780 | 0.02 | 0.21 | 2.47E-255 | 1.12E-254 | 0.21 | 2.47E-255 |
| | *O. sativa* | LOC_Os02g21920 | | | 0.01 | 0.21 | 1.25E-159 | 2.04E-159 | 0.21 | 1.25E-159 |
| | *S. lycopersicum* | Solyc02g091540.2 | | | 0.02 | 0.12 | 8.46E-80 | 2.13E-79 | 0.12 | 8.46E-80 |
| | | Solyc04g007840.2 | | | 0.02 | 0.26 | 5.35E-109 | 1.44E-108 | 0.26 | 5.35E-109 |
| | | Solyc04g074100.2 | | | 0.01 | 0.28 | 8.34E-253 | 4.34E-252 | 0.28 | 8.34E-253 |
| | | POPTR_0004s08870 | | | 0.02 | 0.22 | 6.83E-97 | 2.88E-96 | 0.22 | 6.83E-97 |

| | | | | | | | | | | |
|---|---|---|---|---|---|---|---|---|---|---|
| | *P. trichocarpa* | POPTR_0008s13080 | | | 0.02 | 0.33 | 1.06E-165 | 5.35E-165 | 0.33 | 1.06E-165 |
| | | POPTR_0010s12090 | | | 0.02 | 0.28 | 3.61E-245 | 2.40E-244 | 0.28 | 3.61E-245 |
| HG13 | *A. thaliana* | AT1G48600 | NMT2 | PMEAMT | 0.00 | 0.03 | 2.22E-84 | 5.56E-84 | 0.03 | 2.22E-84 |
| | | AT1G73600 | | S-adenosyl-L-methionine-dependent | 0.00 | 0.03 | 1.93E-100 | 5.23E-100 | 0.03 | 1.93E-100 |
| | | AT3G18000 | NMT1 | XPL1 | 0.00 | 0.03 | 9.49E-165 | 3.13E-164 | 0.03 | 9.49E-165 |
| | *O. sativa* | LOC_Os01g50030 | | | 0.00 | 0.02 | 1.87E-73 | 2.55E-73 | 0.02 | 1.87E-73 |
| | | LOC_Os05g47540 | | Os05g0548900 protein | 0.00 | 0.03 | 1.91E-41 | 2.47E-41 | 0.03 | 1.91E-41 |
| | *P. trichocarpa* | POPTR_0012s04490 | | | 0.00 | 0.05 | 1.08E-134 | 4.98E-134 | 0.05 | 1.08E-134 |
| | *V. vinifera* | VIT_17s0000g02430 | | Putative uncharacterized protein | 0.00 | 0.04 | 1.96E-178 | 1.40E-177 | 0.04 | 1.96E-178 |
| HG14 | *A. thaliana* | AT3G01470 | HAT5 | Homeobox-leucine zipper protein HAT5 | 0.00 | 0.05 | 0.00E+00 | 0.00E+00 | 0.05 | 0.00E+00 |
| | *O. sativa* | LOC_Os02g49700 | HOX16 | Homeobox-leucine zipper protein HOX16 | 0.00 | 0.05 | 0.00E+00 | 0.00E+00 | 0.05 | 0.00E+00 |
| | | LOC_Os08g32080 | HOX5 | Homeobox-leucine zipper protein HOX5 | 0.01 | 0.05 | 0.00E+00 | 0.00E+00 | 0.05 | 0.00E+00 |
| | | LOC_Os09g21180 | | Os09g0379600 protein | 0.07 | 0.04 | 0.00E+00 | 0.00E+00 | 0.04 | 0.00E+00 |
| | *S. lycopersicum* | Solyc02g086930.2 | | Homeodomain leucine zipper protein | 0.00 | 0.05 | 0.00E+00 | 0.00E+00 | 0.05 | 0.00E+00 |
| | | Solyc03g113270.2 | | Homeobox | 0.00 | 0.07 | 9.98E-204 | 3.71E-203 | 0.07 | 9.98E-204 |
| | *P. trichocarpa* | POPTR_0012s07270 | | | 0.06 | 0.08 | 2.70E-255 | 1.83E-254 | 0.08 | 2.70E-255 |
| | | POPTR_0015s07640 | | Putative uncharacterized protein | 0.06 | 0.08 | 6.66E-234 | 4.33E-233 | 0.08 | 6.66E-234 |
| | *V. vinifera* | VIT_01s0026g01550 | | Putative uncharacterized protein | 0.00 | 0.05 | 1.66E-228 | 1.35E-227 | 0.05 | 1.66E-228 |
| | | VIT_14s0066g01440 | ATHB-1 | Putative uncharacterized protein | 0.00 | 0.05 | 0.00E+00 | 0.00E+00 | 0.05 | 0.00E+00 |
| | | VIT_17s0000g05630 | | Putative uncharacterized protein | 0.02 | 0.08 | 1.48E-243 | 1.54E-242 | 0.08 | 1.48E-243 |
| HG15.1 | *A. thaliana* | AT1G29950 | BHLH144 | Transcription factor bHLH144 | 0.00 | 0.10 | 1.54E-135 | 4.72E-135 | 0.10 | 1.54E-135 |
| | | AT5G09460 | BHLH143 | Transcription factor bHLH143 | 0.00 | 0.05 | 1.64E-256 | 7.66E-256 | 0.05 | 1.64E-256 |
| | | AT5G50010 | BHLH145 | Transcription factor bHLH145 | 0.00 | 0.04 | 1.47E-54 | 3.29E-54 | 0.04 | 1.47E-54 |
| | | AT5G64340 | SAC51 | Transcription factor SAC51 | 0.00 | 0.03 | 2.68E-225 | 1.05E-224 | 0.03 | 2.68E-225 |
| | *O. sativa* | LOC_Os01g43680 | | Os01g0626900 protein | 0.10 | 0.11 | 1.91E-127 | 2.92E-127 | 0.11 | 1.91E-127 |
| | | LOC_Os02g21090 | | Os02g0315600 protein | 0.05 | 0.11 | 5.65E-38 | 7.06E-38 | 0.11 | 5.65E-38 |
| | | LOC_Os03g27390 | | Os03g0391700 protein | 0.00 | 0.04 | 9.46E-272 | 2.45E-271 | 0.04 | 9.46E-272 |
| | | LOC_Os03g39432 | | Helix-loop-helix DNA-binding domain containing | 0.03 | 0.10 | 4.12E-97 | 5.84E-97 | 0.10 | 4.12E-97 |
| | *P. trichocarpa* | POPTR_0007s03530 | | | 0.00 | 0.04 | 1.44E-274 | 1.20E-273 | 0.04 | 1.44E-274 |
| | | POPTR_0011s02710 | | basic helix-loop-helix (bHLH) DNA-binding | 0.00 | 0.07 | 1.84E-265 | 1.41E-264 | 0.07 | 1.84E-265 |
| HG15.2 | *A. thaliana* | AT5G09460 | BHLH143 | Transcription factor bHLH143 | 0.00 | 0.39 | 1.17E-02 | 1.55E-02 | 0.39 | 1.17E-02 |
| | | AT5G50010 | BHLH145 | Transcription factor bHLH145 | 0.00 | 0.03 | 4.65E-34 | 1.00E-33 | 0.03 | 4.65E-34 |
| | | AT5G64340 | SAC51 | Transcription factor SAC51 | 0.00 | 0.06 | 2.68E-03 | 3.75E-03 | 0.06 | 2.68E-03 |
| | *P. trichocarpa* | POPTR_0007s03530 | | | 0.00 | 0.04 | 1.17E-12 | 2.92E-12 | 0.04 | 1.17E-12 |
| HG15.3 | *A. thaliana* | AT1G29950 | BHLH144 | Transcription factor bHLH144 | 0.01 | 0.16 | 0.00E+00 | 0.00E+00 | 0.16 | 0.00E+00 |
| | | AT5G09460 | BHLH143 | Transcription factor bHLH143 | 0.04 | 0.07 | 0.00E+00 | 0.00E+00 | 0.07 | 0.00E+00 |
| | | AT5G50010 | BHLH145 | Transcription factor bHLH145 | 0.03 | 0.07 | 0.00E+00 | 0.00E+00 | 0.07 | 0.00E+00 |
| | | AT5G64340 | SAC51 | Transcription factor SAC51 | 0.04 | 0.07 | 0.00E+00 | 0.00E+00 | 0.07 | 0.00E+00 |
| | *O. sativa* | LOC_Os01g43680 | | Os01g0626900 protein | 0.01 | 0.19 | 0.00E+00 | 0.00E+00 | 0.19 | 0.00E+00 |
| | | LOC_Os02g21090 | | Os02g0315600 protein | 0.00 | 0.18 | 3.72E-149 | 5.94E-149 | 0.18 | 3.72E-149 |
| | | LOC_Os03g27390 | | Os03g0391700 protein | 0.01 | 0.06 | 0.00E+00 | 0.00E+00 | 0.06 | 0.00E+00 |
| | | LOC_Os03g39432 | | Helix-loop-helix DNA-binding domain containing | 0.00 | 0.16 | 7.78E-294 | 2.54E-293 | 0.16 | 7.78E-294 |
| | *P. trichocarpa* | POPTR_0007s03530 | | | 0.04 | 0.07 | 0.00E+00 | 0.00E+00 | 0.07 | 0.00E+00 |
| | | POPTR_0011s02710 | | basic helix-loop-helix (bHLH) DNA-binding | 0.01 | 0.17 | 0.00E+00 | 0.00E+00 | 0.17 | 0.00E+00 |
| | | POPTR_0017s07520 | | | 0.04 | 0.07 | 0.00E+00 | 0.00E+00 | 0.07 | 0.00E+00 |
| HG16 | *A. thaliana* | AT3G51630 | WNK5 | Probable serine/threonine-protein kinase WNK5 | 0.00 | 0.06 | 5.82E-245 | 2.45E-244 | 0.06 | 5.82E-245 |
| | *P. trichocarpa* | POPTR_0013s15110 | | | 0.00 | 0.04 | 0.00E+00 | 0.00E+00 | 0.04 | 0.00E+00 |
| | *V. vinifera* | VIT_06s0004g07920 | WNK4 | Putative uncharacterized protein | 0.00 | 0.01 | 0.00E+00 | 0.00E+00 | 0.01 | 0.00E+00 |
| | | VIT_08s0058g01130 | WNK5 | Putative uncharacterized protein | 0.00 | 0.03 | 0.00E+00 | 0.00E+00 | 0.03 | 0.00E+00 |
| HG16.2 | *A. thaliana* | AT3G51630 | WNK5 | Probable serine/threonine-protein kinase WNK5 | 0.00 | 0.10 | 3.02E-55 | 6.85E-55 | 0.10 | 3.02E-55 |
| | *P. trichocarpa* | POPTR_0013s15110 | | | 0.04 | 0.26 | 4.42E-55 | 1.57E-54 | 0.26 | 4.42E-55 |
| HG17 | *A. thaliana* | AT1G58120 | | | 0.08 | 0.10 | 0.00E+00 | 0.00E+00 | 0.10 | 0.00E+00 |
| | | AT3G53400 | | Peptide upstream protein | 0.12 | 0.14 | 7.09E-301 | 4.41E-300 | 0.14 | 7.09E-301 |
| | | AT5G01710 | | Methyltransferase | 0.06 | 0.08 | 1.68E-236 | 6.89E-236 | 0.08 | 1.68E-236 |
| | | AT5G03190 | CPUORF47 | Conserved peptide upstream open reading frame 47 | 0.12 | 0.08 | 5.32E-300 | 3.19E-299 | 0.08 | 5.32E-300 |
| | *S. lycopersicum* | Solyc01g105720.2 | | | 0.06 | 0.10 | 0.00E+00 | 0.00E+00 | 0.10 | 0.00E+00 |
| | *P. trichocarpa* | POPTR_0006s13070 | | | 0.13 | 0.09 | 4.25E-280 | 3.63E-279 | 0.09 | 4.25E-280 |
| | | POPTR_0007s03500 | | | 0.07 | 0.11 | 0.00E+00 | 0.00E+00 | 0.11 | 0.00E+00 |
| | *V. vinifera* | VIT_12s0028g03930 | | Putative uncharacterized protein | 0.06 | 0.13 | 0.00E+00 | 0.00E+00 | 0.13 | 0.00E+00 |
| HG18 | *A. thaliana* | AT4G36990 | HSFB1 | Heat stress transcription factor B-1 | 0.01 | 0.04 | 1.96E-276 | 1.03E-275 | 0.04 | 1.96E-276 |
| | *O. sativa* | LOC_Os09g28354 | HSFB1 | Heat stress transcription factor B-1 | 0.03 | 0.04 | 9.21E-198 | 1.73E-197 | 0.04 | 9.21E-198 |
| | *S. lycopersicum* | Solyc02g090820.2 | | | 0.01 | 0.05 | 1.76E-244 | 8.56E-244 | 0.05 | 1.76E-244 |
| | | Solyc04g016000.2 | | | 0.04 | 0.06 | 9.05E-226 | 3.71E-225 | 0.06 | 9.05E-226 |
| | *P. trichocarpa* | POPTR_0006s04770 | | Heat Stress Transcription Factor family protein | 0.04 | 0.11 | 5.71E-206 | 3.16E-205 | 0.11 | 5.71E-206 |
| | | POPTR_0016s05680 | | Heat Stress Transcription Factor family protein | 0.04 | 0.11 | 1.38E-208 | 7.78E-208 | 0.11 | 1.38E-208 |
| | *V. vinifera* | VIT_07s0031g00670 | HSF4 | Putative uncharacterized protein | 0.01 | 0.03 | 1.90E-249 | 2.07E-248 | 0.03 | 1.90E-249 |
| | | VIT_08s0007g08750 | AT-HSFB3 | Putative uncharacterized protein | 0.04 | 0.15 | 4.28E-192 | 3.16E-191 | 0.15 | 4.28E-192 |
| HG19 | *A. thaliana* | AT5G53590 | | At5g53590 | 0.10 | 0.05 | 0.00E+00 | 0.00E+00 | 0.05 | 0.00E+00 |
| | *O. sativa* | LOC_Os10g36703 | | Os10g0510500 protein | 0.30 | 0.04 | 0.00E+00 | 0.00E+00 | 0.04 | 0.00E+00 |
| | *S. lycopersicum* | Solyc01g091030.2 | | | 0.00 | 0.03 | 0.00E+00 | 0.00E+00 | 0.03 | 0.00E+00 |
| | | Solyc01g096340.2 | | | 0.06 | 0.04 | 2.51E-238 | 1.15E-237 | 0.04 | 2.51E-238 |
| | *P. trichocarpa* | POPTR_0002s14620 | | Auxin-responsive family protein | 0.04 | 0.03 | 0.00E+00 | 0.00E+00 | 0.03 | 0.00E+00 |
| | | POPTR_0014s06240 | | | 0.04 | 0.03 | 0.00E+00 | 0.00E+00 | 0.03 | 0.00E+00 |
| | | POPTR_0015s00920 | | | 0.07 | 0.04 | 0.00E+00 | 0.00E+00 | 0.04 | 0.00E+00 |
| | *V. vinifera* | VIT_15s0048g00530 | | Putative uncharacterized protein | 0.05 | 0.03 | 0.00E+00 | 0.00E+00 | 0.03 | 0.00E+00 |
| | | VIT_16s0098g01150 | | Putative uncharacterized protein | 0.06 | 0.07 | 0.00E+00 | 0.00E+00 | 0.07 | 0.00E+00 |
| HG20 | *A. thaliana* | AT2G37480 | | Putative uncharacterized protein At2g37490 | 0.03 | 0.36 | 2.87E-109 | 8.31E-109 | 0.36 | 2.87E-109 |
| | | AT3G53670 | | Putative uncharacterized protein At3g53670; F4P12.370 | 0.05 | 0.34 | 1.62E-58 | 3.74E-58 | 0.34 | 1.62E-58 |
| | *P. trichocarpa* | POPTR_0006s08210 | | | 0.00 | 0.10 | 8.48E-29 | 2.61E-28 | 0.10 | 8.48E-29 |
| | *V. vinifera* | VIT_08s0040g03310 | | | 0.05 | 0.31 | 5.10E-158 | 3.37E-157 | 0.31 | 5.10E-158 |
| HG21 | *A. thaliana* | AT1G25470 | ERF116 | Ethylene-responsive transcription factor ERF116 | 0.00 | 0.43 | 3.14E-27 | 6.44E-27 | 0.43 | 3.14E-27 |
| | | AT1G68550 | ERF118 | Ethylene-responsive transcription factor ERF118 | 0.01 | 0.41 | 1.16E-68 | 2.78E-68 | 0.41 | 1.16E-68 |
| | | AT3G25890 | ERF119 | Ethylene-responsive transcription factor ERF119 | 0.01 | 0.17 | 4.00E-121 | 1.18E-120 | 0.17 | 4.00E-121 |
| | *O. sativa* | LOC_Os09g13940 | | Os09g0309700 protein | 0.09 | 0.21 | 9.16E-04 | 1.01E-03 | 0.21 | 9.16E-04 |
| | *P. trichocarpa* | POPTR_0010s13540 | | | 0.00 | 0.48 | 7.22E-119 | 3.13E-118 | 0.48 | 7.22E-119 |
| HG23 | *A. thaliana* | AT1G64630 | WNK10 | Probable serine/threonine-protein kinase WNK10 | 0.29 | 0.08 | 3.44E-136 | 1.07E-135 | 0.08 | 3.44E-136 |
| | *P. trichocarpa* | POPTR_0001s11210 | | Kinase family protein | 0.05 | 0.07 | 1.29E-120 | 5.69E-120 | 0.07 | 1.29E-120 |
| | | POPTR_0003s14520 | | Kinase family protein | 0.06 | 0.08 | 1.87E-85 | 7.77E-85 | 0.08 | 1.87E-85 |
| | *V. vinifera* | VIT_02s0025g02360 | | Putative uncharacterized protein | 0.06 | 0.08 | 1.32E-129 | 7.89E-129 | 0.08 | 1.32E-129 |
| HG24 | *A. thaliana* | AT3G22970 | | | 0.15 | 0.21 | 2.67E-05 | 4.20E-05 | 0.21 | 2.67E-05 |
| | | AT4G14620 | | | 0.00 | 0.27 | 2.58E-32 | 5.49E-32 | 0.27 | 2.58E-32 |
| | *P. trichocarpa* | POPTR_0008s15960 | | | 0.02 | 0.29 | 8.15E-44 | 2.74E-43 | 0.29 | 8.15E-44 |
| | | POPTR_0010s09080 | | | 0.01 | 0.26 | 7.20E-55 | 2.53E-54 | 0.26 | 7.20E-55 |
| HG25 | *A. thaliana* | AT3G45240 | GRIK1 | Serine/threonine-protein kinase GRIK1 | 0.00 | 0.04 | 2.18E-161 | 6.92E-161 | 0.04 | 2.18E-161 |
| | | AT5G60550 | GRIK2 | Serine/threonine-protein kinase GRIK2 | 0.00 | 0.04 | 1.68E-161 | 5.44E-161 | 0.04 | 1.68E-161 |
| | *V. vinifera* | VIT_06s0004g02990 | | Putative uncharacterized protein | 0.00 | 0.05 | 4.14E-241 | 3.85E-240 | 0.05 | 4.14E-241 |
| HG26 | *A. thaliana* | AT3G10910 | ATL72 | RING-H2 finger protein ATL72 | 0.27 | 0.16 | 2.31E-21 | 4.51E-21 | 0.16 | 2.31E-21 |
| | *P. trichocarpa* | POPTR_0019s08600 | | | 0.07 | 0.26 | 5.18E-63 | 1.93E-62 | 0.26 | 5.18E-63 |
| | *V. vinifera* | VIT_13s0067g02880 | | Putative uncharacterized protein | 0.03 | 0.22 | 3.01E-81 | 1.64E-80 | 0.22 | 3.01E-81 |
| HG27 | *A. thaliana* | AT4G30960 | CIPK6 | CBL-interacting serine/threonine-protein kinase 6 | 0.01 | 0.22 | 1.39E-187 | 5.18E-187 | 0.22 | 1.39E-187 |
| | *O. sativa* | LOC_Os08g34240 | | Non-specific serine/threonine protein kinase | 0.01 | 0.25 | 3.84E-118 | 5.65E-118 | 0.25 | 3.84E-118 |
| | | LOC_Os09g25100 | | Non-specific serine/threonine protein kinase | 0.01 | 0.21 | 6.86E-175 | 1.17E-174 | 0.21 | 6.86E-175 |
| | *P. trichocarpa* | POPTR_0006s20030 | | Non-specific serine/threonine protein kinase | 0.01 | 0.23 | 2.15E-216 | 1.26E-215 | 0.23 | 2.15E-216 |
| | | POPTR_0018s11770 | | | 0.01 | 0.23 | 2.80E-165 | 1.40E-164 | 0.23 | 2.80E-165 |
| | *V. vinifera* | VIT_09s0070g00160 | | Non-specific serine/threonine protein kinase | 0.01 | 0.24 | 1.21E-134 | 7.57E-134 | 0.24 | 1.21E-134 |
| HG28 | *A. thaliana* | AT1G67480 | | F-box/kelch-repeat protein At1g67480 | 0.01 | 0.34 | 9.60E-20 | 1.85E-19 | 0.34 | 9.60E-20 |
| | *O. sativa* | LOC_Os02g30210 | | Kelch repeat-containing F-box protein-like | 0.01 | 0.33 | 1.04E-07 | 1.20E-07 | 0.33 | 1.04E-07 |
| | | LOC_Os04g31120 | | OSJNBb0089B03.12 protein | 0.01 | 0.33 | 6.16E-10 | 7.46E-10 | 0.33 | 6.16E-10 |
| | *P. trichocarpa* | POPTR_0010s06910 | | | 0.01 | 0.20 | 1.18E-136 | 5.51E-136 | 0.20 | 1.18E-136 |
| | | POPTR_0013s10040 | | | 0.01 | 0.34 | 2.81E-204 | 1.50E-203 | 0.34 | 2.81E-204 |
| HG29 | *A. thaliana* | AT2G22500 | PUMP5 | UCP5 | 0.00 | 0.16 | 0.00E+00 | 0.00E+00 | 0.16 | 0.00E+00 |
| | *O. sativa* | LOC_Os08g37370 | | Os08g0478700 protein | 0.00 | 0.15 | 0.00E+00 | 0.00E+00 | 0.15 | 0.00E+00 |
| | *P. trichocarpa* | POPTR_0002s10460 | | Mitochondrial substrate carrier family protein | 0.00 | 0.15 | 0.00E+00 | 0.00E+00 | 0.15 | 0.00E+00 |
| | *V. vinifera* | VIT_18s0001g07320 | | Putative uncharacterized protein | 0.00 | 0.15 | 0.00E+00 | 0.00E+00 | 0.15 | 0.00E+00 |
| HG30 | *A. thaliana* | AT2G11890 | TTM3 | Triphosphate tunel metalloenzyme 3 | 0.04 | 0.27 | 1.91E-271 | 9.73E-271 | 0.27 | 1.91E-271 |
| HG31 | *A. thaliana* | AT5G01810 | CIPK15 | CBL-interacting serine/threonine-protein kinase 15 | 0.29 | 0.41 | 3.31E-03 | 4.55E-03 | 0.41 | 3.31E-03 |
| | *V. vinifera* | VIT_13s0067g02480 | | Non-specific serine/threonine protein kinase | 0.00 | 0.47 | 4.55E-67 | 2.43E-66 | 0.47 | 4.55E-67 |
| HG32 | *O. sativa* | LOC_Os03g21620 | | Os03g0334000 protein | 0.00 | 0.34 | 9.49E-263 | 2.30E-262 | 0.34 | 9.49E-263 |
| | *S. lycopersicum* | Solyc03g095510.2 | | | 0.00 | 0.33 | 6.68E-293 | 4.01E-292 | 0.33 | 6.68E-293 |
| | *P. trichocarpa* | POPTR_0006s11010 | | Serine/threonine-protein kinase AtPK19 | 0.00 | 0.38 | 1.33E-260 | 9.50E-260 | 0.38 | 1.33E-260 |
| | | POPTR_0016s14530 | | Serine/threonine-protein kinase AtPK19 | 0.00 | 0.35 | 1.56E-256 | 1.08E-255 | 0.35 | 1.56E-256 |
| | *V. vinifera* | VIT_08s0058g01440 | | Putative uncharacterized protein | 0.00 | 0.33 | 3.27E-280 | 4.10E-279 | 0.33 | 3.27E-280 |

| | | | | | | | | | | |
|---|---|---|---|---|---|---|---|---|---|---|
| HG33 | A. thaliana | AT1G30270 | CIPK23 | CBL-interacting serine/threonine-protein kinase 23 | 0.03 | 0.03 | 0.00E+00 | 0.00E+00 | 0.03 | 0.00E+00 |
| | S. lycopersicum | Solyc02g021440.2 | | CBL-interacting serine/threonine-protein kinase 23 | 0.03 | 0.02 | 4.03E-293 | 2.62E-292 | 0.02 | 4.03E-293 |
| HG34 | A. thaliana | AT2G27350 | | OTU-containing deubiquitinating enzyme OTU6 | 0.06 | 0.23 | 2.50E-89 | 6.36E-89 | 0.23 | 2.50E-89 |
| | S. lycopersicum | Solyc06g074220.2 | | | 0.00 | 0.25 | 2.99E-174 | 9.33E-174 | 0.25 | 2.99E-174 |
| | P. trichocarpa | POPTR_0004s20770 | | | 0.01 | 0.25 | 1.98E-212 | 1.14E-211 | 0.25 | 1.98E-212 |
| | | POPTR_0009s16170 | | OTU-like cysteine protease family protein | 0.01 | 0.24 | 2.41E-222 | 1.47E-221 | 0.24 | 2.41E-222 |
| | V. vinifera | VIT_06s0080g00850 | | Putative uncharacterized protein | 0.01 | 0.21 | 6.02E-242 | 5.81E-241 | 0.21 | 6.02E-242 |
| HG35 | A. thaliana | AT2G42880 | MPK20 | Mitogen-activated protein kinase 20 | 0.03 | 0.07 | 1.57E-202 | 5.98E-202 | 0.07 | 1.57E-202 |
| | O. sativa | LOC_Os06g26340 | MPK11 | Mitogen-activated protein kinase 11 | 0.00 | 0.04 | 6.99E-221 | 1.34E-220 | 0.04 | 6.99E-221 |
| | S. lycopersicum | Solyc07g056350.2 | | Mitogen-activated protein kinase 20 | 0.00 | 0.12 | 7.74E-124 | 2.16E-123 | 0.12 | 7.74E-124 |
| | P. trichocarpa | POPTR_0005s22350 | | Mitogen-activated protein kinase | 0.00 | 0.15 | 1.51E-202 | 7.95E-202 | 0.15 | 1.51E-202 |
| HG36 | A. thaliana | AT3G15430 | | MJK13.9 protein | 0.04 | 0.20 | 1.84E-104 | 5.06E-104 | 0.20 | 1.84E-104 |
| | P. trichocarpa | POPTR_0001s41320 | | | 0.04 | 0.24 | 5.13E-64 | 1.94E-63 | 0.24 | 5.13E-64 |
| | V. vinifera | VIT_19s0014g03080 | | | 0.05 | 0.23 | 6.87E-142 | 4.42E-141 | 0.23 | 6.87E-142 |
| HG37 | A. thaliana | AT3G55050 | PP2C6 | Probable protein phosphatase 2C 48 | 0.02 | 0.22 | 1.85E-12 | 3.33E-12 | 0.22 | 1.85E-12 |
| HG38 | A. thaliana | AT4G10170 | PHYL1.1 | Phytolongin Phyl1.1 | 0.12 | 0.15 | 9.62E-30 | 1.99E-29 | 0.15 | 9.62E-30 |
| | V. vinifera | VIT_18s0089g00870 | | Putative uncharacterized protein | 0.15 | 0.21 | 1.55E-63 | 7.93E-63 | 0.21 | 1.55E-63 |
| HG39 | A. thaliana | AT4G12790 | | AT4G12790 protein | 0.07 | 0.19 | 1.15E-58 | 2.68E-58 | 0.19 | 1.15E-58 |
| HG40 | A. thaliana | AT5G02480 | | AT5g02480/T22P11_70 | 0.16 | 0.15 | 2.54E-93 | 6.56E-93 | 0.15 | 2.54E-93 |
| HG41 | A. thaliana | AT5G09330 | NAC082 | NAC domain-containing protein 82 | 0.00 | 0.20 | 1.26E-176 | 4.52E-176 | 0.20 | 1.26E-176 |
| | | AT5G64060 | anac103 | NAC domain containing protein 103 | 0.00 | 0.21 | 1.26E-80 | 3.12E-80 | 0.21 | 1.26E-80 |
| | O. sativa | LOC_Os05g35170 | | NAM-like protein | 0.13 | 0.04 | 1.76E-02 | 1.88E-02 | 0.04 | 1.76E-02 |
| | S. lycopersicum | Solyc03g078120.2 | | | 0.00 | 0.19 | 8.22E-108 | 2.14E-107 | 0.19 | 8.22E-108 |
| | V. vinifera | VIT_04s0044g01220 | | Putative uncharacterized protein | 0.00 | 0.20 | 4.41E-172 | 3.08E-171 | 0.20 | 4.41E-172 |
| HG42 | A. thaliana | AT5G27920 | | | 0.02 | 0.22 | 8.12E-05 | 1.26E-04 | 0.22 | 8.12E-05 |
| | P. trichocarpa | POPTR_0005s02230 | | | 0.11 | 0.24 | 7.84E-28 | 2.39E-27 | 0.24 | 7.84E-28 |
| | | POPTR_0013s01460 | | | 0.05 | 0.22 | 2.24E-20 | 6.27E-20 | 0.22 | 2.24E-20 |
| HG43.1 | A. thaliana | AT4G17980 | anac071 | NAC domain containing protein 71 | 0.00 | 0.19 | 1.04E-23 | 2.08E-23 | 0.19 | 1.04E-23 |
| | | AT5G46590 | anac096 | NAC domain-containing protein 96 | 0.00 | 0.07 | 1.82E-95 | 4.78E-95 | 0.07 | 1.82E-95 |
| | O. sativa | LOC_Os10g42130 | | Os10g0571600 protein | 0.05 | 0.05 | 8.00E-120 | 1.20E-119 | 0.05 | 8.00E-120 |
| | S. lycopersicum | Solyc08g077110.2 | | | 0.00 | 0.09 | 2.57E-191 | 8.72E-191 | 0.09 | 2.57E-191 |
| | V. vinifera | VIT_02s0012g01040 | | Putative uncharacterized protein | 0.00 | 0.05 | 2.08E-243 | 2.09E-242 | 0.05 | 2.08E-243 |
| HG43.2 | P. trichocarpa | POPTR_0003s08830 | | | 0.00 | 0.11 | 1.52E-05 | 3.30E-05 | 0.11 | 1.52E-05 |
| HG44 | P. trichocarpa | POPTR_0009s01700 | | auxin response factor 4 | 0.00 | 0.10 | 4.81E-14 | 1.22E-13 | 0.10 | 4.81E-14 |
| | V. vinifera | VIT_06s0004g03130 | ARF4 | Auxin response factor | 0.00 | 0.10 | 4.86E-05 | 1.23E-04 | 0.10 | 4.86E-05 |
| HG45 | A. thaliana | AT5G63640 | | ENTH/VHS/GAT family protein | 0.02 | 0.29 | 2.27E-09 | 4.01E-09 | 0.29 | 2.27E-09 |
| | P. trichocarpa | POPTR_0004s14580 | | VHS domain-containing family protein | 0.03 | 0.31 | 4.00E-11 | 9.72E-11 | 0.31 | 4.00E-11 |
| HG46 | A. thaliana | AT1G14560 | | CoAc1 | 0.06 | 0.33 | 9.14E-23 | 1.81E-22 | 0.33 | 9.14E-23 |
| | S. lycopersicum | Solyc08g062860.2 | | | 0.06 | 0.21 | 4.26E-224 | 1.66E-223 | 0.21 | 4.26E-224 |
| | P. trichocarpa | POPTR_0006s24280 | | Mitochondrial substrate carrier family protein | 0.04 | 0.22 | 3.14E-263 | 2.29E-262 | 0.22 | 3.14E-263 |
| | V. vinifera | VIT_04s0008g01060 | | Putative uncharacterized protein | 0.14 | 0.22 | 2.03E-269 | 2.42E-268 | 0.22 | 2.03E-269 |
| HG47 | A. thaliana | AT1G68100 | IAR1 | IAA-alanine resistance protein 1 | 0.19 | 0.10 | 4.19E-251 | 1.85E-250 | 0.10 | 4.19E-251 |
| | O. sativa | LOC_Os08g36420 | | IAA-alanine resistance protein 1 | 0.23 | 0.11 | 8.44E-256 | 1.86E-255 | 0.11 | 8.44E-256 |
| | S. lycopersicum | Solyc04g008340.2 | | | 0.19 | 0.10 | 1.09E-254 | 6.05E-254 | 0.10 | 1.09E-254 |
| | V. vinifera | VIT_01s0011g06340 | IAR1 | Putative uncharacterized protein | 0.15 | 0.09 | 8.95E-281 | 1.18E-279 | 0.09 | 8.95E-281 |
| HG48 | A. thaliana | AT1G72820 | | At1g72820/F3N23_2 | 0.05 | 0.27 | 5.30E-04 | 7.74E-04 | 0.27 | 5.30E-04 |
| | P. trichocarpa | POPTR_0018s05460 | | Mitochondrial substrate carrier family protein | 0.13 | 0.06 | 1.16E-04 | 2.36E-04 | 0.06 | 1.16E-04 |
| | V. vinifera | VIT_04s0008g01660 | | Putative uncharacterized protein | 0.11 | 0.05 | 4.92E-52 | 2.29E-51 | 0.05 | 4.92E-52 |
| HG49 | A. thaliana | AT5G14720 | | Protein kinase family protein | 0.00 | 0.02 | 1.49E-04 | 2.30E-04 | 0.02 | 1.49E-04 |
| | S. lycopersicum | Solyc02g086790.2 | | | 0.18 | 0.05 | 1.40E-18 | 3.04E-18 | 0.05 | 1.40E-18 |
| | V. vinifera | VIT_14s0066g01170 | | Putative uncharacterized protein | 0.00 | 0.21 | 1.54E-50 | 6.92E-50 | 0.28 | 3.34E-45 |
| HG50.1 | S. lycopersicum | Solyc10g005080.2 | | | 0.01 | 0.45 | 6.97E-09 | 1.39E-08 | 0.45 | 6.97E-09 |
| | P. trichocarpa | POPTR_0002s18170 | | | 0.00 | 0.42 | 4.45E-08 | 1.02E-07 | 0.42 | 4.45E-08 |
| HG50.2 | O. sativa | LOC_Os08g06110 | | Os08g0157600 protein | 0.30 | 0.16 | 7.60E-08 | 8.90E-08 | 0.16 | 7.60E-08 |
| HG51 | A. thaliana | AT4G15180 | ATXR3 | Histone-lysine N-methyltransferase ATXR3 | 0.00 | 0.10 | 1.71E-04 | 2.61E-04 | 0.10 | 1.71E-04 |
| | V. vinifera | VIT_12s0059g02740 | | Putative uncharacterized protein | 0.00 | 0.15 | 1.66E-24 | 6.71E-24 | 0.15 | 1.66E-24 |
| HG52.1 | P. trichocarpa | POPTR_0006s10720 | | Transducin family protein | 0.02 | 0.17 | 8.36E-24 | 2.43E-23 | 0.17 | 8.36E-24 |
| | | POPTR_0016s13950 | | Transducin family protein | 0.02 | 0.17 | 8.36E-24 | 2.43E-23 | 0.17 | 8.36E-24 |
| | V. vinifera | VIT_08s0058g00890 | | Putative uncharacterized protein | 0.02 | 0.15 | 1.16E-33 | 4.87E-33 | 0.15 | 1.16E-33 |
| HG52.2 | S. lycopersicum | Solyc09g014780.2 | | | 0.00 | 0.10 | 1.98E-03 | 3.21E-03 | 0.10 | 1.98E-03 |
| HG53 | A. thaliana | AT1G62400 | HT1 | Serine/threonine-protein kinase HT1 | 0.12 | 0.11 | 1.27E-68 | 3.02E-68 | 0.11 | 1.27E-68 |
| HG54 | A. thaliana | AT1G65320 | CBSX6 | CBS domain-containing protein CBSX6 | 0.06 | 0.35 | 3.94E-02 | 4.94E-02 | 0.35 | 3.94E-02 |
| | S. lycopersicum | Solyc04g056630.2 | | CBS domain-containing protein CBSX6 | 0.03 | 0.22 | 2.36E-03 | 3.75E-03 | 0.22 | 2.36E-03 |
| | V. vinifera | VIT_04s0023g00310 | | | 0.16 | 0.19 | 3.83E-17 | 1.48E-16 | 0.19 | 3.83E-17 |
| HG55 | O. sativa | LOC_Os03g61760 | SPL6 | Squamosa promoter-binding-like protein 6 | 0.08 | 0.06 | 4.64E-169 | 7.73E-169 | 0.06 | 4.64E-169 |
| | P. trichocarpa | POPTR_0002s18970 | | | 0.02 | 0.04 | 1.05E-229 | 6.67E-229 | 0.04 | 1.05E-229 |
| | | POPTR_0014s10960 | | | 0.02 | 0.04 | 2.72E-227 | 1.69E-226 | 0.04 | 2.72E-227 |
| | V. vinifera | VIT_05s0020g00700 | | Putative uncharacterized protein | 0.00 | 0.04 | 1.05E-239 | 9.10E-239 | 0.04 | 1.05E-239 |
| | | VIT_07s0005g02260 | | Putative uncharacterized protein | 0.00 | 0.04 | 3.30E-251 | 3.76E-250 | 0.04 | 3.30E-251 |
| HG56.1 | S. lycopersicum | Solyc02g076920.2 | | | 0.00 | 0.03 | 9.68E-172 | 2.90E-171 | 0.03 | 9.68E-172 |
| | | Solyc07g053290.2 | | | 0.06 | 0.08 | 5.47E-76 | 1.33E-75 | 0.08 | 5.47E-76 |
| | P. trichocarpa | POPTR_0004s05490 | | | 0.03 | 0.08 | 1.54E-145 | 7.45E-145 | 0.08 | 1.54E-145 |
| | V. vinifera | VIT_10s0003g01170 | | Putative uncharacterized protein | 0.01 | 0.09 | 8.12E-90 | 4.63E-89 | 0.09 | 8.12E-90 |
| HG56.2 | P. trichocarpa | POPTR_0004s05490 | | | 0.00 | 0.03 | 2.60E-20 | 7.19E-20 | 0.03 | 2.60E-20 |
| | V. vinifera | VIT_10s0003g01170 | | Putative uncharacterized protein | 0.02 | 0.02 | 1.42E-87 | 7.92E-87 | 0.02 | 1.42E-87 |
| HG57 | A. thaliana | AT1G54095 | | At1g54095 | 0.00 | 0.06 | 2.29E-02 | 2.93E-02 | 0.06 | 2.29E-02 |
| | | AT1G72510 | | At1g72510 | 0.02 | 0.25 | 1.59E-05 | 2.55E-05 | 0.25 | 1.59E-05 |
| | P. trichocarpa | POPTR_0001s16710 | | | 0.00 | 0.17 | 2.74E-53 | 9.54E-53 | 0.17 | 2.74E-53 |
| | | POPTR_0003s06580 | | | 0.03 | 0.06 | 3.60E-75 | 1.42E-74 | 0.06 | 3.60E-75 |
| | | POPTR_0006s23570 | | | 0.00 | 0.05 | 9.56E-20 | 2.60E-19 | 0.05 | 9.56E-20 |
| HG58 | A. thaliana | AT2G24530 | | Putative uncharacterized protein | 0.00 | 0.11 | 1.95E-08 | 3.35E-08 | 0.11 | 1.95E-08 |
| | V. vinifera | VIT_04s0008g02410 | | Putative uncharacterized protein | 0.00 | 0.45 | 2.82E-07 | 8.62E-07 | 0.45 | 2.82E-07 |
| HG59 | A. thaliana | AT2G35940 | BLH1 | BEL1-like homeodomain protein 1 | 0.02 | 0.44 | 3.24E-09 | 5.68E-09 | 0.44 | 3.24E-09 |
| | P. trichocarpa | POPTR_0006s21950 | | | 0.01 | 0.46 | 2.13E-11 | 5.21E-11 | 0.46 | 2.13E-11 |
| | | POPTR_0016s07040 | | BEL1-like homeodomain 1 family protein | 0.01 | 0.44 | 2.39E-17 | 6.32E-17 | 0.44 | 2.39E-17 |
| HG60 | A. thaliana | AT3G12570 | FYD | FYD | 0.15 | 0.04 | 3.31E-04 | 4.92E-04 | 0.04 | 3.31E-04 |
| | P. trichocarpa | POPTR_0008s05420 | | | 0.11 | 0.02 | 1.18E-04 | 2.37E-04 | 0.02 | 1.18E-04 |
| | | POPTR_0010s21340 | | | 0.12 | 0.05 | 3.35E-04 | 6.54E-04 | 0.05 | 3.35E-04 |
| HG61 | A. thaliana | AT5G55600 | | AT5g55600/MDF20_4 | 0.13 | 0.48 | 4.49E-04 | 6.62E-04 | 0.48 | 4.49E-04 |
| | P. trichocarpa | POPTR_0001s37450 | | agenet domain-containing protein / bromo- | 0.00 | 0.50 | 2.78E-05 | 5.91E-05 | 0.50 | 2.78E-05 |
| HG62 | S. lycopersicum | Solyc04g005430.2 | | | 0.02 | 0.36 | 4.47E-06 | 8.10E-06 | 0.36 | 4.47E-06 |
| | | Solyc05g012450.2 | | | 0.02 | 0.33 | 1.66E-12 | 3.41E-12 | 0.33 | 1.66E-12 |
| | P. trichocarpa | POPTR_0008s09440 | | | 0.00 | 0.31 | 1.72E-04 | 3.44E-04 | 0.31 | 1.72E-04 |
| HG63 | S. lycopersicum | Solyc04g071860.2 | | S-acyltransferase | 0.00 | 0.03 | 3.52E-75 | 8.32E-75 | 0.03 | 3.52E-75 |
| | V. vinifera | VIT_07s0031g01960 | | S-acyltransferase | 0.00 | 0.03 | 7.02E-119 | 4.10E-118 | 0.03 | 7.02E-119 |
| HG64 | P. trichocarpa | POPTR_0002s08640 | | Phosphatidylinositol-4-phosphate 5-kinase family protein | 0.00 | 0.04 | 3.95E-29 | 1.23E-28 | 0.04 | 3.95E-29 |
| | V. vinifera | VIT_18s0001g03290 | | Putative uncharacterized protein | 0.00 | 0.05 | 6.75E-41 | 2.87E-40 | 0.05 | 6.75E-41 |
| HG65 | P. trichocarpa | POPTR_0002s09080 | | Serine hydroxymethyltransferase | 0.15 | 0.05 | 4.26E-05 | 8.92E-05 | 0.05 | 4.26E-05 |
| | V. vinifera | VIT_12s0034g02380 | | Serine hydroxymethyltransferase | 0.18 | 0.05 | 5.24E-05 | 1.32E-04 | 0.05 | 5.24E-05 |
| | | VIT_18s0001g04340 | | Putative uncharacterized protein | 0.12 | 0.20 | 3.07E-04 | 7.21E-04 | 0.20 | 3.07E-04 |
| HG66 | P. trichocarpa | POPTR_0004s20000 | | | 0.02 | 0.23 | 1.98E-78 | 7.91E-78 | 0.23 | 1.98E-78 |
| | | POPTR_0009s15140 | | | 0.02 | 0.27 | 1.36E-65 | 5.21E-65 | 0.27 | 1.36E-65 |
| | V. vinifera | VIT_06s0061g01250 | | | 0.00 | 0.24 | 6.57E-66 | 3.44E-65 | 0.24 | 6.57E-66 |
| HG67 | P. trichocarpa | POPTR_0006s23550 | | | 0.00 | 0.01 | 1.03E-38 | 3.38E-38 | 0.01 | 1.03E-38 |
| | | POPTR_0018s08370 | | | 0.00 | 0.01 | 9.40E-30 | 2.96E-29 | 0.01 | 9.40E-30 |
| | V. vinifera | VIT_04s0008g02100 | CID7 | Putative uncharacterized protein | 0.00 | 0.17 | 9.77E-43 | 4.23E-42 | 0.17 | 9.77E-43 |
| HG68 | P. trichocarpa | POPTR_0011s08940 | | VITAMIN C DEFECTIVE 2 family protein | 0.00 | 0.18 | 5.04E-03 | 8.82E-03 | 0.18 | 5.04E-03 |
| | V. vinifera | VIT_19s0090g01000 | | Putative uncharacterized protein | 0.00 | 0.27 | 6.43E-03 | 1.25E-02 | 0.27 | 6.43E-03 |
| HG69 | P. trichocarpa | POPTR_0013s05200 | | | 0.23 | 0.09 | 4.30E-26 | 1.29E-25 | 0.09 | 4.30E-26 |
| | V. vinifera | VIT_08s0032g00280 | | | 0.13 | 0.06 | 6.64E-47 | 2.92E-46 | 0.06 | 6.64E-47 |
| HG70 | A. thaliana | AT3G52490 | SMXL3 | Protein SMAX1-LIKE 3 | 0.00 | 0.48 | 1.73E-05 | 2.74E-05 | 0.48 | 1.73E-05 |
| HG71 | A. thaliana | AT3G61970 | NGA2 | NGA2 | 0.11 | 0.45 | 3.53E-02 | 4.46E-02 | 0.45 | 3.53E-02 |
| HG72.1 | A. thaliana | AT4G34610 | BLH6 | BEL1-like homeodomain protein 6 | 0.00 | 0.02 | 1.81E-04 | 2.75E-04 | 0.02 | 1.81E-04 |
| HG72.2 | V. vinifera | VIT_03s0038g00050 | | Putative uncharacterized protein | 0.00 | 0.17 | 4.01E-04 | 9.23E-04 | 0.17 | 4.01E-04 |
| HG73 | A. thaliana | AT5G53660 | GRF7 | Growth-regulating factor 7 | 0.00 | 0.41 | 1.94E-08 | 3.35E-08 | 0.41 | 1.94E-08 |
| HG74 | O. sativa | LOC_Os04g36058 | | OSJNBa0064D20.12 protein | 0.00 | 0.19 | 1.06E-05 | 1.18E-05 | 0.19 | 1.06E-05 |
| HG75 | O. sativa | LOC_Os11g16280 | | | 0.08 | 0.13 | 1.63E-144 | 2.55E-144 | 0.13 | 7.00E-123 |
| HG76 | S. lycopersicum | Solyc02g091660.2 | | | 0.00 | 0.06 | 1.11E-02 | 1.64E-02 | 0.06 | 1.11E-02 |
| HG77 | S. lycopersicum | Solyc03g118620.2 | | | 0.13 | 0.06 | 1.40E-18 | 3.04E-18 | 0.06 | 1.40E-18 |
| HG78 | S. lycopersicum | Solyc04g079900.2 | | | 0.15 | 0.22 | 1.70E-02 | 2.41E-02 | 0.22 | 1.70E-02 |
| HG79 | S. lycopersicum | Solyc08g082320.2 | | | 0.15 | 0.34 | 3.05E-02 | 4.10E-02 | 0.34 | 3.05E-02 |
| HG80 | P. trichocarpa | POPTR_0001s30040 | | | 0.03 | 0.30 | 2.31E-45 | 7.83E-45 | 0.30 | 2.31E-45 |
| | | POPTR_0004s21550 | | Nodule inception family protein | 0.01 | 0.25 | 1.55E-49 | 5.33E-49 | 0.25 | 1.55E-49 |

| | | | | | | | | | | |
|---|---|---|---|---|---|---|---|---|---|---|
| | | POPTR_0009s16830 | | RWP-RK domain-containing family protein | 0.03 | 0.28 | 1.43E-33 | 4.64E-33 | 0.28 | 1.43E-33 |
| HG81 | *P. trichocarpa* | POPTR_0001s36900 | | Auxin response factor | 0.00 | 0.13 | 3.90E-107 | 1.67E-106 | 0.13 | 3.90E-107 |
| | | POPTR_0011s09450 | | Auxin response factor | 0.00 | 0.16 | 3.22E-40 | 1.07E-39 | 0.16 | 3.22E-40 |
| HG82 | *P. trichocarpa* | POPTR_0001s38480 | | predicted protein, mRNA | 0.14 | 0.42 | 7.06E-08 | 1.59E-07 | 0.42 | 7.06E-08 |
| HG83 | *P. trichocarpa* | POPTR_0001s41040 | | | 0.17 | 0.10 | 7.75E-03 | 1.28E-02 | 0.10 | 7.75E-03 |
| HG84 | *P. trichocarpa* | POPTR_0002s10310 | | | 0.00 | 0.36 | 2.22E-02 | 3.50E-02 | 0.36 | 2.22E-02 |
| HG85 | *P. trichocarpa* | POPTR_0002s12870 | | Phosphatase 2C family protein | 0.03 | 0.50 | 3.06E-04 | 6.01E-04 | 0.50 | 3.06E-04 |
| HG86 | *P. trichocarpa* | POPTR_0003s08760 | | Glycine cleavage system protein H | 0.00 | 0.44 | 3.97E-04 | 7.71E-04 | 0.44 | 3.97E-04 |
| HG87 | *P. trichocarpa* | POPTR_0004s03900 | | Dof-type zinc finger domain-containing family | 0.03 | 0.47 | 4.82E-18 | 1.29E-17 | 0.47 | 4.82E-18 |
| | | POPTR_0011s04730 | | Dof-type zinc finger domain-containing family | 0.20 | 0.42 | 3.23E-20 | 8.87E-20 | 0.42 | 3.23E-20 |
| HG88 | *P. trichocarpa* | POPTR_0004s05670 | | | 0.00 | 0.32 | 2.89E-04 | 5.73E-04 | 0.32 | 2.89E-04 |
| | | POPTR_0011s07260 | | | 0.00 | 0.36 | 3.22E-03 | 5.79E-03 | 0.36 | 3.22E-03 |
| HG89 | *P. trichocarpa* | POPTR_0004s19490 | | | 0.00 | 0.28 | 3.59E-03 | 6.39E-03 | 0.28 | 3.59E-03 |
| | | POPTR_0009s14620 | | | 0.00 | 0.41 | 5.32E-03 | 9.25E-03 | 0.41 | 5.32E-03 |
| HG90 | *P. trichocarpa* | POPTR_0005s02640 | | Formin-like protein | 0.23 | 0.29 | 4.03E-03 | 7.09E-03 | 0.29 | 4.03E-03 |
| HG91 | *P. trichocarpa* | POPTR_0005s07290 | | | 0.10 | 0.42 | 1.18E-04 | 2.37E-04 | 0.42 | 1.18E-04 |
| | | POPTR_0007s05010 | | Putative uncharacterized protein | 0.10 | 0.35 | 1.29E-02 | 2.08E-02 | 0.35 | 1.29E-02 |
| HG92.1 | *P. trichocarpa* | POPTR_0005s11030 | | Phosphatase 2C family protein | 0.02 | 0.24 | 1.60E-31 | 5.10E-31 | 0.24 | 1.60E-31 |
| | | POPTR_0007s09220 | | | 0.00 | 0.22 | 5.71E-26 | 1.69E-25 | 0.22 | 5.71E-26 |
| HG92.2 | *V. vinifera* | VIT_03s0038g02650 | | Putative uncharacterized protein | 0.00 | 0.02 | 4.34E-21 | 1.73E-20 | 0.02 | 4.34E-21 |
| HG93 | *P. trichocarpa* | POPTR_0005s11480 | | | 0.10 | 0.15 | 5.94E-04 | 1.14E-03 | 0.15 | 5.94E-04 |
| HG94 | *P. trichocarpa* | POPTR_0005s27810 | | RNA recognition motif-containing family protein | 0.00 | 0.43 | 2.43E-03 | 4.48E-03 | 0.43 | 2.43E-03 |
| HG95 | *P. trichocarpa* | POPTR_0006s07660 | | | 0.07 | 0.41 | 5.68E-14 | 1.43E-13 | 0.41 | 5.68E-14 |
| HG96 | *P. trichocarpa* | POPTR_0006s13860 | | | 0.00 | 0.05 | 4.74E-16 | 1.24E-15 | 0.05 | 4.74E-16 |
| | | POPTR_0018s07360 | | | 0.00 | 0.05 | 1.61E-27 | 4.85E-27 | 0.05 | 1.61E-27 |
| HG97 | *P. trichocarpa* | POPTR_0006s14740 | | CCAAT-binding transcription factor family protein | 0.06 | 0.33 | 8.94E-11 | 2.16E-10 | 0.33 | 8.94E-11 |
| HG98 | *P. trichocarpa* | POPTR_0006s22890 | | RNA recognition motif-containing family protein | 0.03 | 0.43 | 2.96E-18 | 7.98E-18 | 0.43 | 2.96E-18 |
| HG99 | *P. trichocarpa* | POPTR_0006s27080 | | | 0.00 | 0.29 | 7.77E-03 | 1.28E-02 | 0.29 | 7.77E-03 |
| HG100 | *P. trichocarpa* | POPTR_0008s10350 | | DNA replication complex GINS protein SLD5 | 0.00 | 0.44 | 7.12E-03 | 1.19E-02 | 0.44 | 7.12E-03 |
| HG101.1 | *P. trichocarpa* | POPTR_0008s15440 | | | 0.00 | 0.14 | 6.29E-03 | 1.06E-02 | 0.14 | 6.29E-03 |
| | | POPTR_0010s09580 | | | 0.00 | 0.13 | 8.65E-03 | 1.41E-02 | 0.13 | 8.65E-03 |
| HG101.2 | *V. vinifera* | VIT_05s0020g03670 | | | 0.00 | 0.17 | 1.35E-04 | 3.26E-04 | 0.17 | 1.35E-04 |
| HG102 | *P. trichocarpa* | POPTR_0008s19310 | | Leucine-rich repeat family protein | 0.12 | 0.29 | 1.89E-05 | 4.06E-05 | 0.29 | 1.89E-05 |
| HG103 | *P. trichocarpa* | POPTR_0009s15460 | | Elongation factor Tu family protein | 0.00 | 0.10 | 1.48E-14 | 3.81E-14 | 0.07 | 1.27E-08 |
| HG104 | *P. trichocarpa* | POPTR_0010s01710 | | Kinesin-like protein | 0.00 | 0.33 | 6.12E-03 | 1.03E-02 | 0.33 | 6.12E-03 |
| HG105 | *P. trichocarpa* | POPTR_0010s09600 | | | 0.03 | 0.46 | 8.15E-06 | 1.78E-05 | 0.46 | 8.15E-06 |
| HG106 | *P. trichocarpa* | POPTR_0010s11700 | | | 0.00 | 0.01 | 1.94E-84 | 7.85E-84 | 0.01 | 1.94E-84 |
| HG107 | *P. trichocarpa* | POPTR_0013s08000 | | | 0.00 | 0.36 | 2.99E-05 | 6.29E-05 | 0.36 | 2.99E-05 |
| HG108 | *P. trichocarpa* | POPTR_0013s13670 | | | 0.00 | 0.48 | 2.64E-02 | 4.11E-02 | 0.48 | 2.64E-02 |
| HG109 | *P. trichocarpa* | POPTR_0014s02470 | | | 0.14 | 0.23 | 9.97E-05 | 2.04E-04 | 0.23 | 9.97E-05 |
| HG110 | *P. trichocarpa* | POPTR_0014s03370 | | | 0.00 | 0.34 | 3.98E-07 | 8.81E-07 | 0.34 | 3.98E-07 |
| HG111 | *P. trichocarpa* | POPTR_0014s09590 | | | 0.00 | 0.13 | 2.59E-23 | 7.45E-23 | 0.13 | 2.59E-23 |
| HG112 | *P. trichocarpa* | POPTR_0014s15970 | | | 0.05 | 0.42 | 3.38E-03 | 6.04E-03 | 0.42 | 3.38E-03 |
| HG113 | *P. trichocarpa* | POPTR_0019s11580 | | | 0.05 | 0.27 | 2.55E-05 | 5.45E-05 | 0.29 | 8.90E-05 |
| HG114 | *P. trichocarpa* | POPTR_0019s13560 | | Glycosyltransferase family 14 family protein | 0.00 | 0.36 | 6.57E-05 | 1.36E-04 | 0.36 | 6.57E-05 |
| HG115 | *V. vinifera* | VIT_01s0010g03770 | | | 0.14 | 0.39 | 1.91E-02 | 3.37E-02 | 0.39 | 1.91E-02 |
| HG116 | *V. vinifera* | VIT_01s0137g00230 | | | 0.00 | 0.35 | 1.31E-04 | 3.18E-04 | 0.35 | 1.31E-04 |
| HG117 | *V. vinifera* | VIT_01s0146g00180 | | Putative uncharacterized protein | 0.11 | 0.04 | 2.35E-202 | 1.79E-201 | 0.04 | 2.35E-202 |
| HG118 | *V. vinifera* | VIT_04s0008g01420 | | Putative uncharacterized protein | 0.00 | 0.32 | 7.62E-03 | 1.46E-02 | 0.32 | 7.62E-03 |
| HG119 | *V. vinifera* | VIT_04s0008g04480 | | Putative uncharacterized protein | 0.00 | 0.24 | 2.96E-05 | 7.59E-05 | 0.24 | 2.96E-05 |
| HG120 | *V. vinifera* | VIT_05s0020g00460 | | Putative uncharacterized protein | 0.06 | 0.47 | 1.56E-03 | 3.31E-03 | 0.47 | 1.56E-03 |
| HG121 | *V. vinifera* | VIT_05s0029g00470 | | | 0.03 | 0.36 | 4.67E-57 | 2.21E-56 | 0.36 | 4.67E-57 |
| HG122 | *V. vinifera* | VIT_06s0004g02980 | | Putative uncharacterized protein | 0.15 | 0.06 | 5.07E-18 | 1.99E-17 | 0.06 | 5.07E-18 |
| HG123 | *V. vinifera* | VIT_11s0016g01790 | | | 0.10 | 0.44 | 1.84E-06 | 5.31E-06 | 0.44 | 1.84E-06 |
| HG124 | *V. vinifera* | VIT_11s0016g02880 | | Putative uncharacterized protein | 0.00 | 0.04 | 5.32E-04 | 1.20E-03 | 0.04 | 5.32E-04 |
| HG125 | *V. vinifera* | VIT_14s0066g00370 | | Putative uncharacterized protein | 0.24 | 0.18 | 2.46E-12 | 8.69E-12 | 0.18 | 2.46E-12 |
| HG126 | *V. vinifera* | VIT_14s0108g00970 | | Putative uncharacterized protein | 0.00 | 0.32 | 8.18E-05 | 2.01E-04 | 0.32 | 8.18E-05 |
| HG127.1 | *V. vinifera* | VIT_18s0001g08100 | | | 0.21 | 0.46 | 1.68E-04 | 4.02E-04 | 0.46 | 1.68E-04 |
| HG127.2 | *V. vinifera* | VIT_04s0008g04480 | | Putative uncharacterized protein | 0.20 | 0.43 | 3.11E-04 | 7.23E-04 | 0.43 | 3.11E-04 |
| HG128 | *V. vinifera* | VIT_18s0001g10680 | PAH2 | Putative uncharacterized protein | 0.00 | 0.32 | 5.29E-03 | 1.05E-02 | 0.32 | 5.29E-03 |
| HG129 | *V. vinifera* | VIT_18s0001g13200 | CKX5 | Putative uncharacterized protein | 0.13 | 0.41 | 6.05E-08 | 1.97E-07 | 0.41 | 6.05E-08 |
| HG130 | *V. vinifera* | VIT_18s0157g00020 | GI | Putative uncharacterized protein | 0.00 | 0.07 | 1.03E-16 | 3.92E-16 | 0.07 | 1.03E-16 |

† Gene description is based on information contained in Ensembl Plants (https://plants.ensembl.org/index.html).

Supplementary Table S2. Taxonomic range of sequence conservation of the CPuORFs.

| HG number | Species | Gene ID | Taxonomic category ** | | | | | | | | | | | | |
|---|---|---|---|---|---|---|---|---|---|---|---|---|---|---|---|
| | | | Lamiids | Asterids* | Malvids | Fabids | Eudicots* | Commelinids | Monocots* | Angiospermae* | Gymnospermae | Polypodiopsida | Embryophyta* | Streptophyta* | Viridiplantae* |
| HG1 | *A. thaliana* | AT1G75390 | 4 | 6 | 3 | 7 | 6 | 3 | 6 | 4 | 0 | 0 | 0 | 0 | 0 |
| | | AT2G18160 | 4 | 6 | 3 | 7 | 6 | 3 | 6 | 4 | 0 | 0 | 0 | 0 | 0 |
| | | AT3G62420 | 3 | 6 | 3 | 6 | 6 | 3 | 6 | 3 | 0 | 0 | 0 | 0 | 0 |
| | | AT4G34590 | 4 | 6 | 3 | 7 | 6 | 3 | 5 | 4 | 0 | 0 | 0 | 0 | 0 |
| | | AT5G49450 | 3 | 6 | 3 | 6 | 6 | 3 | 6 | 3 | 0 | 0 | 0 | 0 | 0 |
| | *O. sativa* | LOC_Os02g03960 | 4 | 6 | 4 | 7 | 6 | 2 | 6 | 4 | 0 | 0 | 0 | 0 | 0 |
| | | LOC_Os03g19370 | 3 | 6 | 4 | 6 | 6 | 2 | 6 | 3 | 0 | 0 | 0 | 0 | 0 |
| | | LOC_Os05g03860 | 4 | 6 | 4 | 7 | 6 | 2 | 6 | 4 | 0 | 0 | 0 | 0 | 0 |
| | | LOC_Os08g26880 | 4 | 6 | 4 | 7 | 6 | 2 | 6 | 4 | 0 | 0 | 0 | 0 | 0 |
| | | LOC_Os09g13570 | 4 | 6 | 4 | 7 | 6 | 2 | 6 | 4 | 0 | 0 | 0 | 0 | 0 |
| | | LOC_Os12g37410 | 4 | 6 | 4 | 7 | 6 | 2 | 6 | 4 | 0 | 0 | 0 | 0 | 0 |
| | *S. lycopersicum* | Solyc01g079480.2 | 3 | 6 | 4 | 7 | 6 | 3 | 6 | 4 | 0 | 0 | 0 | 0 | 0 |
| | | Solyc01g100460.2 | 2 | 5 | 4 | 6 | 6 | 3 | 6 | 4 | 0 | 0 | 0 | 0 | 0 |
| | | Solyc01g109880.2 | 3 | 6 | 4 | 7 | 6 | 3 | 6 | 4 | 0 | 0 | 0 | 0 | 0 |
| | *P. trichocarpa* | POPTR_0002s19720 | 4 | 6 | 4 | 6 | 6 | 3 | 6 | 4 | 0 | 0 | 0 | 0 | 0 |
| | | POPTR_0004s16560 | 4 | 6 | 4 | 6 | 6 | 3 | 5 | 4 | 0 | 0 | 0 | 0 | 0 |
| | | POPTR_0005s25290 | 4 | 6 | 4 | 6 | 6 | 3 | 6 | 4 | 0 | 0 | 0 | 0 | 0 |
| | | POPTR_0007s13380 | 4 | 6 | 4 | 6 | 6 | 3 | 6 | 4 | 0 | 0 | 0 | 0 | 0 |
| | | POPTR_0008s10620 | 3 | 5 | 4 | 5 | 6 | 3 | 6 | 3 | 0 | 0 | 0 | 0 | 0 |
| | | POPTR_0009s12250 | 4 | 6 | 4 | 6 | 6 | 3 | 5 | 4 | 0 | 0 | 0 | 0 | 0 |
| | | POPTR_0010s15280 | 3 | 6 | 4 | 5 | 6 | 3 | 6 | 3 | 0 | 0 | 0 | 0 | 0 |
| | | POPTR_0014s11590 | 4 | 6 | 4 | 6 | 6 | 3 | 6 | 4 | 0 | 0 | 0 | 0 | 0 |
| | *V. vinifera* | VIT_03s0038g04450 | 4 | 6 | 4 | 7 | 5 | 3 | 6 | 4 | 0 | 0 | 0 | 0 | 0 |
| | | VIT_04s0023g02430 | 4 | 6 | 4 | 7 | 5 | 3 | 6 | 4 | 0 | 0 | 0 | 0 | 0 |
| | | VIT_05s0077g01140 | 4 | 6 | 4 | 7 | 5 | 3 | 6 | 4 | 0 | 0 | 0 | 0 | 0 |
| | | VIT_07s0005g01450 | 4 | 6 | 4 | 7 | 5 | 3 | 6 | 4 | 0 | 0 | 0 | 0 | 0 |
| | | VIT_14s0060g01210 | 4 | 6 | 4 | 7 | 5 | 3 | 6 | 4 | 0 | 0 | 0 | 0 | 0 |
| HG2 | *A. thaliana* | AT1G06150 | 3 | 6 | 3 | 6 | 6 | 3 | 5 | 0 | 4 | 2 | 0 | 0 | 0 |
| | | AT2G27230 | 3 | 6 | 3 | 6 | 6 | 3 | 5 | 0 | 4 | 2 | 0 | 0 | 0 |
| | | AT2G31280 | 3 | 6 | 3 | 6 | 6 | 3 | 5 | 0 | 4 | 2 | 0 | 0 | 0 |
| | *O. sativa* | LOC_Os11g06010 | 3 | 6 | 4 | 6 | 6 | 2 | 5 | 0 | 3 | 0 | 0 | 0 | 0 |
| | | LOC_Os12g06330 | 3 | 6 | 4 | 6 | 6 | 2 | 5 | 0 | 3 | 0 | 0 | 0 | 0 |
| | *S. lycopersicum* | Solyc06g074110.2 | 2 | 6 | 4 | 6 | 6 | 3 | 5 | 0 | 4 | 2 | 0 | 0 | 0 |
| | | Solyc09g011170.2 | 2 | 6 | 4 | 6 | 6 | 3 | 5 | 0 | 3 | 1 | 0 | 0 | 0 |
| | | Solyc09g066280.2 | 2 | 6 | 4 | 6 | 6 | 3 | 5 | 0 | 4 | 1 | 0 | 0 | 0 |
| | *P. trichocarpa* | POPTR_0001s22450 | 3 | 6 | 4 | 5 | 6 | 3 | 5 | 0 | 4 | 1 | 0 | 0 | 0 |
| | | POPTR_0005s24390 | 3 | 6 | 4 | 5 | 6 | 3 | 5 | 0 | 4 | 2 | 0 | 0 | 0 |
| | | POPTR_0006s09100 | 3 | 6 | 4 | 5 | 6 | 3 | 5 | 0 | 4 | 1 | 0 | 0 | 0 |
| | *V. vinifera* | VIT_08s0007g00940 | 3 | 6 | 4 | 6 | 5 | 3 | 5 | 0 | 3 | 0 | 0 | 0 | 0 |

| | | | | | | | | | | | | | | | |
|---|---|---|---|---|---|---|---|---|---|---|---|---|---|---|---|
| HG2.2 | *P. trichocarpa* | POPTR_0001s22450 | 0 | 1 | 2 | 5 | 0 | 0 | 0 | 0 | 0 | 0 | 0 | 0 | 0 |
| HG3 | *A. thaliana* | AT3G02470 | 4 | 6 | 3 | 6 | 6 | 3 | 5 | 3 | 6 | 4 | 1 | 3 | 2 |
| | | AT3G25570 | 4 | 6 | 3 | 6 | 6 | 3 | 5 | 3 | 6 | 4 | 1 | 3 | 1 |
| | | AT5G15950 | 4 | 6 | 3 | 6 | 6 | 3 | 5 | 3 | 6 | 4 | 1 | 3 | 2 |
| | *O. sativa* | LOC_Os02g39790 | 4 | 6 | 4 | 6 | 6 | 2 | 5 | 3 | 5 | 4 | 1 | 3 | 3 |
| | | LOC_Os04g42090 | 4 | 6 | 4 | 6 | 6 | 2 | 5 | 3 | 6 | 4 | 1 | 2 | 2 |
| | | LOC_Os09g25620 | 4 | 6 | 4 | 6 | 6 | 2 | 5 | 3 | 5 | 4 | 1 | 3 | 1 |
| | *S. lycopersicum* | Solyc01g010050.2 | 3 | 6 | 4 | 6 | 6 | 3 | 5 | 3 | 6 | 4 | 1 | 3 | 3 |
| | *P. trichocarpa* | POPTR_0004s10660 | 4 | 6 | 4 | 5 | 6 | 3 | 5 | 3 | 6 | 4 | 1 | 3 | 5 |
| | | POPTR_0010s14390 | 4 | 6 | 4 | 5 | 6 | 3 | 5 | 3 | 6 | 4 | 1 | 3 | 3 |
| | | POPTR_0017s14280 | 4 | 6 | 4 | 5 | 6 | 3 | 5 | 3 | 6 | 4 | 1 | 3 | 0 |
| | | POPTR_0018s11040 | 4 | 6 | 4 | 5 | 6 | 3 | 5 | 3 | 6 | 4 | 1 | 3 | 4 |
| HG4 | *A. thaliana* | AT4G25670 | 3 | 5 | 3 | 7 | 6 | 3 | 5 | 2 | 3 | 0 | 0 | 0 | 0 |
| | | AT4G25690 | 3 | 5 | 3 | 7 | 6 | 3 | 5 | 2 | 3 | 0 | 0 | 0 | 0 |
| | | AT5G52550 | 3 | 5 | 3 | 7 | 6 | 3 | 5 | 2 | 3 | 0 | 0 | 0 | 0 |
| | *O. sativa* | LOC_Os02g01360 | 3 | 5 | 4 | 7 | 6 | 2 | 5 | 2 | 3 | 0 | 0 | 0 | 0 |
| | *P. trichocarpa* | POPTR_0004s07800 | 3 | 4 | 4 | 6 | 6 | 3 | 5 | 2 | 3 | 0 | 0 | 0 | 0 |
| | | POPTR_0017s01640 | 3 | 4 | 4 | 6 | 6 | 3 | 5 | 2 | 3 | 0 | 0 | 0 | 0 |
| HG5 | *A. thaliana* | AT5G07840 | 3 | 6 | 3 | 6 | 6 | 3 | 4 | 2 | 3 | 1 | 0 | 0 | 0 |
| | | AT5G61230 | 3 | 6 | 3 | 6 | 6 | 3 | 4 | 3 | 3 | 0 | 0 | 0 | 0 |
| | *O. sativa* | LOC_Os02g01240 | 3 | 6 | 4 | 6 | 6 | 2 | 4 | 2 | 3 | 0 | 0 | 0 | 0 |
| | *P. trichocarpa* | POPTR_0001s01190 | 3 | 6 | 4 | 5 | 6 | 3 | 4 | 2 | 3 | 2 | 0 | 0 | 0 |
| | | POPTR_0003s10350 | 3 | 6 | 4 | 5 | 6 | 3 | 4 | 2 | 3 | 0 | 0 | 0 | 0 |
| | *V. vinifera* | VIT_02s0087g00100 | 3 | 5 | 4 | 6 | 5 | 3 | 4 | 2 | 3 | 2 | 0 | 0 | 0 |
| HG6 | *A. thaliana* | AT2G43020 | 1 | 1 | 2 | 4 | 2 | 1 | 2 | 1 | 0 | 0 | 0 | 0 | 0 |
| | | AT3G59050 | 0 | 1 | 2 | 2 | 0 | 0 | 0 | 1 | 0 | 0 | 0 | 0 | 0 |
| | *S. lycopersicum* | Solyc02g081390.2 | 2 | 6 | 4 | 4 | 5 | 3 | 4 | 1 | 1 | 0 | 0 | 0 | 0 |
| | *P. trichocarpa* | POPTR_0004s07430 | 3 | 6 | 3 | 3 | 5 | 0 | 2 | 0 | 1 | 0 | 0 | 0 | 0 |
| | *V. vinifera* | VIT_04s0043g00220 | 3 | 6 | 4 | 4 | 4 | 3 | 4 | 1 | 1 | 0 | 0 | 0 | 0 |
| HG7 | *A. thaliana* | AT1G36730 | 4 | 6 | 3 | 7 | 6 | 3 | 5 | 1 | 3 | 1 | 1 | 0 | 0 |
| | *O. sativa* | LOC_Os06g48350 | 4 | 6 | 4 | 6 | 6 | 2 | 5 | 1 | 4 | 2 | 1 | 0 | 0 |
| | | LOC_Os09g15770 | 4 | 6 | 4 | 6 | 6 | 2 | 5 | 1 | 4 | 2 | 1 | 1 | 0 |
| | *S. lycopersicum* | Solyc03g034440.2 | 3 | 6 | 4 | 7 | 6 | 3 | 5 | 1 | 5 | 2 | 1 | 1 | 0 |
| | *P. trichocarpa* | POPTR_0004s11110 | 4 | 6 | 4 | 6 | 6 | 3 | 5 | 1 | 5 | 2 | 1 | 0 | 0 |
| | | POPTR_0005s14880 | 4 | 6 | 4 | 6 | 6 | 3 | 5 | 1 | 5 | 1 | 1 | 0 | 0 |
| | *V. vinifera* | VIT_14s0006g01990 | 4 | 6 | 4 | 7 | 5 | 3 | 5 | 1 | 5 | 2 | 1 | 1 | 0 |
| | | VIT_18s0001g04160 | 4 | 6 | 4 | 7 | 5 | 3 | 5 | 1 | 4 | 2 | 1 | 0 | 0 |
| HG8 | *A. thaliana* | AT3G12010 | 3 | 2 | 3 | 6 | 6 | 3 | 4 | 0 | 3 | 2 | 3 | 2 | 0 |
| | *O. sativa* | LOC_Os10g26140 | 3 | 2 | 4 | 6 | 6 | 2 | 4 | 0 | 3 | 2 | 3 | 2 | 0 |
| | *P. trichocarpa* | POPTR_0006s20960 | 3 | 2 | 4 | 5 | 6 | 3 | 4 | 0 | 3 | 2 | 3 | 2 | 0 |
| | *A. thaliana* | AT1G64140 | 4 | 6 | 3 | 6 | 6 | 3 | 4 | 2 | 4 | 0 | 0 | 0 | 0 |
| | | AT5G09670 | 3 | 5 | 3 | 6 | 6 | 3 | 4 | 1 | 4 | 0 | 0 | 0 | 0 |
| | | AT5G64550 | 4 | 5 | 3 | 7 | 6 | 3 | 4 | 1 | 4 | 0 | 0 | 0 | 0 |
| | | LOC_Os01g43370 | 3 | 4 | 4 | 5 | 6 | 2 | 4 | 2 | 4 | 0 | 0 | 0 | 0 |
| | | LOC_Os02g15880 | 3 | 4 | 4 | 5 | 6 | 2 | 4 | 2 | 4 | 0 | 0 | 0 | 0 |

| | | | | | | | | | | | | | | | |
|---|---|---|---|---|---|---|---|---|---|---|---|---|---|---|---|
| HG9 | *O. sativa* | LOC_Os02g36590 | 3 | 5 | 4 | 6 | 6 | 2 | 4 | 1 | 4 | 0 | 0 | 0 | 0 |
| | | LOC_Os04g38520 | 3 | 5 | 4 | 6 | 6 | 2 | 4 | 1 | 4 | 0 | 0 | 0 | 0 |
| | | LOC_Os04g54830 | 3 | 5 | 4 | 6 | 6 | 2 | 4 | 1 | 4 | 0 | 0 | 0 | 0 |
| | | LOC_Os06g33180 | 3 | 5 | 4 | 6 | 6 | 2 | 4 | 2 | 4 | 0 | 0 | 0 | 0 |
| | *P. trichocarpa* | POPTR_0001s10090 | 3 | 5 | 3 | 5 | 6 | 3 | 4 | 1 | 4 | 0 | 0 | 0 | 0 |
| | | POPTR_0001s21410 | 4 | 5 | 4 | 6 | 6 | 3 | 4 | 1 | 4 | 0 | 0 | 0 | 0 |
| | | POPTR_0003s01750 | 4 | 5 | 4 | 6 | 6 | 3 | 4 | 1 | 4 | 0 | 0 | 0 | 0 |
| | | POPTR_0003s13400 | 4 | 6 | 4 | 5 | 6 | 3 | 4 | 2 | 4 | 0 | 0 | 0 | 0 |
| | *V. vinifera* | VIT_02s0025g00710 | 4 | 6 | 4 | 6 | 5 | 3 | 4 | 2 | 4 | 0 | 0 | 0 | 0 |
| | | VIT_11s0065g00090 | 4 | 6 | 4 | 6 | 4 | 3 | 5 | 2 | 4 | 0 | 0 | 0 | 0 |
| HG9.2 | *P. trichocarpa* | POPTR_0003s01750 | 1 | 2 | 2 | 6 | 5 | 1 | 3 | 1 | 1 | 1 | 0 | 0 | 0 |
| HG10 | *A. thaliana* | AT4G19110 | 3 | 3 | 2 | 6 | 3 | 3 | 4 | 0 | 4 | 0 | 0 | 0 | 0 |
| | | AT5G45430 | 3 | 3 | 2 | 6 | 3 | 3 | 4 | 0 | 4 | 0 | 0 | 0 | 0 |
| | *O. sativa* | LOC_Os02g47220 | 3 | 1 | 3 | 4 | 2 | 2 | 2 | 0 | 0 | 0 | 0 | 0 | 0 |
| | | LOC_Os06g02550 | 3 | 3 | 3 | 6 | 3 | 2 | 4 | 0 | 2 | 0 | 0 | 0 | 0 |
| HG11 | *A. thaliana* | AT4G12430 | 3 | 6 | 3 | 5 | 5 | 3 | 4 | 2 | 3 | 0 | 0 | 0 | 0 |
| | | AT4G22590 | 3 | 5 | 3 | 5 | 5 | 3 | 4 | 2 | 3 | 0 | 0 | 0 | 0 |
| | *O. sativa* | LOC_Os02g44230 | 3 | 5 | 4 | 5 | 5 | 2 | 4 | 1 | 2 | 0 | 0 | 0 | 0 |
| | | LOC_Os10g40550 | 3 | 5 | 4 | 5 | 5 | 2 | 4 | 2 | 3 | 0 | 0 | 0 | 0 |
| | *S. lycopersicum* | Solyc03g083960.2 | 2 | 6 | 4 | 5 | 6 | 3 | 4 | 2 | 3 | 0 | 0 | 0 | 0 |
| | *P. trichocarpa* | POPTR_0003s11190 | 3 | 6 | 4 | 5 | 6 | 3 | 4 | 2 | 3 | 0 | 0 | 0 | 0 |
| | | POPTR_0012s14700 | 3 | 6 | 4 | 4 | 6 | 3 | 4 | 2 | 3 | 0 | 0 | 0 | 0 |
| | | POPTR_0015s14810 | 3 | 6 | 4 | 4 | 6 | 3 | 4 | 2 | 3 | 0 | 0 | 0 | 0 |
| | *V. vinifera* | VIT_02s0154g00110 | 3 | 6 | 4 | 5 | 5 | 3 | 4 | 2 | 3 | 0 | 0 | 0 | 0 |
| HG12 | *A. thaliana* | AT1G23150 | 4 | 6 | 3 | 5 | 6 | 3 | 4 | 1 | 3 | 0 | 0 | 0 | 0 |
| | | AT1G70780 | 4 | 6 | 3 | 6 | 6 | 3 | 4 | 1 | 3 | 0 | 0 | 0 | 0 |
| | *O. sativa* | LOC_Os02g21920 | 4 | 6 | 4 | 5 | 6 | 2 | 4 | 1 | 2 | 0 | 0 | 0 | 0 |
| | *S. lycopersicum* | Solyc02g091540.2 | 2 | 5 | 4 | 5 | 5 | 3 | 3 | 0 | 1 | 0 | 0 | 0 | 0 |
| | | Solyc04g007840.2 | 3 | 6 | 4 | 5 | 6 | 3 | 5 | 1 | 2 | 0 | 0 | 0 | 0 |
| | | Solyc04g074100.2 | 3 | 6 | 4 | 6 | 6 | 3 | 5 | 1 | 3 | 0 | 0 | 0 | 0 |
| | *P. trichocarpa* | POPTR_0004s08870 | 4 | 5 | 4 | 4 | 4 | 2 | 4 | 1 | 2 | 0 | 0 | 0 | 0 |
| | | POPTR_0008s13080 | 4 | 6 | 4 | 5 | 6 | 3 | 5 | 1 | 3 | 0 | 0 | 0 | 0 |
| | | POPTR_0010s12090 | 4 | 6 | 4 | 5 | 6 | 3 | 5 | 1 | 3 | 0 | 0 | 0 | 0 |
| HG13 | *A. thaliana* | AT1G48600 | 3 | 2 | 3 | 3 | 1 | 1 | 0 | 0 | 2 | 0 | 2 | 0 | 0 |
| | | AT1G73600 | 3 | 2 | 2 | 6 | 2 | 2 | 1 | 0 | 2 | 0 | 0 | 0 | 0 |
| | | AT3G18000 | 3 | 3 | 3 | 5 | 4 | 1 | 1 | 0 | 2 | 0 | 2 | 0 | 0 |
| | *O. sativa* | LOC_Os01g50030 | 3 | 0 | 3 | 3 | 3 | 1 | 1 | 0 | 2 | 0 | 0 | 0 | 0 |
| | | LOC_Os05g47540 | 3 | 2 | 2 | 3 | 1 | 0 | 0 | 0 | 2 | 0 | 0 | 0 | 0 |
| | *P. trichocarpa* | POPTR_0012s04490 | 3 | 3 | 3 | 3 | 4 | 2 | 1 | 0 | 2 | 0 | 0 | 0 | 0 |
| | *V. vinifera* | VIT_17s0000g02430 | 3 | 3 | 3 | 6 | 3 | 3 | 1 | 0 | 2 | 0 | 0 | 0 | 0 |
| | *A. thaliana* | AT3G01470 | 4 | 6 | 3 | 6 | 5 | 3 | 6 | 1 | 5 | 0 | 0 | 0 | 0 |
| | *O. sativa* | LOC_Os02g49700 | 4 | 6 | 4 | 6 | 5 | 2 | 6 | 1 | 5 | 0 | 0 | 0 | 0 |
| | | LOC_Os08g32080 | 4 | 6 | 4 | 6 | 5 | 2 | 5 | 1 | 4 | 0 | 0 | 0 | 0 |
| | | LOC_Os09g21180 | 4 | 3 | 4 | 6 | 5 | 2 | 5 | 1 | 5 | 0 | 0 | 0 | 0 |
| | *S. lycopersicum* | Solyc02g086930.2 | 3 | 6 | 4 | 6 | 5 | 3 | 5 | 1 | 5 | 0 | 0 | 0 | 0 |

| | | | | | | | | | | | | | | | |
|---|---|---|---|---|---|---|---|---|---|---|---|---|---|---|---|
| HG14 | *S. lycopersicum* | Solyc03g113270.2 | 2 | 6 | 3 | 5 | 5 | 2 | 6 | 1 | 0 | 0 | 0 | 0 | 0 |
| | *P. trichocarpa* | POPTR_0012s07270 | 4 | 6 | 3 | 5 | 5 | 2 | 6 | 1 | 0 | 0 | 0 | 0 | 0 |
| | | POPTR_0015s07640 | 3 | 6 | 3 | 5 | 5 | 2 | 6 | 1 | 0 | 0 | 0 | 0 | 0 |
| | *V. vinifera* | VIT_01s0026g01550 | 4 | 6 | 3 | 6 | 4 | 2 | 6 | 1 | 0 | 0 | 0 | 0 | 0 |
| | | VIT_14s0066g01440 | 4 | 6 | 4 | 6 | 4 | 3 | 5 | 1 | 5 | 0 | 0 | 0 | 0 |
| | | VIT_17s0000g05630 | 3 | 6 | 3 | 6 | 4 | 2 | 6 | 1 | 0 | 0 | 0 | 0 | 0 |
| HG15.1 | *A. thaliana* | AT1G29950 | 1 | 2 | 1 | 6 | 6 | 3 | 2 | 0 | 0 | 0 | 0 | 0 | 0 |
| | | AT5G09460 | 3 | 3 | 3 | 7 | 3 | 3 | 4 | 1 | 4 | 0 | 0 | 0 | 0 |
| | | AT5G50010 | 1 | 0 | 3 | 4 | 2 | 0 | 2 | 0 | 1 | 0 | 0 | 0 | 0 |
| | | AT5G64340 | 3 | 3 | 3 | 7 | 3 | 3 | 4 | 1 | 3 | 0 | 0 | 0 | 0 |
| | *O. sativa* | LOC_Os01g43680 | 2 | 2 | 0 | 6 | 5 | 2 | 4 | 0 | 3 | 0 | 0 | 0 | 0 |
| | | LOC_Os02g21090 | 1 | 2 | 0 | 4 | 2 | 2 | 1 | 0 | 0 | 0 | 0 | 0 | 0 |
| | | LOC_Os03g27390 | 3 | 3 | 4 | 6 | 3 | 2 | 4 | 1 | 4 | 0 | 0 | 0 | 0 |
| | | LOC_Os03g39432 | 3 | 2 | 1 | 6 | 3 | 2 | 2 | 0 | 0 | 0 | 0 | 0 | 0 |
| | *P. trichocarpa* | POPTR_0007s03530 | 3 | 3 | 4 | 6 | 4 | 3 | 4 | 1 | 4 | 0 | 0 | 0 | 0 |
| | | POPTR_0011s02710 | 3 | 4 | 3 | 5 | 6 | 3 | 5 | 0 | 3 | 0 | 0 | 0 | 0 |
| HG15.2 | *A. thaliana* | AT5G09460 | 0 | 0 | 2 | 4 | 0 | 0 | 0 | 0 | 0 | 0 | 0 | 0 | 0 |
| | | AT5G50010 | 1 | 0 | 3 | 6 | 1 | 0 | 0 | 0 | 0 | 0 | 0 | 0 | 0 |
| | | AT5G64340 | 0 | 0 | 1 | 2 | 0 | 0 | 0 | 0 | 0 | 0 | 0 | 0 | 0 |
| | *P. trichocarpa* | POPTR_0007s03530 | 1 | 0 | 3 | 4 | 0 | 0 | 0 | 0 | 0 | 0 | 0 | 0 | 0 |
| HG15.3 | *A. thaliana* | AT1G29950 | 3 | 6 | 3 | 6 | 6 | 3 | 5 | 1 | 4 | 4 | 0 | 0 | 0 |
| | | AT5G09460 | 3 | 6 | 3 | 6 | 6 | 3 | 5 | 1 | 4 | 4 | 0 | 0 | 0 |
| | | AT5G50010 | 3 | 6 | 3 | 6 | 5 | 3 | 5 | 1 | 4 | 4 | 0 | 0 | 0 |
| | | AT5G64340 | 3 | 6 | 3 | 6 | 6 | 3 | 5 | 1 | 4 | 4 | 0 | 0 | 0 |
| | *O. sativa* | LOC_Os01g43680 | 3 | 6 | 4 | 6 | 6 | 2 | 5 | 1 | 4 | 4 | 0 | 0 | 0 |
| | | LOC_Os02g21090 | 3 | 6 | 4 | 6 | 6 | 2 | 5 | 1 | 4 | 4 | 0 | 0 | 0 |
| | | LOC_Os03g27390 | 3 | 6 | 4 | 6 | 6 | 2 | 5 | 1 | 4 | 4 | 0 | 0 | 0 |
| | | LOC_Os03g39432 | 3 | 5 | 4 | 6 | 6 | 2 | 5 | 1 | 4 | 4 | 0 | 0 | 0 |
| | *P. trichocarpa* | POPTR_0007s03530 | 3 | 6 | 4 | 6 | 6 | 3 | 5 | 1 | 4 | 4 | 0 | 0 | 0 |
| | | POPTR_0011s02710 | 3 | 5 | 4 | 5 | 6 | 3 | 5 | 1 | 4 | 4 | 0 | 0 | 0 |
| | | POPTR_0017s07520 | 3 | 6 | 4 | 6 | 6 | 3 | 5 | 1 | 4 | 4 | 0 | 0 | 0 |
| HG16 | *A. thaliana* | AT3G51630 | 3 | 5 | 3 | 7 | 5 | 3 | 3 | 1 | 3 | 0 | 0 | 0 | 0 |
| | *P. trichocarpa* | POPTR_0013s15110 | 3 | 5 | 4 | 6 | 5 | 3 | 3 | 1 | 4 | 0 | 0 | 0 | 0 |
| | *V. vinifera* | VIT_06s0004g07920 | 4 | 5 | 4 | 7 | 4 | 3 | 3 | 1 | 4 | 0 | 0 | 0 | 0 |
| | | VIT_08s0058g01130 | 4 | 5 | 4 | 7 | 4 | 3 | 3 | 1 | 4 | 0 | 0 | 0 | 0 |
| HG16.2 | *A. thaliana* | AT3G51630 | 3 | 4 | 2 | 5 | 2 | 0 | 0 | 0 | 0 | 0 | 0 | 0 | 0 |
| | *P. trichocarpa* | POPTR_0013s15110 | 3 | 4 | 4 | 6 | 5 | 2 | 3 | 0 | 0 | 0 | 0 | 0 | 0 |
| HG17 | *A. thaliana* | AT1G58120 | 3 | 6 | 3 | 6 | 6 | 3 | 4 | 3 | 4 | 2 | 0 | 0 | 0 |
| | | AT3G53400 | 3 | 6 | 3 | 6 | 6 | 3 | 5 | 1 | 4 | 2 | 0 | 0 | 0 |
| | | AT5G01710 | 3 | 6 | 3 | 6 | 6 | 3 | 5 | 2 | 4 | 3 | 0 | 0 | 0 |
| | | AT5G03190 | 3 | 6 | 3 | 6 | 6 | 3 | 5 | 1 | 4 | 3 | 0 | 0 | 0 |
| | *S. lycopersicum* | Solyc01g105720.2 | 2 | 6 | 4 | 6 | 6 | 3 | 5 | 3 | 4 | 2 | 0 | 0 | 0 |
| | *P. trichocarpa* | POPTR_0006s13070 | 3 | 6 | 4 | 5 | 6 | 3 | 5 | 1 | 4 | 2 | 0 | 0 | 0 |
| | | POPTR_0007s03500 | 3 | 6 | 4 | 5 | 6 | 3 | 4 | 1 | 4 | 2 | 0 | 0 | 0 |
| | *V. vinifera* | VIT_12s0028g03930 | 3 | 6 | 4 | 6 | 5 | 3 | 4 | 2 | 4 | 2 | 0 | 0 | 0 |

| | | | | | | | | | | | | | | | |
|---|---|---|---|---|---|---|---|---|---|---|---|---|---|---|---|
| HG18 | *A. thaliana* | AT4G36990 | 4 | 6 | 3 | 6 | 6 | 3 | 2 | 2 | 0 | 0 | 0 | 0 | 0 |
| | *O. sativa* | LOC_Os09g28354 | 4 | 6 | 4 | 6 | 6 | 2 | 2 | 1 | 0 | 0 | 0 | 0 | 0 |
| | *S. lycopersicum* | Solyc02g090820.2 | 3 | 6 | 4 | 6 | 6 | 3 | 2 | 2 | 0 | 0 | 0 | 0 | 0 |
| | | Solyc04g016000.2 | 3 | 4 | 4 | 6 | 6 | 3 | 2 | 2 | 0 | 0 | 0 | 0 | 0 |
| | *P. trichocarpa* | POPTR_0006s04770 | 3 | 5 | 4 | 5 | 5 | 3 | 2 | 1 | 0 | 0 | 0 | 0 | 0 |
| | | POPTR_0016s05680 | 3 | 5 | 4 | 5 | 5 | 3 | 2 | 1 | 0 | 0 | 0 | 0 | 0 |
| | *V. vinifera* | VIT_07s0031g00670 | 4 | 6 | 4 | 6 | 5 | 3 | 2 | 2 | 0 | 0 | 0 | 0 | 0 |
| | | VIT_08s0007g08750 | 3 | 4 | 4 | 6 | 4 | 3 | 2 | 1 | 0 | 0 | 0 | 0 | 0 |
| HG19 | *A. thaliana* | AT5G53590 | 4 | 5 | 3 | 6 | 5 | 3 | 4 | 2 | 5 | 0 | 0 | 0 | 0 |
| | *O. sativa* | LOC_Os10g36703 | 4 | 5 | 4 | 6 | 5 | 2 | 4 | 2 | 5 | 0 | 0 | 0 | 0 |
| | *S. lycopersicum* | Solyc01g091030.2 | 3 | 6 | 4 | 6 | 5 | 3 | 4 | 2 | 5 | 0 | 0 | 0 | 0 |
| | | Solyc01g096340.2 | 3 | 5 | 4 | 6 | 4 | 3 | 3 | 2 | 2 | 0 | 0 | 0 | 0 |
| | *P. trichocarpa* | POPTR_0002s14620 | 4 | 6 | 4 | 5 | 5 | 3 | 4 | 2 | 5 | 0 | 0 | 0 | 0 |
| | | POPTR_0014s06240 | 4 | 6 | 4 | 5 | 5 | 3 | 4 | 3 | 5 | 0 | 0 | 0 | 0 |
| | | POPTR_0015s00920 | 4 | 5 | 4 | 5 | 5 | 3 | 4 | 2 | 5 | 0 | 0 | 0 | 0 |
| | *V. vinifera* | VIT_15s0048g00530 | 4 | 6 | 4 | 6 | 4 | 3 | 4 | 3 | 5 | 0 | 0 | 0 | 0 |
| | | VIT_16s0098g01150 | 4 | 5 | 4 | 6 | 4 | 3 | 4 | 2 | 5 | 0 | 0 | 0 | 0 |
| HG20 | *A. thaliana* | AT2G37480 | 3 | 5 | 3 | 5 | 3 | 2 | 4 | 0 | 0 | 0 | 0 | 0 | 0 |
| | | AT3G53670 | 3 | 4 | 3 | 5 | 3 | 2 | 4 | 0 | 0 | 0 | 0 | 0 | 0 |
| | *P. trichocarpa* | POPTR_0006s08210 | 1 | 2 | 3 | 4 | 1 | 2 | 0 | 0 | 0 | 0 | 0 | 0 | 0 |
| | *V. vinifera* | VIT_08s0040g03310 | 3 | 4 | 4 | 5 | 3 | 2 | 4 | 0 | 0 | 0 | 0 | 0 | 0 |
| HG21 | *A. thaliana* | AT1G25470 | 4 | 6 | 3 | 6 | 6 | 1 | 1 | 0 | 0 | 0 | 0 | 0 | 0 |
| | | AT1G68550 | 4 | 6 | 3 | 6 | 6 | 1 | 2 | 0 | 0 | 0 | 0 | 0 | 0 |
| | | AT3G25890 | 3 | 6 | 3 | 6 | 5 | 0 | 0 | 0 | 0 | 0 | 0 | 0 | 0 |
| | *O. sativa* | LOC_Os09g13940 | 0 | 0 | 0 | 0 | 0 | 2 | 4 | 0 | 0 | 0 | 0 | 0 | 0 |
| | *P. trichocarpa* | POPTR_0010s13540 | 3 | 6 | 4 | 5 | 6 | 1 | 2 | 0 | 0 | 0 | 0 | 0 | 0 |
| HG23 | *A. thaliana* | AT1G64630 | 2 | 4 | 3 | 6 | 6 | 2 | 1 | 0 | 0 | 0 | 0 | 0 | 0 |
| | *P. trichocarpa* | POPTR_0001s11210 | 3 | 6 | 4 | 5 | 5 | 2 | 2 | 0 | 0 | 0 | 0 | 0 | 0 |
| | | POPTR_0003s14520 | 3 | 5 | 4 | 5 | 5 | 2 | 3 | 1 | 0 | 0 | 0 | 0 | 0 |
| | *V. vinifera* | VIT_02s0025g02360 | 3 | 5 | 4 | 6 | 5 | 2 | 3 | 0 | 0 | 0 | 0 | 0 | 0 |
| HG24 | *A. thaliana* | AT3G22970 | 0 | 2 | 0 | 3 | 1 | 0 | 0 | 0 | 0 | 0 | 0 | 0 | 0 |
| | | AT4G14620 | 2 | 4 | 3 | 5 | 1 | 0 | 0 | 0 | 0 | 0 | 0 | 0 | 0 |
| | *P. trichocarpa* | POPTR_0008s15960 | 3 | 4 | 2 | 4 | 2 | 0 | 0 | 0 | 0 | 0 | 0 | 0 | 0 |
| | | POPTR_0010s09080 | 3 | 5 | 2 | 4 | 3 | 0 | 0 | 0 | 0 | 0 | 0 | 0 | 0 |
| HG25 | *A. thaliana* | AT3G45240 | 4 | 4 | 3 | 6 | 5 | 1 | 1 | 1 | 1 | 0 | 0 | 0 | 0 |
| | | AT5G60550 | 4 | 4 | 3 | 7 | 5 | 0 | 1 | 1 | 1 | 0 | 0 | 0 | 0 |
| | *V. vinifera* | VIT_06s0004g02990 | 4 | 5 | 4 | 7 | 5 | 1 | 1 | 2 | 2 | 0 | 0 | 0 | 0 |
| HG26 | *A. thaliana* | AT3G10910 | 2 | 3 | 2 | 5 | 1 | 0 | 0 | 0 | 0 | 0 | 0 | 0 | 0 |
| | *P. trichocarpa* | POPTR_0019s08600 | 2 | 5 | 3 | 5 | 1 | 0 | 0 | 0 | 0 | 0 | 0 | 0 | 0 |
| | *V. vinifera* | VIT_13s0067g02880 | 2 | 4 | 4 | 6 | 2 | 0 | 0 | 0 | 0 | 0 | 0 | 0 | 0 |
| HG27 | *A. thaliana* | AT4G30960 | 4 | 6 | 3 | 7 | 6 | 3 | 4 | 2 | 0 | 0 | 0 | 0 | 0 |
| | *O. sativa* | LOC_Os08g34240 | 4 | 6 | 4 | 7 | 6 | 2 | 5 | 2 | 0 | 0 | 0 | 0 | 0 |
| | | LOC_Os09g25100 | 4 | 6 | 4 | 7 | 6 | 2 | 5 | 2 | 0 | 0 | 0 | 0 | 0 |
| | *P. trichocarpa* | POPTR_0006s20030 | 4 | 6 | 4 | 6 | 6 | 3 | 4 | 2 | 0 | 0 | 0 | 0 | 0 |
| | | POPTR_0018s11770 | 4 | 6 | 4 | 6 | 6 | 3 | 4 | 1 | 0 | 0 | 0 | 0 | 0 |

| | | | | | | | | | | | | | | | |
|---|---|---|---|---|---|---|---|---|---|---|---|---|---|---|---|
| | *V. vinifera* | VIT_09s0070g00160 | 4 | 6 | 4 | 7 | 5 | 3 | 5 | 2 | 0 | 0 | 0 | 0 | 0 |
| HG28 | *A. thaliana* | AT1G67480 | 4 | 5 | 3 | 7 | 4 | 3 | 5 | 0 | 0 | 0 | 0 | 0 | 0 |
| | *O. sativa* | LOC_Os02g30210 | 2 | 3 | 4 | 7 | 5 | 2 | 5 | 0 | 0 | 0 | 0 | 0 | 0 |
| | | LOC_Os04g31120 | 1 | 2 | 4 | 7 | 5 | 2 | 5 | 0 | 0 | 0 | 0 | 0 | 0 |
| | *P. trichocarpa* | POPTR_0010s06910 | 3 | 5 | 4 | 6 | 4 | 1 | 2 | 0 | 0 | 0 | 0 | 0 | 0 |
| | | POPTR_0013s10040 | 4 | 5 | 4 | 6 | 5 | 3 | 5 | 0 | 1 | 0 | 0 | 0 | 0 |
| HG29 | *A. thaliana* | AT2G22500 | 4 | 5 | 3 | 7 | 6 | 3 | 5 | 1 | 3 | 0 | 0 | 0 | 0 |
| | *O. sativa* | LOC_Os08g37370 | 4 | 5 | 4 | 7 | 6 | 2 | 5 | 1 | 3 | 0 | 0 | 0 | 0 |
| | *P. trichocarpa* | POPTR_0002s10460 | 4 | 4 | 4 | 6 | 6 | 3 | 5 | 1 | 3 | 0 | 0 | 0 | 0 |
| | *V. vinifera* | VIT_18s0001g07320 | 4 | 5 | 4 | 7 | 5 | 3 | 5 | 1 | 3 | 0 | 0 | 0 | 0 |
| HG30 | *A. thaliana* | AT2G11890 | 3 | 6 | 2 | 6 | 5 | 3 | 4 | 2 | 3 | 0 | 1 | 2 | 1 |
| HG31 | *A. thaliana* | AT5G01810 | 0 | 0 | 3 | 4 | 1 | 1 | 0 | 0 | 0 | 0 | 0 | 0 | 0 |
| | *V. vinifera* | VIT_13s0067g02480 | 4 | 6 | 4 | 6 | 5 | 2 | 4 | 1 | 0 | 0 | 0 | 0 | 0 |
| HG32 | *O. sativa* | LOC_Os03g21620 | 3 | 3 | 4 | 6 | 5 | 2 | 5 | 3 | 2 | 0 | 0 | 0 | 0 |
| | *S. lycopersicum* | Solyc03g095510.2 | 2 | 4 | 4 | 6 | 6 | 3 | 5 | 3 | 2 | 0 | 0 | 0 | 0 |
| | *P. trichocarpa* | POPTR_0006s11010 | 3 | 4 | 4 | 5 | 5 | 3 | 5 | 3 | 2 | 0 | 0 | 0 | 0 |
| | | POPTR_0016s14530 | 3 | 4 | 4 | 5 | 5 | 3 | 5 | 3 | 2 | 0 | 0 | 0 | 0 |
| | *V. vinifera* | VIT_08s0058g01440 | 3 | 4 | 4 | 6 | 4 | 3 | 5 | 3 | 2 | 0 | 0 | 0 | 0 |
| HG33 | *A. thaliana* | AT1G30270 | 4 | 5 | 3 | 6 | 5 | 2 | 5 | 1 | 4 | 0 | 0 | 0 | 0 |
| | *S. lycopersicum* | Solyc02g021440.2 | 3 | 4 | 4 | 6 | 5 | 1 | 5 | 1 | 4 | 0 | 0 | 0 | 0 |
| HG34 | *A. thaliana* | AT2G27350 | 3 | 2 | 3 | 5 | 5 | 1 | 4 | 1 | 0 | 0 | 0 | 0 | 0 |
| | *S. lycopersicum* | Solyc06g074220.2 | 2 | 3 | 4 | 5 | 6 | 3 | 4 | 1 | 4 | 0 | 0 | 0 | 0 |
| | *P. trichocarpa* | POPTR_0004s20770 | 3 | 3 | 4 | 4 | 6 | 2 | 4 | 2 | 4 | 0 | 0 | 0 | 0 |
| | | POPTR_0009s16170 | 3 | 3 | 4 | 4 | 6 | 2 | 4 | 2 | 4 | 0 | 0 | 0 | 0 |
| | *V. vinifera* | VIT_06s0080g00850 | 3 | 3 | 4 | 5 | 5 | 3 | 4 | 2 | 4 | 0 | 0 | 0 | 0 |
| HG35 | *A. thaliana* | AT2G42880 | 3 | 6 | 2 | 5 | 5 | 3 | 5 | 2 | 4 | 0 | 0 | 0 | 0 |
| | *O. sativa* | LOC_Os06g26340 | 3 | 6 | 4 | 5 | 5 | 2 | 5 | 2 | 4 | 0 | 0 | 0 | 0 |
| | *S. lycopersicum* | Solyc07g056350.2 | 2 | 6 | 4 | 5 | 5 | 3 | 5 | 2 | 4 | 0 | 0 | 0 | 0 |
| | *P. trichocarpa* | POPTR_0005s22350 | 3 | 6 | 4 | 4 | 5 | 3 | 5 | 2 | 4 | 0 | 0 | 0 | 0 |
| HG36 | *A. thaliana* | AT3G15430 | 4 | 4 | 3 | 6 | 4 | 0 | 0 | 0 | 0 | 0 | 0 | 0 | 0 |
| | *P. trichocarpa* | POPTR_0001s41320 | 3 | 2 | 3 | 5 | 4 | 0 | 0 | 0 | 0 | 0 | 0 | 0 | 0 |
| | *V. vinifera* | VIT_19s0014g03080 | 4 | 4 | 4 | 6 | 4 | 0 | 0 | 0 | 0 | 0 | 0 | 0 | 0 |
| HG37 | *A. thaliana* | AT3G55050 | 0 | 3 | 3 | 3 | 1 | 0 | 0 | 0 | 0 | 0 | 0 | 0 | 0 |
| HG38 | *A. thaliana* | AT4G10170 | 2 | 3 | 2 | 4 | 4 | 0 | 0 | 0 | 0 | 0 | 0 | 0 | 0 |
| | *V. vinifera* | VIT_18s0089g00870 | 2 | 3 | 3 | 4 | 3 | 0 | 0 | 0 | 0 | 0 | 0 | 0 | 0 |
| HG39 | *A. thaliana* | AT4G12790 | 3 | 2 | 2 | 6 | 4 | 1 | 2 | 1 | 1 | 0 | 0 | 0 | 0 |
| HG40 | *A. thaliana* | AT5G02480 | 3 | 4 | 2 | 5 | 3 | 2 | 1 | 0 | 0 | 0 | 0 | 0 | 0 |
| HG41 | *A. thaliana* | AT5G09330 | 3 | 6 | 3 | 6 | 5 | 3 | 4 | 0 | 0 | 0 | 0 | 0 | 0 |
| | | AT5G64060 | 3 | 5 | 2 | 6 | 3 | 2 | 3 | 0 | 0 | 0 | 0 | 0 | 0 |
| | *O. sativa* | LOC_Os05g35170 | 0 | 0 | 0 | 0 | 0 | 2 | 0 | 0 | 0 | 0 | 0 | 0 | 0 |
| | *S. lycopersicum* | Solyc03g078120.2 | 2 | 6 | 4 | 5 | 2 | 3 | 4 | 1 | 0 | 0 | 0 | 0 | 0 |
| | *V. vinifera* | VIT_04s0044g01220 | 3 | 5 | 4 | 6 | 3 | 2 | 3 | 0 | 0 | 0 | 0 | 0 | 0 |
| HG42 | *A. thaliana* | AT5G27920 | 0 | 0 | 2 | 1 | 2 | 0 | 0 | 0 | 0 | 0 | 0 | 0 | 0 |
| | *P. trichocarpa* | POPTR_0005s02230 | 1 | 1 | 3 | 3 | 4 | 0 | 1 | 0 | 0 | 0 | 0 | 0 | 0 |
| | | POPTR_0013s01460 | 1 | 1 | 3 | 2 | 3 | 0 | 1 | 0 | 0 | 0 | 0 | 0 | 0 |

| | | | | | | | | | | | | | | | |
|---|---|---|---|---|---|---|---|---|---|---|---|---|---|---|---|
| HG43.1 | *A. thaliana* | AT4G17980 | 1 | 4 | 2 | 3 | 3 | 1 | 0 | 0 | 0 | 0 | 0 | 0 | 0 |
| | | AT5G46590 | 2 | 5 | 3 | 5 | 5 | 3 | 4 | 1 | 0 | 0 | 0 | 0 | 0 |
| | *O. sativa* | LOC_Os10g42130 | 3 | 4 | 4 | 5 | 5 | 2 | 4 | 1 | 0 | 0 | 0 | 0 | 0 |
| | *S. lycopersicum* | Solyc08g077110.2 | 2 | 5 | 4 | 5 | 5 | 3 | 4 | 1 | 0 | 0 | 0 | 0 | 0 |
| | *V. vinifera* | VIT_02s0012g01040 | 3 | 5 | 4 | 5 | 4 | 3 | 4 | 1 | 0 | 0 | 0 | 0 | 0 |
| HG43.2 | *P. trichocarpa* | POPTR_0003s08830 | 0 | 1 | 1 | 2 | 0 | 0 | 0 | 0 | 0 | 0 | 0 | 0 | 0 |
| HG44 | *P. trichocarpa* | POPTR_0009s01700 | 2 | 1 | 3 | 3 | 3 | 0 | 0 | 0 | 0 | 0 | 0 | 0 | 0 |
| | *V. vinifera* | VIT_06s0004g03130 | 2 | 1 | 2 | 4 | 1 | 0 | 0 | 0 | 0 | 0 | 0 | 0 | 0 |
| HG45 | *A. thaliana* | AT5G63640 | 0 | 0 | 1 | 4 | 1 | 0 | 0 | 0 | 0 | 0 | 0 | 0 | 0 |
| | *P. trichocarpa* | POPTR_0004s14580 | 0 | 0 | 3 | 4 | 1 | 0 | 0 | 0 | 0 | 0 | 0 | 0 | 0 |
| HG46 | *A. thaliana* | AT1G14560 | 2 | 4 | 3 | 5 | 3 | 0 | 0 | 0 | 0 | 0 | 0 | 0 | 0 |
| | *S. lycopersicum* | Solyc08g062860.2 | 2 | 5 | 4 | 6 | 5 | 2 | 5 | 3 | 3 | 1 | 0 | 0 | 0 |
| | *P. trichocarpa* | POPTR_0006s24280 | 3 | 5 | 4 | 5 | 5 | 2 | 5 | 3 | 3 | 1 | 0 | 0 | 0 |
| | *V. vinifera* | VIT_04s0008g01060 | 3 | 5 | 4 | 6 | 4 | 2 | 5 | 3 | 3 | 1 | 0 | 0 | 0 |
| HG47 | *A. thaliana* | AT1G68100 | 4 | 6 | 3 | 6 | 5 | 3 | 3 | 1 | 0 | 0 | 0 | 0 | 0 |
| | *O. sativa* | LOC_Os08g36420 | 4 | 5 | 4 | 5 | 6 | 2 | 4 | 1 | 0 | 0 | 0 | 0 | 0 |
| | *S. lycopersicum* | Solyc04g008340.2 | 3 | 5 | 4 | 6 | 6 | 3 | 4 | 1 | 0 | 0 | 0 | 0 | 0 |
| | *V. vinifera* | VIT_01s0011g06340 | 4 | 5 | 4 | 6 | 5 | 3 | 3 | 1 | 0 | 0 | 0 | 0 | 0 |
| HG48 | *A. thaliana* | AT1G72820 | 1 | 0 | 1 | 5 | 0 | 0 | 0 | 0 | 0 | 0 | 0 | 0 | 0 |
| | *P. trichocarpa* | POPTR_0018s05460 | 0 | 1 | 1 | 1 | 0 | 0 | 0 | 0 | 0 | 0 | 0 | 0 | 0 |
| | *V. vinifera* | VIT_04s0008g01660 | 3 | 4 | 2 | 5 | 2 | 0 | 0 | 0 | 0 | 0 | 0 | 0 | 0 |
| HG49 | *A. thaliana* | AT5G14720 | 0 | 0 | 1 | 2 | 0 | 0 | 0 | 0 | 0 | 0 | 0 | 0 | 0 |
| | *S. lycopersicum* | Solyc02g086790.2 | 0 | 0 | 4 | 2 | 1 | 0 | 1 | 0 | 0 | 0 | 0 | 0 | 0 |
| | *V. vinifera* | VIT_14s0066g01170 | 0 | 4 | 4 | 5 | 3 | 0 | 1 | 0 | 0 | 0 | 0 | 0 | 0 |
| HG50.1 | *S. lycopersicum* | Solyc10g005080.2 | 2 | 2 | 2 | 6 | 0 | 0 | 0 | 0 | 0 | 0 | 0 | 0 | 0 |
| | *P. trichocarpa* | POPTR_0002s18170 | 2 | 2 | 3 | 3 | 0 | 0 | 0 | 0 | 0 | 0 | 0 | 0 | 0 |
| HG50.2 | *O. sativa* | LOC_Os08g06110 | 0 | 0 | 0 | 0 | 0 | 2 | 3 | 0 | 0 | 0 | 0 | 0 | 0 |
| HG51 | *A. thaliana* | AT4G15180 | 0 | 0 | 3 | 2 | 0 | 0 | 0 | 0 | 0 | 0 | 0 | 0 | 0 |
| | *V. vinifera* | VIT_12s0059g02740 | 2 | 2 | 4 | 5 | 2 | 0 | 0 | 0 | 0 | 0 | 0 | 0 | 0 |
| HG52.1 | *P. trichocarpa* | POPTR_0006s10720 | 0 | 6 | 4 | 4 | 4 | 2 | 2 | 0 | 0 | 0 | 0 | 0 | 0 |
| | | POPTR_0016s13950 | 0 | 6 | 4 | 4 | 4 | 2 | 2 | 0 | 0 | 0 | 0 | 0 | 0 |
| | *V. vinifera* | VIT_08s0058g00890 | 1 | 6 | 4 | 5 | 4 | 2 | 2 | 0 | 0 | 0 | 0 | 0 | 0 |
| HG52.2 | *S. lycopersicum* | Solyc09g014780.2 | 2 | 0 | 0 | 0 | 0 | 0 | 0 | 0 | 0 | 0 | 0 | 0 | 0 |
| HG53 | *A. thaliana* | AT1G62400 | 3 | 2 | 3 | 5 | 2 | 0 | 0 | 0 | 0 | 0 | 0 | 0 | 0 |
| HG54 | *A. thaliana* | AT1G65320 | 2 | 0 | 0 | 0 | 0 | 0 | 0 | 0 | 1 | 0 | 0 | 0 | 0 |
| | *S. lycopersicum* | Solyc04g056630.2 | 1 | 1 | 1 | 0 | 0 | 0 | 0 | 0 | 0 | 0 | 0 | 0 | 0 |
| | *V. vinifera* | VIT_04s0023g00310 | 2 | 1 | 2 | 1 | 1 | 0 | 0 | 0 | 0 | 0 | 0 | 0 | 0 |
| HG55 | *O. sativa* | LOC_Os03g61760 | 3 | 5 | 3 | 5 | 3 | 2 | 4 | 1 | 0 | 0 | 0 | 0 | 0 |
| | *P. trichocarpa* | POPTR_0002s18970 | 3 | 3 | 4 | 5 | 5 | 3 | 4 | 1 | 0 | 0 | 0 | 0 | 0 |
| | | POPTR_0014s10960 | 3 | 3 | 4 | 5 | 5 | 3 | 4 | 1 | 0 | 0 | 0 | 0 | 0 |
| | *V. vinifera* | VIT_05s0020g00700 | 3 | 4 | 4 | 6 | 4 | 3 | 4 | 1 | 0 | 0 | 0 | 0 | 0 |
| | | VIT_07s0005g02260 | 3 | 4 | 4 | 6 | 4 | 3 | 4 | 1 | 0 | 0 | 0 | 0 | 0 |
| HG56.1 | *S. lycopersicum* | Solyc02g076920.2 | 2 | 5 | 3 | 5 | 6 | 2 | 3 | 0 | 0 | 0 | 0 | 0 | 0 |
| | | Solyc07g053290.2 | 2 | 5 | 3 | 4 | 2 | 2 | 4 | 1 | 1 | 0 | 0 | 0 | 0 |
| | *P. trichocarpa* | POPTR_0004s05490 | 3 | 5 | 3 | 4 | 5 | 3 | 4 | 0 | 0 | 0 | 0 | 0 | 0 |

| | | | | | | | | | | | | | | | |
|---|---|---|---|---|---|---|---|---|---|---|---|---|---|---|---|
| | *V. vinifera* | VIT_10s0003g01170 | 3 | 5 | 3 | 5 | 4 | 3 | 4 | 0 | 0 | 0 | 0 | 0 | 0 |
| HG56.2 | *P. trichocarpa* | POPTR_0004s05490 | 1 | 2 | 2 | 3 | 1 | 0 | 0 | 0 | 0 | 0 | 0 | 0 | 0 |
| | *V. vinifera* | VIT_10s0003g01170 | 3 | 5 | 3 | 4 | 4 | 0 | 0 | 0 | 0 | 0 | 0 | 0 | 0 |
| HG57 | *A. thaliana* | AT1G54095 | 0 | 0 | 2 | 0 | 0 | 0 | 0 | 0 | 0 | 0 | 0 | 0 | 0 |
| | | AT1G72510 | 0 | 2 | 0 | 4 | 0 | 0 | 0 | 0 | 0 | 0 | 0 | 0 | 0 |
| | *P. trichocarpa* | POPTR_0001s16710 | 3 | 2 | 2 | 4 | 4 | 0 | 0 | 0 | 0 | 0 | 0 | 0 | 0 |
| | | POPTR_0003s06580 | 3 | 3 | 4 | 4 | 3 | 0 | 1 | 0 | 0 | 0 | 0 | 0 | 0 |
| | | POPTR_0006s23570 | 3 | 1 | 2 | 4 | 0 | 0 | 0 | 0 | 0 | 0 | 0 | 0 | 0 |
| HG58 | *A. thaliana* | AT2G24530 | 0 | 0 | 2 | 3 | 0 | 0 | 0 | 0 | 0 | 0 | 0 | 0 | 0 |
| | *V. vinifera* | VIT_04s0008g02410 | 0 | 1 | 3 | 4 | 0 | 0 | 0 | 0 | 0 | 0 | 0 | 0 | 0 |
| HG59 | *A. thaliana* | AT2G35940 | 0 | 3 | 1 | 6 | 1 | 0 | 0 | 0 | 0 | 0 | 0 | 0 | 0 |
| | *P. trichocarpa* | POPTR_0006s21950 | 1 | 3 | 3 | 5 | 1 | 0 | 0 | 0 | 0 | 0 | 0 | 0 | 0 |
| | | POPTR_0016s07040 | 1 | 4 | 2 | 5 | 2 | 0 | 0 | 0 | 0 | 0 | 0 | 0 | 0 |
| HG60 | *A. thaliana* | AT3G12570 | 0 | 1 | 0 | 2 | 0 | 0 | 0 | 0 | 0 | 0 | 0 | 0 | 0 |
| | *P. trichocarpa* | POPTR_0008s05420 | 0 | 0 | 2 | 1 | 0 | 0 | 0 | 0 | 0 | 0 | 0 | 0 | 0 |
| | | POPTR_0010s21340 | 0 | 0 | 2 | 1 | 0 | 0 | 0 | 0 | 0 | 0 | 0 | 0 | 0 |
| HG61 | *A. thaliana* | AT5G55600 | 0 | 0 | 2 | 2 | 0 | 0 | 0 | 0 | 0 | 0 | 0 | 0 | 0 |
| | *P. trichocarpa* | POPTR_0001s37450 | 0 | 0 | 3 | 1 | 0 | 0 | 0 | 0 | 0 | 0 | 0 | 0 | 0 |
| HG62 | *S. lycopersicum* | Solyc04g005430.2 | 2 | 1 | 2 | 3 | 0 | 0 | 0 | 0 | 0 | 0 | 0 | 0 | 0 |
| | | Solyc05g012450.2 | 2 | 3 | 1 | 3 | 0 | 0 | 0 | 0 | 0 | 0 | 0 | 0 | 0 |
| | *P. trichocarpa* | POPTR_0008s09440 | 1 | 0 | 2 | 3 | 0 | 0 | 0 | 0 | 0 | 0 | 0 | 0 | 0 |
| HG63 | *S. lycopersicum* | Solyc04g071860.2 | 0 | 5 | 2 | 5 | 4 | 0 | 1 | 0 | 0 | 0 | 0 | 0 | 0 |
| | *V. vinifera* | VIT_07s0031g01960 | 2 | 6 | 2 | 6 | 4 | 0 | 3 | 0 | 0 | 0 | 0 | 0 | 0 |
| HG64 | *P. trichocarpa* | POPTR_0002s08640 | 0 | 0 | 3 | 5 | 2 | 0 | 0 | 0 | 0 | 0 | 0 | 0 | 0 |
| | *V. vinifera* | VIT_18s0001g03290 | 0 | 1 | 3 | 6 | 2 | 0 | 0 | 0 | 0 | 0 | 0 | 0 | 0 |
| HG65 | *P. trichocarpa* | POPTR_0002s09080 | 0 | 1 | 1 | 1 | 0 | 0 | 0 | 0 | 0 | 0 | 0 | 0 | 0 |
| | *V. vinifera* | VIT_12s0034g02380 | 0 | 1 | 1 | 1 | 0 | 0 | 0 | 0 | 0 | 0 | 0 | 0 | 0 |
| | | VIT_18s0001g04340 | 0 | 0 | 1 | 2 | 0 | 0 | 0 | 0 | 0 | 0 | 0 | 0 | 0 |
| HG66 | *P. trichocarpa* | POPTR_0004s20000 | 3 | 3 | 3 | 3 | 3 | 1 | 1 | 1 | 1 | 0 | 0 | 0 | 0 |
| | | POPTR_0009s15140 | 3 | 3 | 3 | 3 | 3 | 0 | 1 | 1 | 0 | 0 | 0 | 0 | 0 |
| | *V. vinifera* | VIT_06s0061g01250 | 3 | 3 | 3 | 4 | 2 | 0 | 1 | 1 | 0 | 0 | 0 | 0 | 0 |
| HG67 | *P. trichocarpa* | POPTR_0006s23550 | 1 | 1 | 4 | 5 | 0 | 0 | 0 | 0 | 0 | 0 | 0 | 0 | 0 |
| | | POPTR_0018s08370 | 0 | 1 | 4 | 5 | 0 | 0 | 0 | 0 | 0 | 0 | 0 | 0 | 0 |
| | *V. vinifera* | VIT_04s0008g02100 | 4 | 5 | 0 | 0 | 3 | 0 | 2 | 0 | 0 | 0 | 0 | 0 | 0 |
| HG68 | *P. trichocarpa* | POPTR_0011s08940 | 0 | 1 | 0 | 3 | 0 | 0 | 0 | 0 | 0 | 0 | 0 | 0 | 0 |
| | *V. vinifera* | VIT_19s0090g01000 | 3 | 1 | 0 | 2 | 0 | 0 | 0 | 0 | 0 | 0 | 0 | 0 | 0 |
| HG69 | *P. trichocarpa* | POPTR_0013s05200 | 0 | 3 | 1 | 2 | 2 | 0 | 0 | 0 | 1 | 0 | 0 | 0 | 0 |
| | *V. vinifera* | VIT_08s0032g00280 | 1 | 3 | 2 | 3 | 2 | 0 | 0 | 0 | 2 | 0 | 1 | 0 | 0 |
| HG70 | *A. thaliana* | AT3G52490 | 0 | 2 | 1 | 3 | 2 | 0 | 0 | 0 | 0 | 0 | 0 | 0 | 0 |
| HG71 | *A. thaliana* | AT3G61970 | 0 | 0 | 2 | 1 | 1 | 0 | 0 | 0 | 0 | 0 | 0 | 0 | 0 |
| HG72.1 | *A. thaliana* | AT4G34610 | 1 | 0 | 1 | 1 | 0 | 0 | 0 | 0 | 0 | 0 | 0 | 0 | 0 |
| HG72.2 | *V. vinifera* | VIT_03s0038g00050 | 2 | 0 | 1 | 2 | 0 | 0 | 0 | 0 | 0 | 0 | 0 | 0 | 0 |
| HG73 | *A. thaliana* | AT5G53660 | 1 | 0 | 3 | 3 | 2 | 0 | 0 | 0 | 0 | 0 | 0 | 0 | 0 |
| HG74 | *O. sativa* | LOC_Os04g36058 | 0 | 0 | 0 | 0 | 1 | 2 | 1 | 1 | 0 | 0 | 0 | 0 | 0 |
| HG75 | *O. sativa* | LOC_Os11g16280 | 2 | 2 | 2 | 4 | 3 | 2 | 3 | 0 | 1 | 0 | 0 | 0 | 0 |

| | | | | | | | | | | | | | | | |
|---|---|---|---|---|---|---|---|---|---|---|---|---|---|---|---|
| HG76 | *S. lycopersicum* | Solyc02g091660.2 | 0 | 2 | 0 | 0 | 0 | 0 | 0 | 0 | 0 | 0 | 0 | 0 | 0 |
| HG77 | *S. lycopersicum* | Solyc03g118620.2 | 1 | 2 | 1 | 2 | 2 | 0 | 0 | 0 | 0 | 0 | 0 | 0 | 0 |
| HG78 | *S. lycopersicum* | Solyc04g079900.2 | 0 | 0 | 1 | 1 | 0 | 0 | 0 | 0 | 0 | 0 | 0 | 0 | 0 |
| HG79 | *S. lycopersicum* | Solyc08g082320.2 | 0 | 0 | 0 | 3 | 0 | 0 | 0 | 0 | 0 | 0 | 0 | 0 | 0 |
| HG80 | *P. trichocarpa* | POPTR_0001s30040 | 3 | 4 | 3 | 1 | 3 | 0 | 0 | 0 | 0 | 0 | 0 | 0 | 0 |
| | | POPTR_0004s21550 | 3 | 2 | 3 | 1 | 3 | 0 | 0 | 0 | 0 | 0 | 0 | 0 | 0 |
| | | POPTR_0009s16830 | 3 | 4 | 1 | 1 | 2 | 0 | 0 | 0 | 0 | 0 | 0 | 0 | 0 |
| HG81 | *P. trichocarpa* | POPTR_0001s36900 | 3 | 6 | 4 | 5 | 4 | 0 | 0 | 0 | 0 | 0 | 0 | 0 | 0 |
| | | POPTR_0011s09450 | 1 | 3 | 4 | 4 | 1 | 0 | 0 | 0 | 0 | 0 | 0 | 0 | 0 |
| HG82 | *P. trichocarpa* | POPTR_0001s38480 | 2 | 1 | 2 | 2 | 0 | 0 | 0 | 0 | 0 | 0 | 0 | 0 | 0 |
| HG83 | *P. trichocarpa* | POPTR_0001s41040 | 0 | 0 | 2 | 0 | 0 | 0 | 0 | 0 | 0 | 0 | 0 | 0 | 0 |
| HG84 | *P. trichocarpa* | POPTR_0002s10310 | 0 | 0 | 1 | 2 | 0 | 0 | 0 | 0 | 0 | 0 | 0 | 0 | 0 |
| HG85 | *P. trichocarpa* | POPTR_0002s12870 | 0 | 0 | 2 | 4 | 2 | 0 | 0 | 0 | 0 | 0 | 0 | 0 | 0 |
| HG86 | *P. trichocarpa* | POPTR_0003s08760 | 1 | 1 | 2 | 3 | 0 | 0 | 0 | 0 | 0 | 0 | 0 | 0 | 0 |
| HG87 | *P. trichocarpa* | POPTR_0004s03900 | 1 | 2 | 3 | 4 | 3 | 0 | 0 | 0 | 0 | 0 | 0 | 0 | 0 |
| | | POPTR_0011s04730 | 1 | 2 | 4 | 4 | 3 | 0 | 0 | 0 | 0 | 0 | 0 | 0 | 0 |
| HG88 | *P. trichocarpa* | POPTR_0004s05670 | 0 | 0 | 1 | 2 | 0 | 0 | 0 | 0 | 0 | 0 | 0 | 0 | 0 |
| | | POPTR_0011s07260 | 0 | 0 | 1 | 2 | 0 | 0 | 0 | 0 | 0 | 0 | 0 | 0 | 0 |
| HG89 | *P. trichocarpa* | POPTR_0004s19490 | 0 | 0 | 2 | 0 | 0 | 0 | 0 | 0 | 0 | 0 | 0 | 0 | 0 |
| | | POPTR_0009s14620 | 0 | 0 | 2 | 0 | 0 | 0 | 0 | 0 | 0 | 0 | 0 | 0 | 0 |
| HG90 | *P. trichocarpa* | POPTR_0005s02640 | 0 | 1 | 2 | 2 | 1 | 0 | 0 | 0 | 0 | 0 | 0 | 0 | 0 |
| HG91 | *P. trichocarpa* | POPTR_0005s07290 | 0 | 1 | 1 | 3 | 0 | 0 | 0 | 0 | 0 | 0 | 0 | 0 | 0 |
| | | POPTR_0007s05010 | 0 | 0 | 1 | 3 | 0 | 0 | 0 | 0 | 0 | 0 | 0 | 0 | 0 |
| HG92.1 | *P. trichocarpa* | POPTR_0005s11030 | 3 | 2 | 2 | 4 | 2 | 0 | 0 | 0 | 0 | 0 | 0 | 0 | 0 |
| | | POPTR_0007s09220 | 2 | 1 | 2 | 4 | 2 | 0 | 0 | 0 | 0 | 0 | 0 | 0 | 0 |
| HG92.2 | *V. vinifera* | VIT_03s0038g02650 | 3 | 4 | 2 | 6 | 2 | 0 | 1 | 0 | 0 | 0 | 0 | 0 | 0 |
| HG93 | *P. trichocarpa* | POPTR_0005s11480 | 0 | 0 | 1 | 0 | 2 | 0 | 0 | 0 | 0 | 0 | 0 | 0 | 0 |
| HG94 | *P. trichocarpa* | POPTR_0005s27810 | 0 | 0 | 1 | 2 | 0 | 0 | 0 | 0 | 0 | 0 | 0 | 0 | 0 |
| HG95 | *P. trichocarpa* | POPTR_0006s07660 | 1 | 1 | 1 | 4 | 2 | 1 | 1 | 0 | 0 | 0 | 0 | 0 | 0 |
| HG96 | *P. trichocarpa* | POPTR_0006s13860 | 0 | 2 | 3 | 2 | 1 | 0 | 0 | 0 | 0 | 0 | 0 | 0 | 0 |
| | | POPTR_0018s07360 | 0 | 3 | 3 | 3 | 2 | 0 | 0 | 0 | 0 | 0 | 0 | 0 | 0 |
| HG97 | *P. trichocarpa* | POPTR_0006s14740 | 2 | 3 | 1 | 2 | 2 | 0 | 0 | 0 | 0 | 0 | 0 | 0 | 0 |
| HG98 | *P. trichocarpa* | POPTR_0006s22890 | 1 | 2 | 2 | 4 | 3 | 2 | 0 | 1 | 0 | 0 | 0 | 0 | 0 |
| HG99 | *P. trichocarpa* | POPTR_0006s27080 | 0 | 0 | 1 | 0 | 1 | 0 | 0 | 0 | 0 | 0 | 0 | 0 | 0 |
| HG100 | *P. trichocarpa* | POPTR_0008s10350 | 0 | 0 | 2 | 1 | 0 | 0 | 0 | 0 | 0 | 0 | 0 | 0 | 0 |
| HG101.1 | *P. trichocarpa* | POPTR_0008s15440 | 0 | 0 | 0 | 2 | 0 | 0 | 0 | 0 | 0 | 0 | 0 | 0 | 0 |
| | | POPTR_0010s09580 | 0 | 0 | 0 | 2 | 0 | 0 | 0 | 0 | 0 | 0 | 0 | 0 | 0 |
| HG101.2 | *V. vinifera* | VIT_05s0020g03670 | 0 | 0 | 0 | 3 | 0 | 0 | 0 | 0 | 0 | 0 | 0 | 0 | 0 |
| HG102 | *P. trichocarpa* | POPTR_0008s19310 | 0 | 1 | 0 | 3 | 2 | 0 | 0 | 0 | 0 | 0 | 0 | 0 | 0 |
| HG103 | *P. trichocarpa* | POPTR_0009s15460 | 2 | 2 | 1 | 0 | 3 | 0 | 1 | 0 | 0 | 0 | 0 | 0 | 0 |
| HG104 | *P. trichocarpa* | POPTR_0010s01710 | 0 | 1 | 2 | 0 | 0 | 0 | 0 | 0 | 0 | 0 | 0 | 0 | 0 |
| HG105 | *P. trichocarpa* | POPTR_0010s09600 | 2 | 1 | 1 | 3 | 0 | 0 | 0 | 0 | 0 | 0 | 0 | 0 | 0 |
| HG106 | *P. trichocarpa* | POPTR_0010s11700 | 3 | 5 | 1 | 4 | 4 | 0 | 0 | 0 | 0 | 0 | 0 | 0 | 0 |
| HG107 | *P. trichocarpa* | POPTR_0013s08000 | 0 | 0 | 2 | 4 | 0 | 0 | 0 | 0 | 0 | 0 | 0 | 0 | 0 |
| HG108 | *P. trichocarpa* | POPTR_0013s13670 | 0 | 0 | 2 | 1 | 0 | 0 | 0 | 0 | 0 | 0 | 0 | 0 | 0 |

| | | | | | | | | | | | | | | | |
|---|---|---|---|---|---|---|---|---|---|---|---|---|---|---|---|
| HG109 | *P. trichocarpa* | POPTR_0014s02470 | 0 | 0 | 1 | 1 | 1 | 0 | 1 | 0 | 0 | 0 | 0 | 0 | 0 |
| HG110 | *P. trichocarpa* | POPTR_0014s03370 | 0 | 2 | 1 | 3 | 0 | 0 | 0 | 0 | 0 | 0 | 0 | 0 | 0 |
| HG111 | *P. trichocarpa* | POPTR_0014s09590 | 0 | 0 | 3 | 4 | 1 | 0 | 0 | 0 | 0 | 0 | 0 | 0 | 0 |
| HG112 | *P. trichocarpa* | POPTR_0014s15970 | 0 | 2 | 0 | 3 | 2 | 0 | 0 | 0 | 0 | 0 | 0 | 0 | 0 |
| HG113 | *P. trichocarpa* | POPTR_0019s11580 | 0 | 2 | 1 | 2 | 1 | 0 | 0 | 0 | 0 | 0 | 0 | 0 | 0 |
| HG114 | *P. trichocarpa* | POPTR_0019s13560 | 0 | 0 | 2 | 0 | 2 | 1 | 0 | 0 | 0 | 0 | 0 | 0 | 0 |
| HG115 | *V. vinifera* | VIT_01s0010g03770 | 0 | 0 | 0 | 1 | 1 | 0 | 0 | 0 | 0 | 0 | 0 | 0 | 0 |
| HG116 | *V. vinifera* | VIT_01s0137g00230 | 0 | 0 | 2 | 3 | 0 | 0 | 0 | 0 | 0 | 0 | 0 | 0 | 0 |
| HG117 | *V. vinifera* | VIT_01s0146g00180 | 4 | 3 | 4 | 5 | 3 | 2 | 3 | 2 | 2 | 0 | 0 | 0 | 0 |
| HG118 | *V. vinifera* | VIT_04s0008g01420 | 0 | 0 | 0 | 3 | 0 | 0 | 0 | 0 | 0 | 0 | 0 | 0 | 0 |
| HG119 | *V. vinifera* | VIT_04s0008g04480 | 1 | 0 | 2 | 0 | 2 | 0 | 0 | 0 | 0 | 0 | 0 | 0 | 0 |
| HG120 | *V. vinifera* | VIT_05s0020g00460 | 0 | 1 | 0 | 2 | 0 | 1 | 0 | 0 | 0 | 0 | 0 | 0 | 0 |
| HG121 | *V. vinifera* | VIT_05s0029g00470 | 3 | 2 | 3 | 5 | 4 | 0 | 0 | 0 | 0 | 0 | 0 | 0 | 0 |
| HG122 | *V. vinifera* | VIT_06s0004g02980 | 1 | 3 | 2 | 3 | 0 | 0 | 0 | 0 | 0 | 0 | 0 | 0 | 0 |
| HG123 | *V. vinifera* | VIT_11s0016g01790 | 2 | 3 | 1 | 4 | 1 | 0 | 0 | 0 | 0 | 0 | 0 | 0 | 0 |
| HG124 | *V. vinifera* | VIT_11s0016g02880 | 0 | 0 | 1 | 2 | 0 | 0 | 0 | 0 | 0 | 0 | 0 | 0 | 0 |
| HG125 | *V. vinifera* | VIT_14s0066g00370 | 1 | 0 | 0 | 4 | 1 | 0 | 0 | 0 | 0 | 0 | 0 | 0 | 0 |
| HG126 | *V. vinifera* | VIT_14s0108g00970 | 0 | 0 | 0 | 4 | 0 | 0 | 0 | 0 | 0 | 0 | 0 | 0 | 0 |
| HG127.1 | *V. vinifera* | VIT_18s0001g08100 | 1 | 0 | 1 | 3 | 0 | 0 | 0 | 0 | 0 | 0 | 0 | 0 | 0 |
| HG127.2 | *V. vinifera* | VIT_04s0008g04480 | 0 | 0 | 0 | 4 | 0 | 0 | 0 | 0 | 0 | 0 | 0 | 0 | 0 |
| HG128 | *V. vinifera* | VIT_18s0001g10680 | 0 | 0 | 1 | 1 | 0 | 0 | 0 | 0 | 0 | 0 | 0 | 0 | 0 |
| HG129 | *V. vinifera* | VIT_18s0001g13200 | 0 | 1 | 1 | 2 | 1 | 0 | 0 | 0 | 0 | 0 | 0 | 0 | 0 |
| HG130 | *V. vinifera* | VIT_18s0157g00020 | 0 | 0 | 3 | 4 | 1 | 1 | 0 | 0 | 0 | 0 | 0 | 0 | 0 |

** Numbers of orders from which uORF-tBLASTn and mORF-tBLASTn hits were extracted are indicated. The presence of the [illegible]
hits in each of the 13 categories is indicated by highlighting cells in red.

* Orders included in lower taxonomic categories were excluded.

### Supplementary Table S3

Supplementary Table S3. Plasmids used in this study and primers used for plasmid construction

| Plasmid | Construct | Primer | |
|---|---|---|---|
| | | Forward | Reverse |
| pNH92 | *35S::HG46(WT):FLuc* | HG46 for | HG46 rev |
| pNH93 | *35S::HG55(WT):FLuc* | HG55 for | HG55 rev |
| pNH94 | *35S::HG57(WT):FLuc* | HG57 for | HG57 rev |
| pNH95 | *35S::HG65(WT):FLuc* | HG65 for | HG65 rev |
| pNH96 | *35S::HG66(WT):FLuc* | HG66 for | HG66 rev |
| pNH97 | *35S::HG80(WT):FLuc* | HG80 for | HG80 rev |
| pNH98 | *35S::HG81(WT):FLuc* | HG81 for | HG81 rev |
| pNH99 | *35S::HG87(WT):FLuc* | HG87 for | HG87 rev |
| pNH100 | *35S::HG88(WT):FLuc* | HG88 for | HG88 rev |
| pNH101 | *35S::HG103(WT):FLuc* | HG103 for | HG103 rev |
| pNH102 | *35S::HG107(WT):FLuc* | 35S XbaI SLiCE-F | FLUC SalI SLiCE-R |
| pNH103 | *35S::HG46(fs):FLuc* | HG46 fs for1 | HG46 fs rev1 |
| | | HG46 fs for2 | HG46 fs rev2 |
| pNH104 | *35S::HG55(fs):FLuc* | HG55 fs for1 | HG55 fs rev1 |
| pNH105 | *35S::HG57(fs):FLuc* | HG57 fs for1 | HG57 fs rev1 |
| | | HG57 fs for2 | HG57 fs rev2 |
| pNH106 | *35S::HG063(fs):FLuc* | HG65 fs for1 | HG65 fs rev1 |
| | | HG65 fs for2 | HG65 fs rev2 |
| pNH107 | *35S::HG066(fs):FLuc* | HG66 fs for1 | HG66 fs rev1 |
| | | HG66 fs for2 | HG66 fs rev2 |
| pNH108 | *35S::HG080(fs):FLuc* | HG80 fsf for1 | HG80 fs rev1 |
| | | HG80 fs for2 | HG80 fs rev2 |
| pNH109 | *35S::HG81(fs):FLuc* | HG81 fs for1 | HG81 fs rev1 |
| pNH110 | *35S::HG87(fs):FLuc* | HG87 fs for1 | HG87 fs rev1 |
| | | HG87 fs for2 | HG87 fs rev2 |
| pNH111 | *35S::HG88(fs):FLuc* | HG88 fs for1 | HG88 fs rev1 |
| | | HG88 fs for2 | HG88 fs rev2 |
| pNH113 | *35S::HG103(fs):FLuc* | HG103 fs for1 | HG103 fs rev1 |
| | | HG103 fs for2 | HG103 fs rev2 |
| pNH112 | *35S::HG107(fs):FLuc* | HG107 fs for | HG107 fs rev |

### Supplementary Table S4

Supplymentary Table S4. Resourse numbers of *Poplus nigra* full-length cDNA clones used for cloning of 5′-UTRs.

| *Poplus trichocarpa* gene name | Homology Group (HG) | *Poplus nigra* fulle-length cDNA resource number |
|---|---|---|
| POPTR_0006s24280 | HG46 | pds25559 |
| POPTR_0014s10960 | HG55 | pds10965 |
| POPTR_0006s23570 | HG57 | pds14390 |
| POPTR_0002s09080 | HG65 | pds12940 |
| POPTR_0009s15140 | HG66 | pds13862 |
| POPTR_0009s16830 | HG80 | pds15817 |
| POPTR_0001s36900 | HG81 | pds28294 |
| POPTR_0011s04730 | HG87 | pds26157 |
| POPTR_0004s05670 | HG88 | pds14623 |
| POPTR_0009s15460 | HG103 | pds23234 |

### Supplementary Table S5

Supplementary Table S5. Primers used in this study

| Name | Primer sequence |
|---|---|
| HG46 for | 5´-CTCTCTAGAAGAAGCCAAAAAAAAAAGAAAAGATACA-3´ |
| HG46 rev | 5´-TCTGTCGACCTCATTTCCCCAAAATTCCAATTTC-3´ |
| HG55 for | 5´-CACGGGGGACTCTAGACTCTTTCTATGTACTATACCTCTCACC-3´ |
| HG55 rev | 5´-GTCTTCCATGGTCGACTCCATAACGAGTACCCAAAAGATC-3´ |
| HG57 for | 5´-CTCTCTAGAAACAGATACAACACAATCTCCATAC-3´ |
| HG57 rev | 5´-TCTGTCGACGCCATTCGTTTCCCTTAAATC-3´ |
| HG65 for | 5´-CTCTCTAGAACTGGTATCTCTCTCTCCCTTTTCTA-3´ |
| HG65 rev | 5´-TCTGTCGACCCCATCTTAAAACCCTAAAACA-3´ |
| HG66 for | 5´-CTCTCTAGAAGCCCAGAATGTCCATCTCCTA-3´ |
| HG66 rev | 5´-TCTGTCGACGGCATAATTGTAAGCGACAGTA-3´ |
| HG80 for | 5´-CTCTCTAGAGAGATTATAATCAAGGTGGTCAATTGA-3´ |
| HG80 rev | 5´-TCTGTCGACTGCATTGAAAATTTGTATCCAACAAC-3´ |
| HG81 for | 5´-CTCTCTAGAAAATGTCCTCTTCATTGATTGAGA-3´ |
| HG81 rev | 5´-TCTGTCGACCTCATTAAGACTGAGAGAGGGGGA-3´ |
| HG87 for | 5´-CACGGGGGACTCTAGAAAGGAAAGGGTGCTGAGTATATCA-3´ |
| HG87 rev | 5´-GTCTTCCATGGTCGACCCCATCTCTTAAAACACTACAAGA-3´ |
| HG88 for | 5´-CTCTCTAGATGATTAAACAATTTCGAAGACTTTCC-3´ |
| HG88 rev | 5´-TCTGTCGACGGCATGATGCAAAGAATGTAGC-3´ |
| HG103 for | 5´-CTCTCTAGAGACAACCCTCTCCAAACTC-3´ |
| HG103 rev | 5´-TCTGTCGACTCCATCTCTAATAAAAATAAAAATTGG-3´ |
| 35S XbaI SLiCE-F | 5´-AGAACACGGGGGACTCTAGA-3´ |
| FLUC SalI SLiCE-R | 5´-GGCGTCTTCCATGGTCGA-3´ |
| HG46 fs for1 | 5´-TGTGAACTTTCCCATTCGCTTCTGGGTG-3´ |
| HG46 fs rev1 | 5´-CACCCAGAAGCGAATGGGAAAGTTCACA-3´ |
| HG46 fs for2 | 5´-TCAAGCTTCAGAGTGGGATAATTACTATTATCAC-3´ |
| HG46 fs rev2 | 5´-GTGATAATAGTAATTATCCCACTCTGAAGCTTGA-3´ |
| HG55 fs for1 | 5´-TGTACTTTTGGGGAAGAAGCGACCGCTGATTAG-3´ |
| HG55 fs rev1 | 5´-GCTTCTTCCCCAAAAGTACAAAAAAACCAACCCACACACC-3´ |
| HG57 fs for1 | 5´-GTTTGGAGTTCCTTTGTTCAAGGA-3´ |
| HG57 fs rev1 | 5´-TCCTTGAACAAAGGAACTCCAAAC-3´ |
| HG57 fs for2 | 5´-GGAGAAAGAGCAGATAGATATTAGCTCA-3´ |
| HG57 fs rev2 | 5´-TGAGCTAATATCTATCTGCTCTTTCTCC-3´ |
| HG65 fs for1 | 5´-TAACACATTAAGATTCGTTACTCCCG-3´ |
| HG65 fs rev1 | 5´-CGGGAGTAACGAATCTTAATGTGTTA-3´ |
| HG65 fs for2 | 5´-TTGTTTCTACTTCTTACCCCTGAAA-3´ |
| HG65 fs rev2 | 5´-TTTCAGGGGTAAGAAGTAGAAACAA-3´ |
| HG66 fs for1 | 5´-AACGCTCCTCTCTGCTCTCGGTTTC-3´ |
| HG66 fs rev1 | 5´-GAAACCGAGAGCAGAGAGGAGCGTTG-3´ |
| HG66 fs for2 | 5´-CTACCGCCGTCTTGAGGTAACC-3´ |
| HG66 fs rev2 | 5´-GGTTACCTCAAGACGGCGGTAG-3´ |
| HG80 fs for1 | 5´-CATATTGGGTTCTTCGGATACAAGC-3´ |
| HG80 fs rev1 | 5´-GCTTGTATCCGAAGAACCCAATATG-3´ |
| HG80 fs for2 | 5´-TCTTTTGTGTTTCGGTCGTGATAAAAG-3´ |
| HG80 fs rev2 | 5´-CTTTTATCACGACCGAAACACAAAAGA-3´ |
| HG81 fs for1 | 5´-CGTGTTTTGTAGCAAAACCGGTTTCTGGTTGAGGATTTA-3´ |
| HG81 fs rev1 | 5´-CGGTTTTGCTACAAAACACGATCGTTTCAGCCTCA-3´ |
| HG87 fs for1 | 5´-TCACTTGAGCACATATTTCACTGTTC-3´ |
| HG87 fs rev1 | 5´-GAACAGTGAAATATGTGCTCAAGTGA-3´ |
| HG87 fs for2 | 5´-CCAAGGTCCAAGGCTAGCCAGTTA-3´ |
| HG87 fs rev2 | 5´-TAACTGGCTAGCCTTGGACCTTGG-3´ |
| HG88 fs for1 | 5´-TAATGGCAATGCATTATATTGCTGTC-3´ |
| HG88 fs rev1 | 5´-GACAGCAATATAATGCATTGCCATTA-3´ |
| HG88 fs for2 | 5´-TGCAGAGAGAGACAACTTGCTTTTG-3´ |
| HG88 fs rev2 | 5´-CAAAAGCAAGTTGTCTCTCTCTGCA-3´ |

### Supplementary Table S5 (continued)

| Name | Primer sequence |
|---|---|
| HG103 fs for1 | 5´-CAAGAACAACAACACAAGAAAATCCA-3´ |
| HG103 fs rev1 | 5´-TGGATTTTCTTGTGTTGTTGTTCTTG-3´ |
| HG103 fs for2 | 5´-GAACCTAACAGACCATAAAAAATCTC-3´ |
| HG103 fs rev2 | 5´-GAGATTTTTTATGGTCTGTTAGGTTC-3´ |
| HG107 fs for | 5´-TCCGGTTCATCCGCTTCTCTCTCTTCTTCTCTGGAGGCCTGATTGA-3´ |
| HG107 fs rev | 5´-GAGAAGCGGATGAACCGGAAAGGGTGTAAGTGGA-3´ |

The recognition sites of restriction enzymes are underlined.
